# Roles of RodZ and Class A PBP1b in the Assembly and Regulation of the Peripheral Peptidoglycan Elongasome in Ovoid-Shaped Cells of *Streptococcus pneumoniae* D39

**DOI:** 10.1101/2022.06.01.494439

**Authors:** Melissa M. Lamanna, Irfan Manzoor, Merrin Joseph, Ziyun A. Ye, Mattia Benedet, Alessia Zanardi, Zhongqing Ren, Xindan Wang, Orietta Massidda, Ho-Ching T. Tsui, Malcolm E. Winkler

**Author notes:** Co-corresponding authors: Malcolm E. Winkler Ho-Ching T. Tsui.

## Abstract

RodZ of rod-shaped bacteria functions to link MreB filaments to the Rod peptidoglycan (PG) synthase complex that moves circumferentially perpendicular to the long cell axis, creating hoop-like sidewall PG. Ovoid-shaped bacteria, such as *Streptococcus pneumoniae* (pneumococcus; *Spn*) that lack MreB, use a different modality for peripheral PG elongation that emanates from the midcell of dividing cells. Yet, *S. pneumoniae* encodes a RodZ homolog similar to RodZ in rod-shaped bacteria. We show here that the helix-turn-helix and transmembrane domains of RodZ(*Spn*) are essential for growth at 37°C. Δ*rodZ* mutations are suppressed by Δ*pbp1a*, *mpgA*(Y488D), and Δ*khpA* mutations that suppress Δ*mreC*, but not Δ*cozE*. Consistent with a role in PG elongation, RodZ(*Spn*) co-localizes with MreC and aPBP1a throughout the cell cycle and forms complexes and interacts with PG elongasome proteins and regulators. Depletion of RodZ(*Spn*) results in aberrantly shaped, non-growing cells and mislocalization of elongasome proteins MreC, PBP2b, and RodA. Moreover, Tn-seq reveals that RodZ(*Spn*), but not MreCD(*Spn*), displays a specific synthetic-viable genetic relationship with aPBP1b, whose function is unknown. We conclude that RodZ(*Spn*) acts as a scaffolding protein required for elongasome assembly and function and that aPBP1b, like aPBP1a, plays a role in elongasome regulation and possibly peripheral PG synthesis.

**Graphical Summary:** 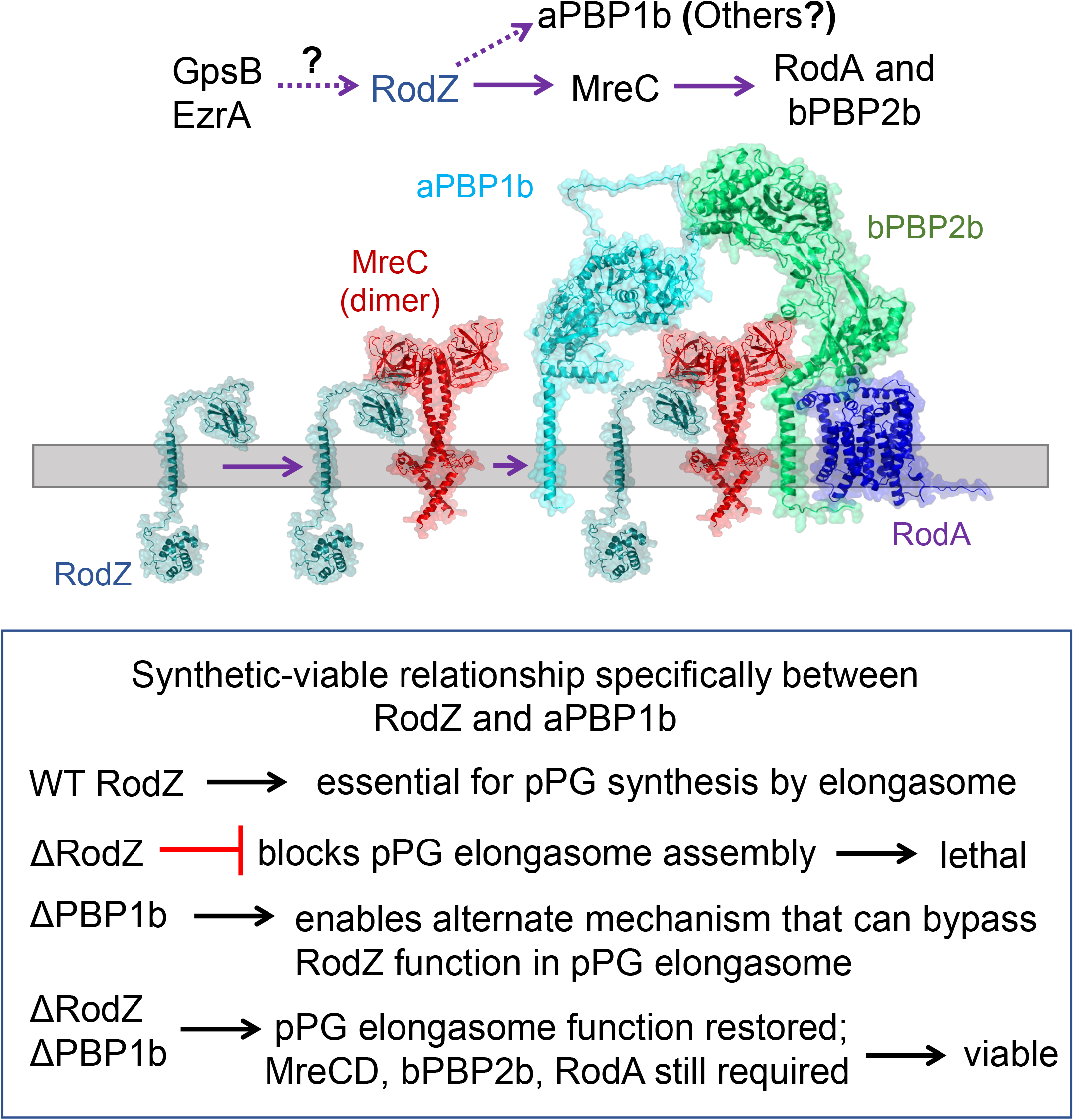

## 1 INTRODUCTION

The peptidoglycan (PG) mesh, which consists of peptide-crosslinked glycan chains, determines the shape of eubacteria, contributing to their colonization and survival in different environmental niches (Daitch & Goley, 2020, Egan *et al*., 2020, Kumar *et al*., 2022, Rohs & Bernhardt, 2021, Young, 2006). PG also protects bacteria from turgor pressure and serves as a scaffold for the attachment of extracellular proteins and exopolysaccharide capsules and wall-teichoic acids of Gram-positive bacteria, which lack an outer membrane (Rajagopal & Walker, 2017, Vollmer *et al*., 2019). PG synthesis has been a major target for many classes of antibiotics, starting with the β-lactam penicillin (Bush & Bradford, 2016); however, resistance to cell-wall targeted antibiotics is now a serious, widespread health problem (CDC, 2019, Hakenbeck, 2014, Hakenbeck *et al*., 2012). Because of its extracellular location, absence in eukaryotic hosts, and many vulnerable enzymatic and regulatory steps, PG synthesis remains a leading target for the discovery and development of new classes of antibiotics (CDC, 2019, den Blaauwen *et al*., 2014, Lewis, 2020, Ling *et al*., 2015, Sham *et al*., 2012).

Formation of ovoid-shaped (ovococcal) bacteria (Zapun *et al*., 2008), such as the major respiratory pathogen *Streptococcus pneumoniae* (pneumococcus; *Spn*) (Weiser *et al*., 2018), requires two modes of PG synthesis (reviewed in (Briggs *et al*., 2021, Massidda *et al*., 2013, Perez *et al*., 2021a, Vollmer *et al*., 2019)). Septal PG (sPG) synthesis separates dividing pneumococcal cells at midcell into two daughter cells, whereas peripheral PG (pPG) synthesis is a form of sidewall PG synthesis that also emanates from the midcell division ring of dividing pneumococcal cells. All protein components for both modes of PG synthesis are initially organized by FtsZ, FtsA, and EzrA into a single ring at the equators of predivisional pneumococcal cells (Perez *et al*., 2019). In the course of division, the sPG synthesis machine moves with the constricting FtsZ ring at the leading edge of the closing septal annulus, separate from the pPG synthesis machine that remains at the outer edge of the septal disk (Briggs *et al*., 2021). This dual pattern of PG synthesis was recently visualized as two concentric midcell rings by high-resolution structured-illumination microscopy (3D-SIM) (Perez *et al*., 2021a) and direct stochastic optical reconstruction microscopy (dSTORM) (Trouve *et al*., 2021) microscopy of vertically oriented pneumococcal cells. At the start of division, pPG synthesis likely begins slightly before sPG synthesis, but throughout most of the cell cycle, sPG and pPG synthesis and PG remodeling at midcell are concurrent and highly coordinated (Briggs *et al*., 2021, Perez *et al*., 2021a, Trouve *et al*., 2021, Tsui *et al*., 2014, Wheeler *et al*., 2011).

Midcell localization of sPG and pPG synthesis in ovococci is fundamentally different in many ways from the patterns of sPG and sidewall PG synthesis used by rod-shaped bacteria (Rohs & Bernhardt, 2021). In *Bacillus subtilis*, which like *S. pneumoniae* is a low-GC Gram-positive bacterium, a wall of sPG is synthesized during septal closure without surface constriction between daughter cells that are later separated by PG hydrolases (Errington & Wu, 2017, Straume *et al*., 2021). Gram-positive coccoid bacteria, such as *Staphylococcus aureus*, also synthesize a septal cell wall between daughter cells, which are later separated by a rapid PG hydrolytic popping mechanism (Lund *et al*., 2018, Saraiva *et al*., 2020, Straume *et al*., 2021). In Gram-negative *Escherichia coli*, septal closure and cell separation are largely concurrent (Rohs & Bernhardt, 2021), similar to what is observed for *S. pneumoniae*. However, in *E. coli,* the regulation of sPG synthesis occurs by a different mechanism than in *S. pneumoniae*, which lacks FtsN-mediated activation of essential Class B PBP3 (FtsI) transpeptidase activity required for septal closure in *E. coli* (Briggs *et al*., 2021, Pichoff *et al*., 2019, Rohs & Bernhardt, 2021).

Early in cell division of rod-shaped bacteria, preseptal PG synthesis pushes sidewall PG outward from the Z-ring (Aaron *et al*., 2007, Pazos *et al*., 2018, van Teeseling, 2021), resembling the pPG synthesis that occurs throughout the pneumococcal cell cycle (Briggs *et al*., 2021). However, following preseptal PG synthesis, the Rod-complex elongasome, containing conditionally essential MreB and the sidewall PG synthase complex, assembles along the curved cylindrical body of rod-shaped cells (Bratton *et al*., 2018, Hussain *et al*., 2018, Morgenstein *et al*., 2015, Rohs & Bernhardt, 2021). MreB is an actin-like homolog that polymerizes into multiple, short, curved filaments along the cell membrane, perpendicular to the cell long axis, which is the direction of maximum negative Gaussian curvature (Bratton *et al*., 2018, Hussain *et al*., 2018, Morgenstein *et al*., 2015, Rohs & Bernhardt, 2021). MreB filaments are linked to the sidewall PG synthase complex by RodZ, whose cytoplasmic helix-turn-helix (HTH) interacts with MreB inside the cell (Ago & Shiomi, 2019, Bendezu *et al*., 2009, Morgenstein *et al*., 2015, van den Ent *et al*., 2010), and whose cytoplasmic and transmembrane (TM) domains (see Fig. 1 and S1) interact with the largely extracellular PG synthase proteins, including MreC and MreD (positive regulators) (Rohs & Bernhardt, 2021, Rohs *et al*., 2018, Rohs *et al*., 2021), RodA (SEDS glycosyl transferase) (Meeske *et al*., 2016, Sjodt *et al*., 2018, Sjodt *et al*., 2020), and an essential Class B penicillin-binding protein (bPBP2 transpeptidase in *E. coli*) (Rohs & Bernhardt, 2021, Sjodt *et al*., 2020). The interaction between RodZ and MreB can modulate the density and length of MreB filaments (Bratton *et al*., 2018, Colavin *et al*., 2018, Hussain *et al*., 2018). In *E. coli*, a limiting amount of bPBP2 seems to bind to PG and recruit the rest of the Rod elongasome (Ozbaykal *et al*., 2020). Movement of the assembled Rod elongasome is driven by sidewall PG synthesis itself, rather than by ATPase-dependent treadmilling of MreB (Rohs & Bernhardt, 2021), with the MreB filaments serving as curvature-sensing rudders (Hussain *et al*., 2018). Finally, in contrast to the parallel PG synthesis by the Rod elongasome, synthesis of sidewall PG by Class A PBPs appears to be largely non-ordered, possibly filling in or reinforcing gaps left by the Rod system (Cho *et al*., 2016, Dion *et al*., 2019, Lamanna & Maurelli, 2022, Rohs & Bernhardt, 2021).

**Figure 1.**
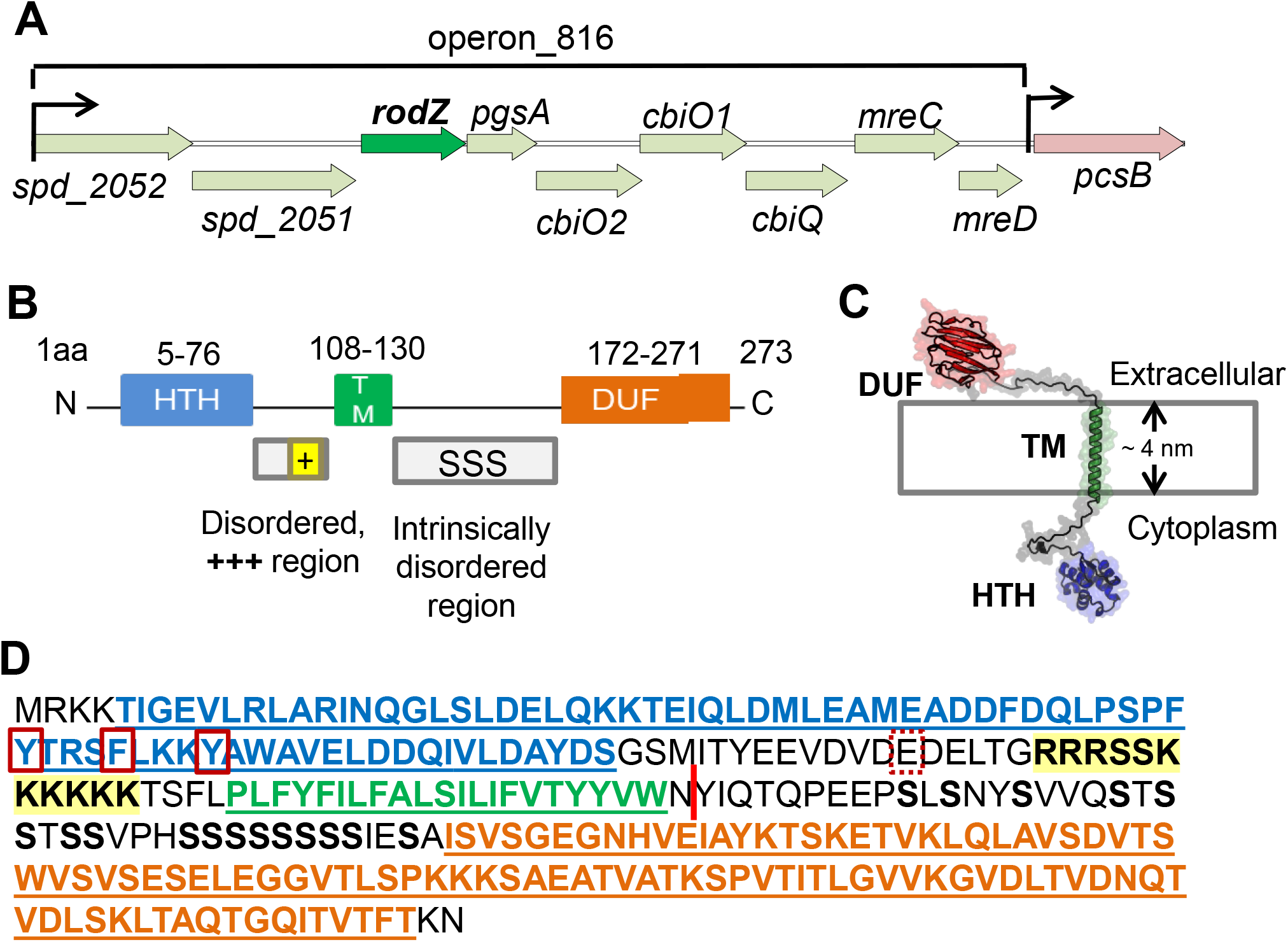
Location and domains of RodZ(*Spn*). (A) *rodZ* (*spd_2050*) is predicted to be a member of operon_816 in the *Spn* D39 chromosome (Slager *et al*., 2018). Operon_816 consists of *spd_2052* (putative Zinc protease), *spd_2051* (putative M16 family peptidase), *rodZ*, *pgsA* (CDP-diacylglycerol-glycerol-3-phosphate 3-phosphatidyltransferase), *cbiO2, cbiO1, cbiQ* (putative ATPase and transmembrane components of a cobalt ABC transporters), *mreC*, and *mreD*. (B) 2D protein structure of RodZ(*Spn*). Black line indicates residues that are not part of known domains. The intracellular helix-turn-helix (HTH) domain, transmembrane (TM), and extracellular domain of unknown function (DUF 4115) are depicted as blue, green, and orange, respectively, and intrinsically disordered regions are represented as gray boxes. The HTH and DUF domains are predicted to be in alpha helices and beta sheets, respectively, by AlphaFold2 (Jumper *et al*., 2021). TM domain was determined with TMHMM server. The positively charged juxta-membrane region of the intracellular linker is shown as a yellow box (+++). SSS symbolizes multiple serine residues in the extracellular linker. (C). Predicted 3D structure of RodZ(*Spn*) generated using the AlphaFold2 webserver. (D) Amino acid sequence of RodZ(*Spn*). Color coding is as described in (B), except that the multiple serine residues are bolded and the positive juxta-membrane is both bolded and highlighted in yellow. Y51 and F55 within Helix 4 of the HTH domain (red boxes) correspond to the positions of aromatic amino acids that interact with MreB in *E. coli* (see Fig. S1) (van den Ent *et al*., 2010). E89 (dotted box) corresponds to the position of S85 in RodZ(*Bsu*) that may be phosphorylated (Sun & Garner, 2020). The red bar between N131 and Y132 marks the first TA site in a TAT (Y132) codon with a Tn-Mariner insertion recovered by Tn-seq of the WT strain (Fig. 2). The junction of the Tn insertion creates a TAA stop codon, indicating that RodZ(M1-N131) is viable.

In contrast to rod-shaped bacteria, the pPG elongasome of *S. pneumoniae* is zonal and confined to the midcell of dividing pneumococcal cells. As division proceeds, the pPG elongasome locates to an outer ring of PG synthesis at the edge of the septal annulus (Briggs *et al*., 2021, Perez *et al*., 2021a, Trouve *et al*., 2021). Homologs of MreC, MreD, an essential Class B PBP (bPBP2b), and RodA have been associated with the pneumococcal pPG elongasome by genetic, physiological, and bacterial two-hybrid (B2H) experiments (Berg *et al*., 2013, Land & Winkler, 2011, Massidda *et al*., 2013, Philippe *et al*., 2014, Stamsas *et al*., 2017, Straume *et al*., 2017, Tsui *et al*., 2014, Zheng *et al*., 2017). In addition, Class A aPBP1a and CozE have been linked to pneumococcal pPG elongasome function through a synthetic-viable genetic relationship, in that Δ*pbp1a* or *cozE* depletion suppresses Δ*mreCD* (Fenton *et al*., 2016, Land & Winkler, 2011, Tsui *et al*., 2016). Complexes containing aPBP1a and pPG elongasome proteins have also been detected by co-immunoprecipitation (co-IP) and bacterial two-hybrid (B2H) assays (Fenton *et al*., 2016, Stamsas *et al*., 2017)}. Similarly, aPBP1a is synthetically viable with muramidase MpgA (formerly MltG(*Spn*), in that Δ*pbp1a* suppresses Δ*mpgA* (Taguchi *et al*., 2021, Tsui *et al*., 2016). Each of these proteins localizes to the midcell of dividing pneumococcal cells, consistent with the zonal mechanism of pPG elongation (Briggs *et al*., 2021, Land *et al*., 2013, Tsui *et al*., 2016, Tsui *et al*., 2014).

*S. pneumoniae*, like most non-rod-shaped bacteria, encodes a RodZ homolog (Fig. 1 and S1), despite the absence of an MreB homolog. The secondary structure of RodZ(*Spn*) is remarkably similar to RodZ homologs in rod-shaped bacteria (Fig. 1 and S1) (Ago & Shiomi, 2019, Alyahya *et al*., 2009, Bendezu *et al*., 2009, Shiomi *et al*., 2008). RodZ(*Spn*) contains a cytoplasmic N-terminal HTH domain of the XRE family, which contains five helices that often mediate protein interactions (Aravind *et al*., 2005). The HTH domain is connected to a TM domain by a disordered “juxtamembrane” domain, which is positively charged in RodZ(*Spn*) and RodZ(*Eco*) (Fig. 1D and S1) (Bendezu *et al*., 2009). Non-conserved Ser85 in this linker region of RodZ(*Bsu*) (Fig. 1 and S1B) has been reported to be phosphorylated in a preliminary report (Sun & Garner, 2020)}. The non-conserved extracellular linker of RodZ(*Spn*) contains a large number of repeated Ser residues and connects the TM domain to a domain of unknown function (DUF#4115) that is predicted by AlphaFold2 to fold into a beta-strand structure (Fig. 1C), similar to DUF determined for RodZ(*Bsu*) (Ago & Shiomi, 2019). *rodZ* is essential or conditionally essential in *E. coli*, *B. subtilis*, and *Caulobacter crescentus* (Alyahya *et al*., 2009, Bendezu *et al*., 2009, Muchova *et al*., 2013). Structure-function mutagenesis shows that the cytoplasmic HTH and TM domains of RodZ are essential for its function in *E. coli* and *C. crescentus*, whereas extracellular domains can be deleted without severe cell growth and morphology phenotypes (Alyahya *et al*., 2009, Bendezu *et al*., 2009, Morgenstein *et al*., 2015). In this regard, RodZ of *Rickettsia* and *Chlamydia* species lack an extracellular domain (Kemege *et al*., 2015). These obligate intracellular pathogens lack FtsZ and use a modified Rod complex consisting of homologs of MreB, RodZ, MreC, a bPBP, and RodA to synthesize midcell PG (Liechti *et al*., 2016, Ouellette *et al*., 2020, Ranjit *et al*., 2020).

In this paper, we demonstrate by Tn-seq, transformation assays, and protein depletion that RodZ(*Spn*) is conditionally essential in serotype-2 D39 strains of *S. pneumoniae* at 37°C. Tn-seq and structure-function analyses show that this essentiality requires the HTH and TM domains of RodZ(*Spn*), but not the extracellular linker or DUF domains. Suppression patterns of Δ*rodZ* and Δ*mreC*, but not Δ*cozE*, mutants phenocopy each other, linking RodZ(*Spn*) to the pPG elongasome. The conclusion that RodZ(*Spn*) is a member of the pPG elongasome is supported by interaction studies using co-IP and B2H assays and by microscopic co-localization of RodZ(*Spn*) and MreC(*Spn*) or aPBP1a(*Spn*) throughout the pneumococcal cell cycle. Depletion of RodZ(*Spn*) stops growth and results in viable, rounded, heterogeneous cells with a qualitatively different appearance from cells depleted for MreC or bPBP2b (Berg *et al*., 2013, Land & Winkler, 2011, Tsui *et al*., 2014). Depletion of RodZ(*Spn*) or MreC(*Spn*) further reveals a hierarchy for pPG elongasome assembly. Finally, Tn-seq experiments show the unexpected result that Class A aPBP1b, whose function is not known, is in a synthetic-viable genetic relationship with RodZ, but not MreCD, whereas Class A aPBP1a is in a synthetic-viable genetic relationship with MreC, MreD, and RodZ. Together, these results show that RodZ(*Spn*) still acts as an essential scaffold protein through its HTH and TM domains for pneumococcal pPG elongasome assembly and function, despite the absence of MreB. This study also shows that diverse cell morphology and genetic phenotypes result when different members of the pPG elongasome are absent or depleted. Last, this work shows that aPBP1b and aPBP1a play different roles in modulating the function of the *S. pneumoniae* pPG elongasome and possibly participate in pPG synthesis. These results are discussed in terms of a model in which failsafe mechanisms can bypass or regulate the function of the pneumococcal core RodZ-MreCD-bPBP2b-RodA elongasome to ensure viability.

## 2 RESULTS

### 2.1 RodZ is conditionally essential in *Streptococcus pneumoniae* D39 at 37°C

RodZ(*Spn*) is annotated as “probably not essential” in serotype 2 D39 strains based on recent genomics approaches (see PneumoBrowse site) (Slager *et al*., 2018). Likewise, a Tn-seq screen of serotype 4 TIGR4 strains recovered insertions in *rodZ* that seemed to grow in laboratory media (van Opijnen & Camilli, 2012). Δ*rodZ* mutants were reported in unencapsulated (Δ*cps*) R6 laboratory strains (Martin-Galiano *et al*., 2014, Stamsas *et al*., 2017, Straume *et al*., 2017), whose progenitor is strain D39 (Lanie *et al*., 2007). However, we previously reported that we could not obtain Δ*rodZ* mutants in an unencapsulated (Δ*cps*) derivative constructed in progenitor strain D39 (Tsui *et al*., 2016).

We performed a series of experiments to reconcile these conflicting previous results, leading to the conclusion that *rodZ* is essential for growth at 37°C in unencapsulated and encapsulated D39 strains, although poor growth occurs at 32°C (Fig. S2). Tn-seq analysis showed that insertions occur in the non-essential carboxyl-terminal DUF-domain half of *rodZ*(*Spn*), but are not recovered in the essential transmembrane and amino-terminal domains (Fig. 1D and 2A (WT)). This pattern of insertions relative to RodZ(*Spn*) domain functions is taken up further below and underlies why a substantial number of non-essential *rodZ* insertions were detected in previous Tn-seq experiments. In addition, the Tn-seq profile here (Fig 2A) confirms previous Tn-seq (Fenton *et al*., 2016) and complementation results (see below; (Rued *et al*., 2017)) showing that *mreC* is essential in *S. pneumoniae* D39, contrary to a conclusion in (Straume *et al*., 2017).

**Figure 2.**
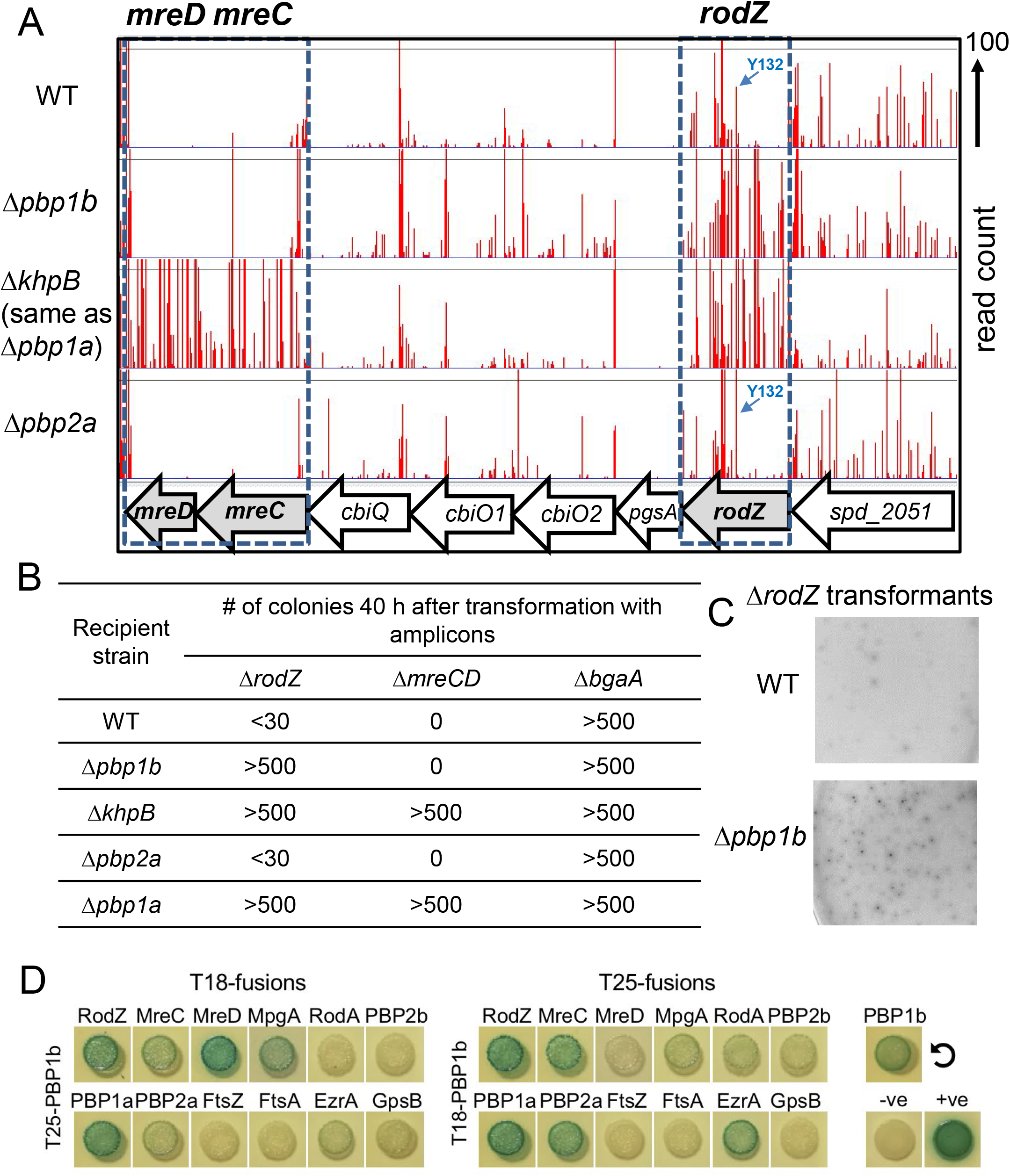
Tn-seq analysis reveals suppression of Δ*rodZ,* but not Δ*mreCD,* lethality by Δ*pbp1b* deletion. (A) Tn-Seq transposon insertion profile for the genome region covering *spd_2051, rodZ, pgsA, cbiO2, cbiO1, cbiQ, mreC, and mreD* of mini-Mariner *Malgellan6* transposon (Tn) into the genomes of WT (D39 Δ*cps rpsL1*, IU1824), Δ*pbp1b* (IU14697), Δ*khpB* (IU10592), or Δ*pbp2a* (IU13256) strains. *In vitro* transposition reactions, containing purified genomic DNA, *Magellan6* plasmid DNA, and purified MarC9 mariner transposase, transformation, harvesting of transposon-inserted mutants, NextSeq 75 high-output sequencing, and analysis were performed as described in *Experimental procedures*. (B) Transformation assay confirming that Δ*pbp1b* suppresses RodZ, but not MreCD, essentiality. Results were obtained 40 h after transformation of WT, Δ*pbp1b*, Δ*khpB*, Δ*pbp2a,* or Δ*pbp1a* (IU6741) strains with linearized Δ*rodZ*::P_c_-*aad9*, Δ*mreCD*<>*aad9,* or positive control Δ*bgaA*::Pc-*erm* amplicons. Numbers of transformants were normalized to correspond to 1 mL of transformation mixture. Similar results were obtained in two or more independent experiments. Similar results were obtained after 24 h of incubation, except that colonies Δ*rodZ* Δ*pbp1b* transformants were fainter in appearance than at 40 h, and <10 Δ*rodZ* colonies were obtained in the WT and Δ*pbp2a* backgrounds (data not shown). (C) Appearance of colonies of the WT or Δ*pbp1b* strain 40 h after transformation with the Δ*rodZ*::P_c_-*aad9* amplicon. (D) aPBP1b interacts with RodZ, MreC, MreD, EzrA, MpgA, aPBP1a, and aPBP2a as well as with itself in B2H assays. Agar plates were photographed after 40 h at 30°C.

Transformation assays confirmed that Δ*rodZ* was not viable in unencapsulated WT D39 strains at 37°C, unless it was complemented in a merodiploid strain by an ectopic copy of *rodZ*^+^ that was under control of a zinc-inducible promoter (Table 1, rows 1-4). At the lower temperature of 32°C, we did observe small colonies of Δ*rodZ* mutants in merodiploid strains lacking the -Zn inducer for *rodZ*^+^ expression (Fig. S2A) or in transformation assays (Fig. S2C, line 3). However, we found that slow growth at 32°C is a phenotype of mutants lacking other components of the pneumococcal pPG elongasome, including MreC, bPBP2b, or RodA (Fig. S2B; Fig. S2C, lines 4-6). In contrast to the unencapsulated D39 background, Δ*rodZ* transformed into the R6 laboratory strain (Table 1, line 17), as reported previously (Stamsas *et al*., 2017, Straume *et al*., 2017); however, R6 derivatives contain mutations in *pbp1a* that suppress mutations in genes encoding the peripheral PG (pPG) synthesis elongasome machine (Land & Winkler, 2011, Tsui *et al*., 2016), including Δ*mreC* (Table 1, line 17). Finally, comparable experiments at 37°C showed that Δ*rodZ* could not be transformed into encapsulated D39 progenitor strains (Table 1, lines 18 and 19). Numerous mutations suppress Δ*rodZ* in *S. pneumoniae* (see next section) and likely account for the small number of colonies that arose in the D39 strains transformed with Δ*rodZ* amplicons. The essentiality of *rodZ* in the encapsulated D39 strains was corroborated in merodiploid strains, where hundreds of colonies were obtained only in the complementation strain (+Zn), but not in the depleted Δ*rodZ* strain (-Zn; Table 1, lines 20 and 21). We conclude that *rodZ* is essential for growth of unencapsulated or encapsulated D39 strains of *S. pneumoniae* at the optimal culture temperature of 37°C. Because capsule partially masks primary phenotypes of PG synthesis mutants in *S. pneumoniae* and complicates microscopy due to cell chaining (Barendt *et al*., 2009), the rest of these studies of *rodZ* physiology and function were performed in the D39 unencapsulated genetic background.

**Table 1.**
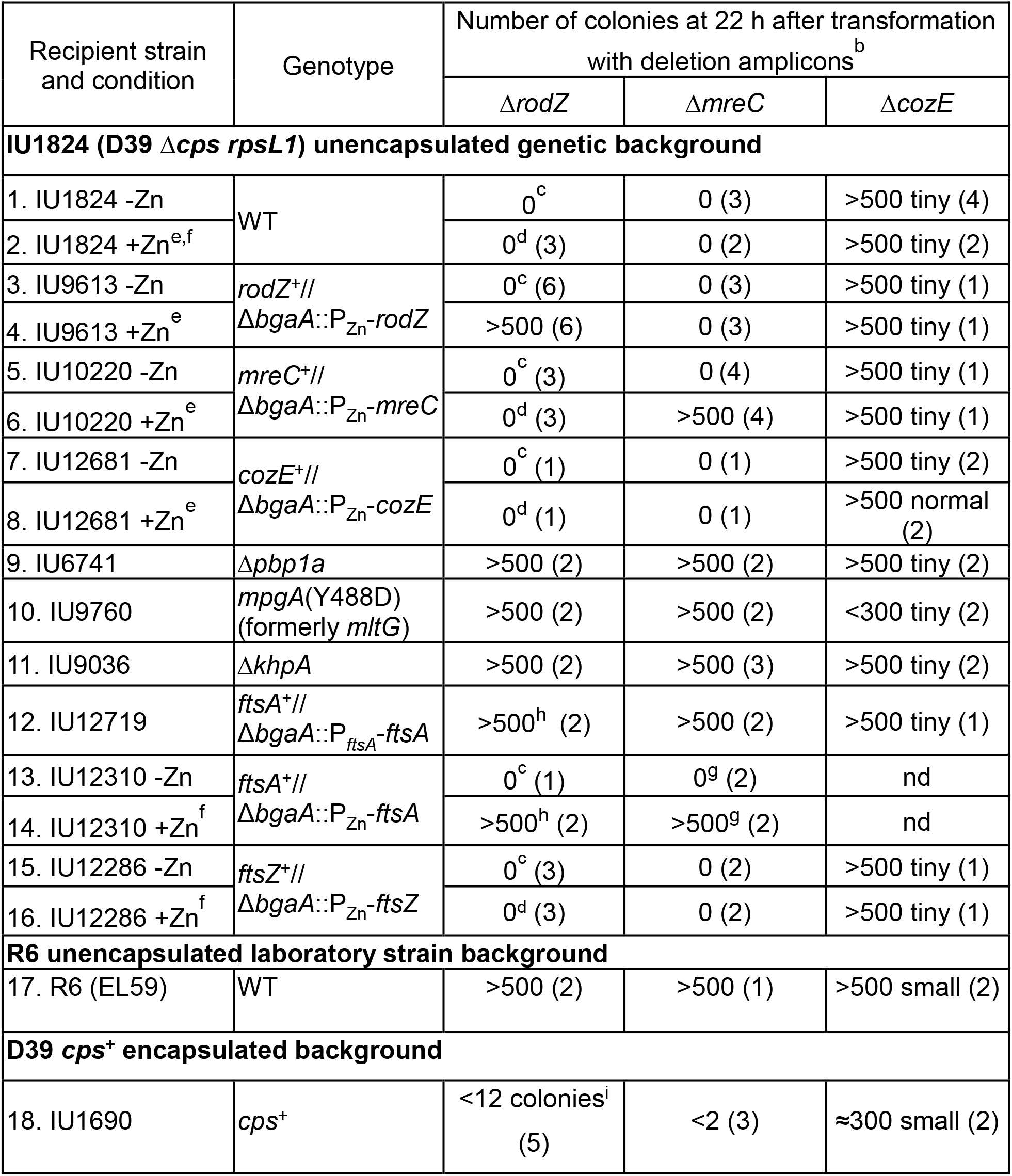
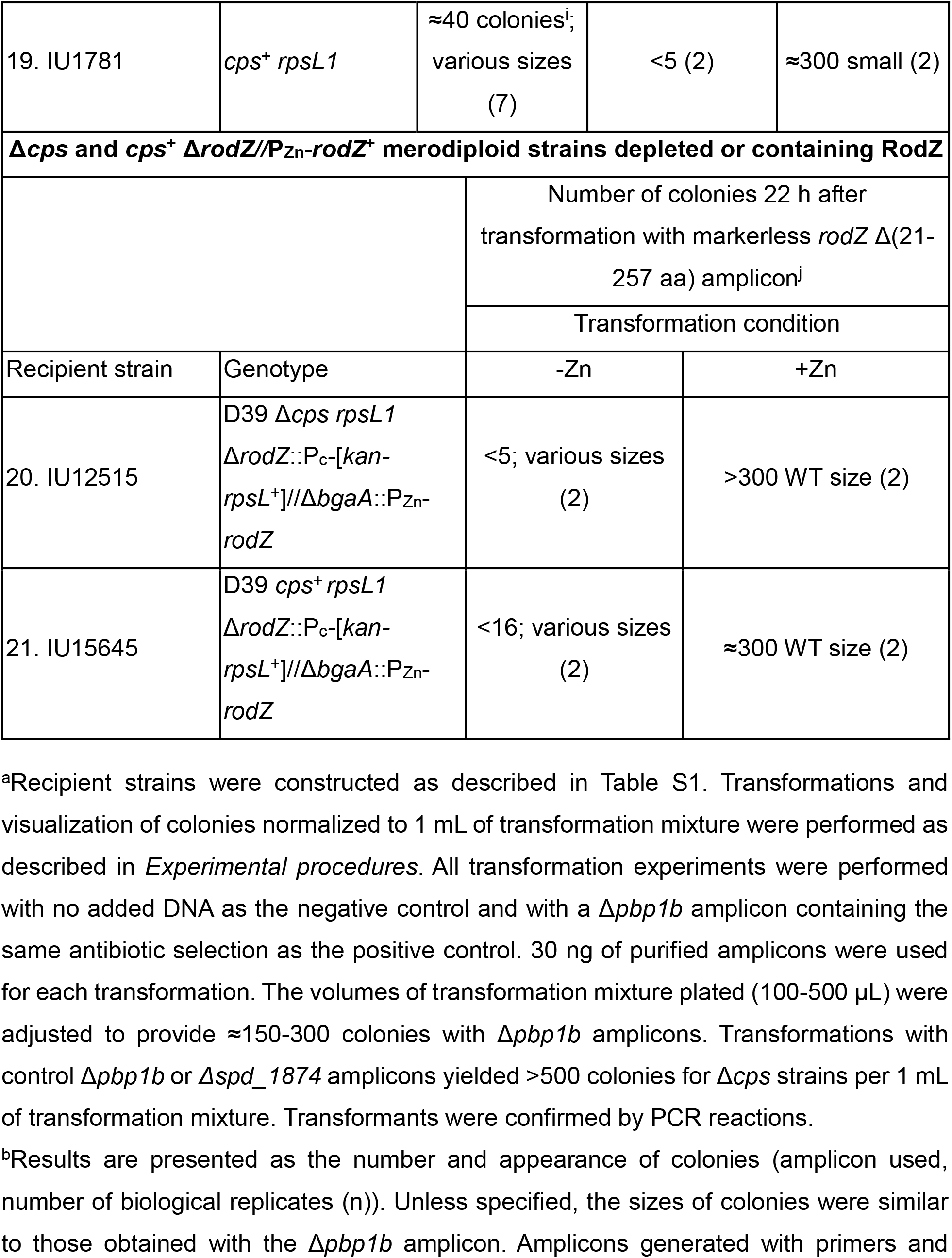
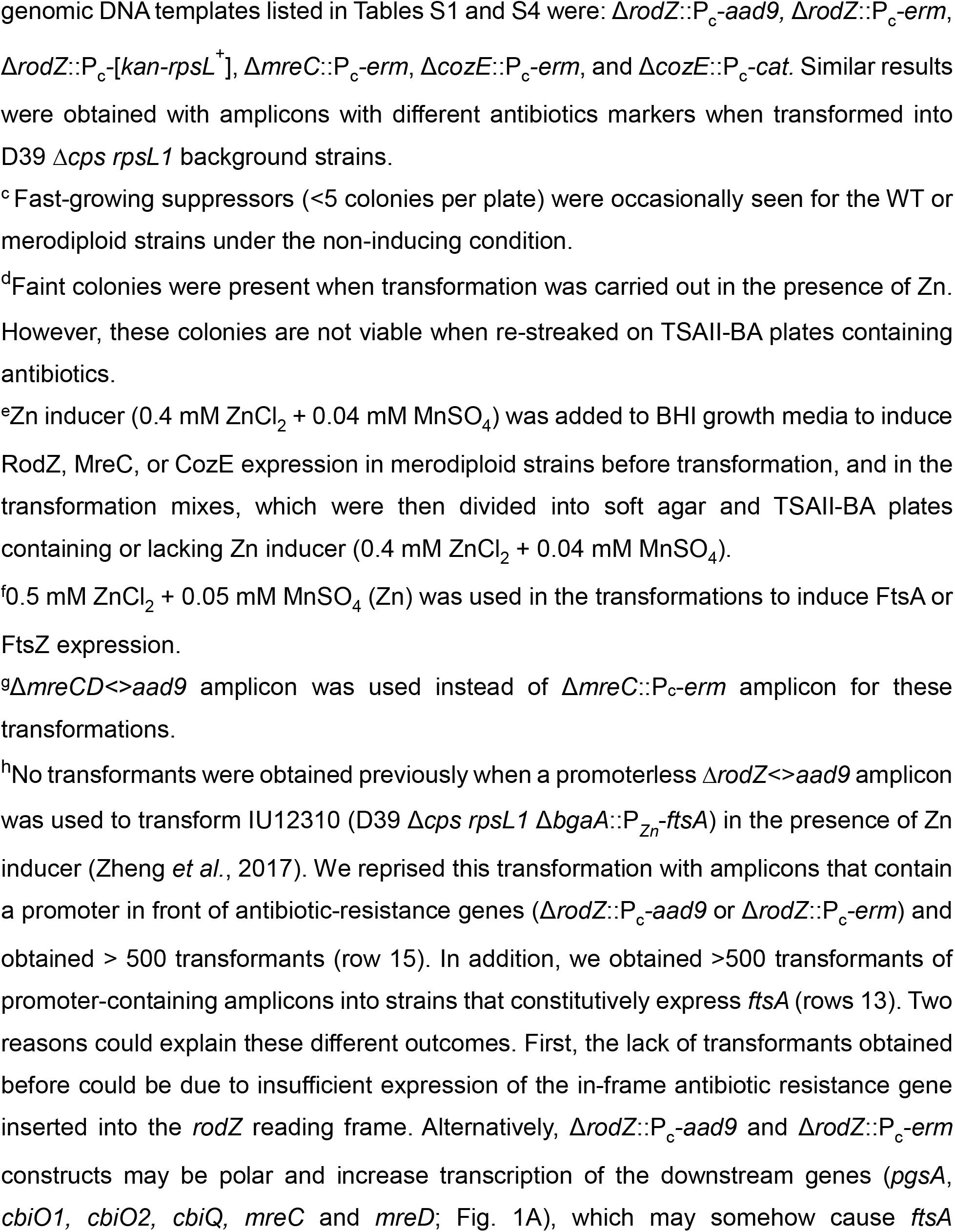
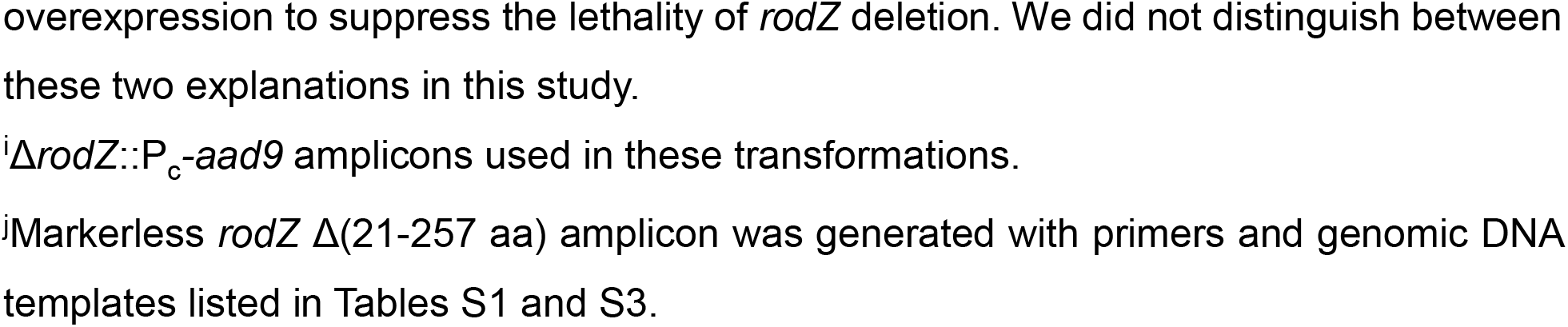
Δ*rodZ* has a similar suppression pattern as Δ*mreC*, but not to Δ*cozE*, in transformation assays at 37°C^a^

### 2.2 Δ*rodZ* has a similar suppression pattern as Δ*mreC*, but not to Δ*cozE*, in transformation assays at 37°C

Mutations in essential genes of the pneumococcal PG elongasome complex, including *mreC*, *pbp2b*, and *rodA*, are suppressed by several kinds of mutations (Land & Winkler, 2011, Stamsas *et al*., 2017, Tsui *et al*., 2016, Zheng *et al*., 2017). For example, Δ*mreC* is suppressed by Δ*pbp1a* (Class A aPBP1a), *mpgA*(Y488D) (reduced activity of MpgA (formerly MltG(*Spn*)) muramidase, Δ*khpA* (RNA binding regulator), and overexpression of FtsA (division actin-homolog) (Table 1, lines 9-14) (Land & Winkler, 2011, Stamsas *et al*., 2017, Taguchi *et al*., 2021, Tsui *et al*., 2016, Zheng *et al*., 2017). To link RodZ to MreC and pPG synthesis, we showed that Δ*rodZ* has the same suppression pattern as Δ*mreC*. Δ*rodZ* or Δ*mreC* is complemented in merodiploid strains by ectopic expression (+Zn) of *rodZ*^+^ or *mreC*^+^, respectively (Table 1, lines 4 and 6). Like Δ*mreC*, Δ*rodZ* is suppressed by Δ*pbp1a*, *mpgA*(Y488D), Δ*khpA*, and overexpression of FtsA (Table 1, lines 9-14). This identical suppression pattern provides strong evidence that RodZ functions in the pPG synthesis elongasome, along with MreC, bPBP2b, and RodA. In addition, overexpression of MreC or FtsZ in merodiploid strains did not bypass the requirement for RodZ (Table 1, lines 6 and 16). Likewise, the requirement for MreC was not bypassed by overexpression of RodZ or FtsZ (Table 1, lines 4 and 16).

We used transformation assays to study two other aspects of RodZ function. CozE was discovered in a Tn-seq of essential genes that become dispensable in a mutant lacking aPBP1a (Fenton *et al*., 2016). Localization and interaction studies indicated that CozE is a member of the MreCD complex in the pneumococcal elongasome. Therefore, we fully expected Δ*cozE* mutants to show the same suppression patterns as Δ*mreC* and Δ*rodZ* mutants. Unexpectedly, we found that Δ*cozE* significantly reduced, but did not abolish, colony growth, indicating that *cozE* is dispensable under the conditions used here (Table 1, line 1 and 2). This non-essentiality was recapitulated in Tn-seq experiments showing that insertion in *cozE* are recovered in the WT strain propagated in BHI broth (see below; data not shown). The colony growth defect of a Δ*cozE* mutant was fully complemented by ectopic expression of *cozE*^+^ in a merodiploid strain (Table 1; line 8). However, the Δ*cozE* colony growth was not ostensibly improved by any of the mutations that suppressed both Δ*mreC* and Δ*rodZ*, including Δ*pbp1a* (Table 1, lines 9-16). Thus, loss of CozE is not equivalent to loss of MreC or RodZ under some growth conditions, suggesting different functions in the elongasome and/or cell growth. This difference was not studied further here.

Last, *rodZ* is immediately upstream of essential *pgsA*, which encodes phosphatidylglycerol phosphate synthase, required for phospholipid synthesis (Fig 1A). In *B. subtilis,* insertion mutations in *rodZ* can have polar effects on *pgsA* expression (van Beilen *et al*., 2016). In this study of *S. pneumoniae*, *rodZ* mutant growth and cell morphology phenotypes are complemented by expression of an ectopic copy of *rodZ*^+^ and markerless Δ*rodZ* mutations are used for most experiments (Table 1, lines 4, 20, and 21; see below). Conversely, ectopic expression of *pgsA*^+^ (+Zn) complemented Δ*pgsA* for growth in a Δ*pgsA*//P_Zn_-*pgsA*^+^ merodiploid (Table S5, line 11), but overexpression of PgsA in a *pgsA^+^*//P_Zn_-*pgsA*^+^ merodiploid strain (+Zn) did not allow growth of Δ*rodZ* transformants (Table 1, line 9). We conclude that phenotypes attributed to mutations in *rodZ* in this study are not caused by polarity on *pgsA*.

### 2.3 RodZ is required for normal cell shape and morphology of *S. pneumoniae*

To study primary *rodZ* mutant phenotypes without accumulating suppressors (Table 1), RodZ was depleted in Δ*rodZ*//P_Zn_-*rodZ*^+^ and Δ*rodZ*//P_Zn_-*rodZ-*FLAG merodiploid mutants (-Zn), and cultures were sampled at various times after depletion (Fig. 3 and 4, where F is used as an abbreviation for the FLAG tag here and elsewhere). RodZ depletion caused a decrease in growth rate after ≈4.5 h, followed by a decrease in culture density at ≈7 h (Fig. 3A and 4A). In controls for depletion experiments, cells expressing functional RodZ-FLAG from the native locus grew at the same rate in BHI broth lacking or containing the Zn inducer (0.4 mM ZnCl_2_ + 0.04 mM MnSO_4_), where 1/10 MnSO_4_ was added to lessen Zn^2+^ toxicity (Fig. S3A} (Jacobsen *et al*., 2011, Perez *et al*., 2021b, Tsui *et al*., 2016), and cell morphology and cellular RodZ-FLAG amount were not appreciably altered by Zn addition (Fig. S3B and S3C). By 4 h of RodZ depletion, cells tended to become larger and more heterogeneous in shape, with some cells becoming round and others exhibiting pointed ends (Fig. 3B, 3C, and 4B). By 6 h of RodZ depletion, cell width and size increased significantly, and cells became more spherical with an average aspect ratio approaching 1 (Fig. 3B and 3C). Quantitative western blotting showed that RodZ-FLAG was not detectable by 3 h of depletion and that RodZ depletion for 4 h did not alter the cellular amounts of MreC, bPBP2b, or bPBP2x (Fig. 4D). Depletion of RodZ also led to a moderate increase in the number of cells in chains (Fig. S4).

**Figure 3.**
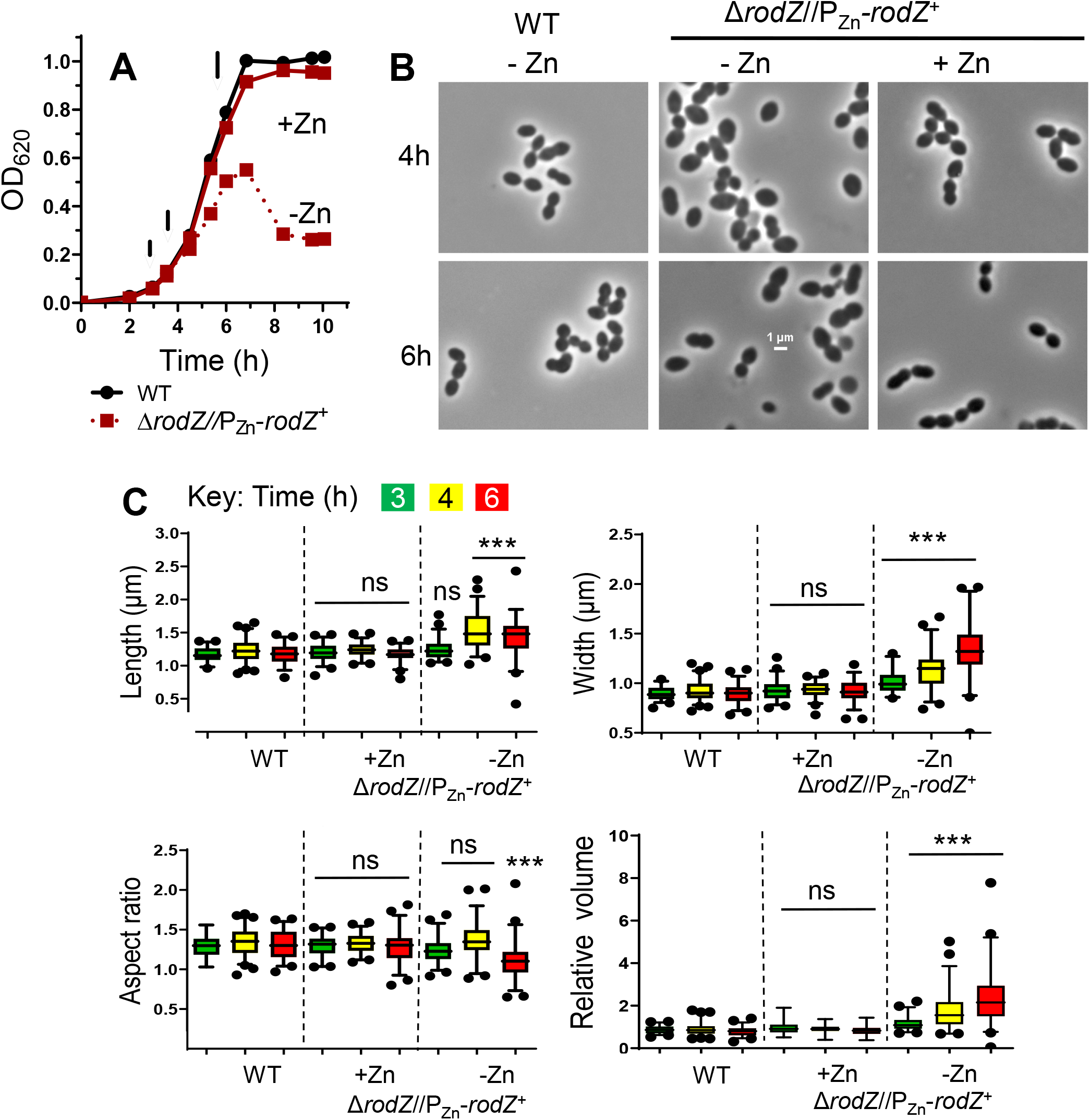
Depletion of RodZ results in cell rounding, indicative of a defect in peripheral PG synthesis. (A). Representative growth curves of IU1824 (WT) and IU12738 (Δ*rodZ*//P_Zn_-*rodZ*^+^) with or lacking Zn inducer (0.4 mM ZnCl_2_ + 0.04 mM MnSO_4_). IU1824 and IU12738 were grown overnight in BHI broth at 37°C lacking and with Zn inducer, respectively. Samples were re-suspended in fresh BHI ±Zn to an OD_620_ of ≈0.003. Arrows indicate times at which samples were taken for microscopy. (B) Representative micrographs of IU1824 (WT) and IU12738 (Δ*rodZ*//P_Zn_-*rodZ*^+^) grown in the presence or absence of Zn inducer after 4h or 6 h of growth. All images are at the same magnification (scale bar = 1 µm). (C) Box-and-whiskers plots (5 to 95 percentile) of cell length, width, aspect ratio, and relative volume measured for IU1824 grown in the absence of Zn, and IU12738 grown in the presence or absence of Zn for 3h, 4h, and 6h. For each time point ≈50-80 cells per sample were measured, and statistical analysis was conducted using one-way ANOVA Kruskal-Wallis test in GraphPad Prism. Statistical comparisons were carried out with IU12738 grown in the presence or absence of Zn vs the WT control at the respective time points. ***, p<0.001; ns, non-significant. Results shown are representative from one of at least three independent biological replicates.

**Figure 4.**
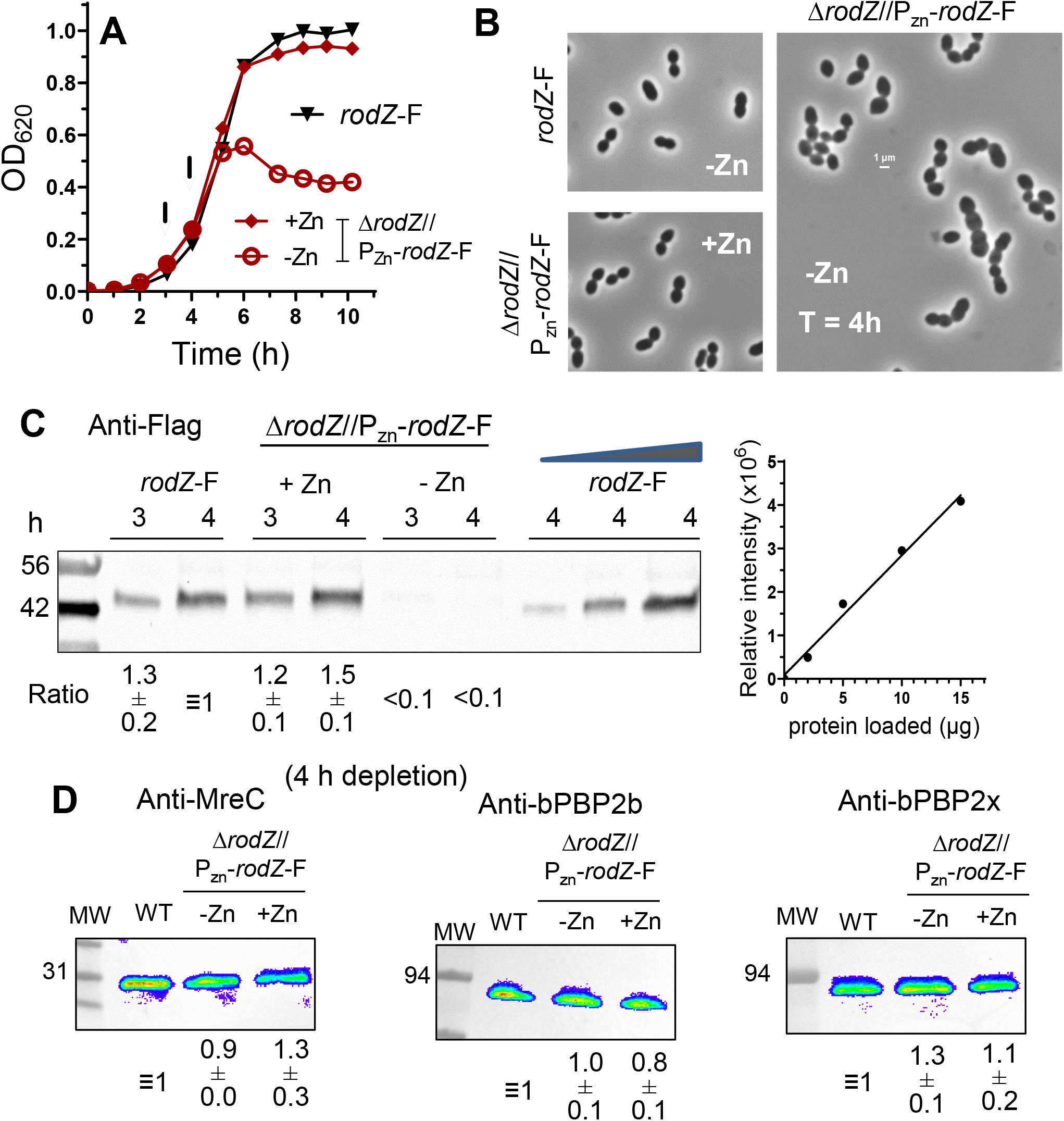
RodZ levels decrease to an undetectable level upon depletion for 3 h. (A) Representative growth curves of *rodZ^+^*-FLAG (IU14594), where F is used as an abbreviation for the FLAG tag, and depletion strain Δ*rodZ*//P_Zn-_*rodZ*^+^-FLAG (IU10947). IU14594 and IU10947 were grown overnight in BHI with and lacking Zn inducer (0.4 mM ZnCl_2_ + 0.04mM MnSO_4_), respectively, and diluted into BHI with no Zn for IU14594, and into BHI with or lacking Zn for IU10947. Cultures were sampled at 3 or 4 h for Western analysis (arrows). (B) Representative micrographs of IU10947 and IU14594 at 4 h. Scale bar = 1 µm. (C) Representative quantitative Western blot showing RodZ-F amount expressed from the native chromosomal site in IU14594, or from the ectopic site in the presence or absence of Zn inducer in IU10947 (Δ*rodZ*//P_Zn-_*rodZ*^+^-F) at 3h and 4h of growth. 10 µg of crude cell lysates were loaded in the left 6 lanes, and 2, 5, or 15 µg were loaded in the right three lanes to generate a standard curve for quantitation. SDS-PAGE and western blotting were carried out as described in *Experimental procedures* using Licor IR Dye800 CW secondary antibody detected with Azure biosystem 600. Signals obtained with anti-FLAG antibody were normalized with total protein stain in each lane using Totalstain Q-NC (Azure Scientific). Ratios indicate RodZ-F protein amounts (average ± SEM) from 3h or 4h samples obtained from IU10947 relative to IU14594 at 4h. (D) MreC, bPBP2b, and bPBP2x protein levels are not altered upon RodZ depletion. Protein samples were obtained from IU14594 (*rodZ*^+^-F WT), or IU10947 (Δ*rodZ*//P_Zn-_ *rodZ*^+^-F) grown in the presence or absence of Zn inducer for 4 h. 3 µg of crude cell lysates were loaded in each lane. SDS-PAGE and Western blotting were carried out with primary antibodies to MreC, bPBP2b, or bPBP2x. Chemiluminescence signals obtained with secondary HRP-conjugated antibodies were detected using an IVIS imaging system. Ratios indicate protein amounts (average ± SEM) in IU10947 (Δ*rodZ*//P_Zn-_*rodZ*^+^-F) relative to those in IU14594 (WT) from 2 independent biological replicates.

Depletion of RodZ for 4 h or 6 h is bacteriostatic and did not lead to a loss of cell viability, as determined by live-dead staining (Fig. 5) or by recovery of CFUs following RodZ depletion for at least 7 h (Fig. S5). Finally, we tested whether overexpression of RodZ affected growth or cell morphology of *S. pneumoniae*, as happens in *E. coli* and *C. crescentus* (Alyahya *et al*., 2009, Bendezu *et al*., 2009, Shiomi *et al*., 2008). Overexpression of RodZ-FLAG by ≈2.5 fold (Fig. S6C) did not have an appreciable effect on pneumococcal growth (Fig. S6A) or cell morphology (Fig. S6B and S6D), although more lysed cells were observed when RodZ-FLAG was overexpressed. Altogether, these results indicate that RodZ is required for normal morphology of pneumococcal cells. Notably, cell shape and size tend to be more heterogeneous for RodZ depletion than for depletion of other pneumococcal elongasome components, such as MreC, bPBP2b, or FtsEX, which results in chains of relatively uniform, spherical cells at 4 h (Berg *et al*., 2013, Sham *et al*., 2013, Tsui *et al*., 2014).

**Figure 5.**
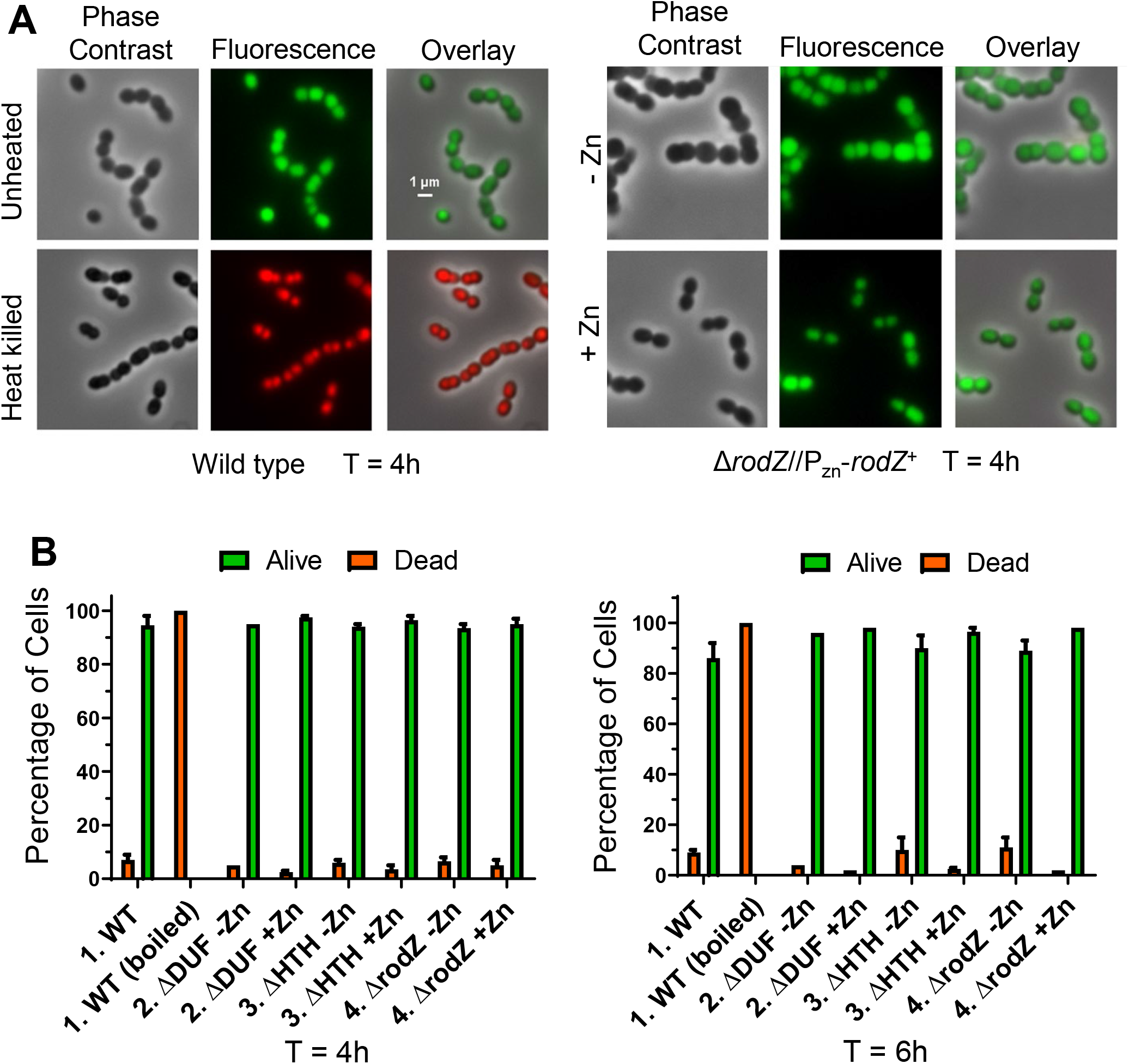
Cells depleted of RodZ for 6 h remain viable. (A) Representative micrographs of WT (IU1824, left panels) or Δ*rodZ*//P_Zn_-*rodZ*^+^ merodiploid strain (IU12738, right panels) after 4 h of growth in the absence or presence of Zn inducer. Cells were stained with the Live-Dead BacLight Bacterial Viability Kit (Syto9 and Propidium Iodide) as described in *Experimental procedures*. Left, majority of WT cells during exponential growth displayed the green “live” staining pattern, while cells heat-killed by boiling for 5 min at 95°C displayed the red “dead” staining, indicative of membrane permeabilization. Scale bar = 1 µm. (B) Quantitation of the percentage of live or dead cells of exponentially growing or boiled WT strain (IU1824), and *rodZ*(ΔDUF)//P_Zn_-r*odZ* (IU12699), *rodZ* (ΔHTH)//P_Zn_-r*odZ* (IU12696), or Δ*rodZ*//P_Zn_-r*odZ* (IU12738) strains grown in the presence or absence of Zn inducer. 200 cells were examined and categorized for each sample. Data were from 2 independent experiments, except for the 6h time points of IU12699 and IU12738, which are from a single experiment.

### 2.4 The RodZ(*Spn*) HTH and TM domains, but not the DUF domain, are required for normal growth

RodZ(*Spn*) has the same overall domain structure as RodZ from bacteria that express MreB homologs (*Introduction*; Fig. 1 and S1). Tn-seq of the WT strain demonstrated that viable insertions are obtained in the C-terminus of *rodZ*(*Spn*) starting with TA in the TAT (Y132) codon, resulting in a stop codon after AAC (N131) (Fig. 1D and 2A). Thus, the entire extracellular region of RodZ(*Spn*) from aa Y132-N273, including DUF4115 and the disordered extracellular domain proximal to the membrane are dispensable for growth in BHI broth at 37°C. This conclusion was confirmed by deletion mutations constructed in a *rodZ*//P_Zn_-*rodZ^+^* merodiploid strain (Table 2, lines 6 and 7; Fig. 6). RodZ(M1-Q134) and RodZ(M1-T135) mutants formed normal-sized colonies on TSAII-BA plates, as did other deletion mutants of the extracellular domains (Table 2; Fig. 6A). Cells of RodZ(ΔDUF) and RodZ(M1-Q195; lacking DUF and the C-terminal region) resemble WT cells in BHI broth at 37°C (Fig. 6B and S7). By contrast, RodZ(M1-Q134) and RodZ(M1-T135) mutants form wider, larger cells than WT, indicative of partial RodZ function or instability. C-terminal FLAG-tagged WT RodZ was readily detected by western blotting (Fig. 4C). Curiously, truncated RodZ(M1-Q195) with a C-terminal FLAG-tag could not be detected by western blotting (Fig. 6), despite not showing growth or morphology phenotypes, suggesting C-terminal degradation in the absence of the DUF domain.

**Figure 6.**
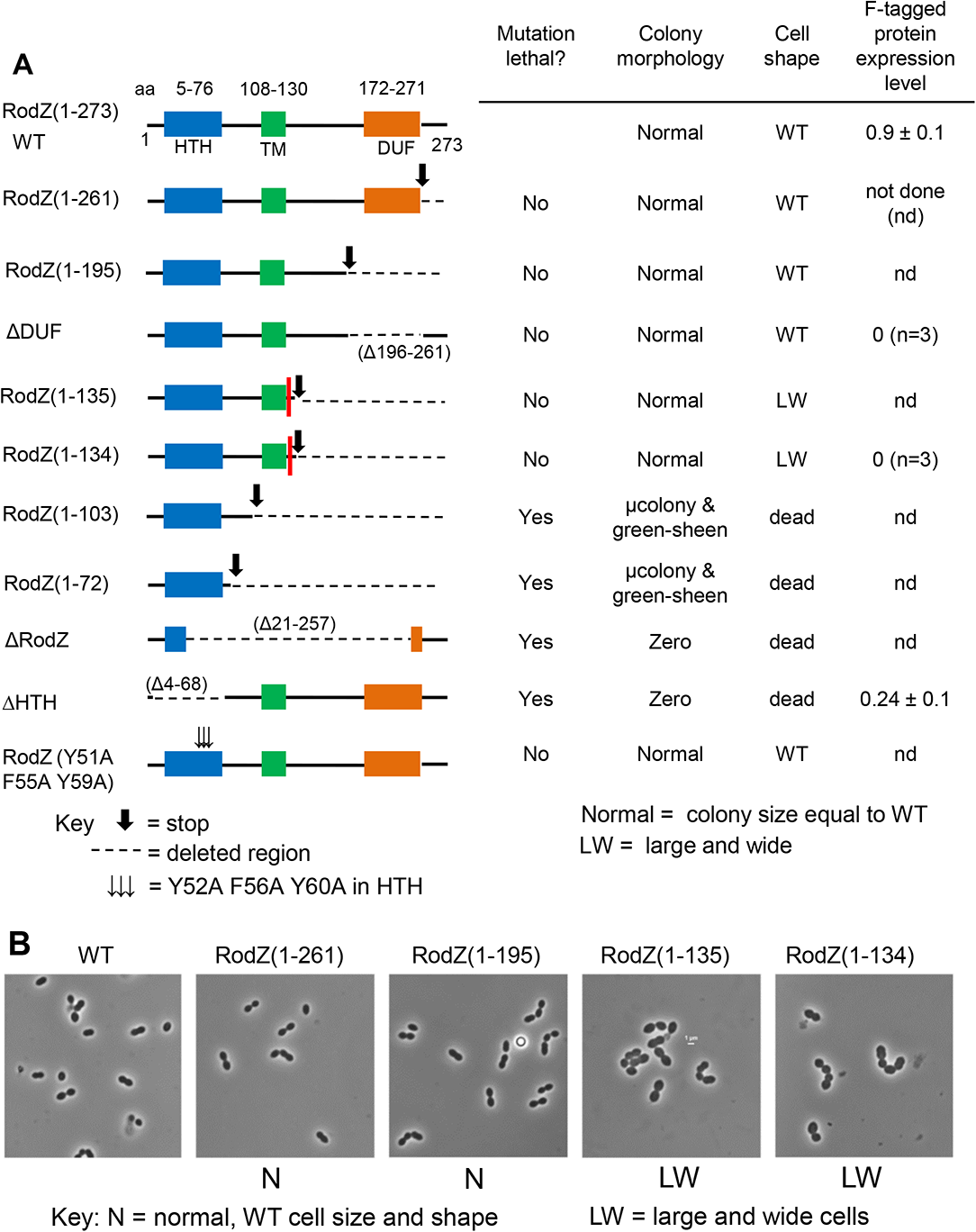
Amino Acids 1-131 of RodZ are required for growth of *Spn*. (A) Amplicons harboring *rodZ* truncation alleles were transformed into merodiploid strain IU12515 (Δ*rodZ*::P_c_*-*[*kan-rpsL*^+^] //P_Zn_-*rodZ^+^*) to replace the Janus cassette (Δ*rodZ*::P_c_*-* [*kan-rpsL*^+^]) as described in *Experimental procedures*. Effects of RodZ truncations were determined by transformation assays on blood agar plates with or lacking Zn inducer. Colony numbers, sizes, and morphologies were evaluated in comparison with *rodZ*^+^ transformants, after 20-24 h incubation at 37°C (see legend to Table 1 for experimental details). “µcolonies” (micro colonies) are barely visible by eye, but readily observed using a low power microscope. “Green-sheen” refers to a shiny green pattern observed on top of the blood agar that may be due to partial hemolysis. Similar results were obtained in two independent transformation experiments. The red bar between N131 and Y132 in the RodZ(1-135) and RodZ(1-134) entries marks the first TA site with a TnMariner insertion recovered by Tn-seq of the WT strain (see Figure 2). Relative amounts of corresponding truncated RodZ proteins fused to a C-terminus FLAG tag were determined by quantitative western blotting probed with anti-FLAG antibody as described in *Experimental procedures*. Proteins samples were obtained from strains IU14594 (*rodZ*-F at native chromosomal locus), IU13457 (*rodZ-*F//P_Zn_-*rodZ*^+^), IU13655 (*rodZ*(ΔDUF)*-*F//P_Zn_-*rodZ*^+^), IU13660 ((*rodZ*(1-134)*-*F//P_Zn_-*rodZ*^+^), and IU13705 (*rodZ*(ΔHTH)*-*F//P_Zn_-*rodZ*^+^) (see Table S1). Strains were grown in BHI broth supplemented with Zn inducer overnight, followed by growth for 4 h in BHI media lacking or containing of Zn inducer. Values in the last column are amounts of truncated F-tagged RodZ variants grown in the absence of Zn relative to the amount of RodZ-F in IU14594. Although IU13655 (*rodZ*(ΔDUF)*-*F//P_Zn_-*rodZ*^+^) and IU13660 ((*rodZ*(1-134)*-*F//P_Zn_-*rodZ*^+^) were viable without Zn, *rodZ*(ΔDUF)*-*F and *rodZ*(1-134)*-*F were undetectable in samples grown with or without Zn, consistent with cleavage of the FLAG tag off the truncated RodZ variants. (B) Representative micrographs of IU1824 (WT parent), and *rodZ* truncation mutants IU12794 (*rodZ*(1-261)//P_Zn_-*rodZ*^+^), IU12797 (*rodZ*(1-195)//P_Zn_-*rodZ*^+^), IU12799 (*rodZ*(1-135)//P_Zn_-*rodZ*^+^), and IU12803 (*rodZ*(1-134)//P_Zn_-*rodZ*^+^), which grew in the absence of Zn inducer. Cells were imaged during exponential growth at an OD_620_ ≈0.1-0.15 after ≈2.5-3.0 h of growth. Representative growth curves of truncated RodZ variants are shown in Figure S9D. Micrographs are at the same magnification (scale bar in panel 4) and representative of two independent biological replicates.

**Table 2.**
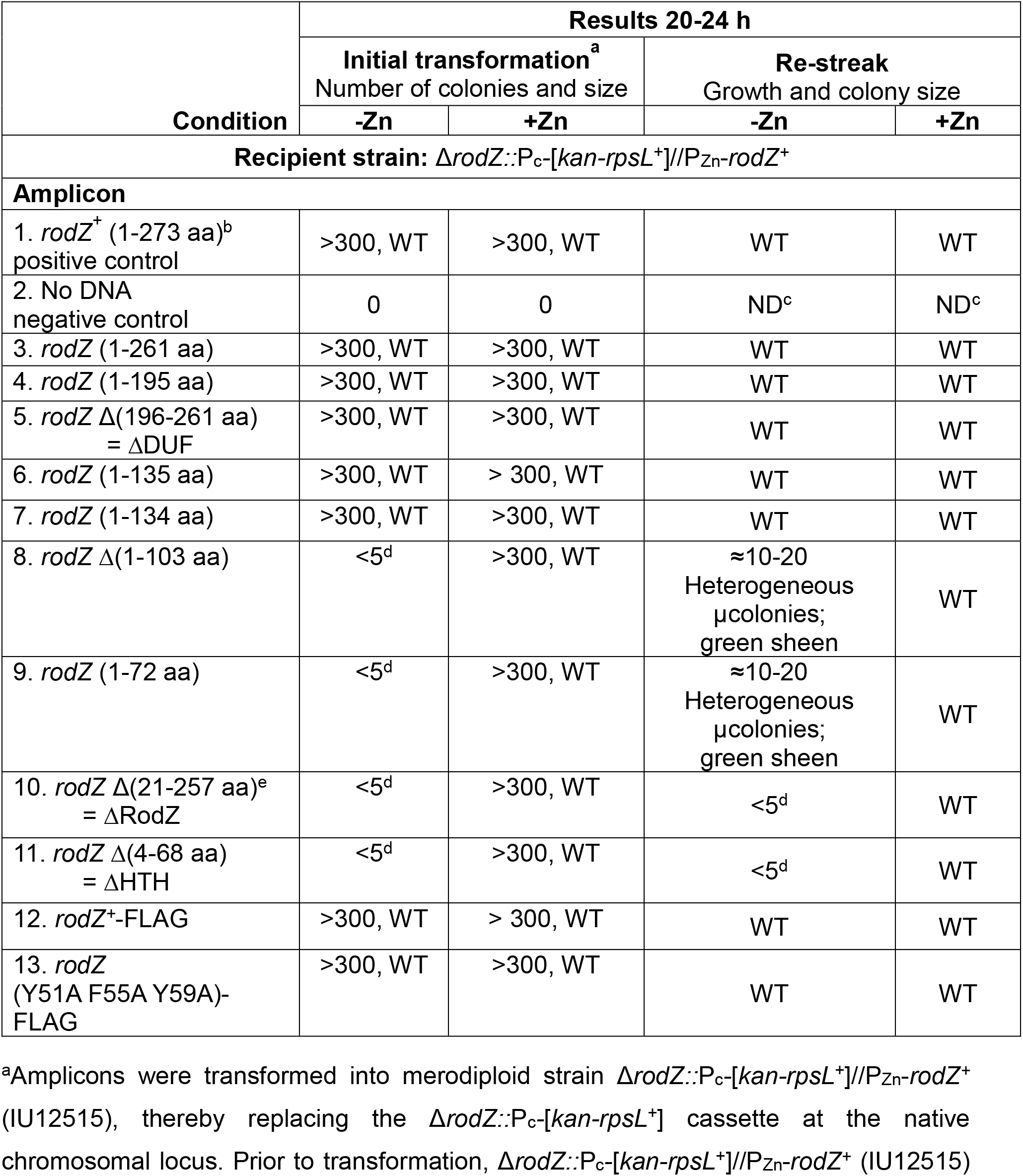
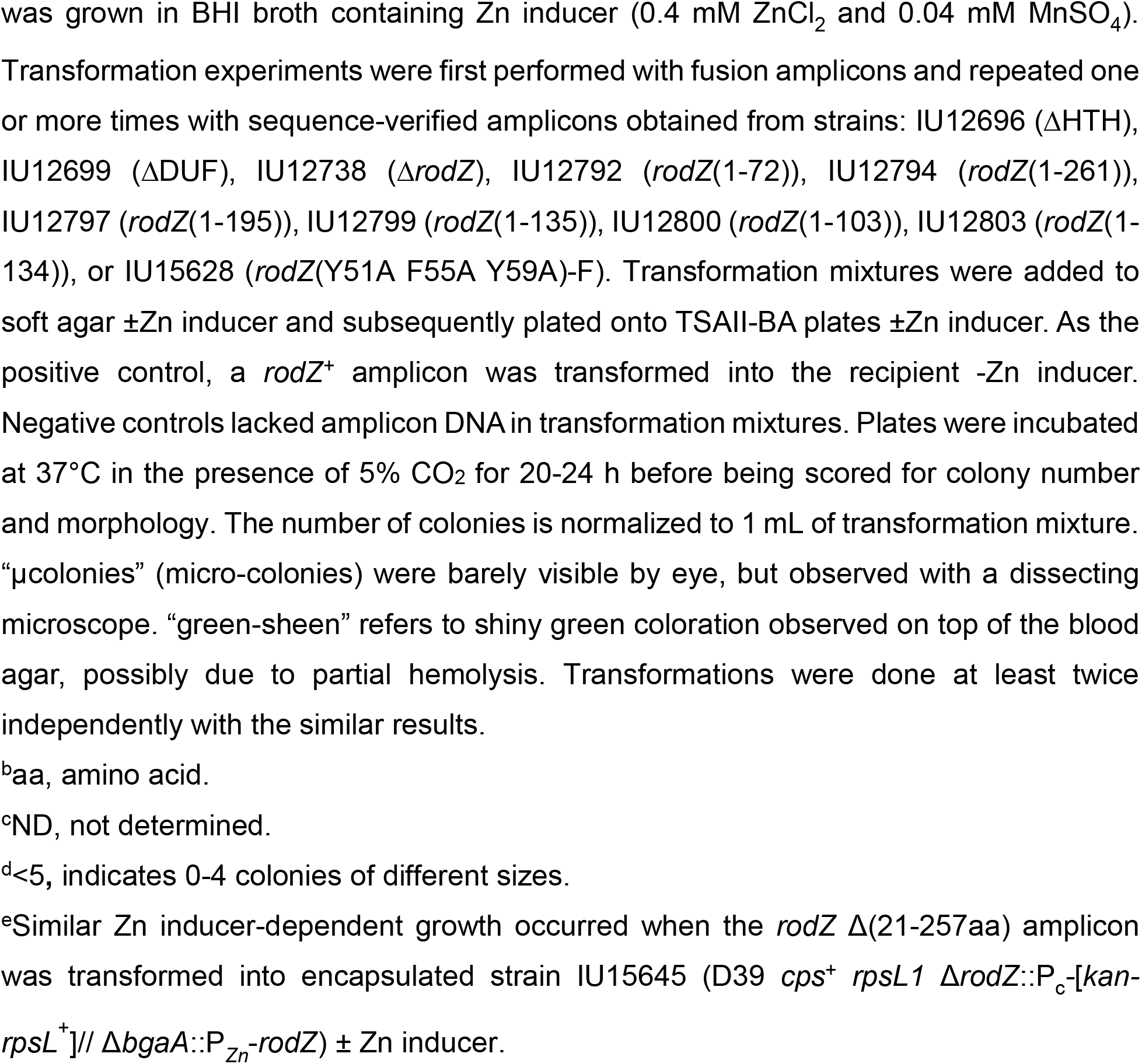
The RodZ(*Spn*) HTH and TM domains are essential, while the extracellular domain, including DUF, is dispensable

Transformation assays and growth characterization indicated that mutants lacking the TM, HTH, or the cytoplasmic linker domain of RodZ(*Spn*) are not viable (Table 2; Fig. 2A, 6A, and S8). RodZ(ΔHTH) mutant protein with a C-terminal FLAG-tag was detected in 4-h depletion experiments (Fig. 6A), consistent with lack of RodZ(ΔHTH) function rather than protein instability underlying its null mutant phenotype. Changes of aromatic amino acids in RodZ(*Spn*) at three positions in Helix 4 of the HTH domain, which correspond to the MreB interaction interface in *E. coli* (van den Ent *et al*., 2010), do not show growth phenotypes in *S. pneumoniae* (Fig. 6A and S9). In addition, amino-acid changes in the membrane-proximal pedestal region of bPBP2b failed to suppress Δ*rodZ* lethality in *S. pneumoniae* (Table S6), unlike the corresponding amino acid changes in *E. coli* bPBP2 that did suppress Δ*rodZ* phenotypes (Rohs *et al*., 2018). Finally, phosphorylation of S85 in RodZ(*Bsu*) was proposed to increase MreB filament density and growth (Sun & Garner, 2020). S85 of RodZ(*Bsu*) corresponds in alignment to E89 of RodZ(*Spn*) (dotted box, Fig. 1D; Fig. S1), which is not immediately adjacent to other serine or threonine residues. Phostag-PAGE analysis failed to detect phosphorylation of functional RodZ-HA^3^ in *S. pneumoniae* (Fig. S10). From these combined results, we conclude that the cytoplasmic HTH and linker, but not the extracellular domains, of RodZ(*Spn*) are required for growth at 37°C under the conditions tested here and that amino acids important for RodZ function in *E. coli* and *B. subtilis* are not required in *S. pneumoniae* (see *Discussion*).

### 2.5 RodZ(*Spn*) localizes with known pPG elongasome proteins throughout the pneumococcal cell cycle

The identical suppression pattern of Δ*rodZ* and Δ*mreC* in *S. pneumoniae* (Table 1) supports the hypothesis that RodZ is a member of the pPG elongasome. This hypothesis is further corroborated by protein co-localization analyses using immunofluorescence microscopy (IFM) as described in *Experimental procedures*. Strains expressing RodZ-FLAG^3^ constructs or other epitope-tagged proteins from their native chromosomal loci were functional and did not exhibit aberrant growth or cell morphologies (Fig. S11). We used a previously published method to compare the average locations of two fluorescent epitope-tagged proteins relative to DAPI-stained nucleoids at four stages of division in pneumococcal cells growing exponentially in BHI broth at 37°C (Fig. 7) (Land *et al*., 2013, Tsui *et al*., 2014). This method also allows statistical comparisons of average midcell widths at different cell division stages. By this analysis, RodZ co-localizes throughout the cell cycle with MreC and aPBP1a (Fig. 7A-7D), which have been implicated in pPG elongation in *S. pneumoniae* (Briggs *et al*., 2021, Fenton *et al*., 2016, Land & Winkler, 2011, Philippe *et al*., 2014, Straume *et al*., 2017, Tsui *et al*., 2016). All three proteins localize at the midcell equator in Stage 1 cells, remain at the midcell septum in Stage 2 and 3 cells, and only appear at the new equators of daughter cells late in division at Stage 4. In contrast to RodZ, nascent FtsZ-rings move outward toward the future sites of the new equators throughout division and largely leave the septum in Stage 3 and 4 cells (Fig. 7E and 7F) (Perez *et al*., 2019). We conclude that RodZ co-localizes with components of the pPG elongasome, which overlaps FtsZ localization in Stage 1 cells, but is different in later stages of the cell cycle. This conclusion is corroborated independently by high-resolution 3D-SIM of RodZ and FtsZ in cells at different division stages (Fig. S12).

**Figure 7.**
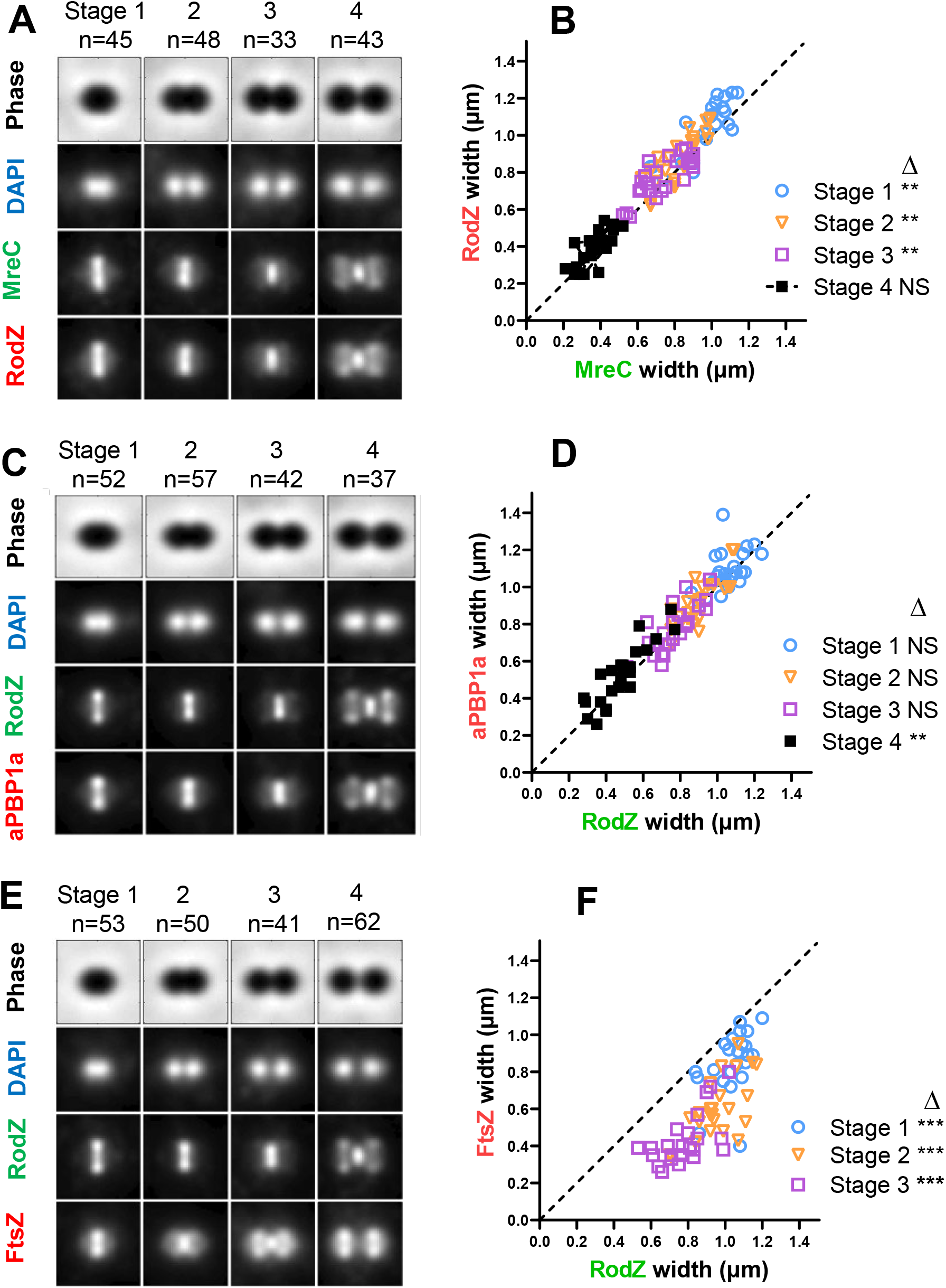
RodZ localizes with MreC and aPBP1a of the peripheral PG synthesis machine. (A) Composite image displaying localization patterns of MreC and RodZ through four stages of pneumococcal growth and division. Images were obtained by dual-labeling immunofluorescence microscopy (IFM). To construct composite images, n > 30 cells from each division stage were averaged and quantified as described in *Experimental procedures.* (A) IU7113 (*mreC*-L-F^3^ *rodZ*-Myc) IFM was probed with DAPI (DNA), anti-Myc, and anti-FLAG antibodies as detailed in *Experimental procedures.* (B) Scatter plot of the paired widths of RodZ compared to MreC constructed using the IMA-GUI program as described in *Experimental procedures*. The dotted line intercepts the origin with a slope of 1, indicating the expected distribution if MreC and RodZ widths were identical. Differences between the paired widths were calculated, and the null hypothesis was tested by the 1-sample student’s *t* test, where ** indicates P <0.01. (C) Composite image of RodZ and aPBP1a localization in IFM of IU7515 (*pbp1a*-L-F^3^ *rodZ*-Myc) probed with DAPI, anti-Myc, and anti-FLAG antibodies. (D) Scatter plot of paired width analysis of aPBP1a compared to RodZ. (E) Composite image of RodZ and FtsZ localization in IFM of IU7072 (*rodZ*-L-F^3^ *ftsZ*-Myc) probed with DAPI, anti-Myc, and anti-FLAG scatter plot of paired width analysis of FtsZ compared to RodZ. ***P value <0.001. Data were obtained from two independent experiments for each comparison.

### 2.6 RodZ(*Spn*) forms complexes and interacts with proteins in the septal PG (sPG) and pPG synthesis machines and with PG synthesis regulators

To gain more information about RodZ function in *S. pneumoniae*, we performed pairwise co-IP experiments using RodZ-FLAG and RodZ-FLAG^3^ as bait proteins that were probed in western blots for complex formation with proteins involved in PG elongation, septation, or cell division. Representative co-IP results are shown in Figure 8A, quantitated in Table 3, and summarized in Figure 8B and 8C. Additional supporting data are in Figures S13 and S14. Strong complex formation was detected between RodZ and pPG elongasome proteins MreC and bPBP2b at some stage of division in non-synchronized cell cultures (Table 3). The experiment to probe for complexes between RodZ and RodA could not be performed, because cells expressing RodZ-FLAG^3^ and HaloTag-RodA (HT-RodA) showed a synthetic lysis phenotype not observed in cells separately expressing the fusion proteins (Fig. S14A and S14B). Complexes were also detected between RodZ and protein regulators of PG synthesis (GpsB; StkP (Ser/Thr protein kinase); DivIVA), Class A PBPs (aPBP1a; aPBP2a), and MpgA (PG muramidase) (Briggs *et al*., 2021, Massidda *et al*., 2013). Consistent with these results, MreC, MpgA, or aPBP1a, each of which has been linked to pPG elongation in *S. pneumoniae* (Briggs *et al*., 2021, Fenton *et al*., 2016, Land & Winkler, 2011, Massidda *et al*., 2013, Philippe *et al*., 2014, Taguchi *et al*., 2021, Tsui *et al*., 2016), pulled down the same set of proteins (Table 3). RodZ was also detected in complexes with the sPG synthesis proteins bPBP2x (Fig. 8A) and FtsW (Fig. S14C) at some stage of cell division, including the initial equatorial ring of newly divided cells (Briggs *et al*., 2021). In contrast, marginal or no complexes were detected between RodZ and FtsA, FtsZ, PhpP (protein phosphatase), or KhpAB (RNA-binding regulator) (Fig. 8 and S13C; Table 3) (Massidda *et al*., 2013, Mura *et al*., 2017, Perez *et al*., 2019, Rued *et al*., 2017, Stamsas *et al*., 2017, Zheng *et al*., 2017). Thus, an *in vivo* complex containing RodZ and KhpB was not detected, despite a previous report of an interaction in a B2H assay (Winther *et al*., 2021).

**Figure 8.**
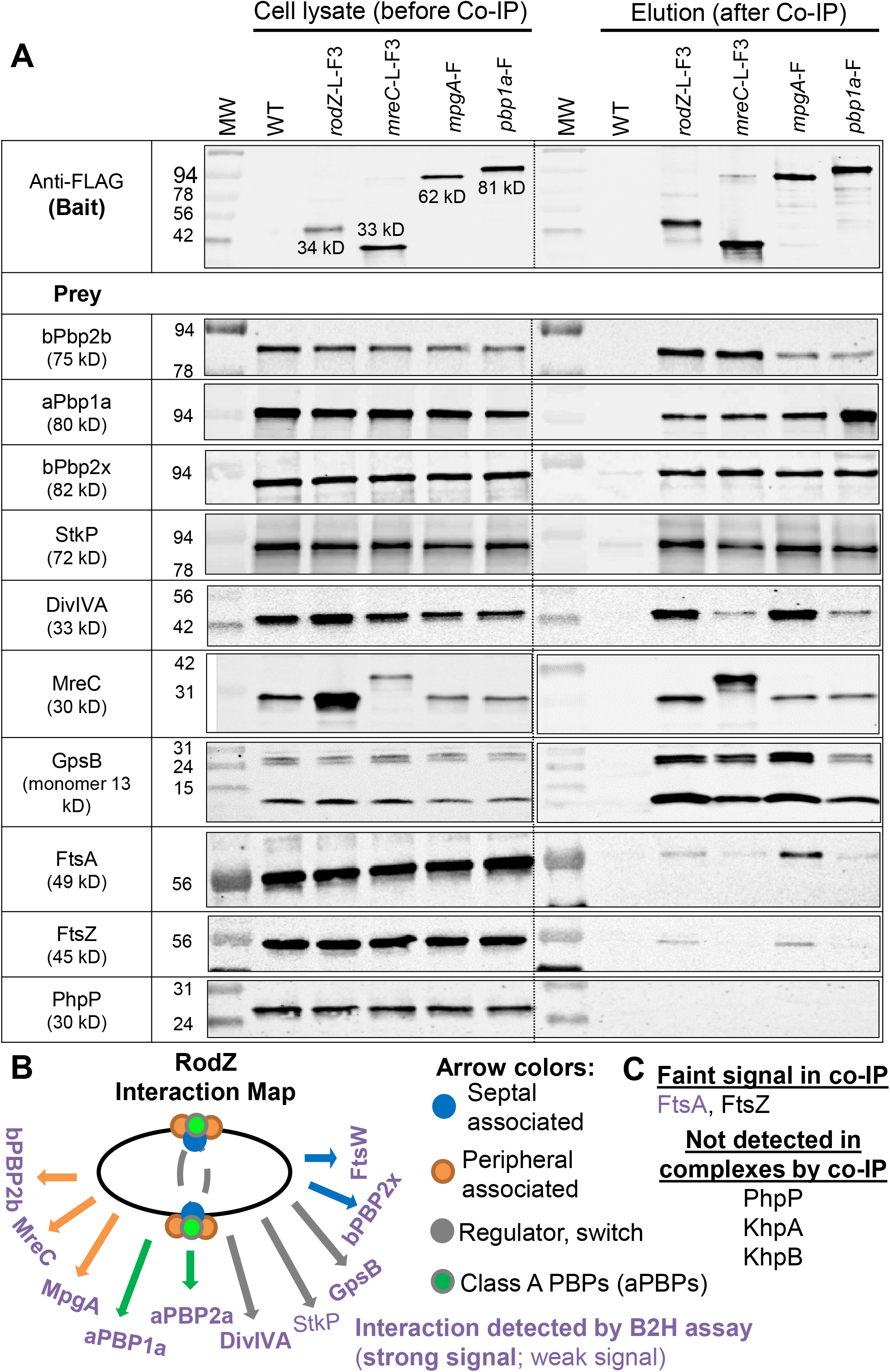
RodZ, MreC, MpgA (formerly MltG), and aPbp1a are in complexes with components of the peripheral and septal PG machines, class A PBPs, and cellular regulators StkP, GpsB, and DivIVA. Co-IP experiments using non-FLAG-tagged WT strain (IU1945) or FLAG-tagged strains RodZ-L-F^3^ (IU6291), MreC-L-F^3^ (IU4970), MpgA-F (IU7403), or PBP1a-F (IU5840) as bait were probed with native antibodies to detect prey proteins bPBP2b, aPBP1a, bPBP2x, StkP, DivIVA, MreC, GpsB, FtsA, FtsZ and PhpP, as described in *Experimental procedures*. Prey proteins are detected in all cell lysates (input; left lanes). In elution output samples (right lanes), prey proteins are undetectable for the WT non-FLAG-tagged control strain, but are present in different relative amounts in samples of the FLAG-tagged strains. Top blot was probed with anti-FLAG primary antibody for detection bait proteins. For most blots, 4 µl (4-6 µg) of each lysate sample (input) were loaded on the left lanes, while 15 µL of each elution output sample (after mixing 1:1 2x Laemmli buffer) was loaded onto the right lanes. For detection of GpsB, 6 µl (6 µg) of lysate sample and 25 µL of output were loaded. Two bands are detected with anti-GpsB antibody in the input and output samples, possibly due to failure of heating to reverse crosslinking of GpsB monomers. The bottom band corresponds to GpsB monomer (≈13 kDa), whereas the top band is likely a GpsB dimer (≈26 kDa). The bands detected with anti-MreC or anti-aPBP1a in MreC-L-F3 or Pbp1a-F strains were F-tagged bait proteins. For detection of MreC-L-F3 or Pbp1a-F in output samples, 3 µl of samples were loaded to each lane, and proteins were detected with anti-MreC or anti-PBP1a. The relative amount of MreC was 5-9-fold higher in the input lysate of *rodZ*-L-F^3^-P_c_*erm* (IU4970; MreC row) compared to that from untagged WT strain (shown in adjacent lane) or lysate obtained from the markerless *rodZ*-F strain (IU14594, data not shown), suggesting that the P_c_ promoter present in the *rodZ*-L-F^3^-P_c_*erm* construct leads to overexpression of downstream genes, including *mreC*. Nevertheless, the Co-IP results using the *rodZ*-F markerless strain with no antibiotics cassette (IU14594) were similar to those for the *rodZ*-L-F^3^-P_c-_*erm* strain (IU4970) (data not shown). Co-IP experiments were performed 2-6 times with similar results (See Table 2 for quantitation). (B) Interaction map of RodZ in cells detected by co-IP. (C) Proteins that were weakly or not detected in complex with RodZ.

**Table 3.**
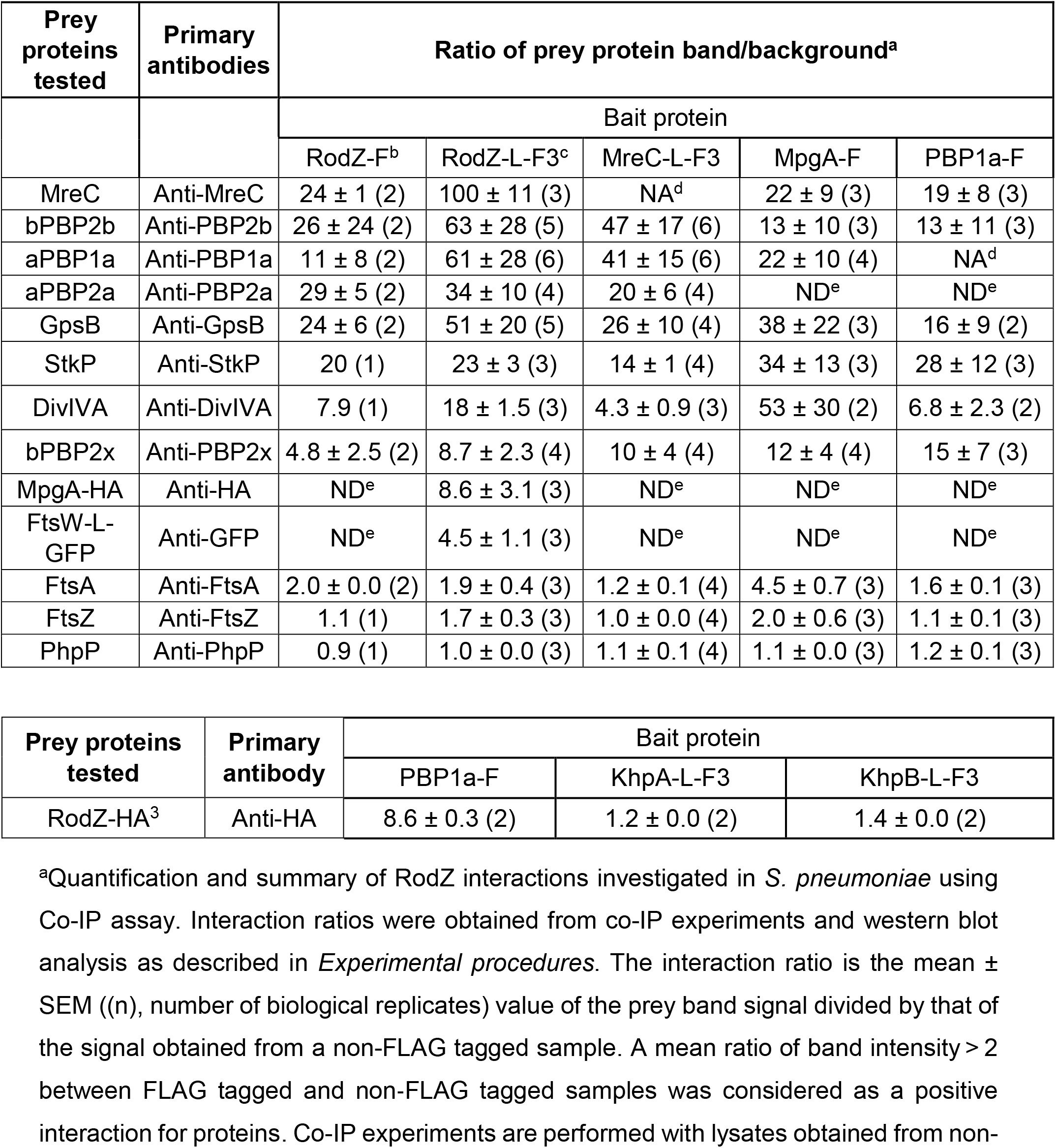
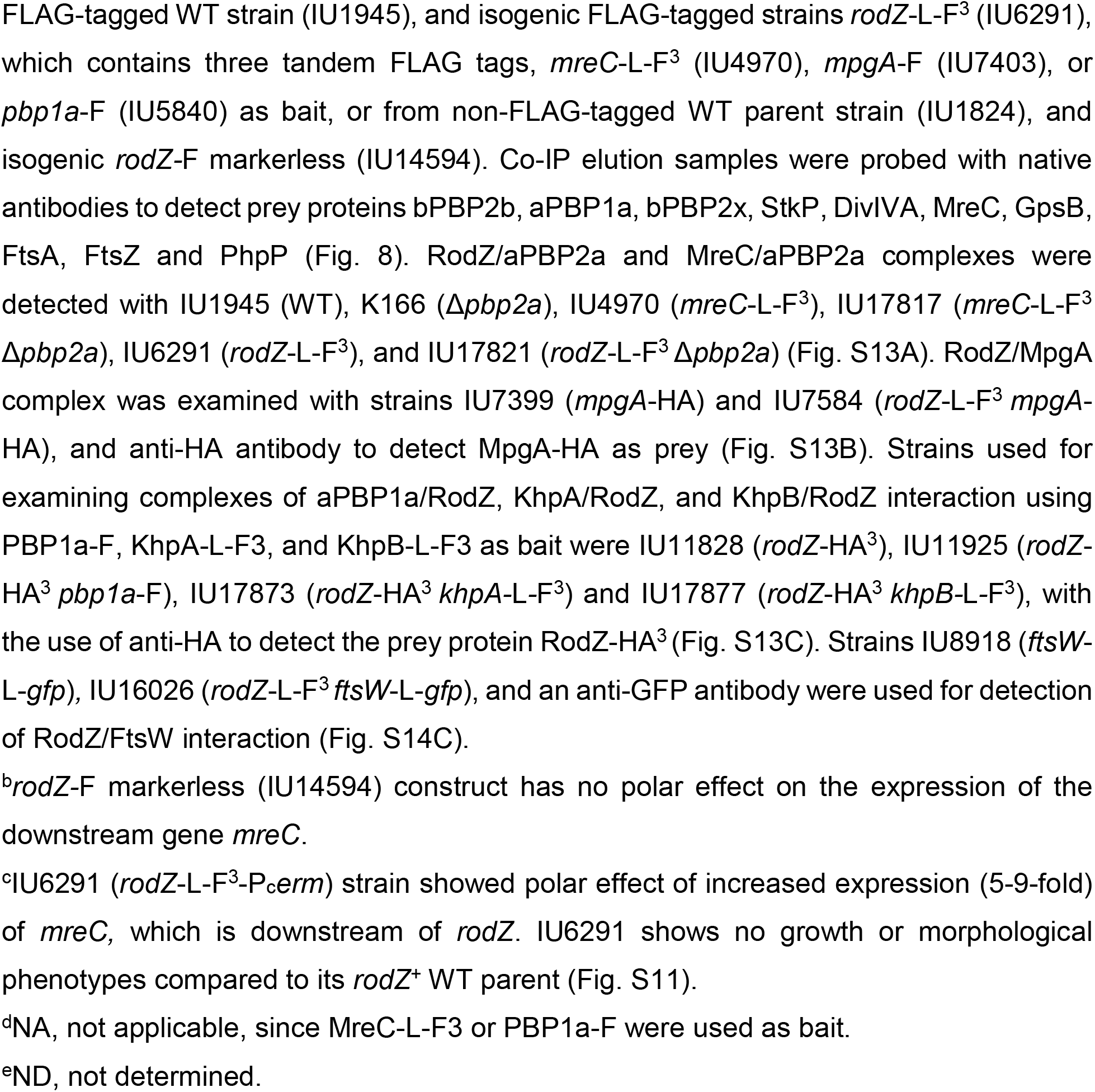
RodZ, MreC, MpgA, and aPBP1a are in complexes with key components of the peripheral and septal PG synthesis machines and division regulators StkP, GpsB, and DivIVA

B2H assays were performed in *E. coli* to test for direct interactions between WT RodZ (*Spn*) or truncated constructs lacking the HTH or DUF domain of RodZ(*Spn*) and the set of proteins mentioned above, as well as additional proteins not analyzed by co-IP (Fig. 9 and S15; Table S7). B2H assays revealed RodZ(*Spn*) self-interaction and a strong signal of interactions, usually bidirectional, between RodZ(*Spn*) and GpsB, MreC, MreD, MpgA, bPBP2b, RodA, aPBP1a, aPBP2a, bPBP2x, FtsW, EzrA, DivIVA, or aPBP1b (Fig. 2D, 8C, and 9A). Weaker signals of unidirectional interaction or no interaction were detected by B2H between RodZ(*Spn*) and StkP, FtsA, or FtsZ (Fig. 9A). For comparison, B2H assays were performed to determine direct interactions of pPG elongasome proteins MreC(*Spn*) or MreD(*Spn*) with Class A PBPs. MreC interacts with itself and shows bidirectional interactions with aPBP1a, aPBP2a, or aPBP1b, whereas MreD also self-interacts and shows bidirectional interactions with aPBP1a, but unidirectional interactions with aPBP2a or aPBP1b (Fig. 2D and 9B).

**Figure 9.**
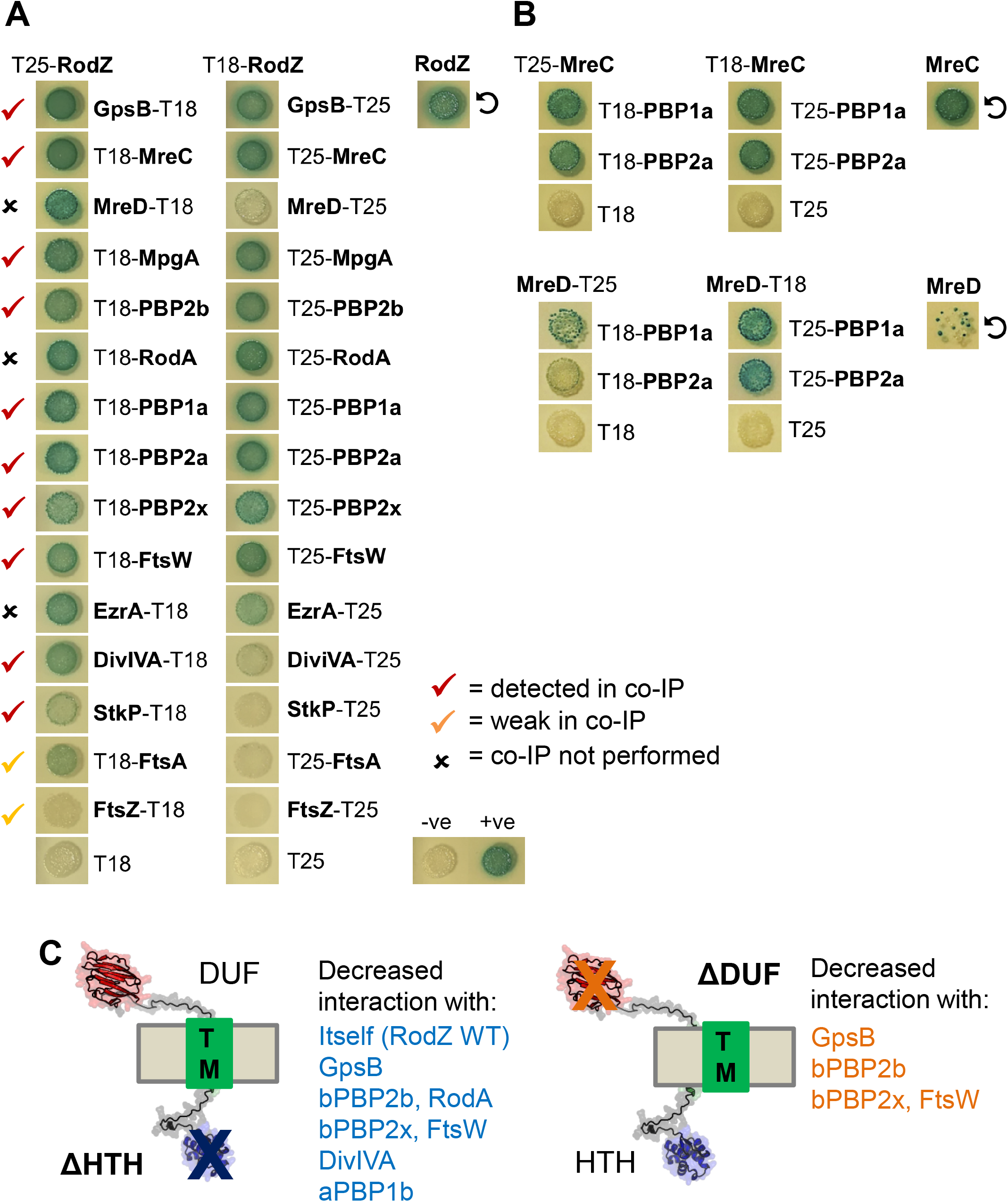
RodZ interacts with numerous cell elongation and division proteins as well as with itself in B2H assays. (A) RodZ interacts with GpsB, MreC, MreD, MpgA, bPBP2b, RodA, aPBP1a, aPBP2a, bPBP2x, FtsW, EzrA, and DivIVA in both directions, and with StkP and FtsA with a lower signal and only in one direction. Also, RodZ self-interaction is shown. Agar plates were photographed after 40 h at 30°C. See Figure S15 for earlier time points at 24, 30 and 36 h. (B) MreC and MreD interact with aPBP1a and aPBP2a and also self-interact. Agar plates were photographed after 40 h at 30°C. The punctate appearance of the spot showing MreD self-interaction is likely due to high toxicity of the *S. pneumoniae mreD* hybrid constructs in *E. coli.* (C) Summary of decreased interactions of *Spn* RodZ(ΔHTH) and RodZ(ΔDUF) compared to RodZ WT with certain PG synthesis and division proteins in B2H assays. Data are shown in Figure S15.

Finally, B2H assays were used to determine whether the absence of the HTH or DUF domain reduces binding to the above-mentioned proteins. Compared to WT RodZ(*Spn*), the absence of the HTH or DUF domain does not completely abolish the interactions between the truncated RodZ variants and any of the numerous partners tested (Fig. S15; Table S7). For many of them, including MreC, MreD, MpgA, aPBP2a, and EzrA, the signal was unchanged compared to WT RodZ(*Spn*) at the endpoint of the assay (data not shown). Yet, the absence of the HTH domain significantly reduces the interactions of RodZ(*Spn*) with itself and several proteins, including GpsB, bPBP2b, RodA, bPBP2x, FtsW, DivIVA, and aPBP1b (Fig. 9C and S15; Table S7). Although not essential in *S. pneumoniae* (Fig. 2 and 6; Table 2), the absence of the extracellular DUF domain also reduces the interactions with GpsB, bPBP2b, bPBP2x, or FtsW in the B2H assay (Fig. 9C and S15; Table S7). Together, we conclude that RodZ(*Spn*) is in complexes with numerous pPG elongasome proteins, PG synthesis regulatory proteins, and a few sPG synthesis proteins, possibly through direct interactions in some cases. Implications of these complexes to pneumococcal PG synthesis and division are considered further in *Discussion*.

### 2.7 Depletion of RodZ(*Spn*) mislocalizes MreC, bPBP2b, and RodA, but not other pPG and sPG synthesis proteins

We tested the hypothesis that RodZ(*Spn*) organizes the assembly of pPG elongasome proteins. We first determined whether incorporation of a fluorescent D-amino acid (FDAA) changes in a *rodZ*(ΔDUF), *rodZ*(ΔHTH), or Δ*rodZ* merodiploid mutant after ectopically expressed WT RodZ is depleted (Fig. S16). FDAA incorporation indicates regions of active PBP transpeptidase activity during PG synthesis (Boersma *et al*., 2015, Tsui *et al*., 2014), but does not distinguish between sPG and pPG synthesis at the midcell of *S. pneumoniae* cells (Perez *et al*., 2021a). As expected, FDAA labeling in Δ*rodZ*(DUF) cells depleted for RodZ (-Zn) is the same as that of cells expressing RodZ (+Zn or WT). FDAA is also similar in Δ*rodZ*(HTH) or Δ*rodZ* cells depleted of RodZ (-Zn inducer), although the RodZ depletion changes the cell size and morphology (Fig. S16). Results presented next show that RodZ depletion disrupts normal localization of MreC and the bPBP2b:RodA pPG synthase. Therefore, we interpret the FDAA labeling at midcell and equators of Δ*rodZ*(HTH) or Δ*rodZ* cells depleted of RodZ to reflect sPG synthesis, which is not disrupted by RodZ depletion. We conclude that RodZ depletion does not lead to widespread mislocalization of sPG synthesis, as occurs upon FtsZ, FtsA, or EzrA depletion (Mura *et al*., 2017, Perez *et al*., 2021b).

We next constructed Δ*rodZ*//P_Zn_-*rodZ^+^* merodiploid strains expressing from native chromosomal loci twelve other PG synthesis and division proteins fused to epitope tags, fluorescent reporter proteins, or a HaloTag (HT) (Fig S17-S22). Apart from three exceptions, we did not observe pronounced fusion-associated phenotypes that suppressed or exacerbated growth defects upon RodZ depletion (-Zn) in these strains. A sfGFP-MpgA fusion suppressed Δ*rodZ* lethality (Fig. S18A and S18B), likely because of reduced MpgA enzymatic activity, which is known to bypass the requirement for the pPG elongasome in *S. pneumoniae* (Taguchi *et al*., 2021, Tsui *et al*., 2016). Conversely, GFP-MpgA or HT-bPBP2x fusion exacerbated the drop in OD_620_ upon RodZ depletion, without overtly changing localization of the fusion proteins after 4 h of RodZ depletion (Fig. S18C and S22A).

Of the twelve proteins tested, aberrant localization upon RodZ depletion was only observed for MreC (Fig. 10A, 11, and S21), bPBP2b (Fig. 10B, S21, and S22B), and RodA (Fig. S22C) (summarized in Fig. 12). Mislocalization of MreC, bPBP2b, and RodA upon RodZ depletion was demonstrated by demographic analysis (Fig. 11, S21, and S22) and confirmed independently by IFM for MreC and bPBP2b (Fig. 10). By contrast, MpgA (Fig. S18) and aPBP1a (Fig. S20) (pPG synthesis); bPBP2x (Fig. S19 and S22D) (sPG synthesis); FtsZ (Fig. S19), MapZ (Fig. S19), EzrA (Fig. S19), and FtsA (Fig. S20) (Z-ring organization); and StkP (Fig. S20) and DivIVA (Fig. S19) (pPG and sPG synthesis) localize normally at midcell upon RodZ depletion (Fig. 12) (for functions, see (Briggs *et al*., 2021, Massidda *et al*., 2013, Straume *et al*., 2021)). We conclude that RodZ(*Spn*) is required for normal assembly and localization of MreC, bPBP2b, and RodA in the pPG elongasome.

**Figure 10.**
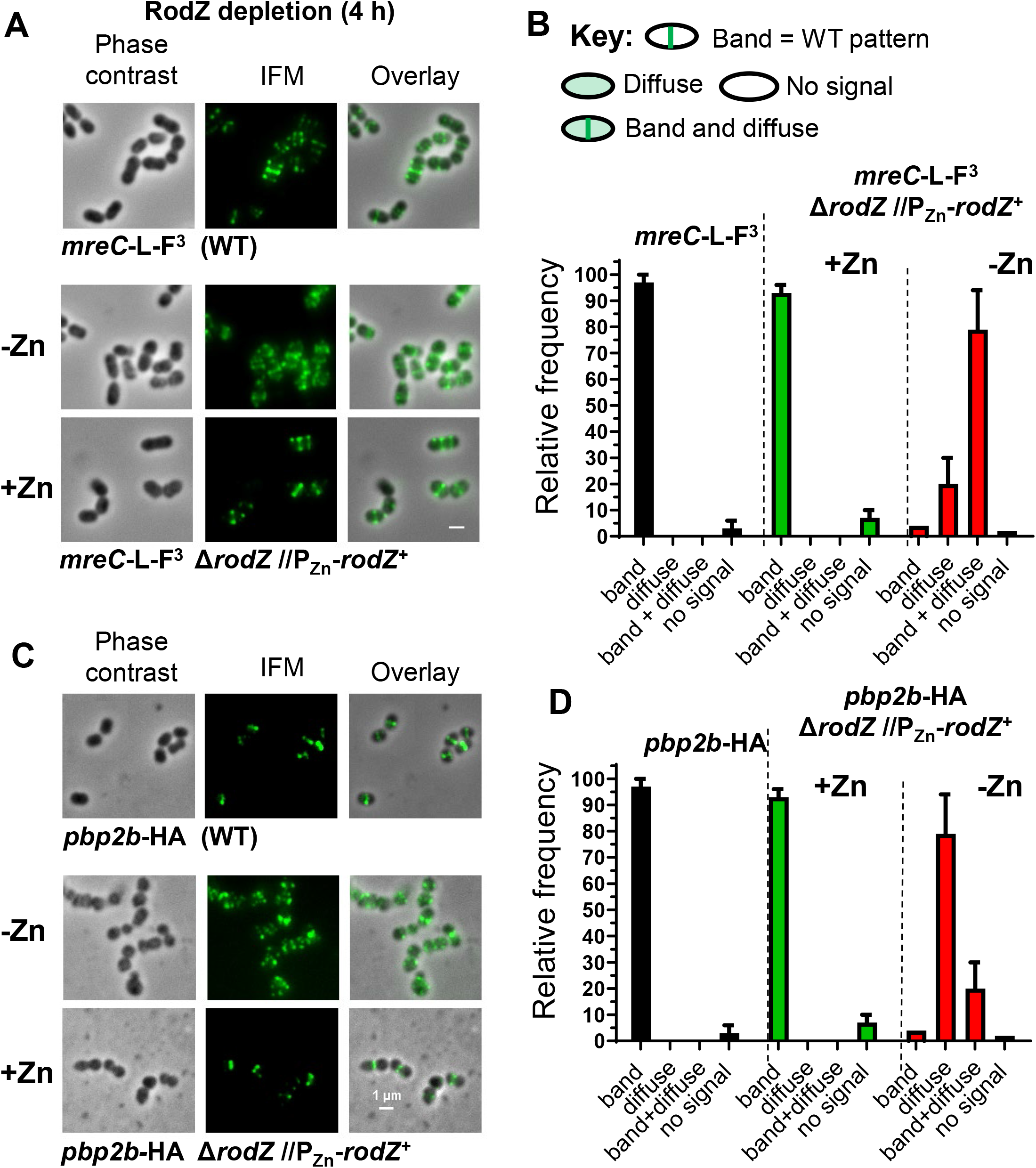
Depletion of RodZ leads to the mislocalization of MreC and bPBP2b detected by IFM. Representative images showing localization of MreC (A) or bPbp2b (C) after depletion of RodZ for 4 h of growth, which reduced RodZ to an undetectable amount (Fig. 3C). Phase contrast and 2D IFM was performed as described in *Experimental procedures* using antibody to the FLAG or HA tags. Strains used: (A) WT IU14458 (*mreC*-L-F^3^) and merodiploid strain IU14158 (*mreC*-L-F^3^ Δ*rodZ*//P_Zn_*-rodZ*^+^); (C): WT IU14455 (*pbp2B*-HA) and merodiploid strain IU14131 (*pbp2B*-HA Δ*rodZ*//P_Zn_*-rodZ*^+^). Quantification of localization patterns of MreC (B) and bPBP2b (D) observed at 4h in the WT and after RodZ depletion. For each sample and condition, 100 cells were manually examined and classified from two independent experiments.

**Figure 11.**
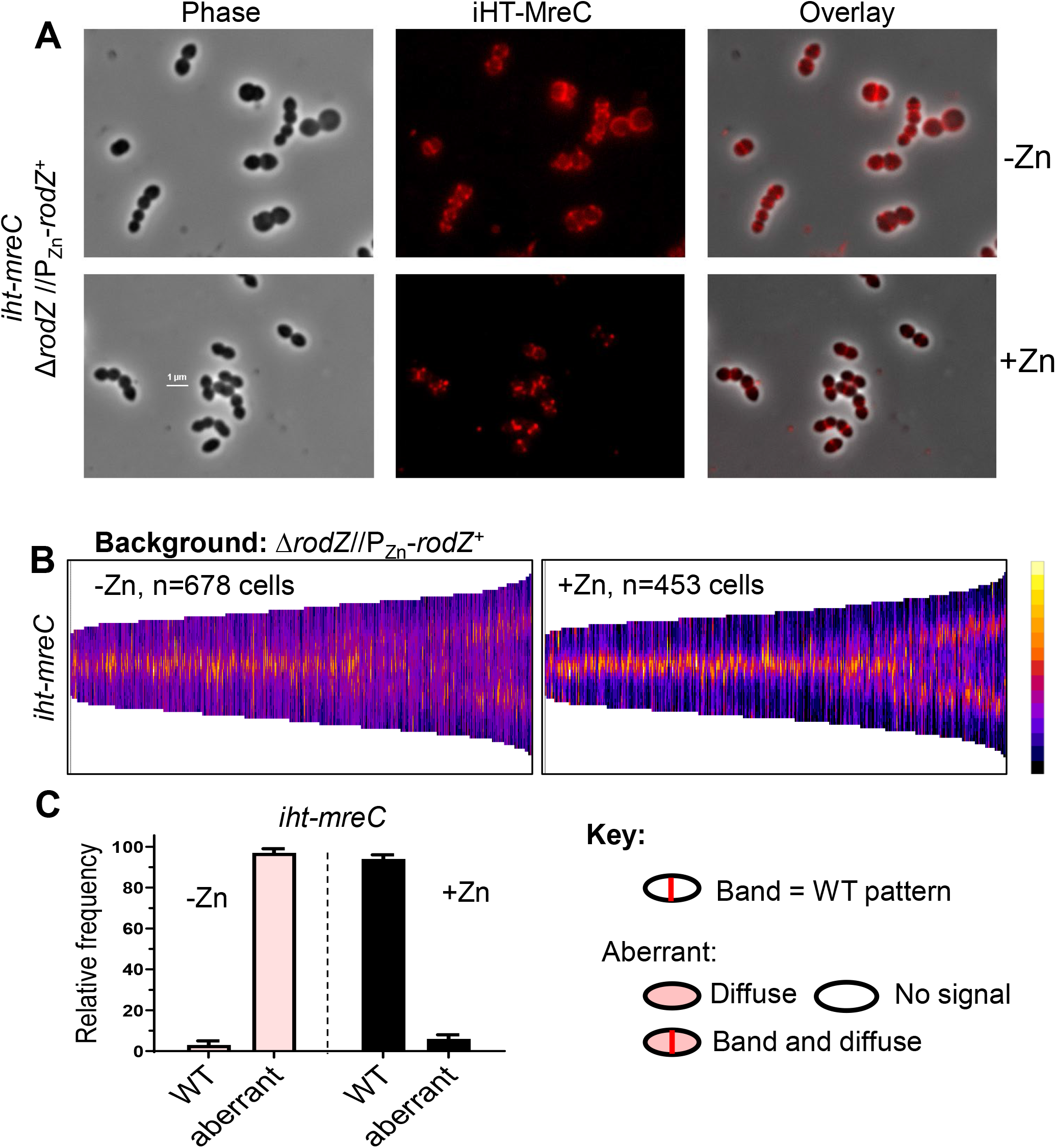
Depletion of RodZ leads to the mislocalization of MreC detected by 2D-FM. IU16920 (*iht-mreC* Δ*rodZ*//P_Zn_-*rodZ*^+^) was grown overnight in the presence of Zn inducer (0.4 mM ZnCl_2_ + 0.04 mM MnSO_4_) and diluted into fresh medium to OD_620_ ≈0.003 containing (complementation) or lacking (depletion) Zn inducer. At 4 h, localization of iHT-MreC was evaluated following saturating labeling of the iHT domain with TMR ligand by 2D-FM as described in *Experimental procedures*. (A) Representative micrographs showing iHT-MreC localization. (B) Demographs displaying fluorescence intensity of iHT-MreC localization in the absence (-Zn) or presence (+Zn) of RodZ for the number of cells (n) aligned and displayed in each demograph. Microscopy and demographs are representative of 3 independent biological replicates. (C) Bar graph displaying iHT-MreC localization patterns in micrographs of 100 cells from two independent experiments for each condition. The key illustrates criteria used to classify localization in the bar graphs.

**Figure 12.**
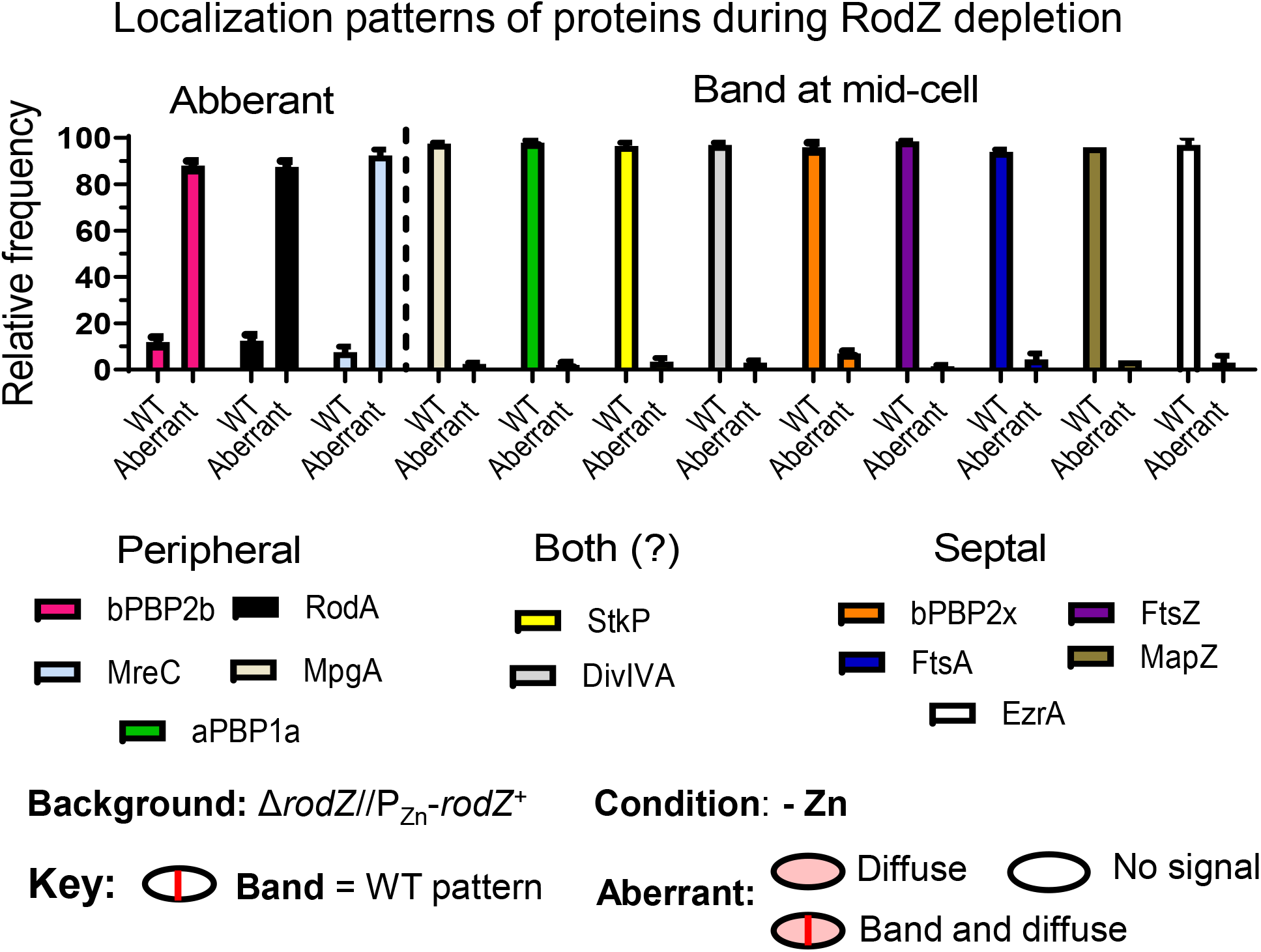
Localization patterns of PG synthesis and division proteins after RodZ depletion (-Zn) for 4 h. Among the peripheral PG synthesis machine components, the morphogenic protein MreC, the PG synthases bPBP2b (TPase) and RodA (GTase) require RodZ for localization, while the localization of MpgA (formerly MltG(*Spn*) muramidase and Class A PBP1a were unchanged by RodZ depletion. MreC, bPBP2b, and RodA localized normally in the presence of Zn inducer (Figure 10, Figure 11, Figure S22). Localization of cell division components (bPBP2x, FtsZ, FtsA, MapZ and EzrA), as well as StkP and DivIVA involved possibly with both septal and peripheral PG synthesis, were unaffected by RodZ depletion. Representative micrographs of localization studies are shown in Figure 10, Figure 11, Figure S18, Figure S19, Figure S20 and Figure S22. 100 cells were categorized by eye within a given field according to the key and averages with SEMs are indicated. Similar patterns were observed in two or more biological replicates. Strains used: IU16058 (*iht-pbp2b*), IU16060 (*iht-rodA*), IU16920 (*iht-mreC*), IU14433 (*gfp-mpgA*), IU14496 (*pbp1a-*FLAg), IU14160 (*stkP-*FLAG^2^), IU12993 (*ftsZ-sfgfp*), IU13061 (*divIVA-gfp*), IU13062 (*gfp-mapZ*), IU13058 (*ezrA-sfgfp*), IU13000 (*isfgfp-pbp2x*), and IU17022 (FLAG*-ftsA*) in the Δ*rodZ*//P_Zn_-*rodZ*^+^ background (see Table S1).

To test this notion and further establish the assembly hierarchy, we determined protein localization upon MreC depletion. We first established that Zn inducer (0.4 mM ZnCl_2_ + 0.4 mM MnSO_4_) does not affect growth or MreC amount in WT cells (Fig. S23). We then showed that MreC depletion for 4 h in a Δ*mreC*//P_Zn_-*mreC^+^* merodiploid strain reduces MreC cellular amount to ≈10% of WT, but does not alter bPBP2b or bPBP2x cellular amount (Fig. S24). Depletion of MreC resulted in mislocalization of bPBP2b detected by demographic analysis of HaloTag-bPBP2b (Fig. 13) and by IFM (Fig. S25C). Likewise, demographic analysis showed that RodA mislocalizes upon MreC depletion (Fig. S26B). In contrast, RodZ (Fig. S25A), aPBP1a (Fig. S25B), and bPBP2x (Fig. S26C) remained at their normal midcell positions upon MreC depletion. Together, these results support an assembly hierarchy wherein RodZ is required for MreC midcell localization, which in turn, is required for midcell localization of bPBP2b and RodA (Fig. 14).

**Figure 13.**
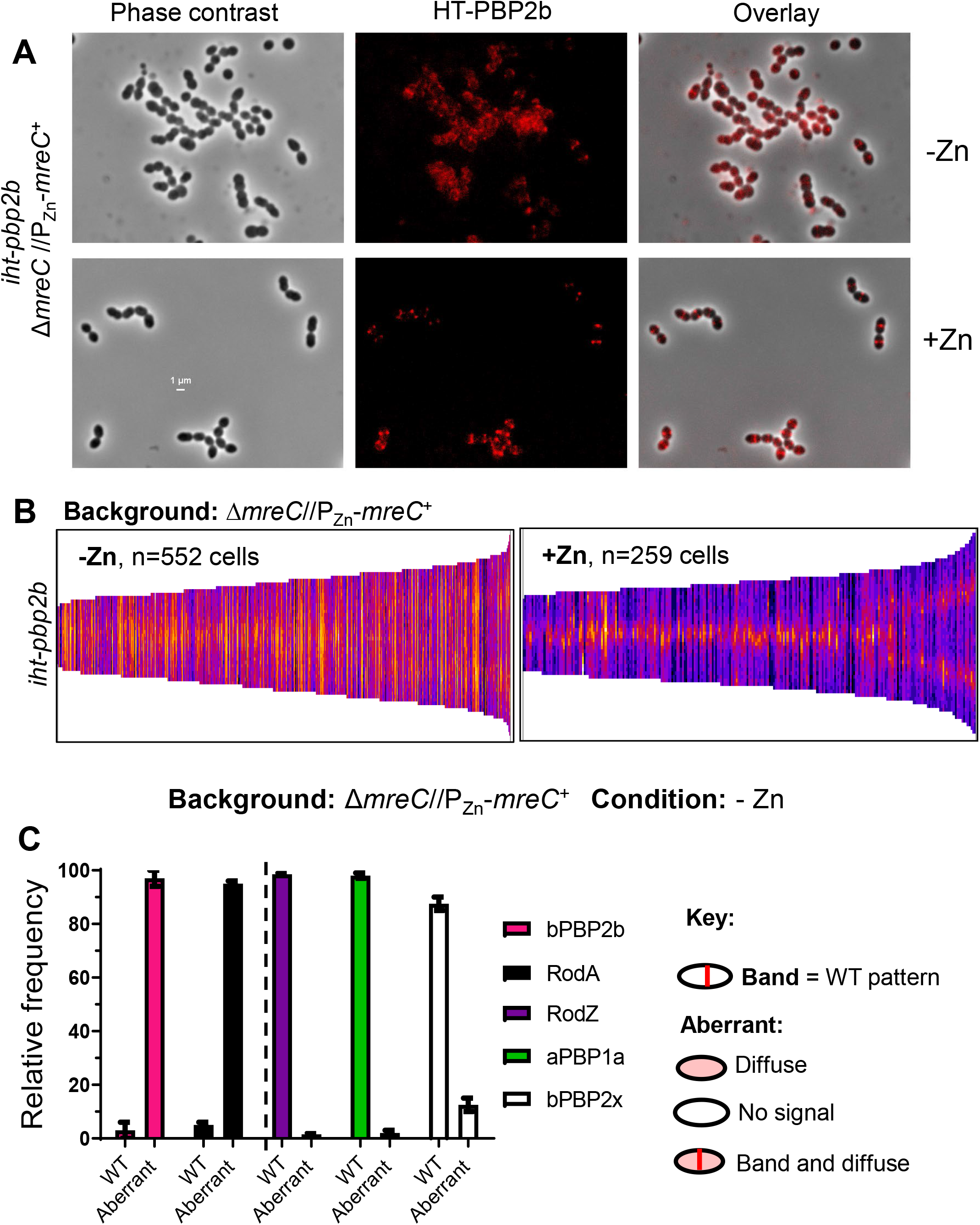
Depletion of MreC leads to mislocalization of bPBP2b and RodA, but not RodZ. For localization of bPBP2b, IU16281 (*iht-pbp2b* Δ*mreC* //P_Zn_*-mreC*^+^) was grown overnight in the presence of Zn inducer (0.4 mMCl_2_ Zn + 0.04 mM MnSO_4_) and diluted into fresh medium containing (complementation) or lacking (depletion) Zn inducer to OD_620_ ≈0.003. After 4 h, iHT-PBP2b was labeled with a saturating concentration of a TMR ligand, and localized in cells by 2D-FM as described in *Experimental procedures*. (A) Representative micrographs of iHT-PBP2b localization under MreC complementation or depletion conditions. (B) Demographs displaying fluorescence intensity of iHT-PBP2b localization upon MreC depletion (-Zn) or in the presence of MreC (+Zn) for the number of cells (n) aligned and displayed in each demograph. Microscopy and demographs are representative of 3 independent biological replicates. (C) Bar graph displaying localization patterns of bPBP2b, RodA, RodZ, aPBP1a, and bPBP2x after MreC depletion (-Zn) in micrographs of 100 cells from two independent experiments for each strain. Averages with SEMs are indicated in the bar graph. The key illustrates criteria used to classify localization in the bar graphs. Strains used: IU16281 (*iht-pbp2b*), IU16283 (*iht-rodA*), IU14598 (*rodZ-*FLAG), IU15901 (*pbp1a-*FLAG), and IU16326 (*iht-pbp2x*) in the Δ*mreC*//P_Zn_-*mreC*^+^ background (see Table S1). Representative micrographs of proteins other than bPBP2b are in Figures S21, S25, and S26.

**Figure 14.**
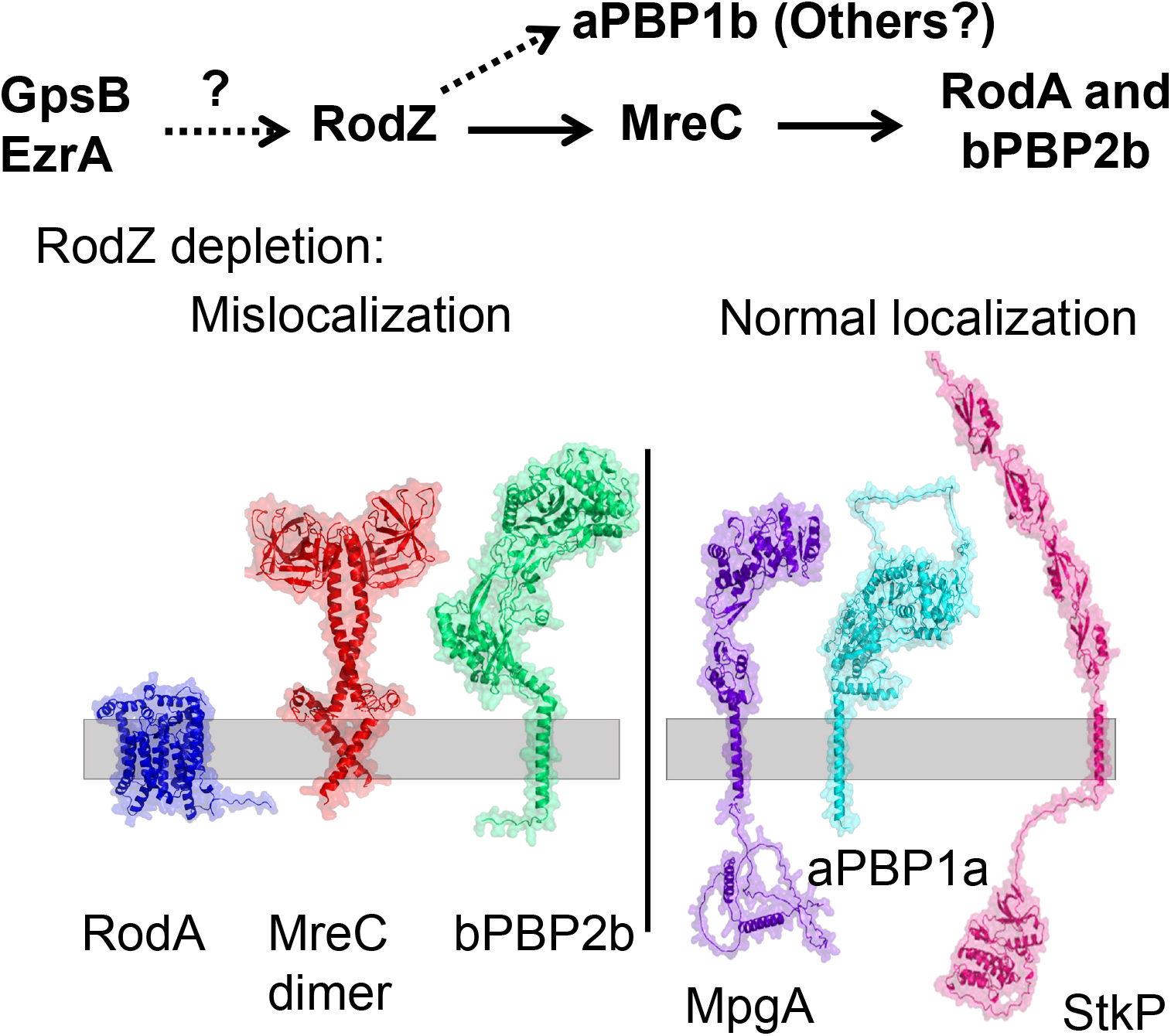
Assembly hierarchy of the pPG elongasome mediated by RodZ in *S. pneumoniae*. Results presented here establish RodZ(*Spn*) as an essential scaffolding protein required for the assembly and function of the pPG elongasome. The assembly hierarchy is based on RodZ depletion experiments, protein interaction assays, and genetic relationships described in *Results*. Depletion of RodZ leads to mislocalization of bPBP2b, RodA, and MreC, which are members of the core pPG elongasome, but not aPBP1a, StkP, FtsA, PBP2x, or MpgA (formerly MltG(*Spn*)). In turn, depletion of MreC leads to mislocalization of bPBP2b and RodA, but not RodZ or bPBP2x. Hence, depletion of RodZ results in incomplete assembly of the pPG elongasome. Predicted structures are illustrated without showing interactions for RodA, MreC dimer, and bPBP2b that mislocalize upon RodZ depletion and some of the proteins (MpgA, aPBP1a and StkP) that do not mislocalize. Structures were predicted using AlphaFold2 (Jumper *et al*., 2021). A synthetic-viable genetic relationship between RodZ(*Spn*) and aPBP1b and interaction experiments described in *Results* indicate that aPBP1b plays a role in pPG elongasome regulation and possibly in pPG synthesis. Finally, interaction experiments show that RodZ(*Spn*) interacts with GpsB and EzrA, which have been proposed to play roles in the interface between cell division and PG synthesis in *S. pneumoniae* (Cleverley *et al*., 2019, Perez *et al*., 2021b, Rued *et al*., 2017). See text for additional details.

### 2.8 RodZ(*Spn*), but not MreCD(*Spn*), displays a synthetic-viable genetic relationship with aPBP1b

Tn-seq analysis indicates the essentiality of the cytoplasmic N-terminal HTH and TM domains of RodZ(*Spn*) (above; Fig. 2A, row 1) and confirmed the essentiality of MreC(*Spn*) and MreD(*Spn*) (see above). Unexpectedly, Tn-seq analysis of a Δ*pbp1b* mutant, which lacks aPBP1b of unknown function, indicates suppression of *rodZ* essentiality (*i.e.*, insertions throughout the HTH and TM domains), but not *mreCD* essentiality (Fig. 2A, row 2). Previous results and those reported here show that Δ*pbp1a* suppresses the requirement for MreC, MreD, and RodZ (Table 1) (Fenton *et al*., 2016, Land & Winkler, 2011, Tsui *et al*., 2016). Likewise, Δ*khpA* and Δ*khpB* mutations, which result in the absence of the major KhpAB RNA-binding regulatory protein (Hor *et al*., 2020, Olejniczak *et al*., 2022), suppress Δ*mreCD* and Δ*rodZ* mutations (Table 1) (Zheng *et al*., 2017). These results are reiterated by Tn-seq analysis (Fig. 2A, row 3). Finally, in contrast to Δ*pbp1b* or Δ*pbp1a*, Tn-seq analysis shows that Δ*pbp2a*, which lacks aPBP2a, fails to suppress the essentiality of *mreC*, *mreD*, or *rodZ* in *S. pneumoniae* (Fig. 2A, row 4). We conclude that there is an unanticipated synthetic-viable genetic relationship between null mutations of *pbp1b* and *rodZ*, but not between *pbp1b* and *mreCD* (Fig. 15A). By contrast, there is a different synthetic-viable genetic relationship between null mutations of *pbp1a* and *mreC, mreD*, or *rodZ* (Fig. 15B; *Discussion*).

**Figure 15.**
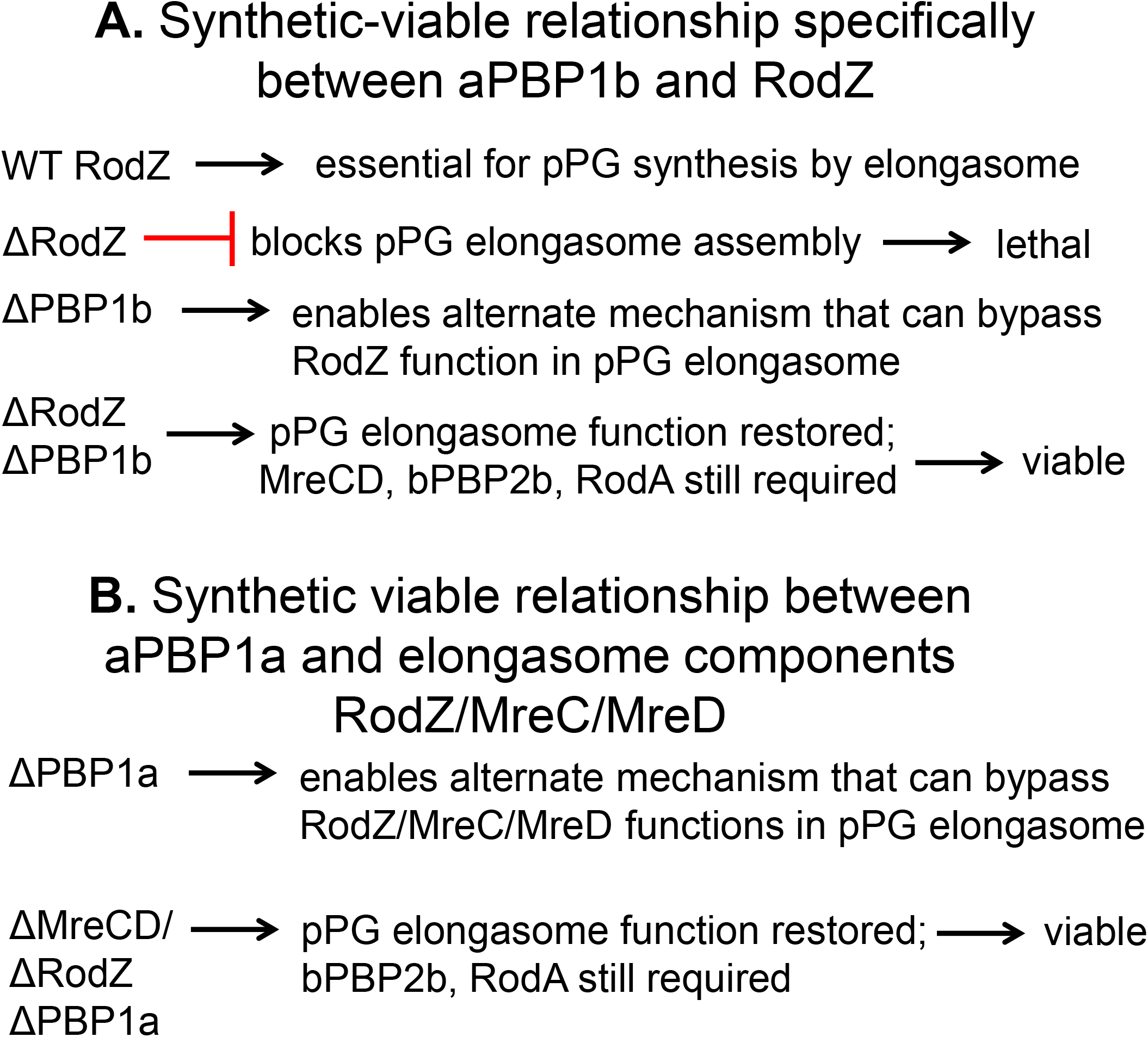
Model for bypass pPG synthesis to account for the synthetic-viable genetic relationships between aPBP1b and aPBP1a and components of the pPG elongasome of *S. pneumoniae*. **(A)** Δ*pbp1b* suppresses Δ*rodZ*, but not Δ*mreCD*, and MreCD bPBP2b, and RodA functions are still required for viability. **(B)** Δ*pbp1a* suppresses Δ*rodZ*, Δ*mreC*, or Δ*mreCD*, and bPBP2b and RodA function are still required for viability. A favored model postulates that some form of pPG synthesis is essential for pneumococcal viability because of the proposed role of pPG synthesis in positioning future equatorial Z-rings in daughter cells. In WT cells, the RodZ-MreCD-bPBP2b-RodA core elongasome carries out this pPG synthesis. However, there are also failsafe mechanisms that can bypass or modulate the function of the core pPG elongasome and restore division and growth. According to this model, Δ*pbp1b* enables alternate activities and/or interactions that can bypass the absence of RodZ (A), and Δ*pbp1a* enables alternate activities and/or interactions that can bypass the absence of RodZ, MreC, and MreD (B). See text for additional details and for alternative models for these synthetic-viable phenotypes.

We confirmed these synthetic-viable relationships detected by Tn-seq by independent transformation assays, in which Δ*rodZ* or Δ*mreCD* amplicons were transformed into deletion mutants of recipient strains (Fig 2B and 2C). Consistent with the Tn-seq results, Δ*pbp1b* suppresses Δ*rodZ*, but not Δ*mreCD,* Δ*pbp1a* or Δ*khpB* suppresses Δ*rodZ* or Δ*mreCD*, and Δ*pbp2a* does not suppress Δ*rodZ* or Δ*mreCD* (Fig. 2B and 2C). This synthetic-viable genetic relationship between aPBP1b and RodZ could reflect a direct interaction between the proteins. To test this idea, we performed B2H assays. Indeed, aPBP1b interacts with itself and shows bidirectional interactions with RodZ, MreC, or aPBP1a (Fig. 2D), as well as unidirectional interactions with MreD, MpgA, aPBP2a, or EzrA. Altogether, these results implicate aPBP1b in the regulation of the pPG elongasome, possibly through direct interaction with RodZ, as discussed below.

## 3 DISCUSSION

The essentiality of RodZ(*Spn*) has been controversial (Stamsas *et al*., 2017, Straume *et al*., 2017, Tsui *et al*., 2016). Δ*rodZ* mutants can be constructed in laboratory strain R6 and its derivatives (Martin-Galiano *et al*., 2014, Straume *et al*., 2017); however, these R6-derived strains contain dozens of mutations compared to progenitor D39 strain, including mutations in *pbp1a* (Land & Winkler, 2011, Lanie *et al*., 2007). Moreover, R6-derived strains have significant differences in shape and the timing of cell division compared to progenitor D39 strains (Trouve *et al*., 2021), indicating that R6-derived strains are not an exact model for mechanisms of PG synthesis and cell division that occur in their virulent progenitor D39 strain. Yet, *rodZ* has also been classified as non-essential in several Tn-seq studies of different *S. pneumoniae* serotype strains (Slager *et al*., 2018, van Opijnen & Camilli, 2012). Tn-seq and deletion analyses (Fig. 2 and 6) presented here show that the entire extracellular domain corresponding to about 48% of *rodZ*(*Spn*) is dispensable, which may account for these previous misclassifications. The essentiality of *rodZ* and other members of the pPG elongasome, including *mreC, mreD*, *pbp2b*, and *rodA*, was confirmed by transformation assays of unencapsulated (Δ*cps*) *S. pneumoniae* D39 grown at 37°C (Table 1; Fig. S2C). *rodZ* was also confirmed to be essential by transformation assays of encapsulated D39 strains (Table 1) and by depletion assays of unencapsulated strains (Fig. 3-5). Analogous to the case in *E. coli* (Bendezu & de Boer, 2008), this essentiality is conditional in unencapsulated D39 strains. Mutants lacking components of the pneumococcal pPG elongasome (MreCD, bPBP2b, RodA, or RodZ) grow poorly at the lower temperature of 32°C under the growth conditions tested (Fig. S2). This residual growth at 32°C, which is close to the that of the nasopharynx, is likely not of physiological significance *in vivo*, because insertions in *mreC*, *pbp2b,* or *rodA* are not recovered in Tn-seq screens of a murine nasal colonization model (van Opijnen & Camilli, 2012).

In rod-shaped bacteria, RodZ acts as the linker between cytoplasmic MreB filaments and the largely extracellular Rod complex containing regulators MreC and MreD and the PG synthase complex consisting of an essential bPBP and SEDS-protein RodA (*Introduction*) (reviewed in (Rohs & Bernhardt, 2021)). Despite the absence of MreB, the overall structure of RodZ(*Spn*) is remarkably similar to that of RodZ in rod-shaped bacteria that have MreB (Fig. 1 and S1) (Ago & Shiomi, 2019, Alyahya *et al*., 2009, Bendezu *et al*., 2009, Shiomi *et al*., 2008). Nevertheless, the TM and cytoplasmic domains, including the HTH domain, are essential for RodZ(*Spn)* function (Fig. 2 and 6; Table 2), similar to their importance in rod-shaped bacteria (Alyahya *et al*., 2009, Bendezu *et al*., 2009, Morgenstein *et al*., 2015, Shiomi *et al*., 2008). In rod-shaped bacteria, the HTH domain of RodZ binds to MreB, and changes in amino acids in one helix (Fig. 1 and S9) alter this interaction (Bendezu *et al*., 2009, Morgenstein *et al*., 2015, van den Ent *et al*., 2010). However, amino acid changes at corresponding positions in RodZ(*Spn*) do not cause a detectable phenotype (Table 2; Fig. 6). In contrast to the HTH and TM domains, the RodZ(*Spn*) β-strand DUF domain and most of the extracellular unstructured linker (beyond amino acid 132) are dispensable for growth, although cell morphology is altered by truncations ending at amino acids 134 and 135 (Fig. 2 and 6; Table 2). Deletion of the RodZ(*Spn*) DUF domain lessens interactions with a couple of proteins in B2H assays, including bPBP2b (Fig. 9C and S15A); however, these interactions, if they exist in pneumococcal cells, are dispensable.

Several lines of evidence presented here strongly link RodZ(*Spn*) to the elongasome that carries out pPG synthesis at the middle of dividing pneumococcal cells (*Introduction*) (Briggs *et al*., 2021, Massidda *et al*., 2013, Perez *et al*., 2021a, Philippe *et al*., 2014, Tsui *et al*., 2014). Transformation assays show that Δ*rodZ*(*Spn*) is suppressed by the same mutations that suppress Δ*mreC*, including Δ*pbp1a*, *mpgA*(Y488D), Δ*khpA*, insertions in *khpB*, and overexpression of FtsA (Fig. 2; Table 1)) (Land & Winkler, 2011, Tsui *et al*., 2016, Zheng *et al*., 2017). MreC and MpgA are members of the pPG elongasome (Briggs *et al*., 2021, Fenton *et al*., 2016, Land & Winkler, 2011, Philippe *et al*., 2014, Straume *et al*., 2017, Tsui *et al*., 2016), and KhpAB and FtsA amounts regulate the requirement for pPG elongasome in *S. pneumoniae* (Zheng *et al*., 2017). Although CozE(*Spn*) also associates with proteins in the pPG elongasome (Fenton *et al*., 2016), it is neither essential, nor is the poor growth of Δ*cozE* mutants suppressed by Δ*pbp1a*, *mpgA*(Y488D), or other mutations that suppress Δ*mreC* and Δ*rodZ* in transformation assays (Table 1). These results suggest different functions of MreC and RodZ from CozE in pPG elongation under the conditions tested here.

In addition, throughout the cell cycle, RodZ(*Spn*) colocalizes with pPG elongasome members MreC and aPBP1a (Fig. 7), which are genetically linked to pPG synthesis (see below) (Briggs *et al*., 2021, Fenton *et al*., 2016, Land & Winkler, 2011, Massidda *et al*., 2013, Tsui *et al*., 2014). These colocalization experiments were performed by IFM of exponentially growing cells expressing C-terminal RodZ-Myc, RodZ-FLAG, or RodZ-L-FLAG^3^ fusions from the *rodZ* native locus. These constructs did not cause observable growth or morphology defects (Fig. S11 and S12). Attempts to construct numerous RodZ(*Spn*) N-terminal, C-terminal, or internal sandwich-fusions to different fluorescent-protein reporters were unsuccessful, because the fusions were lethal, caused cell morphology defects, and/or showed diffuse fluorescence over entire cells (data not shown). Finally, depletion experiments of RodZ(*Spn*) or MreC(*Spn*) demonstrate that RodZ is required for MreC, bPBP2b, and RodA localization, whereas MreC is required for bPBP2b and RodA localization, but not for RodZ localization (Fig. 10-14). Together, these transformation assays, colocalization, and assembly hierarchy results establish that RodZ is a member of the pPG elongasome of *S. pneumoniae*.

RodZ(*Spn*) acting as a scaffold for the MreC/bPBP2b/RodA complex is analogous to its function in rod-shaped bacteria, where an assembly hierarchy has not yet been reported (Ago & Shiomi, 2019, Bendezu *et al*., 2009, Morgenstein *et al*., 2015, Rohs & Bernhardt, 2021). However, there are some specific differences in phenotypes that may reflect the absence of an MreB-dependent mechanism in *S. pneumoniae*. Depletion of RodZ(*Spn*) is bacteriostatic, halts growth, and leads to cell rounding, increased size, and slight chaining (Fig. 3-5 and S4-S5). Nonetheless, cells depleted for RodZ(*Spn*) are more heterogenous in shape and form shorter chains than cells depleted of MreC(*Spn*) (Land & Winkler, 2011), bPBP2b(*Spn*) (Berg *et al*., 2013, Tsui *et al*., 2014), or FtsEX(*Spn*) (Sham *et al*., 2011, Sham *et al*., 2013), which also localizes with the pPG elongasome (Perez *et al*., 2021a). These results suggest that RodZ(*Spn*) may have some functions distinct from MreC(*Spn*), as discussed below for aBPB1b (see Fig. 14). In another example, activated mutants of bPBP2(*Eco*) largely bypass the requirement for regulators MreC, MreD, and RodZ, suggesting that MreB, bPBP2, and RodA form the core of the *E. coli* elongasome (Rohs *et al*., 2018). In contrast, corresponding amino acid changes in the pedestal region of bPBP2b(*Spn*) fail to suppress Δ*rodZ* mutations (Table S6), raising the possibility that bPBP2(*Eco*) activation is dependent on MreB. Other differences in phenotypes suggest functional differences between RodZ in *S. pneumoniae* and *E. coli.* Overexpression of FtsZ(*Spn*) does not suppress Δ*rodZ*(*Spn*) (Table 1), and overexpression of RodZ(*Spn*) does not alter cell shape (Fig. S6), in contrast to overexpression phenotypes reported in *E. coli* and *C. crescentus* (Alyahya *et al*., 2009, Bendezu *et al*., 2009, Shiomi *et al*., 2008).

To further define the role of RodZ(*Spn*) in pPG synthesis by the elongasome, we performed co-IP and B2H assays (summarized in Fig. 8 and 9). B2H and bimolecular fluorescence complementation assays and genetic approaches have indicated that RodZ of rod-shaped bacteria interacts with MreB and with other components of the Rod complex, including MreC, MreD, bPBP2, and RodA (Ago & Shiomi, 2019, Alyahya *et al*., 2009, Beilharz *et al*., 2012, Morgenstein *et al*., 2015, Shiomi *et al*., 2013, van den Ent *et al*., 2010). Likewise, RodZ(*Spn*) is found in complexes in unsynchronized pneumococcal cells with tested proteins bPBP2b, MreC, MpgA, and aPBP1a, which have been implicated in pPG elongation (Fig. 8; Table 3) (Briggs *et al*., 2021, Philippe *et al*., 2014, Straume *et al*., 2021). Direct interactions were detected in B2H assays between RodZ(*Spn*) and these proteins and with other elongasome proteins MreD and RodA (Fig. 9), which were not tested in co-IP assays (Fig 8). In addition, complexes and direct interactions were detected for RodZ(*Spn*) and other aPBPs (aPBP2a and aPBP1b), PG synthesis regulators (GpsB and DivIVA), the sPG synthase components (bPBP2x and FtsW), and EzrA (Fig. 2, 8, and 9; Table 3). Based on B2H assays, many of these interactions are mediated at some level by the cytoplasmic HTH domain of RodZ(*Spn*), but others are not (Fig. 9 and S15; Table S7). Phosphorylation of RodZ(*Spn*) by StkP was not detected by western blotting (Fig. S10) or by phosphoryl-proteomic analysis (Ulrych *et al*., 2021).

The functions and timing of these putative interactions of RodZ(*Spn*) with proteins outside of the canonical elongasome remain to be determined. Pneumococcal GpsB is found in complexes with EzrA, MreC, aPBP2a, bPBP2b, and StkP, which itself is detected in complexes with MreC, bPBP2b, and bPBP2x, at currently unknown stages of cell division (Cleverley *et al*., 2019, Rued *et al*., 2017). The interactions and phenotypes of *gpsB* mutants suggest a model wherein GpsB activates protein phosphorylation by StkP and also balances sPG and pPG synthesis at the midcell of dividing pneumococcal cells (Cleverley *et al*., 2019, Rued *et al*., 2017). Moreover, a low level of bPBP2x is detected in the outer midcell pPG synthesis ring of *S. pneumoniae* (*Introduction*) (Briggs *et al*., 2021, Perez *et al*., 2021a, Tsui *et al*., 2014). On the other hand, strong complexes containing RodZ(*Spn*) and FtsZ were not detected by co-IP (Fig. 8; Table 3) and no interactions between RodZ(*Spn*) and FtsZ were detected by B2H assays (Fig. 9), in contrast to what was reported in *E. coli* (Ago & Shiomi, 2019, Yoshii *et al*., 2019). In this regard, *S. pneumoniae* lacks a homolog of newly characterized ZapG, which interacts with elongasome proteins, including RodZ(*Eco*), and divisome proteins (Mehla *et al*., 2021). At best, only marginal complexes were detected containing RodZ(*Spn*) and FtsA, which is an actin homolog like MreB (Fig. 8). Altogether, the detected interactions and assembly patterns are consistent with RodZ(*Spn*) acting as a scaffold protein that may link GpsB and EzrA to the assembly and function of the pneumococcal pPG elongasome (Fig. 14).

A main finding from this study is the synthetic-viable genetic relationship specifically between RodZ and aPBP1b, but not between RodZ and MreCD in *S. pneumoniae* (Fig. 2 and 15). This is one of the first phenotypes that has been associated with pneumococcal aPBP1b, about which little is known (Briggs *et al*., 2021, Ducret & Grangeasse, 2017, Straume *et al*., 2021). The regulation and functions of Class A PBPs are not generally well understood in *S. pneumoniae* and other bacteria, but likely play roles during normal growth and stress conditions (Briggs *et al*., 2021, Pazos & Vollmer, 2021, Rohs & Bernhardt, 2021, Straume *et al*., 2021, Vigouroux *et al*., 2020). aPBP1a and aPBP2a have a synthetic-lethal relationship, such that Δ*pbp1a* Δ*pbp2a* mutants are inviable (Paik *et al*., 1999, Straume *et al*., 2021). In addition, aPBP1a was shown previously to have a synthetic-viable relationship with the elongasome components MreC, MreD, and RodZ, such that Δ*pbp1a* suppresses and allows growth of strains containing normally lethal Δ*mreC,* Δ*mreCD*, or Δ*rodZ* mutations (Fig. 2; Table 1) (Fenton *et al*., 2016, Land & Winkler, 2011, Tsui *et al*., 2016). Notably, Δ*pbp1a* does not suppress Δ*pbp2b* or Δ*rodA,* indicating that aPBP2b and RodA are still required for viability in the absence of MreCD, RodZ, and aPBP1a (Tsui *et al*., 2016). In contrast to aPBP1a, aPBP1b displays a synthetic-viable relationship only with RodZ, but not with MreCD, such that Δ*pbp1b* suppresses Δ*rodZ*, but does not suppress Δ*mreCD* (Fig. 2A). This synthetic viable relationship was confirmed by transformation assays (Fig. 2B and 2C), and B2H assays indicate direct interactions between aPBP1b and RodZ, MreC, or aPBP1a (Fig. 2D). Tn-seq data confirms that Δ*pbp1b* also does not suppress knock-out insertions in *pbp2b* or *rodA* (data not shown).

Different models can account for the synthetic-viable relationships of aPBP1b and aPBP1a with components of the pPG elongasome. Our favored model postulates that some form of pPG synthesis is essential for pneumococcal viability and that failsafe mechanisms can bypass or modulate the function of the core RodZ-MreCD-bPBP2b-RodA elongasome (Fig. 15). pPG synthesis may be essential, because it drives the composite MapZ/FtsZ/FtsA/EzrA nascent equatorial rings from midcell to the middle of daughter cells (Briggs *et al*., 2021, Fleurie *et al*., 2014, Holeckova *et al*., 2014, Perez *et al*., 2019). The absence of aPBP1b induces the pneumococcal WalRK two-component system (TCS) regulon (Zheng *et al*., 2017), which responds to cell wall stresses (Gutu *et al*., 2010, Tsui *et al*., 2016, Wayne *et al*., 2012). Induction of the WalRK TCS increases transcript amounts of genes encoding PG hydrolases and PG-binding proteins (Ng *et al*., 2005). Another possible change that may occur in the absence of aPBP1b is altered activity and/or interactions of aPBP1a, which associates with the core pPG elongasome and can provide alternate TP and GT activities (Fig. 8 and 9) (Briggs *et al*., 2021, Land & Winkler, 2011, Philippe *et al*., 2014, Tsui *et al*., 2016). In a Δ*rodZ* Δ*pbp1b* mutant, these alternate activities and/or interactions are proposed to bypass the defects in pPG elongasome assembly in the absence of RodZ and allow the aPBP2b:RodA PG synthase and perhaps aPBP1a to carry out sufficient pPG synthesis for division and growth (Fig. 15A).

The absence of aPBP1a also induces the WalRK regulon (Zheng *et al*., 2017), changes cell shape in culture (Land & Winkler, 2011, Tsui *et al*., 2016), and possibly alters the interactions and/or functions aPBP1b, which is associated with the pneumococcal pPG elongasome (Fig. 2). In a Δ*pbp1a* Δ*mreCD* Δ*rodZ* mutant, alternate activities and/or interactions would bypass the defects in pPG elongasome assembly in the absence of RodZ and MreCD and allow the bPBP2b:RodA PG synthase and perhaps aPBP1b to carry out sufficient pPG synthesis for division and growth (Fig. 15B). Another putative component of failsafe, bypass mechanisms for pPG synthesis is the sPG synthase bPBP2x:FtsW. A majority of bPBP2x migrates in the inner ring of sPG synthesis at the leading edge of the closing septal annulus (Briggs *et al*., 2021, Perez *et al*., 2021b, Tsui *et al*., 2014). However, some bPBP2x, and presumably FtsW, remains in the outer pPG synthesis ring and may provide an alternate pathway of pPG synthesis. Consistent with this notion, RodZ(*Spn*) associates with bPBP2x and FtsW (Fig. 8 and 9; Table 3). Finally, although aPBP2a also associates with the pPG elongasome (Fig. 8 and 9; Table 3), there is no genetic evidence that implicates aPBP2a in a bypass mechanism of pPG synthesis (Fig. 2), and Δ*pbp2a* does not induce the WalRK TCS (Zheng *et al*., 2017).

Other genetic patterns strongly support the idea of alternate, bypass mechanisms to maintain pPG synthesis in *S. pneumoniae.* The pPG elongasome-associated muramidase MpgA (formerly MLtG(*Spn*)) (Fig. 8 and 9) is essential; yet, Δ*mpgA* is suppressed by Δ*pbp1a* (Taguchi *et al*., 2021, Tsui *et al*., 2016). Furthermore, a triple Δ*pbp1a* Δ*mpgA* Δ*pbp2b* mutant lacks the core pPG elongasome, but is viable and forms elongated cells (Tsui *et al*., 2016). In addition, a *mpgA*(Y488D) mutant, which expresses an MpgA with greatly reduced enzymatic activity (Taguchi *et al*., 2021, Tsui *et al*., 2016), bypasses the requirement for the core pPG elongasome, in that *mpgA*(Y488D) Δ*pbp2b* and *mpgA*(Y488D) Δ*rodA* mutants are viable and form elongated cells (Tsui *et al*., 2016). Yet, the *mpgA*(Y488D) Δ*pbp2b* Δ*pbp1a* mutant is now inviable, which appears at odds with the phenotype of the Δ*pbp1a* Δ*mpgA* Δ*pbp2b* mutant. This apparent discrepancy can be explained if aPBP1b mediates pPG bypass synthesis in the Δ*pbp1a* Δ*mpgA* Δ*pbp2b* mutant, while aPBP1a mediates pPG bypass synthesis in the *mpgA*(Y488D) Δ*pbp2b* mutant, where aPBP1a bypass activity is dependent on the physical presence, but not the activity, of MpgA(Y488D). Finally, the shape and size of WT and Δ*pbp1b* cells grown in BHI broth are similar to those of Δ*pbp1b* Δ*rodZ* mutants (Fig. 2B and 2C), with some heterogeneity, whereas Δ*pbp1a* Δ*rodZ* cells have the distinctive shorter, narrower shape of Δ*pbp1a* mutant cells compared to WT (data not shown) (Land & Winkler, 2011, Tsui *et al*., 2016). These data indicate that suppression of Δ*rodZ* by Δ*pbp1b* or Δ*pbp1a* are not equivalent, consistent with different mechanisms. Taken together, these results support the hypothesis of multiple bypass pathways for essential pPG synthesis when the core pPG elongasome is incomplete or absent.

Another model for the synthetic-viable relationships of aPBP1b and aPBP1a with components of the pPG elongasome invokes direct regulation of aPBP activity, interactions, and/or localization (Fig. S27) (Land & Winkler, 2011, Tsui *et al*., 2016). In this model, RodZ(*Spn*) acts as a negative regulator of aPBP1b activity, interactions, and/or mislocalization. The absence of RodZ causes aPBP1b misregulation that contributes to cell lethality (Fig. S27A). Likewise, MreC, MreD, and RodZ would negatively regulate aPBP1a activity, interactions, and/or mislocalization. In the absence of MreC, MreD, or RodZ, aPBP1a misregulation contributes to cell lethality (Fig. S27B).

There are issues with this alternative model. First, to date, PBPs have been found to be positively regulated, rather than negatively regulated. For example, in *E. coli* the activities of FtsI(bPBP3), bPBP2, aPBP1a, and aPBP1b are positively activated by FtsN, MreC, LpoA, and LpoB, respectively (Pazos & Vollmer, 2021, Pichoff *et al*., 2019, Rohs & Bernhardt, 2021). In *S. pneumoniae*, aPBP2a is positively regulated by MacP and GpsB (Cleverley *et al*., 2019, Fenton *et al*., 2018). Second, aPBP misregulation alone is insufficient to account for cell lethality, because function of the bPBP2b:RodA pPG synthase is still essential (Fig. 15), despite suppression of Δ*rodZ* or Δ*mreCD* Δ*rodZ* by Δ*pbp1b* or Δ*pbp1a*, respectively (Fig. 2). Third, aPBP1a localizes normally when RodZ or MreC is depleted (Fig. 12 and 13C). This result contrasts with a previous conclusion that aPBP1a mislocalizes in the absence of MreC (Fenton *et al*., 2016). The different results may reflect the ectopic induction of potentially high levels of active GFP-aPBP1a in a Δ*mreC* mutant, which is not tolerated, as opposed to the MreC depletion used here in a strain expressing epitope-tagged aPBP1a-FLAG from its chromosomal locus (Fig. S25B). Moreover, moderate (≈2-fold) overexpression of aPBP1a in a WT strain does not overtly affect normal cell morphology or growth (Averi McFarland; unpublished result). Finally, because the synthetic-viable relationships of the aPBPs with components of the pPG elongasome are different, it is difficult to reconcile a model postulating that absence of aPBP1b or aPBP1a in the Δ*rodZ* or Δ*rodZ* Δ*mreCD* mutant, respectively, solely decreases competition or interference with residual bPBP2b:RodA pPG synthase activity. For this type of model to work, there still needs to be differential regulation of aPBP1b or bPBP1a expression, activity, and/or interactions allowing bypass pPG synthesis. Therefore, current data supports alternate mechanisms leading to pPG bypass synthesis more than other models for these synthetic-viable genetic relationships.

The different synthetic-viable relationships of aPBP1b and aPBP1a with components of the pPG elongasome indicates functional fungibility during pPG synthesis in *S. pneumoniae*. This flexibility points to altered protein interactions and/or regulatory pathways that enable alternate pPG synthesis pathways. Some of these outcomes may be through direct interactions, while others may be indirect through additional proteins induced or regulated by stress responses. Along the same lines, CozE does not have a strong synthetic viable relationship with aPBP1a in transformation assays or in Tn-seq analyses (data not shown) under the conditions used here, and Δ*cozE* was not suppressed by the array of mutations that suppress Δ*rodZ* and Δ*mreC* (Table 1). These shared and different functions and interactions of pPG elongasome members RodZ, MreC, MreD and CozE require further study at different stages of the pneumococcal cell cycle. Taken together, we conclude that both aPBP1a and aPBP1b play roles in the regulation of the pPG elongasome and possibly participate in pPG synthesis in *S. pneumoniae*. The action of RodZ(*Spn*) in assembly and function of the pPG elongasome and the roles of the aPBPs in WT and bypass pPG synthesis are important topics for future studies.

## 4 EXPERIMENTAL PROCDURES

### 4.1 Strain construction and growth conditions

Bacterial strains used in this study are derivatives of the unencapsulated, *S. pneumoniae* serotype 2 strain D39W (Lanie *et al*., 2007, Slager *et al*., 2018), and are listed in Table S1. For strains containing antibiotic markers, linear DNA amplicons synthesized by fusion PCR were transformed into competent pneumococcal cells as described in (Land *et al*., 2013, Tsui *et al*., 2010, Tsui *et al*., 2014). For antibiotic selection Trypticase soy agar II (modified; Becton-Dickinson) and 5% (vol/vol) defibrinated sheep blood (TSAII-BA) plates were supplemented with the following final concentrations of antibiotics: 250 μg kanamycin/mL, 150 μg spectinomycin/mL, 0.3 μg erythromycin/mL, 200 μg streptomycin/mL, or 0.25 μg tetracycline/mL. Strains containing markerless mutations or insertions at the native site in the chromosome, e.g. *iht-pbp2b* markerless, were constructed through two rounds of transformation via the Janus method, as described in (Sung *et al*., 2001). Linkers used in construction of fluorochrome or epitope-tagged fusion proteins are listed in Table S2. All primers and templates used in this study are listed in Table S4. All strains were confirmed via PCR and sequencing. For overnight growth, BHI broth was inoculated with frozen glycerol stocks, serially diluted and propagated overnight for 12-13 hours at 37°C in an atmosphere of 5% CO_2_. Antibiotics were not added to the media. To start experiments, overnight cultures with an OD_620_ of 0.1-0.4 (exponentially growing) were diluted to an OD_620_ of 0.003 in fresh BHI, lacking antibiotics.

### 4.2 Tn-seq transposon library generation and insertion sequencing

Tn-seq was carried out using protocols in (Fenton *et al*., 2016, van Opijnen *et al*., 2015) with the following modifications. A transposon insertion library was generated for each of the following strains: WT D39 Δ*cps rpsL1* (IU1824), isogenic Δ*pbp1b* (IU14697), Δ*khpB* (IU10592), and Δ*pbp2a* (IU13256). Approximately 200,000 (WT, Δ*pbp1b* and Δ*khpB*) or 300,000 (Δ*pbp2a*) transformants were pooled for each library (see Appendix A). Genomic DNA preparations were modified from the instructions provided by Qiagen for Gram-positive bacteria using a DNeasy blood and tissue kit (Qiagen 69504). 5 mL of cultures at OD_620_ =0.4 were centrifuged for 10 min at 5,000 x *g* at room temperature and suspended in 180 µL of enzymatic lysis buffer containing 20 mg/mL lysozyme, and incubated for 30 min at 37°C. 10 µL of RNase A (Qiagen 19101,100 mg/ml) was added, followed by a 5-min incubation at room temperature. Subsequent steps were as specified by the Qiagen manual, except that DNA was eluted with 100 µL of water. Eluted genomic DNA and pMagellan6 DNA prepared with a QIAprep Spin Miniprep kit (27104) were concentrated by ethanol precipitation to concentrations of more than 0.3 µg/µL in ultrapure distilled water. *In vitro* transposition reactions were performed as described previously (van Opijnen *et al*., 2015) with genomic DNA obtained from WT or mutant strains with the following modifications. A reaction mixture of 1 µg genomic DNA, 1 µg pMagellan6 DNA, and 3 μL of purified MarC9 transposase prepared as specified in (van Opijnen *et al*., 2015) was incubated at 30°C for 4h. Transposon junctions were repaired by using 1µL of 3 U/µL T4 DNA polymerase at 12°C for 30 min. All incubation steps were performed in a thermocycler. Ten independent 20 µL-transposition products were prepared each time and stored at −20°C. Starter cultures for transformation were prepared by growing frozen stocks of respective strains in 4 mL of BHI broth at 37°C with CO_2_ to OD_620_ ≈0.15. The cultures were centrifuged for 3 min at 16,000 × *g* at room temperature, and pellets were resuspended in 400 μL BHI broth mixed with 600 μL 25% glycerol. 50-μL aliquots of starter cultures were stored at −80°C.

Transposed DNA was transformed into CSP-1 induced competent WT or mutant strains and plated onto TSAII agar plates containing spectinomycin and catalase. On the day of transformation, recipient strains were grown from frozen starter cultures in 5 mL of BHI broth to OD_620_ ≈0.03-0.04. TSAII agar (BBL 212305) plates containing spectinomycin and catalase were prepared by pouring 17 mL of warm TSAII agar containing 150 μg/mL spectinomycin into each 100 x 15 mm plate. After solidification, 396 µL (13,000U to 15,000U) of catalase solution (Worthington, CAT # LS001896) were spread on the surface of each plate and dried for 30 min in a sterile hood. Transformation mixes were prepared by addition of 40 µL CSP-1 (50 ng/µL), 1 mL heat inactivated horse serum, 45 µL 40% glucose, to 9 mL BHI broth. 300 µL of cell culture at OD_620_ ≈0.03-0.04 were mixed with 700 µL of transformation mix. After incubation at 37°C for 10 min, 3-8 µL of respective transposed DNA were added to each transformation, and the mixtures were incubated at 37°C for 1 h. Transformations containing no DNA or 27 ng of genomic DNA obtained from IU2072 containing *spxR*::Mariner (Ramos-Montanez *et al*., 2008) were used as negative or positive controls. 200 µL of transformed cell culture were spread on the surface of each prepared TSAII/spectinomycin/catalase plate, which were incubated at 37°C in 5% CO_2_ for 20 h.

After 20 h of incubation, colonies were scraped from 50 plates and collected in 20 mL of BHI broth. Cell suspensions were centrifuged for 8 min at 3,000 x *g* at room temperature, and the pellets were resuspended in 3 mL BHI mixed with 2 mL 25% glycerol. From each transformation, twenty 250-µL of transposon library starter culture aliquots were stored at −80°C. For WT, Δ*pbp1b*, Δ*khpB* strains, transposon library starter cultures were obtained from ≈200,000 colonies with 11, 2, and 5 independent transformations, respectively. For Δ*pbp2a* strain, transposon library starter cultures were obtained from ≈300,000 colonies with 4 independent transformations. Transposon library starter cultures from different transformations were thawed and mixed together in proportion to the numbers of transformants obtained from each transformation. The combined starter cultures were diluted to OD_620_ ≈0.005 in 5 mL of BHI broth containing 180 μg/mL spectinomycin and 5μL/mL Ec-oxyrase (Oxyrase, EC0005), and were grown at 37°C with 5 % CO_2_ to OD_620_ ≈0.4. 5 mL of culture at OD_620_ ≈0.4 were used to extract genomic DNA using DNeasy blood and tissue kit. 3 µg of DNA from each sample was used for *MmeI* digestion, followed by ligation to adaptors described in (van Opijnen *et al*., 2015). The samples were further processed according to (Fenton *et al*., 2016) and sequenced on the Illumina NextSeq 500 using a NextSeq 75 high sequencing kit at the Center for Genomics and Bioinformatics, Indiana University Bloomington. Sequencing reads were de-multiplexed and trimmed using the QIAGen CLC genomics workbench (version 11.0.1). Data were mapped and analyzed as described in (Fenton *et al*., 2016). Insertion data were visualized graphically using the Artemis genome browser (version 10.2) (Carver *et al*., 2012). Tn-seq primary data for the region between *mreD* (*spd_2044*) and *spd_2051*, the gene upstream *rodZ* (*spd_2050*), are contained in Appendix A, including run summaries, number of reads per TA site in each gene, and count ratios for each gene in the indicated mutants compared to WT. P values for comparisons of the number of reads per TA site in each gene were calculated by the Mann-Whitney test using GraphPad Prism (9.2.0).

### 4.3 Growth of Zn-dependent depletion and merodiploid strains

Ectopic expression of *rodZ* or *mreC* was achieved from a zinc-inducible promoter (P_Zn_) in the *bgaA* site (Tsui *et al*., 2016, Tsui *et al*., 2014). 0.4 mM of ZnCl_2_ and corresponding 1/10 concentration of MnSO_4_ were added to TSAII-BA plates or BHI broth for inducing conditions. Mn^2+^ was added to Zn^2+^ conditions to prevent zinc toxicity (Jacobsen *et al*., 2011, Perez *et al*., 2021b, Tsui *et al*., 2016). Depletion strains requiring ZnCl_2_ for growth were grown overnight in BHI broth in the presence of Zn inducer (0.4 mM ZnCl_2_ + 0.04 mM MnSO_4_). For depletion/complementation experiments, overnight cultures (OD_620_ of 0.1-0.4) supplemented with inducer were diluted to an OD_620_ of 0.003 in 5 mL fresh BHI with or without inducer. Growth was monitored every 1 hour using a Genesys 2 spectrophotometer (Thermo Scientific) for 10 h. For these experiments, the point of resuspension serves as time zero, (T = 0). All growth curves and microscopy experiments were performed two or more times with similar results.

### 4.4 Transformation assays

Transformations were performed as detailed in (Rued *et al*., 2017, Tsui *et al*., 2016). All amplicons (experimental and control) contained ≈1 kb flanking region and were obtained from PCR reactions using primer pairs and templates listed in Table S4. Recipient strains were grown to OD_620_ ≈0.03 from glycerol ice stock and 100 µL was added to 900 µL of transformation mix containing 10% (wt/vol) heat-treated horse serum, 0.18% (wt/vol) glucose, 100 ng CSP-1 (competence stimulatory peptide, type 1) mL^-1^ and 9 mL of BHI. The mixture was incubated for 10 min at 37°C in the presence of 5% CO_2_. 30 or 100 ng of purified amplicon (for unencapsulated and encapsulated strains, respectively) was added to the transformation mixture and incubated for 1 h at 37°C in the presence of 5% CO_2_. A fraction or the entire final transformation mixture was added to 3 mL of soft agar containing the appropriate antibiotic (72 µL of 0.1 mg erythromycin mL^-1^, 36 µL of 0.1 mg spectinomycin mL^-1^, 60 µL of 100 mg streptomycin ml^-1^, or 120 µL of 50 mg kanamycin mL^-1^) and plated onto TSAII-BA plates. Unless explicitly stated, the numbers of colonies listed are normalized to 1 mL of transformation mixture. Transformants were incubated overnight in the presence of 5% CO_2_ for 20-24 h, at which time colony numbers and morphology were counted and evaluated.

### 4.5 Viable count (CFU) assays

At various time points (3, 4, 5, 6, and 7.5 h) in depletion and complementation experiments, 100 µL of culture was suspended in 900 µL of 1X PBS and serially diluted from 10^-1^ to 10^-7^. 100 µL of three selected dilutions were suspended in 3 mL of molten soft agar and poured onto TSAII-BA plates with or without inducer, 0.4 mM Zn + 0.04 mM Mn. Solidified plates were incubated at 37°C in the presence 5% CO_2_ for 20-24 h. Plates containing 30-300 CFUs were scored and counted with respect to colony number and size. CFU/mL values were calculated for each strain and condition.

### 4.6 Image Acquisition and processing

For 2D-epifluorescence microscopy (eFM), 2D-immunofluorescence microscopy (IFM), and phase-contrast microscopy (PCM) experiments, images were taken using a Nikon E-400 epifluorescence phase-contrast microscope and 100X Nikon Plan Apo oil-immersion objective (numerical aperture, 1.40) connected to a CoolSNAP HQ2 charged-coupled device (CCD) camera (Photometrics). Images were analyzed with NIS-Elements AR software (Nikon). Micrographs were assembled using Adobe Illustrator and all images are to scale.

### 4.7 Cell length and width measurements

Cell lengths and widths of strain growing exponentially in BHI broth were measured as previously described (Perez *et al*., 2021b, Tsui *et al*., 2016). For all strains, only ovoid-shape pre-divisional cells were measured. Unless indicated in the figure legends, more than 50 cells from one experiment were measured, and plotted with box and whiskers plot (5 to 95 percentile whiskers). P values were obtained by one-way ANOVA analysis by using the Kruskal-Wallis test in GraphPad Prism program.

### 4.8 Live-dead staining and eFM

Live/dead staining was performed using the BacLight Bacterial Viability kit (Syto9 and Propidium Iodide), according to the manufacturer’s instructions (ThermoFisher Scientific, Cat. #L7007) and (Perez *et al*., 2021b, Sham *et al*., 2013). Briefly, 500 µL of culture were harvested at 4 and 6 h of growth and centrifuged at 12,000 x *g* for 2.5 min at 25°C. Pellets were re-suspended in 50 μL of BHI broth plus 2 μL of a 1:1 (v/v) mixture of Syto-9 and propidium iodide, and incubated for 5 min in the dark at 22°C. As a control, heat killed cells (95° C for 5 min) were stained as described above for comparison. After incubation, samples were immediately imaged using both the Alexa 488 (EX 460-500, DM 505, and BA 510-560) and Alexa 568 (EX 532-587, DM 595, and BA 608-682) filters. The staining pattern of the sample was revealed by superimposing the corresponding Alexa 488 and Alexa 568 images upon one another within the NIS-Elements AR software (Nikon). A total of 200 cells were categorized based on the staining pattern for quantification of percentage of live vs dead cells.

### 4.9 FDAA (fluorescent D-amino acids) short-pulse labeling and eFM

The FDAA TADA (tetramethylrhodamine 3-amino-D-alanine) synthesized as described in (Kuru *et al*., 2012) was obtained from Michael VanNieuwenhze. Samples were processed as described in (Boersma *et al*., 2015, Perez *et al*., 2021a, Tsui *et al*., 2014) with minor changes. Briefly, at 4 h of growth, 500 µL of culture was harvested and centrifuged at 16,000 x *g* for 5 min at room temperature. Cultures were washed twice with 500 µL ice cold 1X PBS. Pelleted via centrifugation (16,000*×* for 5 min) and re-suspended in 250 µL of BHI broth containing TADA at a final concentration of 500 µM. Working solutions of TADA in BHI were diluted from 500 mM stocks in DMSO, which were stored at −20°C. Samples were incubated for 5 min at 37°C. Cells were centrifuged for 2.5 min at 16 000 x *g* at 4°C, and washed twice in 1 mL of cold 1x PBS. Pellets were resuspended in 1 mL of 4% (wt/vol) paraformaldehyde (EMS; 157-4), followed by 15 min incubation at room temperature and 45 min incubation on ice in the dark. Fixed cells were centrifuged and washed three times with ice cold 1x PBS. After which cells were resuspended in 50-75 µL of Slowfade Gold antifade reagent (Invitrogen; S36936), vortexed briefly, and applied to a glass slide. A glass coverslip was gently placed onto to the slide and the samples were cured overnight at 4° C, in the dark. To visualize TADA labeling, images were taken using eFM with a Texas-Red filter.

### 4.10 HaloTag labeling and eFM

To determine the localization pattern of the HaloTag-fusion proteins, cells were labeled with saturating TMR ligand and viewed by eFM as described in (Perez *et al*., 2019, Perez *et al*., 2021b). Briefly, strains with HT-domain fusions expressed from native chromosomal loci were grown in BHI broth as described above. At 4 h of growth, 0.5 µL of working stock of TMR (500 µM TMR ligand (Promega cat #G8252) in DMSO stored at −20° C) was added to 300 µL of culture (final concentration = 0.83 µM). Tubes were inverted gently three times and then incubated for 15 min at 37 °C in the dark in the absence of CO_2_. Cells were then collected by centrifugation at 14,000 x *g* for 2.5 min, washed once with 500 µL of fresh BHI broth, re-pelleted (14,000 x *g* for 2.5 min), and resuspended in 15-20 µL of fresh BHI broth. Cell shape and fluorescence localization patterns were imaged using PCM and eFM with a Texas-red filter.

### 4.11 Demograph generation

Demographs showing protein fluorescence intensity as a function of cell length were generated using Microbe J (version 5.11s) (Ducret *et al*., 2016, Perez *et al*., 2019, Perez *et al*., 2021b). Demographs were also generated from phase-contrast images corresponding to light scattering caused by the cell body as a function of cell length. For cells displaying WT size and morphology, the following “WT” parameters were used: (area [µm^2^] 0.53-max; length [µm] 0.5-3.2; width ([µm] 0.2-max; circularity [0-1] 0-max; curvature [0-max] 0-max; sinuosity [0-max] 0-max; angularity [rad] 0-0.38; solidity [0-max] 0.75-max; intensity [0-max] 0-6200; Z-score 2.0-max). Within a given WT field of cells, ≈3-5% of the population were excluded from automated selection by the program due to variables such as clustering of cells, size, shape defects, and/or out of plane of focus. These cells were not manually entered into the program. Stages of cells were classified by the degree of separation as described in (Perez *et al*., 2019). For depletion conditions in which gross morphological changes occurred in both cell shape and size, the “WT” parameters did not fit to automated selection, leading to the exclusion of > 30 % of the cells from automated selection. Therefore, a separate set of “mutant” parameters were used: (area [µm^2^] 0.43-5; length [µm] 0.3-5; width ([µm] 0.4-max; circularity [0-1] 0-max; curvature [0-max] 0-0.45; sinuosity [0-max] 0-max; angularity [rad] 0-0.45; solidity [0-max] 0.75-max; intensity [0-max] 0-6200; Z-score 1.0-max). Complementation fields were analyzed using the “mutant” parameters.

### 4.12 IFM of strains expressing single epitope-tagged proteins

IFM of cells harvested after 4 h of growth was performed as described in (Land *et al*., 2013, Tsui *et al*., 2016, Tsui *et al*., 2014). Primary antibodies were: rabbit anti-FLAG polyclonal antibody (Sigma, F7425, 1:100 dilution) or rabbit anti-HA polyclonal antibody (Invitrogen, 71–5500, 1:100 dilution). Secondary antibodies used were: anti-rabbit IgG conjugated to Alexa Fluor 488 (Life Technologies; A11034, 1:100 dilution). Control experiments did not detect labeling in untagged WT IU1824 or untagged depletion IU12738 (Δ*rodZ*//P_Zn_-*rodZ*^+^) and IU12345 (Δ*mreC//*P_Zn_-*mreC*^+^) strains. Protein localization patterns were visualized and scored across multiple fields for depletion/complementation conditions and WT backgrounds. Cells were scored in accordance with the key provided in the appropriate figure legends.

### 4.13 IFM of strains expressing two epitope-tagged proteins

For co-localization IFM studies, double epitope-tagged strains IU7072 (*rodZ-*L-F^3^ *ftsZ-*Myc), IU7113 (*mreC-*L-F^3^ *rodZ-*Myc), and IU7515 (*pbp1a-*L-F^3^ *rodZ-*Myc) were grown exponentially to OD_620_ ≈0.15-0.20 and processed as detailed in (Land *et al*., 2013, Tsui *et al*., 2016, Tsui *et al*., 2014). Primary antibodies used were: anti-FLAG rabbit polyclonal antibody (dilution 1:100) and anti-Myc mouse monoclonal antibody (dilution 1:100). Secondary antibodies used were: goat anti-rabbit conjugated to Alexa Fluor 488 (1:100) or goat anti-mouse-Alexa Fluor 568 (1:100). Primary and secondary antibody incubations were for 2 h at 37°C, and 1 h at 24°C, respectively. DNA in nucleoids was stained with SlowFade Gold Antifade reagent with DAPI (Life Technologies, S36939). Image analysis was performed using a point-and-click IMA-GUI organized in MATLAB (The Mathworks) as described in (Land *et al*., 2013, Tsui *et al*., 2014). Pneumococcal cells were manually aligned and binned into four division stages 1 to 4: pre-, early-, mid- and late-divisional, within the program by eye. The mean cell outline (phase-contrast image) and fluorescence intensities of DNA or tagged-proteins were measured and represented graphically as previously performed in (Land *et al*., 2013). (*n*) indicates the numbers of cells averaged for that particular stage of cellular division. Analysis was conducted using data from two independent biological replicates.

### 4.14 Structured-illumination microscopy (3D-SIM)

IFM images were taken using the Deltavision OMX Super Resolution system located in the Indiana University Bloomington Light Microscopy Imaging Center (LMIC) as detailed in (Tsui *et al*., 2014). Briefly, the system is equipped with four Photometrics Cascade II EMCCD cameras that allow simultaneous four-color imaging, and is controlled by DV-OMX software, with image processing by Applied Precision softWoRx 6.0. software. For information: http://www.indiana.edu/~lmic/microscopes/index.htmL#OMX

### 4.15 Western blotting and immunodetection

Cell lysates were prepared by the SEDS lysis-buffer (0.1% deoxycholate (vol/vol), 150 mM NaCl, 0.2% SDS (vol/vol), 15 mM EDTA pH 8.0)) method as described in (Cleverley *et al*., 2019). Briefly, bacteria were grown exponentially in 5 mL BHI broth to an OD_620_ ≈0.1-0.2. Aliquots of 1.0-2.0 mL were centrifuged (5 min, 16,000 × *g* at 4°C), and cell pellets were washed once with 4°C PBS. Frozen pellets collected from 1.8 mL of cultures at OD_620_ = 0.16 were suspended in 80 µL of SEDS lysis buffer. Samples collected from different volumes or at different OD_620_ readings were resuspended in volumes of SEDS lysis buffer proportional to the culture volumes and cell densities. Cell lysis was performed by incubation at 37°C in a shaking block at 300 rpm for 15 min. Protein concentrations of lysed samples were determined with Bio-Rad DC^TM^ protein assay kit. Samples were denatured with 2x Laemmeli SDS loading buffer (Bio-Rad) plus β-mercaptoethanol (5% vol: vol) at 95°C for 10 min. 3-10 µg of total crude lysate per sample was loaded onto a 4-15% precast gradient SDS-PAGE gel (Bio-Rad) and subjected to electrophoresis. Amounts of crude lysates loaded and primary antibodies are specified for individual experiments.

Sources of antibodies used for western blotting are as below. Primary antibodies used are anti-HaloTag monoclonal antibody (Promega, G921A, 1:1000), and the following polyclonal rabbit antibodies: anti-FLAG (Sigma, F7425, 1:2,000); anti-HA (Invitrogen, 71– 5500, 1:1,000); anti-Myc (ThermoFisher Scientific, PA1-981, 1:1,000); anti-GFP (Invitrogen, A11122, 1:1,400); anti-StkP ((Beilharz *et al*., 2012), 1:10,000); anti-PhpP ((Beilharz *et al*., 2012), 1:5,000); anti-MreC ((Land & Winkler, 2011), 1:5,000); anti-FtsZ ((Lara *et al*., 2005), 1:20,000); anti-FtsA ((Lara *et al*., 2005),1:20,000); anti-bPBP2b ((Perez *et al*., 2021a), 1:10,000); anti-bPBP2x ((Perez *et al*., 2021a), 1:10,000); anti-GpsB ((Cleverley *et al*., 2019), 1:2,000); anti-aPBP2a ((Cleverley *et al*., 2019), 1:5,000); and anti-DivIVA ((Fadda *et al*., 2007), 1:5,000). Anti-aPBP1a (1:5,000) was generated with purified aPBP1a (aa S37 to P719) and showed no signal at 94 kDa in lysate prepared from a Δ*pbp1a* strain. Secondary antibodies used were anti-mouse IgG conjugated to horseradish peroxidase (Invitrogen, SA1-100, 1:3300), anti-rabbit IgG conjugated to horseradish peroxidase (GE healthcare NA93AV, 1:10,000), or Licor IR Dye800 CW goat anti-rabbit (926-32211, 1:14,000). Chemiluminescence signals obtained with secondary HRP-conjugated antibodies were detected using IVIS imaging system as described previously (Wayne *et al.,* 2010). IR signals obtained with Licor IR Dye800 CW secondary antibody was detected with Azure biosystem 600.

### 4.16 Depletion experiments with quantitative western blotting

Depletion of proteins expressed from a Zn^2+^-inducible prompter (P_Zn_) was performed as described in (Perez *et al*., 2021b, Tsui *et al*., 2016) with the following modifications. Merodiploid strains IU12345 (Δ*mreC*//P_Zn_-*mreC*^+^) and IU10947 (Δ*rodZ*//P_Zn_-*rodZ-*F) require 0.4 mM Zn + 0.04 mM Mn in BHI broth for growth and ectopic induction of MreC or RodZ-F, respectively. To measure protein amounts during depletion of MreC, or RodZ-F, IU1824 (WT), IU12345 (Δ*mreC*//P_Zn_-*mreC*^+^), IU14594 (*rodZ*-F) and IU10947 (Δ*rodZ*//P_Zn_-*rodZ-*F) were diluted from overnight cultures (IU12345 and IU10947 supplemented with Zn inducer (0.4 mM ZnCl_2_ +0.04 mM MnSO_4_) and re-suspended to an OD_620_ ≈0.003 in fresh BHI broth ± Zn inducer. Cultures were harvested at 3 or 4 h of growth, and processed for western blotting as described above. To ensure western blots were quantitative, standard curves were generated by loading a range of protein amounts on the lanes, and labeled for anti-MreC, or anti-FLAG for RodZ-F (Perez *et al*., 2021a). A protein amount corresponding to the mid-range of the standard curve was loaded for each targeted protein. 3 µg of cell lysate were loaded for the detection of bPBP2x or bPBP2b, 3 or 6 µg for detection of MreC, and 10 µg for the detection of RodZ-F. For the detection of RodZ-F during depletion, Licor IR Dye800 CW goat anti-rabbit secondary antibody (926-32211)(1:14,000) was used, and IR signal was detected with Azure biosystem 600. In addition, signals obtained with anti-F antibody were normalized with total protein stain in each lane using Totalstain Q-NC reagent from Azure.

### 4.17 Phos-tag SDS-PAGE and western blotting

This method is based on the mobility shift of phosphorylated proteins in SDS-PAGE with polyacrylamide-bound Mn^2+^-Phos-tag (Kinoshita *et al*., 2006). Phosphorylated proteins in gels are visualized as slower migrating bands compared to corresponding unphosphorylated proteins. Phos-tag SDS-PAGE and standard Western blotting were carried out as described previously with modifications (Wayne *et al*., 2012). Overnight BHI broth cultures were diluted and grown up to OD_620_ ≈0.2 in 30 mL of BHI. Cells were centrifuged at 14,500 x *g* for 5 min at 4°C, and all subsequent steps were performed at 4°C. Pellets were lysed using a FastPrep homogenizer (MP biomedicals) in cold lysis buffer (20 mM Tris-HCl pH 7.0 and 1 protease inhibitor tablet (ThermoFisher Scientific) per 10 mL buffer). Cell lysates were resolved by 10 % SDS-PAGE supplemented with 75 μM Phos-tag acrylamide (AAL-107; Wako) and 100 μM MnCl_2_, and standard 10 % SDS-PAGE as control. Volumes of loaded samples were normalized to OD_620_ of harvested cultures (for OD_620_ ≈0.2, 30 µL of the sample was loaded). Gel electrophoresis was carried out for 3h. RodZ-HA^3^ was detected by western blotting as described above using anti-HA as the primary antibody.

### 4.18 Co-immunoprecipitation (Co-IP) assays

Co-IP assays were performed as previously described in (Perez *et al*., 2019, Perez *et al*., 2021b, Rued *et al*., 2017). Briefly, cultures were grown exponentially to an OD_620_ ≈0.2-0.3 in 400 mL of BHI broth, concentrated 20-fold in 4°C PBS, and cross-linked with 0.1 % paraformaldehyde for 1h at 37°C. After quenching with 1 M glycine and a wash with PBS, pellets were re-suspended in 2 mL of cold lysis buffer (50 mM Tris-HCl pH 7.4, 150 mM NaCl, 1 mM EDTA, 1% Triton X100 (v/v) containing protease inhibitor, and homogenized in lysing matrix B tubes (MP Biomedicals) in a FastPrep homogenizer. One mL of lysate (input sample, normalized to ≈2-5 mg/mL) was added to 50 μL of anti-FLAG magnetic beads (Sigma, M8823) and incubate for 2h at 4°C with rotation. 100 μL of FLAG elution solution containing 150 ng 3X FLAG peptide/μL (Sigma, F4799) was used to elute FLAG-tagged proteins and other associated proteins (output elution samples). Input or output samples were mixed with 2X Laemmli sample buffer (Bio-Rad) containing 10% (vol/vol) β-mercaptoethanol (Sigma), and were heated at 95°C for 1 h to break cross-links, with the exception of IU8918 (*ftsW*-L-*gfp*) and IU16126 (*ftsW*-L-*gfp rodZ-*L*-*F^3^), when the samples were not heated. For most samples, 4-6 µL of input mixed with sample buffer was loaded onto pre-cast SDS-PAGE 4-15% gels (Bio-Rad), resulting in ≈4-9 µg of cell lysate. For the output (elutions), 15 or 25 µL of samples mixed with sample buffer were loaded. Proteins were detected using standard western blotting and immunodetection methods described above. All pair-wise co-IP experiments were performed independently 2-6 times.

### 4.19 Bacterial two-hybrid (B2H) assays

B2H assays were performed as described before (Cleverley *et al*., 2019, Perez *et al*., 2021b, Rued *et al*., 2017) with the following modifications. The hybrid plasmids used in the B2H assays are listed in Table S3. For cloning, the target genes were amplified by PCR from *S. pneumoniae* D39 chromosomal DNA (or its derivatives) using the primers listed in Table S4. PCR fragments for *rodZ*(ΔHTH)*, rodZ*(ΔDUF) and *pbp1b* were purified, digested with the appropriate restriction enzymes and cloned into the corresponding sites of the pKT25/pUT18C vectors to generate plasmids encoding the corresponding hybrid proteins fused at the C-terminal ends of the T25/T18 fragments. *E. coli* DH5α or XL1-blue transformants were selected on LB agar plates containing ampicillin (100 μg/mL) or kanamycin (50 μg/mL) and 0.4% glucose to repress leaky expression (Karimova *et al*., 2005). The correct sequence of each construct was verified by double-strand sequencing, using primers listed in Table S4. B2H vectors pKT25/pUT18C containing *rodZ*, *mreC, mpgA, pbp1a, pbp2b, rodA, pbp2a, pbp2x, ftsW,* and *ftsA* and vectors pKNT25/pUT18 containing *ftsZ*, *ezrA, gpsB*, *divIVA*, *stkP* and *mreD* were previously constructed and reported (Cleverley *et al*., 2019, Krupka *et al*., 2012, Perez *et al*., 2021b, Rued *et al*., 2017). Each pair of plasmids was co-transformed into the *E. coli cya*^-^ BTH101 strain and co-transformation mixtures were spotted directly onto LB agar plates supplemented with ampicillin (100 μg/mL), kanamycin (50 μg/mL) and X-Gal (60 μg/mL), followed by incubation at 30°C. Plasmid pairs pKT25/pUT18C and pKT25-zip/pUT18C-zip were used as negative and positive controls, respectively. Plates were inspected and photographed after 24 h and 40 h. In the case of time course experiments, B2H plates were inspected for color development after 24, 30 and 36 h of incubation at 30°C and scored similarly as reported in (Bendezu *et al*., 2009). All the B2H experiments were performed at least twice.

## Supporting information

Tn-seq run summaries, reads, and ratios

## ACKNOWLEDGEMENTS

We thank Kevin Bruce, Jiaqi Zheng, and other members of the Winkler lab for discussions about this work; Jim Powers and Sidney Shaw (Indiana University Bloomington) for advice about light microscopy; Jason Rosch (St. Jude’s) and Tim van Opijnen (Boston College) for Tn-seq protocols; Mike VanNieuwenhze (Indiana University Bloomington) for TADA FDAA reagent; and Suzanne Walker and David Rudner (Harvard Medical School) for antibodies against pneumococcal PG synthesis proteins. This work was supported by NIH Grants R35GM131767 (to MEW), R01GM141242 (to XW), Predoctoral Grant F31AI138430 (to MML), and NIH Equipment Grant S10OD024988 (to the Indiana University Bloomington (IUB) Light Microscopy Imaging Center); and by institutional research funds from the CIBIO Department of the University of Trento (to OM).

## AUTHOR CONTRIBUTIONS

MML, HCTT, and MEW contributed to the conception or design of this study. MML, IM, MJ, ZAY, MB, AZ, ZR, XW. OM, HCTT, and MEW contributed to the acquisition, analysis, and interpretation of the data. MML, OM, HCTT, and MEW contributed to the writing of the manuscript with input from the other authors.

## DATA AVAILABILITY

The data that support the findings of this study are available in Appendix A and from the corresponding authors upon reasonable request.

## ABBREVIATED SUMMARY

This paper establishes RodZ as an essential scaffolding protein required for the assembly and function of the elongasome that synthesizes peripheral peptidoglycan (pPG) in *Streptococcus pneumoniae*, which lacks an MreB homolog. (Top panel) The assembly hierarchy mediated by RodZ(*Spn*). This paper also reports synthetic-viable, suppressor relationships between Class A aPBP1b and aPBP1a and components of the core pPG elongasome. (Bottom panel) Bypass model for the modulation of function of the pPG elongasome to restore viability.

