## Supplemental Tables and Figures for "Roles of RodZ and Class A PBP1b in the Assembly and Regulation of the Peripheral Peptidoglycan Elongasome in Ovoid-Shaped Cells of *Streptococcus pneumoniae* D39"

**Contents:**

**Supplemental Tables S1-S6**
**Supplemental References**
**Supplemental Figures S1-S27 with Legends**

**Table S1.** *Streptococcus pneumoniae* strains used in this study

| ***Streptococcus pneumoniae* strains** | | | |
|---|---|---|---|
| Strain number | Genotype (description)[a] | Antibiotic resistance[b] | Reference or source |
| EL59 | Unencapsulated laboratory strain R6 | None | (Hoskins *et al.*, 2001) |
| E46 | D39 Δ*cps* Δ*bgaA*::P$_c$-*erm* | Erm$^R$ | (Rued *et al.*, 2017) |
| E149 | D39 Δ*cps* Δ*mreC*::P$_c$-*erm* (IU1945 X fusion Δ*mreC*::P$_c$-*erm* amplicon) | Erm$^R$ | This study |
| E177 | D39 Δ*cps* Δ*pbp1a*::P$_c$-*erm* | Erm$^R$ | (Land *et al.*, 2013) |
| E193 | D39 Δ*cps* Δ*pbp1b*::P$_c$-*erm* | Erm$^R$ | (Land *et al.*, 2013) |
| E655 | D39 Δ*cps* Δ*rodZ*::P$_c$-*erm* | Erm$^R$ | Tsui *et al.*, 2016 |
| K49 | D39 Δ*cps* Δ*mreC*::P$_c$-[*kan-rpsL*$^+$] (IU1945 X fusion Δ*mreC*::P$_c$-[*kan-rpsL*$^+$] amplicon) | Kan$^R$ | This study |
| K164 | D39 Δ*cps* Δ*pbp1a*::P$_c$-[*kan-rpsL*$^+$] | Kan$^R$ | (Tsui *et al.*, 2016) |
| K166 | D39 Δ*cps* Δ*pbp2a*::P$_c$-[*kan-rpsL*$^+$] | Kan$^R$ | (Tsui *et al.*, 2016) |
| K180 | D39 Δ*cps* Δ*pbp1b*::P$_c$-[*kan-rpsL*$^+$] | Kan$^R$ | (Tsui *et al.*, 2014) |
| K654 | D39 Δ*cps* Δ*rodZ*::P$_c$-[*kan-rpsL*$^+$] (IU1945 X fusion Δ*rodZ::*P$_c$-[*kan-rpsL*$^+$]) | Kan$^R$ | (Tsui *et al.*, 2016) |
| IU1690 | D39 *cps*$^+$ | None | (Lanie *et al.*, 2007) |
| IU1751 | R6 Δ*mreCD<>aad9* | Spc$^R$ | (Land & Winkler, 2011) |
| IU1781 | D39 *cps*$^+$ *rpsL1* | Str$^R$ | (Ramos-Montañez *et al.*, 2008) |
| IU1824[c] | D39 *rpsL1* Δ*cps*2A'-*cps*2H' = D39 *rpsL1* Δ*cps* | Str$^R$ | (Lanie *et al.*, 2007) |
| IU1945 | D39 Δ*cps 2A'-cps2H'* = D39 Δ*cps* | None | (Lanie *et al.*, 2007) |
| IU4970 | D39 Δ*cps* *mreC*-L$_0$-FLAG$^3$-P$_c$-*erm* | Erm$^R$ | (Land & Winkler, 2011) |
| IU5544 | D39 Δ*cps* *pbp1a*-L$_0$-FLAG$^3$-P$_c$-*erm* | Erm$^R$ | (Land & Winkler, 2011) |

| | | | |
|---|---|---|---|
| IU5648 | D39 Δ*cps rpsL1 divIVA*-P$_c$-[*kan-rpsL*$^+$] | Kan$^R$ | (Perez *et al.*, 2019) |
| IU5840 | D39 Δ*cps pbp1a*-FLAG-P$_c$-*erm* | Erm$^R$ | (Land *et al.*, 2013) |
| IU6291 | D39 Δ*cps rodZ*-L$_0$-FLAG$^3$-P$_c$-*erm*<br>(IU1945 X fusion *rodZ*-L$_0$-FLAG$^3$-P$_c$-*erm*) | Erm$^R$ | This study |
| IU6293 | D39 Δ*cps rodZ*-FLAG-P$_c$-*erm*<br>(IU1945 X fusion *rodZ*-FLAG-P$_c$-*erm*) | Erm$^R$ | This study |
| IU6397 | D39 *rpsL1* Δ*bgaA*::*kan*-t1t2-P$_{ftsA}$-*phoU2* | Kan$^R$ | (Zheng *et al.*, 2016) |
| IU6741 | D39 Δ*cps rpsL1* Δ*pbp1a* markerless | Str$^R$ | (Tsui *et al.*, 2016) |
| IU6933 | D39 Δ*cps pbp2b*-HA-P$_c$-*kan* | Kan$^R$ | (Tsui *et al.*, 2014) |
| IU6962 | D39 Δcps *ftsZ*-Myc-P$_c$-*kan* | Kan$^R$ | (Land *et al.*, 2013) |
| IU6987 | D39 Δ*cps* Δ*rodZ*::P$_c$-*aad9* (IU1945 X fusion Δ*rodZ*::P$_c$-*aad9* amplicon) | Spc$^R$ | This study |
| IU7054 | D39 Δ*cps* Δ*bgaA*::*kan*-t1t2-P$_{ftsA}$-*ftsZ* (IU1945 X fusion amplicon) | Kan$^R$ | This study |
| IU7068 | D39 Δ*cps rodZ*-Myc-P$_c$-*kan*<br>(IU1945 X fusion *rodZ*-Myc-P$_c$-*kan)* | Kan$^R$ | This study |
| IU7072 | D39 Δ*cps rodZ*-L$_0$-FLAG$^3$-P$_c$-*erm ftsZ*-Myc-P$_c$-*kan*<br>(IU6962 X *rodZ*-L$_0$-FLAG$^3$-P$_c$-*kan* from IU6291) | Erm$^R$ Kan$^R$ | This study |
| IU7113 | D39 Δ*cps rodZ*-Myc-P$_c$-*kan mreC*-L$_0$-FLAG$^3$-P$_c$-*erm*<br>(IU7068 X *mreC*-L$_0$-FLAG$^3$-P$_c$-*erm* from IU4970) | Erm$^R$ Kan$^R$ | This study |
| IU7242 | D39 Δ*cps pbp1a*-HA-P$_c$-*kan* | Kan$^R$ | (Rued *et al.*, 2017) |
| IU7397 | D39 Δ*cps* Δ*pbp2b*<>*aad9* //Δ*bga*::*kan*-t1t2-P$_{fcsk}$-*pbp2b*$^+$ | Spc$^R$ Kan$^R$ | (Tsui *et al.*, 2014) |
| IU7399 | D39 Δ*cps mpgA*-HA-P$_c$-*kan* | Kan$^R$ | (Tsui *et al.*, 2016) |
| IU7403 | D39 Δ*cps mpgA*-FLAG -P$_c$-*erm* | Erm$^R$ | (Tsui *et al.*, 2016) |
| IU7426 | D39 Δ*cps pbp2b*-HA$^4$-P$_c$-*kan* | Kan$^R$ | (Tsui *et al.*, 2014) |
| IU7434 | D39 Δ*cps stkP*-FLAG$^2$-P$_c$-*erm* | Erm$^R$ | (Tsui *et al.*, 2014) |
| IU7515 | D39 Δ*cps rodZ*-Myc-P$_c$-*kan pbp1a*-L$_0$-FLAG$^3$-P$_c$-*erm*<br>(IU7068X *pbp1a*-L$_0$-FLAG$^3$-P$_c$-*erm* from IU5544) | Erm$^R$ Kan$^R$ | This study |
| IU7584 | D39 Δ*cps rodZ*-L$_0$-FLAG$^3$-P$_c$-*erm mpgA*-HA-P$_c$-*kan*<br>(IU6291 X *mpgA*-HA-P$_c$-*kan* amplicon from IU7399) | Erm$^R$ Kan$^R$ | This study |
| IU7614 | D39 Δ*cps rpsL1 ftsZ*$^+$-P$_c$-[*kan-rpsL*$^+$] | Kan$^R$ | (Tsui *et al.*, 2016) |
| IU7616 | D39 Δ*cps rpsL1 fts*A$^+$-P$_c$-[*kan-rpsL*$^+$] | Kan$^R$ | (Perez *et al.*, 2019) |

| | | | |
|---|---|---|---|
| IU7814 | D39 Δcps Δ*ftsZ*::*aad9*//Δ*bgaA*::*kan*-t1t2-P$_{ftsA}$-*ftsZ*$^{+}$ | Spc$^{R}$ Kan$^{R}$ | (Perez *et al.*, 2019) |
| IU7850 | D39 Δ*cps rpsL1* Δ*pbp1b*::P$_{c}$-[*kan-rpsL*$^{+}$]( IU1824 X Δ*pbp1b*::P$_{c}$-[*kan-rpsL*$^{+}$] amplicon from K180) | Kan$^{R}$ | This study |
| IU8122 | D39 Δcps Δ*bgaA*::*tet*-P$_{Zn}$-RBS$^{ftsA}$-*ftsZ*$^{+}$ | Tet$^{R}$ | (Zheng *et al.*, 2017) |
| IU8918 | D39 Δ*cps rpsL1 ftsW*-L$_{2}$-*gfp* markerless | Str$^{R}$ | (Perez *et al.*, 2019) |
| IU8921 | D39 Δ*cps rpsL1* P$_{c}$-[*kan-rpsL*$^{+}$]-*pbp2x* | Kan$^{R}$ | (Perez *et al.*, 2019) |
| IU8980 | D39 Δ*cps rpsL1* P$_{c}$-[*kan-rpsL*$^{+}$]-*mpgA* | Kan$^{R}$ | (Tsui *et al.*, 2016) |
| IU8986 | D39 Δ*cps rpsL1 mpgA*$^{+}$-[*kan-rpsL*$^{+}$] | Kan$^{R}$ | (Tsui *et al.*, 2016) |
| IU9023 | D39 Δ*cps rpsL1* P$_{c}$-[*kan-rpsL*$^{+}$]-*pbp2b*$^{+}$ | Kan$^{R}$ | (Perez *et al.*, 2019) |
| IU9036 | D39 Δ*cps rpsL1* Δ*khpA* markerless | Str$^{R}$ | (Zheng *et al.*, 2017) |
| IU9077 | D39 Δ*cps rpsL1 ezrA*-P$_{c}$-[*kan-rpsL*$^{+}$] | Kan$^{R}$ | (Perez *et al.*, 2019) |
| IU9094 | D39 Δ*cps rpsL1* P$_{c}$-[*kan-rpsL*$^{+}$]-*mapZ* | Kan$^{R}$ | (Perez *et al.*, 2019) |
| IU9102 | D39 Δ*cps* Δ*mpgA*::P$_{c}$-*aad9* //Δ*bgaA*::tet-P$_{Zn}$-RBS$^{mpgA}$-*mpgA* | Spc$^{R}$ Tet$^{R}$ | (Tsui *et al.*, 2016) |
| IU9167 | D39 Δ*cps rpsL1 divIVA*-L$_{2}$-*gfp* markerless | Str$^{R}$ | (Perez *et al.*, 2019) |
| IU9182 | D39 Δ*cps rpsL1 gfp*-L$_{1}$-*mapZ* markerless | Str$^{R}$ | (Perez *et al.*, 2019) |
| IU9602 | D39 Δ*cps khpA*-L$_{0}$-FLAG$^{3}$-P$_{c}$-*erm* | Erm$^{R}$ | (Zheng *et al.*, 2017) |
| IU9613 | D39 Δ*cps rpsL1* Δ*bgaA*::*tet*-P$_{Zn}$-RBS$^{ftsA}$-*rodZ*$^{+}$ (IU1824 X fusion Δ*bgaA*::*tet*-P$_{Zn}$-RBS$^{ftsA}$-*rodZ*$^{+}$) | Str$^{R}$ Tet$^{R}$ | This study |
| IU9621 | D39 Δcps rpsL1 Δ*khpA*//Δ*bgaA*::*kan*-t1t2-P$_{ftsA}$-*khpA*$^{+}$ | Str$^{R}$ Kan$^{R}$ | (Zheng *et al.*, 2017) |
| IU9760 | D39 Δ*cps rpsL1 mpgA*(Y488D) markerless (IU8986 X *mpgA*(Y488D) amplicon) | Str$^{R}$ | (Tsui *et al.*, 2016) |
| IU9765 | D39 Δ*cps* Δ*bgaA*::*tet*-P$_{Zn}$-RBS$^{ftsA}$-*rodZ*$^{+}$ | Tet$^{R}$ | (Tsui *et al.*, 2016) |
| IU9767 | D39 Δ*cps rpsL1* P$_{c}$-[*kan-rpsL*$^{+}$]-*ftsA* | Kan$^{R}$ | (Mura *et al.*, 2017) |
| IU9895 | D39 Δ*cps mpgA*(Y488D)-P$_{c}$-*erm* | Erm$^{R}$ | (Tsui *et al.*, 2016) |
| IU9931 | D39 Δ*cps* Δ*rodZ*<>*aad9*//Δ*bgaA*::*tet*-P$_{Zn}$-RBS$^{ftsA}$-*rodZ*$^{+}$ | Spc$^{R}$ Tet$^{R}$ | (Tsui *et al.*, 2016) |
| IU9969 | D39 Δ*cps rpsL1* Flag-*ftsA* markerless | Str$^{R}$ | (Mura *et al.*, 2017) |

| | | | |
|---|---|---|---|
| IU9985 | D39 Δ*cps rpsL1 ftsZ*-$L_2$-*sfgfp* markerless | $Str^R$ | (Perez *et al.*, 2019) |
| IU9990 | D39 Δ*cps* Δ*bgaA*::*tet*-$P_{Zn}$-$RBS^{ftsA}$-*pbp2b*$^+$ (IU1945 X fusion amplicon) | $Tet^R$ | This study |
| IU10035 | D39 Δ*cps rpsL1 gfp*-$L_1$-*ftsA* markerless | $Str^R$ | (Perez *et al.*, 2019) |
| IU10103 | D39 Δ*cps rpsL1* $P_c$-[*kan-rpsL*$^+$]-*mreC*$^+$ (IU1824 X fusion $P_c$-[*kan-rpsL*$^+$]-*mreC*$^+$ amplicon) | $Kan^R$ | This study |
| IU10220 | D39 Δ*cps rpsL1* Δ*bgaA*::*tet*-$P_{Zn}$-$RBS^{ftsA}$-*mreC*$^+$ (IU1824 X fusion Δ*bgaA*::*tet*-$P_{Zn}$-$RBS^{ftsA}$-*mreC*$^+$ ) | $Tet^R$ $Str^R$ | This study |
| IU10222 | D39 Δ*cps* Δ*bgaA*::*tet*-$P_{Zn}$-$RBS^{ftsA}$-*mreC*$^+$ (IU1945 X fusion Δ*bgaA*::*tet*-$P_{Zn}$-$RBS^{ftsA}$-*mreC*$^+$) | $Tet^R$ | This study |
| IU10224 | D39 Δ*cps rpsL1* Δ*bgaA*::*tet*-$P_{Zn}$-$RBS^{ftsA}$-*rodZ*-FLAG (IU1824 X fusion Δ*bgaA*::*tet*-$P_{Zn}$-$RBS^{ftsA}$-*rodZ*-FLAG) | $Tet^R$ $Str^R$ | This study |
| IU10228 | D39 Δ*cps rpsL1 gfp*-$L_1$-*mpgA* markerless | $Str^R$ | (Tsui *et al.*, 2016) |
| IU10254 | D39 Δ*cps rpsL1 ezrA*-$L_2$-*sfgfp* markerless | $Str^R$ | (Perez *et al.*, 2019) |
| IU10294 | D39 Δ*cps rpsL1* Δ*pbp1a* Δ*mpgA*::$P_c$-*erm* Δ*spd_0104*::$P_c$-[*kan-rpsL*$^+$] Δ*spd_1874*::$P_c$-*cat* | $Kan^R$ $Erm^R$ $Cm^R$ | (Tsui *et al.*, 2016) |
| IU10592 | D39 Δ*cps rpsL1* Δ*khpB* | $Str^R$ | (Zheng *et al.*, 2017) |
| IU10664 | D39 Δ*cps khpB*-$L_0$-$FLAG^3$-$P_c$-*erm* | $Erm^R$ | (Zheng *et al.*, 2017) |
| IU10943 | D39 Δ*cps rpsL1 mpgA*(Y488D) markerless Δ*rodA*::$P_c$-*erm* | $Str^R$ $Erm^R$ | (Tsui *et al.*, 2016) |
| IU10947 | D39 Δ*cps rpsL1* Δ*rodZ<>aad9* //Δ*bgaA*::*tet*-$P_{Zn}$-$RBS^{ftsA}$-*rodZ*-FLAG(IU10224 X Δ*rodZ<>aad9* from IU9931) | $Spc^R$ $Tet^R$ | This study |
| IU11005 | D39 Δ*cps rpsL1 sfgfp*-$L_1$-*mpgA* markerless | $Str^R$ | (Perez *et al.*, 2019) |
| IU11119 | D39 Δ*cps ezrA*-$L_0$-*sfgfp*-$P_c$-*cat* | $Cm^R$ | (Perez *et al.*, 2019) |
| IU11157 | D39 Δ*cps rpsL1 isfgfp*-$L_1$-*pbp2x* markerless | $Str^R$ | (Perez *et al.*, 2019) |
| IU11173 | D39 Δ*cps pbp2b<>aad9* //Δ*bgaA*::*tet*-$P_{Zn}$-$RBS^{ftsA}$-*pbp2b*$^+$(IU9990 X *pbp2b<>aad9* from IU7397) | $Spc^R$ $Tet^R$ | This study |
| IU11828 | D39 Δ*cps rodZ*-$HA^3$-Pc-*kan* (IU1945 X fusion amplicon) | $Kan^R$ | This study |
| IU11835 | D39 Δ*cps pbp1a-HA*-$P_c$*kan mreC-L-F*$^3$-$P_c$-*erm* (IU4970 X *pbp1a*-HA-$P_c$-*kan* amplicon from IU7242) | $Kan^R$ $Erm^R$ | This study |
| IU11896 | D39 Δ*cps rodZ*-$L_0$-$FLAG^3$-$P_c$-*erm* *pbp2b*-HA-$P_c$-*kan* (IU6291 x *pbp2b*-HA-$P_c$-*kan* amplicon from IU6933) | $Kan^R$ $Erm^R$ | This study |
| IU11900 | D39 Δ*cps rodZ*-$L_0$-$FLAG^3$-Pc-*erm pbp1a*-HA-$P_c$-*kan* (IU6291 X *pbp1a*-HA-$P_c$-*kan* from IU7242) | $Kan^R$ $Erm^R$ | This study |

| | | | |
|---|---|---|---|
| IU11925 | D39 Δ*cps pbp1a*-FLAG-P$_c$-*erm rodZ*-HA$^3$-P$_c$*kan* (IU5840 X *rodZ*-HA$^3$-P$_c$*kan* from IU11828) | Erm$^R$ Kan$^R$ | This study |
| IU12268 | D39 Δ*cps* Δ*mreC*::P$_c$-*erm*//Δ*bgaA*::*tet*-P$_{Zn}$-RBS$^{ftsA}$-*mreC*$^+$ (IU10222 X Δ*mreC*::P$_c$-*erm* from E149) | Erm$^R$ tet$^R$ | This study |
| IU12272 | D39 Δ*cps rpsL1* Δ*mreC*::P$_c$-[*kan-rpsL*$^+$]//Δ*bgaA*::*tet*-P$_{Zn}$-RBS$^{ftsA}$-m*reC*$^+$(IU10220 X Δ*mreC*::P$_c$-[*kan-rpsL*$^+$] from K49) | Tet$^R$ Kan$^R$ | This study |
| IU12286 | D39 Δ*cps rpsL1* Δ*bgaA*::*tet*-P$_{Zn}$-RBS$^{ftsA}$-*ftsZ* | Str$^R$ Tet$^R$ | (Perez *et al.*, 2019) |
| IU12310 | D39 Δ*cps rpsL1* Δ*bgaA*::*tet*-P$_{Zn}$-RBS$^{ftsA}$-*ftsA* | Str$^R$ Tet$^R$ | (Mura *et al.*, 2017) |
| IU12332 | D39 Δ*cps rpsL1* Δ*cozE*::P$_c$-*erm* Δ*pbp1a* markerless | Str$^R$ Erm$^R$ | (Zheng *et al.*, 2017) |
| IU12345 | D39 Δ*cps rpsL1* Δ*mreC* markerless //Δ*bgaA*::*tet*-P$_{Zn}$-RBS$^{ftsA}$-*mreC*$^+$ (IU12272 X Δ*mreC* markerless fusion amplicon) | Str$^R$ Tet$^R$ | This study |
| IU12515 | D39 Δ*cps rpsL1* Δ*rodZ*::P$_c$-[*kan-rpsL*$^+$] //Δ*bgaA*::*tet*-P$_{Zn}$-RBS$^{ftsA}$-*rodZ*$^+$ (IU9613 X Δ*rodZ*::P$_c$-[*kan-rpsL*$^+$] from K654) | Kan$^R$ Tet$^R$ | This study |
| IU12678 | D39 Δ*cps* Δ*bgaA*::*tet*-P$_{Zn}$-RBS$^{ftsA}$-*cozE* (IU1945 X fusion Δ*bgaA*::*tet*-P$_{Zn}$- RBS$^{ftsA}$-*cozE*$^+$) | Tet$^R$ | This study |
| IU12681 | D39 Δ*cps rpsL1* //Δ*bgaA*::*tet*-P$_{Zn}$-RBS$^{ftsA}$-*cozE*$^+$ (IU1824 X fusion Δ*bgaA*::*tet*-P$_{Zn}$- RBS$^{ftsA}$-*cozE*$^+$) | Str$^R$ Tet$^R$ | This study |
| IU12696 | D39 Δ*cps rpsL1 rodZ*Δ(4-68*)*aa$^d$ markerless //Δ*bgaA*::*tet*-P$_{Zn}$-RBS$^{ftsA}$-*rodZ*$^+$ = ΔHTH (IU12515 X fusion *rodZ*Δ(4-68*)*aa) | Str$^R$ Tet$^R$ | This study |
| IU12699 | D39 Δ*cps rpsL1 rodZ*Δ(196-261)aa markerless //Δ*bgaA*::*tet*-P$_{Zn}$-RBS$^{ftsA}$-*rodZ*$^+$ = ΔDUF (IU12515 X fusion *rodZ*Δ(196-261)aa) amplicon | Str$^R$ Tet$^R$ | This study |
| IU12712 | D39 Δ*cps* Δ*bgaA*::*kan*-t1t2-P$_{ftsA}$-RBS$^{ftsA}$-*ftsA*$^+$ (IU1945 X Δ*bgaA*::*kan*- t1t2-P$_{ftsA}$-RBS$^{ftsA}$-*ftsA* fusion) | Kan$^R$ | This study |
| IU12719 | D39 Δ*cps rpsL1* Δ*bgaA*::*kan*-t1t2-P$_{ftsA}$-RBS$^{ftsA}$-*ftsA* (IU1824 X Δ*bgaA*::*kan*- t1t2-P$_{ftsA}$-RBS$^{ftsA}$-*ftsA*) fusion | Str$^R$ Kan$^R$ | This study |
| IU12738 | D39 Δ*cps rpsL1 rodZ*Δ(21-257*)*aa markerless //Δ*bgaA*::*tet*-P$_{Zn}$-RBS$^{ftsA}$-*rodZ*$^+$ = Δ*rodZ* (IU12515 X fusion *rodZ*Δ(21-257*)*aa amplicon) | Str$^R$ Tet$^R$ | This study |
| IU12788 | D39 Δcps *rpsL1* Δ*bgaA*::*kan*-t1t2-P$_{Zn}$-RBS$_{ftsA}$-*khpA*$^+$ | Kan$^R$ | (Zheng *et al.*, 2017) |
| IU12792 | D39 Δ*cps rpsL1 rodZ*(1-72)aa markerless //Δ*bgaA*::*tet*-P$_{Zn}$-RBS$^{ftsA}$-*rodZ*$^+$ (IU12515 X fusion *rodZ*(1-72)aa amplicon) | Str$^R$ Tet$^R$ | This study |
| IU12794 | D39 Δ*cps rpsL1 rodZ*(1-261)aa markerless //Δ*bgaA*::*tet*-P$_{Zn}$-RBS$^{ftsA}$-*rodZ*$^+$ (IU12515 X fusion *rodZ*(1-261)aa amplicon) | Str$^R$ Tet$^R$ | This study |

| | | | |
|---|---|---|---|
| IU12797 | D39 Δ*cps rpsL1 rodZ*(1-195)aa markerless //Δ*bgaA*::*tet*-$P_{Zn}$-$RBS^{ftsA}$-*rodZ*$^{+}$ (IU12515 X fusion *rodZ*(1-195)aa amplicon) | $Str^R$ $Tet^R$ | This study |
| IU12799 | D39 Δ*cps rpsL1 rodZ*(1-135)aa::TAA-TAG-TGA markerless //Δ*bgaA*::*tet*-$P_{Zn}$-$RBS^{ftsA}$-*rodZ*$^{+}$ (IU12515 X fusion *rodZ(1-135)aa ::*TAA-TAG-TGA) | $Str^R$ $Tet^R$ | This study |
| IU12800 | D39 Δ*cps rpsL1 rodZ*(1-103)aa markerless //Δ*bgaA*::*tet*-$P_{Zn}$-$RBS^{ftsA}$-*rodZ*$^{+}$ (IU12515 X fusion *rodZ*(1-103)aa amplicon) | $Str^R$ $Tet^R$ | This study |
| IU12803 | D39 Δ*cps rpsL1 rodZ*(1-134)aa markerless //ΔbgaA::*tet*-$P_{Zn}$-$RBS^{ftsA}$-*rodZ*$^{+}$ (IU12515 X fusion *rodZ*(1-134)aa amplicon) | $Str^R$ $Tet^R$ | This study |
| IU12915 | D39 Δ*cps rpsL1 ftsZ*-$P_c$-[*kan-rpsL*$^{+}$] Δ*rodZ* markerless //Δ*bgaA*::*tet*-$P_{Zn}$ -$RBS^{ftsA}$-*rodZ*$^{+}$ (IU12738 X *ftsZ*-$P_c$-[*kan-rpsL*$^{+}$] from IU7614) | $Tet^R$ $Kan^R$ | This study |
| IU12917 | D39 Δ*cps rpsL1* $P_c$-*[kan-rpsL*$^{+}$*]-mpgA* Δ*rodZ* markerless //Δ*bgaA*::*tet*-$P_{Zn}$-$RBS^{ftsA}$-*rodZ*$^{+}$ (IU12738 X $P_c$-[*kan-rpsL*$^{+}$]-*mpgA* from IU8980) | $Tet^R$ $Kan^R$ | This study |
| IU12919 | D39 Δ*cps rpsL1* $P_c$-[*kan-rpsL*$^{+}$]-*pbp2x* Δ*rodZ* markerless //Δ*bgaA*::*tet*-$P_{Zn}$-$RBS^{ftsA}$-rodZ$^{+}$ (IU12738 X $P_c$-[*kan-rpsL*$^{+}$] -*pbp2x* from IU8921) | $Tet^R$ $Kan^R$ | This study |
| IU12923 | D39 Δ*cps* Δ*cozE*::$P_c$-*erm* //Δ*bgaA*::*tet*-$P_{Zn}$-$RBS^{ftsA}$-*cozE*$^{+}$ (IU12678 X Δ*cozE*::$P_c$-*erm* from IU12332) | $Erm^R$ $Tet^R$ | This study |
| IU12971 | D39 Δ*cps* Δ*cozE*::$P_c$-*cat*//Δ*bgaA*::*tet*-$P_{Zn}$-$RBS^{ftsA}$-*cozE*$^{+}$ (IU12678 X fusion Δ*cozE*::$P_c$-*cat*) | $Cm^R$ $Tet^R$ | This study |
| IU12993 | D39 Δ*cps rpsL1 ftsZ*-$L_2$-*sfgfp* markerless Δ*rodZ* markerless //Δ*bgaA*::*tet*-$P_{Zn}$-$RBS^{ftsA}$-*rodZ*$^{+}$ (IU12915 X *ftsZ*-L-*sfgfp* amplicon from IU9985) | $Str^R$ $Tet^R$ | This study |
| IU12998 | D39 Δ*cps rpsL1 sfgfp*-$L_1$-*mpgA* markerless Δ*rodZ* markerless //Δ*bgaA::tet*-$P_{Zn}$-$RBS^{ftsA}$-*rodZ*$^{+}$ (IU12917 X *sfgfp*-$L_1$-*mtlG* amplicon from IU11005) | $Str^R$ $Tet^R$ | This study |
| IU13000 | D39 Δ*cps rpsL1 isfgfp*-$L_1$-*pbp2x* markerless Δ*rodZ* markerless //Δ*bgaA*::*tet*-$P_{Zn}$-$RBS^{ftsA}$-*rodZ*$^{+}$ (IU12919 X *isfgfp*-$L_1$-*pbp2x* amplicon from IU11157) | $Str^R$ $Tet^R$ | This study |
| IU13042 | D39 Δ*cps rpsL1 ezrA*-$P_c$-[*kan-rpsL*$^{+}$] Δ*rodZ* markerless //Δ*bgaA*::*tet*-$P_{Zn}$-$RBS^{ftsA}$-*rodZ*$^{+}$ (IU12738 X *ezrA*-$P_c$-[*kan-rpsL*$^{+}$] from IU9077) | $Tet^R$ $Kan^R$ | This study |
| IU13044 | D39 Δ*cps rpsL1 divIVA*-$P_c$-[*kan-rpsL*$^{+}$] Δ*rodZ* markerless //Δ*bgaA*::*tet*-$P_{Zn}$-$RBS^{ftsA}$-*rodZ*$^{+}$ (IU12738 X *divIVA*-$P_c$-[*kan-rpsL*$^{+}$] from IU5648) | $Tet^R$ $Kan^R$ | This study |
| IU13046 | D39 Δ*cps rpsL1* $P_c$-[*kan-rpsL*$^{+}$]-*mapZ* Δ*rodZ* markerless //ΔbgaA::*tet*-$P_{Zn}$-$RBS^{ftsA}$-*rodZ*$^{+}$ (IU12738 X *mapZ*-$P_c$-[*kan-rpsL*$^{+}$] from IU9094) | $Tet^R$ $Kan^R$ | This study |
| IU13058 | D39 Δ*cps rpsL1 ezrA*-$L_2$-*sfgfp* markerless Δ*rodZ* markerless //Δ*bgaA*::*tet*-$P_{Zn}$-*RBS*$^{ftsA}$-*rodZ*$^{+}$ (IU13042 X *ezrA*-$L_2$-*sfgfp* amplicon from IU10254) | $Str^R$ $Tet^R$ | This study |

| | | | |
|---|---|---|---|
| IU13061 | D39 Δ*cps rpsL1 divIVA*-$L_2$-*gfp* markerless Δ*rodZ* markerless //Δ*bgaA*::*tet*-$P_{Zn}$-*RBS*$^{ftsA}$-*rodZ*$^+$ (IU13044 X *divIVA*-$L_2$-*gfp* amplicon from IU9167) | Str$^R$ Tet$^R$ | This study |
| IU13062 | D39 Δ*cps rpsL1 gfp*-$L_1$-*mapZ* markerless Δ*rodZ* markerless //Δ*bgaA*::*tet*-$P_{Zn}$-*RBS*$^{ftsA}$-*rodZ*$^+$ (IU13046 X *gfp*-$L_1$-*mapZ* amplicon from IU9182) | Str$^R$ Tet$^R$ | This study |
| IU13256 | D39 Δ*cps rpsL1* Δ*pbp2a* | Str$^R$ | (Cleverley *et al.*, 2019) |
| IU13440 | D39 Δ*cps pbp2b*-HA-$P_c$-*kan*//Δ*bgaA*::*tet*-$P_{Zn}$-*RBS*$^{ftsA}$-*pbp2b*$^+$(IU11173 X *pbp2b*-HA-$P_c$-*kan* from IU6933) | Tet$^R$ Kan$^R$ | This study |
| IU13454 | D39 Δ*cps rpsL1 rodZ*(ΔHTH)-FLAG-$P_c$-*erm* //Δ*bgaA*::*tet*-$P_{Zn}$-RBS$^{ftsA}$-*rodZ*$^+$ (IU12515 X fusion *rodZ*(ΔHTH-FLAG-$P_c$-*erm*) | Str$^R$ Erm$^R$ Tet$^R$ | This study |
| IU13456 | D39 Δ*cps rpsL1 rodZ*(ΔDUF-FLAG)-$P_c$-*erm* //Δ*bgaA*::*tet*-$P_{Zn}$-RBS$^{ftsA}$-*rodZ*$^+$(IU12515 X fusion *rodZ*(ΔDUF-FLAG)-$P_c$-*erm*) | Str$^R$ Erm$^R$ Tet$^R$ | This study |
| IU13457 | D39 Δ*cps rpsL1 rodZ*-FLAG markerless //Δ*bgaA*::*tet*-$P_{Zn}$-RBS$^{ftsA}$-*rodZ*$^+$(IU12515 X fusion *rodZ*-FLAG markerless) | Str$^R$ Tet$^R$ | This study |
| IU13473 | D39 Δ*cps rpsL1 rodZ*-FLAG-$P_c$-*erm* //Δ*bgaA*::*tet*-$P_{Zn}$-RBS$^{ftsA}$-*rodZ*$^+$ (IU12515 X *rodZ*-FLAG-$P_c$-*erm* from IU6293) | Erm$^R$ Str$^R$ Tet$^R$ | This study |
| IU13555 | D39 Δ*cps rpsL1 rodZ*(1-72aa)-FLAG-$P_c$-*erm* //Δ*bgaA*::*tet*-$P_{Zn}$-RBS$^{ftsA}$-*rodZ*$^+$ (IU12515 x fusion *rodZ*(1-72)aa-FLAG-$P_c$-*erm*) | Erm$^R$ Str$^R$ Tet$^R$ | This study |
| IU13556 | D39 Δ*cps rpsL1 rodZ*(1-134)aa-FLAG-$P_c$-*erm* //Δ*bgaA*::*tet*-$P_{Zn}$-RBS$^{ftsA}$-*rodZ*$^+$ (IU12515 X fusion *rodZ*(1-134)aa-FLAG-$P_c$-*erm*) | Erm$^R$ Str$^R$ Tet$^R$ | This study |
| IU13577 | D39 Δ*cps rpsL1 rodZ(*Δ21-257)-FLAG-$P_c$-*erm* //Δ*bgaA*::*tet*-$P_{Zn}$-RBS$^{ftsA}$-*rodZ*$^+$ (IU12515X fusion Δ*rodZ*-FLAG-$P_c$-*erm)* | Erm$^R$ Str$^R$ Tet$^R$ | This study |
| IU13655 | D39 Δ*cps rpsL1 rodZ*(ΔDUF)-FLAG markerless //Δ*bgaA*::*tet*-$P_{Zn}$-RBS$^{ftsA}$-*rodZ*$^+$ (IU12515 X fusion ΔDUF-FLAG markerless) | Str$^R$ Tet$^R$ | This study |
| IU13656 | D39 Δ*cps rpsL1 rodZ(*Δ21-257)-FLAG markerless //Δ*bgaA*::*tet*-$P_{Zn}$-RBS$^{ftsA}$-*rodZ*$^+$ (IU12515 X fusion Δ*rodZ*-FLAG-markerless) | Str$^R$ Tet$^R$ | This study |
| IU13658 | D39 Δ*cps rpsL1 rodZ*(1-72)aa-FLAG markerless //Δ*bgaA*::*tet*-$P_{Zn}$-RBS$^{ftsA}$-*rodZ*$^+$ (IU12515 X fusion *rodZ*(1-72)aa-FLAG amplicon) | Str$^R$ Tet$^R$ | This study |
| IU13660 | D39 Δ*cps rpsL1 rodZ*(1-134)aa-FLAG markerless //Δ*bgaA*::*tet*-$P_{Zn}$-RBS$^{ftsA}$-*rodZ*$^+$ (IU12515 X fusion *rodZ*(1-134)aa-FLAG) | Str$^R$ Tet$^R$ | This study |
| IU13662 | D39 Δ*cps rpsL1 ftsA'-sfgfp-ftsA'* markerless | Str$^R$ | (Perez *et al.*, 2019) |

| | | | |
|---|---|---|---|
| IU13680 | D39 Δ*cps* Δ*pbp1b*::$P_c$-*aad9*<br>(IU1945 X fusion Δ*pbp1b*::$P_c$-*aad9* amplicon) | $Spc^R$ | This study |
| IU13705 | D39 Δ*cps rpsL1 rodZ*(ΔHTH)-FLAG markerless<br>//Δ*bgaA*::*tet*-$P_{Zn}$-RBS$^{ftsA}$-*rodZ*$^+$<br>(IU12515 X fusion ΔHTH-FLAG markerless) | $Str^R$ $Tet^R$ | This study |
| IU13837 | D39 Δ*cps rpsL1* Δ*bgaA*::$P_c$-*kan*-t1t2-RBS$^{ftsA}$-$P_{Zn}$-<br>*pgsA*$^+$ (IU1824 X fusion amplicon *)* | $Kan^R$ $Str^R$ | This study |
| IU13910 | D39 Δ*cps rpsL1 ht-pbp2x markerless* | $Str^R$ | (Perez *et al.*, 2019) |
| IU13960 | D39 Δ*cps rpsL1* Δ*pgsA*::$P_c$-*erm*<br>//Δ*bgaA::*$P_c$-*kan*-t1t2-RBS$^{ftsA}$-$P_{Zn}$-*pgsA*$^+$<br>(IU13837 X fusion Δ*pgsA*::$P_c$-*erm)* | $Str^R$ $Erm^R$ $Kan^R$ | This study |
| IU14158 | D39 Δ*cps rpsL1 mreC*-$L_0$-FLAG$^3$-$P_c$-*erm*<br>Δ*rodZ* markerless //Δ*bgaA*::*tet*-$P_{Zn}$-*RBS*$^{ftsA}$-*rodZ*$^+$<br>(IU12738 X *mreC*-$L_0$-FLAG$^3$-$P_c$-*erm* from IU4970) | $Str^R$ $Erm^R$ $Tet^R$ | This study |
| IU14160 | D39 Δ*cps rpsL1 stkP*-FLAG$^2$-$P_c$-*erm*<br>Δ*rodZ* markerless //Δ*bgaA*::tet-$P_{Zn}$-*RBS*$^{ftsA}$-*rodZ*$^+$<br>(IU12738 X *stkP*-FLAG$^2$-$P_c$-*erm* from IU7434) | $Str^R$ $Erm^R$ $Tet^R$ | (Tsui *et al.*, 2014) |
| IU14167 | D39 Δ*cps rpsL1 ftsA*-$P_c$-[*kan-rpsL*$^+$]<br>Δ*rodZ* markerless //Δ*bgaA*::*tet*-$P_{Zn}$-*RBS*$^{ftsA}$-*rodZ*$^+$<br>(IU12738 X *ftsA*-$P_c$-[*kan-rpsL*$^+$] from IU7616) | $Kan^R$ $Tet^R$ | This study |
| IU14199 | D39 Δ*cps rpsL1 ftsA'-sfgfp-ftsA'* markerless<br>Δ*rodZ* markerless //Δ*bgaA*::*tet*-$P_{Zn}$-*RBS*$^{ftsA}$-*rodZ*$^+$<br>(IU14167 X *ftsA'-sfgfp-ftsA'* amplicon from IU13662) | $Str^R$ $Tet^R$ | This study |
| IU14431 | D39 Δ*cps rpsL1 pbp2b*-HA-$P_c$-*kan*<br>Δ*rodZ* markerless //Δ*bgaA*::*tet-*$P_{Zn}$*-RBS*$^{ftsA}$-*rodZ*$^+$<br>(IU12738 X *pbp2b*-HA-$P_c$-*kan* from IU6933) | $Str^R$ $Kan^R$ $Tet^R$ | This study |
| IU14433 | D39 Δ*cps rpsL1 gfp*-$L_1$-*mpgA* markerless<br>*ΔrodZ* markerless *//ΔbgaA*::*tet-*$P_{Zn}$*-RBS*$^{ftsA}$-*rodZ*$^+$<br>(IU12917 X *gfp*-$L_1$-*mpgA* markerless from IU10228) | $Str^R$ $Tet^R$ | This study |
| IU14455 | D39 Δ*cps rpsL1 pbp2b*-HA-$P_c$-*kan*<br>(IU1824 X *pbp2b*-HA-$P_c$-*kan* amplicon from IU6933) | $Kan^R$ | This study |
| IU14458 | D39 Δ*cps rpsL1 mreC*-$L_0$-FLAG$^3$ $P_c$-*erm*<br>(IU1824 X *mreC*-$L_0$-FLAG$^3$ amplicon from IU4970) | $Erm^R$ $Str^R$ | This study |
| IU14459 | D39 Δ*cps rpsL1 stkP*-FLAG$^2$-$P_c$-*erm*<br>(IU1824 X *stkP*-FLAG$^2$-$P_c$-*erm* from IU7434) | $Erm^R$ $Str^R$ | This study |
| IU14494 | D39 Δ*cps rpsL1 pbp1a*-FLAG-$P_c$-*erm*<br>(IU1824 X *pbp1a*-FLAG-$P_c$-*erm* from IU5840) | $Erm^R$ $Str^R$ | This study |
| IU14496 | D39 Δ*cps rpsL1 pbp1a*-FLAG $P_c$-*erm*<br>Δ*rodZ* markerless*//ΔbgaA*::*tet*-$P_{Zn}$-RBS$^{ftsA}$-*rodZ*$^+$<br>(IU12738 X *pbp1a*-FLAG-$P_c$-*erm* from IU5840) | $Erm^R$ $Str^R$ $Tet^R$ | This study |
| IU14522 | D39 Δ*cps rpsL1*<br>*rodZ*$^+$-$P_c$-[*kan-rpsL*$^+$]-60bp 3'-*rodZ*$^+$ "direct repeat"<br>(IU1824 X fusion *rodZ*$^+$-$P_c$-[*kan-rpsL*$^+$]-60bp 3'-*rodZ*$^+$*)* | $Kan^R$ | This study |

| | | | |
|---|---|---|---|
| IU14524 | D39 Δ*cps rpsL1 rodZ*$^{+}$-P$_{c}$-[*kan-rpsL*$^{+}$]-60bp 3’-*rodZ*$^{+}$ “direct repeat” // Δ*mreC* markerless//P$_{Zn}$-*mreC*$^{+}$ (IU12345 X*rodZ*$^{+}$-P$_{c}$-[*kan-rpsL*$^{+}$]-60bp 3’-*rodZ*$^{+}$ fusion) | Kan$^{R}$ Tet$^{R}$ | This study |
| IU14528 | D39 Δ*cps rpsL1* Δ*cozE*::P$_{c}$-[*kan-rpsL*$^{+}$] //Δ*bgaA*::*tet*-P$_{Zn}$-RBS$^{ftsA}$-*cozE*$^{+}$ (IU12681 X fusion Δ*cozE*::P$_{c}$-[*kan-rpsL*$^{+}$] | Kan$^{R}$ Tet$^{R}$ | This study |
| IU14594 | D39 Δ*cps rpsL1 rodZ*-FLAG markerless (IU14522 X *rodZ*-FLAG markerless from IU13457) | Str$^{R}$ | This study |
| IU14598 | D39 Δ*cps rpsL1 rodZ*-FLAG markerless Δ*mreC* markerless//Δ*bgaA*::*tet*-P$_{Zn}$-RBS$^{ftsA}$-*mreC*$^{+}$ (IU14524 X *rodZ*-FLAG markerless from IU13457) | Str$^{R}$ Tet$^{R}$ | This study |
| IU14697 | D39 Δ*cps rpsL1* Δ*pbp1b* (IU7850 X fusion amplicon) | Str$^{R}$ | This study |
| IU14738 | D39 Δ*cps rpsL1 iht*-L$_{6}$-*mapZ* markerless | Str$^{R}$ | (Perez *et al.*, 2019) |
| IU14773 | D39 Δ*cps rpsL1 pbp2b*-HA-P$_{c}$-*kan* Δ*mreC* markerless*//*Δ*bgaA*::*tet*-P$_{Zn}$-RBS$^{ftsA}$-*mreC*$^{+}$ (IU12345 X *pbp2b*-HA-P$_{c}$-*kan* from IU6933) | Kan$^{R}$ Tet$^{R}$ | This study |
| IU14927 | D39 Δ*cps rpsL1 iht*-L$_{6}$-*pbp2x* markerless | Str$^{R}$ | (Perez *et al.*, 2019) |
| IU15337 | D39 Δ*cps pbp2b*(Q56L)-HA-P$_{c}$-*kan* //Δbga::tet-P$_{Zn}$-*pbp2b*$^{+}$(IU11173 X fusion amplicon) | Kan$^{R}$ Tet$^{R}$ | This study |
| IU15340 | D39 Δ*cps pbp2b*(T57A)-HA-P$_{c}$-*kan* //Δbga::tet-P$_{Zn}$-*pbp2b*$^{+}$(IU11173 X fusion amplicon) | Kan$^{R}$ Tet$^{R}$ | This study |
| IU15341 | D39 Δ*cps pbp2b*(T57N)-HA-P$_{c}$-*kan* //Δbga::tet-P$_{Zn}$-*pbp2b*$^{+}$(IU11173 X fusion amplicon) | Kan$^{R}$ Tet$^{R}$ | This study |
| IU15343 | D39 Δ*cps pbp2b*(T57R)-HA-P$_{c}$-*kan* //Δbga::tet-P$_{Zn}$-*pbp2b*$^{+}$(IU11173 X fusion amplicon) | Kan$^{R}$ Tet$^{R}$ | This study |
| IU15347 | D39 Δ*cps pbp2b*(I290A)-HA-P$_{c}$-*kan* //Δbga::tet-P$_{Zn}$-*pbp2b*$^{+}$(IU11173 X fusion amplicon) | Kan$^{R}$ Tet$^{R}$ | This study |
| IU15329 | D39 Δ*cps rpsL1 rodZ*-L-FLAG$^{3}$-P$_{c}$-*erm gfp*-L-*mpgA markerless* (IU10228 X *rodZ*-L-FLAG$^{3}$-P$_{c}$-*erm* from IU6291) | Erm$^{R}$ Str$^{R}$ | This study |
| IU15605 | D39 *rpsL1* Δ*bgaA*::*tet*-P$_{Zn}$-RBS$^{ftsA}$-*rodZ*$^{+}$(IU1781 X Δ*bgaA*::*tet*-P$_{Zn}$-RBS$^{ftsA}$-*rodZ*$^{+}$ amplicon from IU9765) | Str$^{R}$ Tet$^{R}$ | This study |
| IU15628 | D39 Δ*cps rpsL1 rodZ*(Y51A F55A Y59A)-Flag // Δ*bgaA*::*tet*-P$_{Zn}$-RBS$^{ftsA}$-*rodZ*$^{+}$ (IU12515 X fusion *rodZ*(Y51A F55A Y59A)-Flag amplicon | Str$^{R}$ Tet$^{R}$ | This study |
| IU15645 | D39 *cps*$^{+}$ *rpsL1* Δ*rodZ*::P$_{c}$-[*kan-rpsL*$^{+}$] //Δ*bgaA*::*tet*-P$_{Zn}$-RBS$^{ftsA}$-*rodZ*$^{+}$(IU15605 X Δ*rodZ*::P$_{c}$-[*kan-rpsL*$^{+}$] from IU12515) | Kan$^{R}$ Tet$^{R}$ | This study |
| IU15901 | D39 Δ*cps rpsL1 pbp1a*-FLAG-P$_{c}$-*erm* Δ*mreC* markerless*//* Δ*bgaA*::*tet*-P$_{Zn}$-RBS$^{ftsA}$-*mreC*$^{+}$ (IU12345 X *pbp1a*-FLAG-P$_{c}$-*erm* from IU14494) | Erm$^{R}$ Str$^{R}$ Tet$^{R}$ | This study |
| IU15907 | D39 Δ*cps rpsL1* P$_{c}$-[*kan-rpsL*$^{+}$]-*rodA*$^{+}$ (IU1824 x fusion P$_{c}$-[*kan-rpsL*$^{+}$]-*rodA* amplicon) | Kan$^{R}$ | This study |

| | | | |
|---|---|---|---|
| IU15928 | D39 Δ*cps rpsL1 iht*-$L_6$-*pbp2b* markerless (IU9023 X fusion *iht*-$L_6$-*pbp2b* amplicon) | $Str^R$ | This study |
| IU15970 | D39 Δ*cps rpsL1 iht*-$L_6$-*rodA* markerless (IU15907 X fusion *iht*-$L_6$-*rodA* amplicon) | $Str^R$ | This study |
| IU15987 | D39 Δ*cps rpsL1* Δ*stkP*::$P_c$-*erm* with suppression mutation | $Erm^R$ $Str^R$ | This study |
| IU16046 | D39 Δ*cps rpsL1* $P_c$-[*kan-rpsL*$^+$]-*pbp2b*$^+$ Δ*rodZ* //Δ*bgaA*::*tet*-$P_{Zn}$-RBS$^{ftsA}$-*rodZ*$^+$ (IU12738 X $P_c$-[*kan-rpsL*$^+$]-*pbp2b*$^+$ from IU9023) | $Kan^R$ $Tet^R$ | This study |
| IU16048 | D39 Δ*cps rpsL1* $P_c$-[*kan-rpsL*$^+$]-*rodA*$^+$ Δ*rodZ* //Δ*bgaA*::*tet*-$P_{Zn}$-RBS$^{ftsA}$-*rodZ*$^+$ (IU12738 X $P_c$-[*kan-rpsL*$^+$]-*rodA*$^+$ from IU15907) | $Kan^R$ $Tet^R$ | This study |
| IU16050 | D39 Δ*cps rpsL1* $P_c$-[*kan-rpsL*$^+$]-*pbp2x*$^+$ Δ*rodZ* //Δ*bgaA*::*tet*-$P_{Zn}$-RBS$^{ftsA}$-*rodZ*$^+$ (IU12738 X $P_c$-[*kan-rpsL*$^+$]-*pbp2x*$^+$ from IU8921) | $Kan^R$ $Tet^R$ | This study |
| IU16058 | D39 Δ*cps rpsL1 iht*-$L_6$-*pbp2b* markerless Δ*rodZ* markerless// Δ*bgaA*::*tet*-$P_{Zn}$-RBS$^{ftsA}$-*rodZ*$^+$ (IU16046 X *iht-pbp2b* from IU15928) | $Str^R$ $Tet^R$ | This study |
| IU16060 | D39 Δ*cps rpsL1 iht-rodA*$^+$ markerless Δ*rodZ* markerless// Δ*bgaA*::*tet*-$P_{Zn}$-RBS$^{ftsA}$-*rodZ*$^+$ (IU16048 X *iht-rodA* from IU15970) | $Str^R$ $Tet^R$ | This study |
| IU16062 | D39 Δ*cps rpsL1 ht-pbp2x* markerless Δ*rodZ* markerless// Δ*bgaA*::*tet*-$P_{Zn}$-RBS$^{ftsA}$-*rodZ*$^+$ (IU16050 X *ht-pbp2x* amplicon from IU13910) | $Str^R$ $Tet^R$ | This study |
| IU16126 | D39 Δ*cps rpsL1 ftsW*-$L_2$-*gfp* markerless *rodZ*-$L_0$-$FLAG^3$-$P_c$-*erm* (IU8918 X *rodZ*-L-$FLAG^3$-$P_c$-*erm* from IU6291) | $Str^R$ $Erm^R$ | This study |
| IU16128 | D39 Δ*cps rpsL1 iht-rodA* markerless *rodZ*-$L_0$-$FLAG^3$-$P_c$-*erm* (IU15970 X *rodZ*-$L_0$-$FLAG^3$-$P_c$-*erm* from IU6291) | $Str^R$ $Erm^R$ | This study |
| IU16252 | D39 Δ*cps rpsL1* [*kan-rpsL*$^+$]-*pbp2b* Δ*mreC* markerless //Δ*bgaA*::*tet*-$P_{Zn}$-RBS$^{ftsA}$-*mreC*$^+$ (IU12345 X [*kan-rpsL*$^+$]-*pbp2b* from IU9023) | $Kan^R$ $Tet^R$ | This study |
| IU16254 | D39 Δ*cps rpsL1* [*kan-rpsL*$^+$]-*rodA* Δ*mreC* markerless //Δ*bgaA*::*tet*-$P_{Zn}$-RBS$^{ftsA}$-*mreC*$^+$ (IU12345 X [*kan-rpsL*$^+$]-*rodA* from IU15907) | $Kan^R$ $Tet^R$ | This study |
| IU16281 | D39 Δ*cps rpsL1 iht*-$L_6$-*pbp2b* markerless Δ*mreC* markerless // Δ*bgaA*::*tet*-$P_{Zn}$-RBS$^{ftsA}$-*mreC*$^+$ (IU16252 X *iht*-$L_6$-*pbp2b* markerless from IU15928) | $Str^R$ $Tet^R$ | This study |
| IU16283 | D39 Δ*cps rpsL1 iht*-$L_6$-*rodA* markerless Δ*mreC* markerless // Δ*bgaA*::*tet*-$P_{Zn}$-RBS$^{ftsA}$-*mreC*$^+$ (IU16254 X *iht*-$L_6$-*rodA* amplicon from IU15970) | $Str^R$ $Tet^R$ | This study |

| | | | |
|---|---|---|---|
| IU16307 | D39 Δ*cps rpsL1* [*kan-rpsL*$^{+}$]-*pbp2x* Δ*mreC* markerless // Δ*bgaA*::*tet*-$P_{Zn}$-RBS$^{ftsA}$-*mreC*$^{+}$ (IU12345 X [*kan-rpsL*$^{+}$]-*pbp2x* from IU8921) | Kan$^{R}$ Tet$^{R}$ | This study |
| IU16326 | D39 Δ*cps rpsL1 iht*-$L_6$-*pbp2x* markerless Δ*mreC* markerless // Δ*bgaA*::*tet*-$P_{Zn}$-RBS$^{ftsA}$-*mreC*$^{+}$ (IU16307 X *iht*-$L_6$-*pbp2x* from IU14927) | Str$^{R}$ Tet$^{R}$ | This study |
| IU16338 | D39 Δ*cps rpsL1 rodZ*-FLAG-$P_c$-*erm* //Δ*bgaA*::*tet*-$P_{Zn}$-RBS$^{ftsA}$-*rodZ*-FLAG (IU10224 X *rodZ*-FLAG-$P_c$-*erm* from IU6293) | Erm$^{R}$ Tet$^{R}$ | This study |
| IU16344 | D39 Δ*cps rpsL1 iht*-$L_6$-*mreC* markerless (IU10103 X *iht*-$L_6$-*mreC* fusion amplicon) | Str$^{R}$ | This study |
| IU16881 | D39 Δ*cps rpsL1* $P_c$-[*kan-rpsL*$^{+}$]-*mreC*$^{+}$ Δ*rodZ* // Δ*bgaA*::*tet*-$P_{Zn}$-RBS$^{ftsA}$-*rodZ*$^{+}$ (IU12738 X [*kan-rpsL*$^{+}$]-*mreC*$^{+}$ from IU10103) | Kan$^{R}$ Tet$^{R}$ | This study |
| IU16920 | D39 Δ*cps rpsL1 iht*-$L_6$-*mreC* markerless Δ*rodZ* // Δ*bgaA*::*tet*-$P_{Zn}$-RBS$^{ftsA}$-*rodZ*$^{+}$ (IU16881 X *iht*-$L_6$-*mreC* amplicon from IU16344) | Str$^{R}$ Tet$^{R}$ | This study |
| IU17010 | D39 Δ*cps rpsL1* [*kan-rpsL*$^{+}$]-*ftsA* markerless Δ*rodZ* markerless // Δ*bgaA*::*tet*-$P_{Zn}$-RBS$^{ftsA}$-*rodZ*$^{+}$ (IU12738 X [*kan-rpsL*$^{+}$]-*ftsA* from IU9767) | Kan$^{R}$ Tet$^{R}$ | This study |
| IU17022 | D39 Δ*cps rpsL1* Flag-ftsA markerless Δ*rodZ* markerless // Δ*bgaA*::*tet*-$P_{Zn}$-RBS$^{ftsA}$-*rodZ*$^{+}$ (IU17010 X Flag-*ftsA* markerless from IU9969) | Str$^{R}$ Tet$^{R}$ | This study |
| IU17024 | D39 Δ*cps rpsL1 gfp-ftsA* markerless Δ*rodZ* markerless // Δ*bgaA*::*tet*-$P_{Zn}$-RBS$^{ftsA}$-*rodZ*$^{+}$ (IU17010 X *gfp-ftsA* markerless from IU10035) | Str$^{R}$ Tet$^{R}$ | This study |
| IU17817 | D39 Δ*cps mreC*-$L_0$-FLAG$^{3}$-$P_c$-*erm* Δ*pbp2a*::$P_c$-[*kan-rpsL*$^{+}$] (IU4970 X Δ*pbp2a*::$P_c$-[*kan-rpsL*$^{+}$] from K166) | Erm$^{R}$ Kan$^{R}$ | This study |
| IU17821 | D39 Δ*cps rodZ*-$L_0$-FLAG$^{3}$-$P_c$-*erm* Δ*pbp2a*::$P_c$-[*kan-rpsL*$^{+}$] (IU6291 X Δ*pbp2a*::$P_c$-[*kan-rpsL*$^{+}$] from K166) | Erm$^{R}$ Kan$^{R}$ | This study |
| IU17873 | D39 Δ*cps khpA*-$L_0$-FLAG$^{3}$-$P_c$-*erm rodZ*-HA$^{3}$-$P_c$*kan* (IU9602 X *rodZ*-HA$^{3}$-$P_c$*kan* from IU11828) | Erm$^{R}$ Kan$^{R}$ | This study |
| IU17877 | D39 Δ*cps khpB*-$L_0$-FLAG$^{3}$-$P_c$-*erm rodZ*-HA$^{3}$-$P_c$*kan* (IU10664 X *rodZ*-HA$^{3}$-$P_c$*kan* from IU11828) | Erm$^{R}$ Kan$^{R}$ | This study |
| IU17883 | D39 Δ*cps rpsL1* Δ*stkP*::$P_c$-*erm rodZ*-HA$^{3}$-$P_c$*kan* with Δ*stkP* suppressor mutation (IU15987 X *rodZ*-HA$^{3}$-$P_c$*kan* from IU11828) | Str$^{R}$ Erm$^{R}$ Kan$^{R}$ | This study |
| IU18579 | D39 Δ*cps rpsL1* Δ*pbp1a* markerless (single colony isolate of IU6741) | Str$^{R}$ | This study |

[a]Strains were constructed as described in the *Experimental Procedures.* :: indicates an insertion into a region, whereas <> indicates an exact reading frame replacement. Linkers used to synthesize fusion amplicons are listed in Supplemental Table S2. Primers used

to synthesize fusion amplicons are listed in Supplemental Table S3. The amino acid sequence of the FLAG epitope is DYKDDDDK (Wayne *et al.*, 2010). The Myc epitope amino acid sequence is EQKLISEEDL (Evan *et al.*, 1985), and the HA epitope amino acid sequence is YPYDVPDYA (Tu *et al.*, 1998).

[b]Antibiotic resistance markers used are: $Erm^R$, erythromycin; $Kan^R$, kanamycin; $Spc^R$, spectinomycin; $Tet^R$, tetracycline; $Str^R$, streptomycin; $Cm^R$, chloramphenicol. Markerless indicates an antibiotic cassette or marker is not present, *e.g.* clean deletion.

[c]IU1824 (D39 Δ*cps rpsL1*) harbors a spontaneous GC→TA drift mutation 4 bp upstream of the -35 box of the $P_{ftsA}$ promoter (-119 bp upstream of the *ftsA* start codon). This region does not contain indirect repeats or an overt regulatory role. Expression of *ftsA* in IU1824 is comparable to levels in IU1945 (D39 Δ*cps*), which lacks the drift mutation (data not shown).

[d]aa, amino acid

48 **Table S2.** Linker sequences used in this study

| Linker | Nucleotide sequence | Linker aa sequence | Reference |
|---|---|---|---|
| $L_0$ | ggttccgctggctccgctgctggttctggc | GSAGSAAGSG | (Wayne *et al.*, 2010) |
| $L_1$ | ctcgagggatccgga | LEGSG | (Fleurie *et al.*, 2014) |
| $L_2$ | aaactagacatcgagttcctgcag | KLDIEFLQ | (Fleurie *et al.*, 2014) |
| $L_6$ | ttggaaggatcaggacaaggaccagga tctggtcaaggttctggt | LEGSGQGPGSGQGSG | (Perez *et al.*, 2019) |

49

**Table S3.** B2H plasmids used in this study

| **Name** | **Relevant characteristics** | **Construct** | **Reference** |
|---|---|---|---|
| ***S. pneumoniae* B2H** | | | |
| pMKV24 | *kan* $P_{lac}$-*cya(T25)-ftsA* | T25-FtsA | (Krupka *et al.*, 2012) |
| pMKV19 | *amp* $P_{lac}$-*cya(T18)-ftsA* | T18-FtsA | (Krupka *et al.*, 2012) |
| pKNT25_*ftsZ* | *kan* $P_{lac}$-*ftsZ-cya(T25)* | FtsZ-T25 | (Rued *et al.*, 2017) |
| pUT18_*ftsZ* | *amp* $P_{lac}$-*ftsZ-cya(T18)* | FtsZ-T18 | (Rued *et al.*, 2017) |
| pKNT25_*ezrA* | *kan* $P_{lac}$-*ezrA-cya(T25)* | EzrA-T25 | (Rued *et al.*, 2017) |
| pUT18_*ezrA* | *amp* $P_{lac}$-*ezrA -cya(T18)* | EzrA-T18 | (Rued *et al.*, 2017) |
| pKNT25_*divIVA* | *kan* $P_{lac}$-*divIVA-cya(T25)* | DivIVA-T25 | (Rued *et al.*, 2017) |
| pUT18_*divIVA* | *amp* $P_{lac}$-*divIVA-cya(T18)* | DivIVA-T18 | (Rued *et al.*, 2017) |
| pKNT25_*gpsB* | *kan* $P_{lac}$-*gpsB-cya(T25)* | GpsB-T25 | (Rued *et al.*, 2017) |
| pUT18_*gpsB* | *amp* $P_{lac}$-*gpsB-cya(T18)* | GpsB-T18 | (Rued *et al.*, 2017) |
| pKNT25_*stkP* | *kan* $P_{lac}$-*stkP-cya(T25)* | StkP-T25 | (Rued *et al.*, 2017) |
| pUT18_*stkP* | *amp* $P_{lac}$-*stkP-cya(T18)* | StkP-T18 | (Rued *et al.*, 2017) |
| pFC113 | *kan* $P_{lac}$-*cya(T25)-mreC* | T25-MreC | (Cleverley *et al.*, 2019) |
| pFC114 | *amp* $P_{lac}$-*cya(T18)-mreC* | T18-MreC | (Cleverley *et al.*, 2019) |
| pFC115 | *kan* $P_{lac}$-*cya(T25)-pbp2a* | T25-PBP2a | (Cleverley *et al.*, 2019) |
| pFC116 | *amp* $P_{lac}$-*cya(T18)-pbp2a* | T18-PBP2a | (Cleverley *et al.*, 2019) |
| pFC123 | *kan* $P_{lac}$-*cya(T25)-pbp1a* | T25-PBP1a | (Cleverley *et al.*, 2019) |
| pFC124 | *amp* $P_{lac}$-*cya(T18)-pbp1a* | T18-PBP1a | (Cleverley *et al.*, 2019) |
| pFC125 | *kan* $P_{lac}$-*cya(T25)-pbp2b* | T25-PBP2b | (Cleverley *et al.*, 2019) |
| pFC126 | *amp* $P_{lac}$-*cya(T18)-pbp2b* | T18-PBP2b | (Cleverley *et al.*, 2019) |
| pFC127 | *kan* $P_{lac}$-*cya(T25)-pbp2x* | T25-PBP2x | (Cleverley *et al.*, 2019) |
| pFC128 | *amp* $P_{lac}$-*cya(T18)-pbp2x* | T18-PBP2x | (Cleverley *et al.*, 2019) |
| pMBM147 | *kan* $P_{lac}$-*cya(T25)-mpgA* | T25-MpgA | (Perez *et al.*, 2021) |
| pMBM148 | *amp* $P_{lac}$-*cya(T18)-mpgA* | T18-MpgA | (Perez *et al.*, 2021) |
| pMBM151 | *kan* $P_{lac}$-*cya(T25)-rodA* | T25-RodA | (Perez *et al.*, 2021) |
| pMBM152 | *amp* $P_{lac}$-*cya(T18)-rodA* | T18-RodA | (Perez *et al.*, 2021) |
| pMBM153 | *kan* $P_{lac}$-*cya(T25)-ftsW* | T25-FtsW | (Perez *et al.*, 2021) |
| pMBM154 | *amp* $P_{lac}$-*cya(T18)-ftsW* | T18-FtsW | (Perez *et al.*, 2021) |
| pDDM169 | *kan* $P_{lac}$-*mreD-cya(T25)* | MreD-T25 | (Perez *et al.*, 2021) |
| pDDM170 | *amp* $P_{lac}$-*mreD-cya(T18)* | MreD-T18 | (Perez *et al.*, 2021) |
| pFC141 | *kan* $P_{lac}$-*cya(T25)-rodZ* | T25-RodZ | (Perez *et al.*, 2021) |
| pFC142 | *amp* $P_{lac}$-*cya(T18)-rodZ* | T18-RodZ | (Perez *et al.*, 2021) |
| pMBM143 | *kan* $P_{lac}$-*cya(T25)-rodZ* ΔHTH | T25-RodZ ΔHTH | This work |
| pMBM144 | *amp* $P_{lac}$-*cya(T18)-rodZ* ΔHTH | T18-RodZ ΔHTH | This work |
| pMBM145 | *kan* $P_{lac}$-*cya(T25)-rodZ* ΔDUF | T25-RodZ ΔDUF | This work |
| pMBM146 | *amp* $P_{lac}$-*cya(T18)-rodZ* ΔDUF | T18-RodZ ΔDUF | This work |
| pAZM201 | *kan Plac-cya(T25)-pbp1b* | T25-PBP1b | This work |
| pAZM202 | *kan Plac-cya(T18)- pbp1b* | T18-PBP1b | This work |

**Table S4.** Oligonucleotide primers used in this study

| Primer | Sequence (5'-3') | Template | Amplicon Product |
|---|---|---|---|
| **For strain constructions** | | | |
| **For construction of E149 (Δ*mreC*::$P_c$-*erm*)** | | | |
| P104 | AATGAGACGTGTTGCCATTGCAGG | D39[a] | upstream + 5' 60 bp of *mreC* |
| P118 | CATTATCCATTAAAAATCAAACGGATCCTACACAAGCA GAACAGTGACAAAAACAATAAT | | |
| kan rpsL forward | TAGGATCCGTTTGATTTTTAATGGATAATG | Pc-*erm* cassette | $P_c$-*erm* |
| kan rpsL reverse | GGGCCCCTTTCCTTATGCTTTTG | | |
| P119 | CAAAAGCATAAGGAAAGGGGCCCGTTAAATTGAGTGC AGATACTCATAATGTAGATGTG | D39 | 3' 57 bp *mreC* + downstream |
| P107 | TGTCGCTTTCTCAGCAGCAAGACT | | |
| **For construction of K49 (Δ*mreC*::$P_c$-[*kan-rpsL*$^+$])** | | | |
| P104 | AATGAGACGTGTTGCCATTGCAGG | D39 | upstream + 5' 60 bp *mreC* |
| P118 | CATTATCCATTAAAAATCAAACGGATCCTACACAAGCA GAACAGTGACAAAAACAATAAT | | |
| kan rpsL forward | TAGGATCCGTTTGATTTTTAATGGATAATG | Pc-[*kan-rpsL*$^+$] cassette | $P_c$-[*kan-rpsL*$^+$] |
| kan rpsL reverse | GGGCCCCTTTCCTTATGCTTTTG | | |
| P119 | CAAAAGCATAAGGAAAGGGGCCCGTTAAATTGAGTGC AGATACTCATAATGTAGATGTG | D39 | 3' 57 bp *mreC* + downstream |
| P107 | TGTCGCTTTCTCAGCAGCAAGACT | | |
| **For construction of IU6291 (*rodZ*-$L_0$-FLAG$^3$-$P_c$-*erm*)** | | | |
| SS01 | GCAACGCAATATGATGCTTTTGAAAATGGTG | D39 | upstream to *rodZ* |
| SS02 | CGGAGCCAGCGGAACCATTTTTAGTAAAGGTTACAGT GATTTGTCCAG | | |
| SS03 | GACAAATCACTGTAACCTTTACTAAAAATGGTTCCGCT GGCTCCGC | IU4970 | $L_0$-FLAG$^3$ $P_c$-*erm* |
| SS04 | TCTTTTTTCATTCGTTTTTCCTTATTTCCTCCCGTTAAA TAATAGATAACTATTAAAAAT | | |
| SS05 | AGTTATCTATTATTTAACGGGAGGAAATAAGGAAAAAC GAATGAAAAAAGAACAAA | D39 | downstream |
| P1385 | ACAACACCTGCAATGGCCACACGTTGCTTT | | |
| **For construction of IU6293 (*rodZ*-FLAG-$P_c$-*erm*)** | | | |
| SS01 | GCAACGCAATATGATGCTTTTGAAAATGGTG | D39 | upstream to *rodZ*-FLAG |
| SS06 | GTTATTTATCATCATCATCTTTATAATCATTTTTAGTAA AGGTTACAGTGATTTGTCCAG | | |
| SS07 | ACAAATCACTGTAACCTTTACTAAAAAT | IU4970 | FLAG-$P_c$-*erm* |
| SS04 | TCTTTTTTCATTCGTTTTTCCTTATTTCCTCCCGTTAAA TAATAGATAACTATTAAAAAT | | |
| SS05 | AGTTATCTATTATTTAACGGGAGGAAATAAGGAAAAAC GAATGAAAAAAGAACAAA | D39 | downstream |

| | | | |
|---|---|---|---|
| P1385 | ACAACACCTGCAATGGCCACACGTTGCTTT | | |
| **For construction of IU6987 (Δ*rodZ*::P$_c$-*aad9*)** | | | |
| TT329 | CAACTGATATAGTTGGAAGTGAGGAGTCCATTTCCC | E655 | upstream + 5' 60 bp of *rodZ* + P$_c$ |
| TT383 | ATGTATTCAAATATATCCTCCTCACTTATTATTTCCTTC CTCTTTTCTACAGTATTTAAA | | |
| TT384 | ACTGTAGAAAAGAGGAAGGAAATAATAAGTGAGGAGG ATATATTTGAATACATACGAACA | IU1751 | *aad9* |
| TT385 | CTTTTGGACGTTTAGTACCGTATTATAATTTTTTTAATC TGTTATTTAAATAGTTTATAG | | |
| TT386 | CTATTTAAATAACAGATTAAAAAAATTATAATACGGTAC TAAACGTCCAAAAGCATAAGG | D39 | 3' 57 bp *rodZ* + downstream |
| P1385 | ACAACACCTGCAATGGCCACACGTTGCTTT | | |
| **For construction of IU7054 (Δ*bgaA*::*kan*-t1t2-P$_{ftsA}$-*ftsZ*)** | | | |
| P146 | TGGCCATTCATCGCTGGTCGTGCTGAAAT | IU6397 | Δ*bgaA*::*kan*-*t1t2*-P$_{ftsA}$ |
| TT393 | CAGCTGTATCAAATGAAAATGTCATTACATCGCTTCCT CTCTATCTTCCAAGT | | |
| TT394 | GGAAGATAGAGAGGAAGCGATGTAATGACATTTTCAT TTGATACAGCTGCTG | D39 | *ftsZ* |
| TT395 | CAACTGGTTTATGAGAAAGTAAGTTCTTCTAACGATTT TTGAAAAATGGAGGTGTATC | | |
| TT396 | CCTCCATTTTTCAAAAATCGTTAGAAGAACTTACTTTCT CATAAACCAGTTGCTG | D39 | *3' bgaA'* |
| CS121 | GCTTTCTTGAGGCAATTCACTTGGTGC | | |
| **For construction of IU7068 (*rodZ*-Myc-P$_c$-*kan*)** | | | |
| SS01 | GCAACGCAATATGATGCTTTTGAAAATGGTG | D39 | upstream to *rodZ* |
| TT402 | GATCTTCTTCAGAAATAAGTTTTTGTTCATTTTTAGTAA AGGTTACAGTGATTTGTCCAG | | |
| TT403 | AAATCACTGTAACCTTTACTAAAAATGAACAAAAACTT ATTTCTGAAGAAGATCTTTAAC | IU6962 | Myc-P$_c$-*kan* |
| TT404 | GTTCTTTTTTCATTCGTTTTTCCCTAAAACAATTCATCC AGTAAAATATAATATTTTATT | | |
| TT405 | AATATTATATTTTACTGGATGAATTGTTTTAGGGAAAAA CGAATGAAAAAAGAACAAATT | D39 | downstream |
| P1385 | ACAACACCTGCAATGGCCACACGTTGCTTT | | |
| **For construction of IU9613 (Δ*bgaA::tet-P$_{Zn}$-RBS$^{ftsA}$-rodZ$^+$*) = P$_{Zn}$-*rodZ*** | | | |
| TT657 | CGCCCCAAGTTCATCACCAATGACATCAAC | IU8122 | *bgaA'* *tet*-P$_{Zn}$-*RBS$^{ftsA}$* |
| TT769 | CCTCTCCAATTGTTTTTTTTCTCATTACATCGCTTCCTC TCTATCTTCCTTGT | | |
| TT770 | GGAAGATAGAGAGGAAGCGATGTAATGAGAAAAAAAA CAATTGGAGAGGTTTTAC | D39 | *rodZ* |
| TT771 | ACTGGTTTATGAGAAAGTAAGTTCTTTTAATTTTTAGTA AAGGTTACAGTGATTTGTCCA | | |
| TT772 | AAATCACTGTAACCTTTACTAAAAATTAAAAGAACTTAC TTTCTCATAAACCAGTTGCTG | D39 | *bgaA'* to downstream |
| CS121 | GCTTTCTTGAGGCAATTCACTTGGTGC | | |

| | | | |
|---|---|---|---|
| **For construction of IU9990 (Δ*bgaA*::*tet*-$P_{Zn}$-$RBS^{ftsA}$-*pbp2b*$^{+}$) = $P_{Zn}$-*pbp2b*$^{+}$** | | | |
| P146 | TGGCCATTCATCGCTGGTCGTGCTGAAAT | IU9613 | *bgaA'* *tet*-$P_{Zn}$-$RBS^{ftsA}$ |
| BR70 | GTTAAATTTTCTCATACAAATCAGTCTCATTACATCGCTTCCTCTCTATCTTCCTTGTTA | | |
| BR69 | AGGAAGATAGAGAGGAAGCGATGTAATGAGACTGATTTGTATGAGAAAATTTAACAGC | D39 | *pbp2b* |
| BR72 | AACTGGTTTATGAGAAAGTAAGTTCTTCTAATTCATTGGATGGTATTTTTGATACAGATT | | |
| BR71 | GTATCAAAAATACCATCCAATGAATTAGAAGAACTTACTTTCTCATAAACCAGTTGCTGC | | *bgaA'* to downstream |
| CS121 | GCTTTCTTGAGGCAATTCACTTGGTGC | | |
| **For construction of IU10103 ($P_c$-[*kan-rpsL*$^{+}$]-*mreC*$^{+}$)** | | | |
| P104 | AATGAGACGTGTTGCCATTGCAGG | D39 | *spd_2046* + 9 bp downstream |
| TT831 | CCATTAAAAATCAAACGGATCCTAAAGCTACTAAGATTTTAAGAAAAATAAACAACAACC | | |
| TT832 | TGTTTATTTTTCTTAAAATCTTAGTAGCTTTAGGATCCGTTTGATTTTTAATGGATAATG | Pc-[*kan-rpsL*$^{+}$] cassette | $P_c$-[*kan-rpsL*$^{+}$] |
| kan rpsL reverse | GGGCCCCTTTCCTTATGCTTTTG | | |
| TT833 | CAAAAGCATAAGGAAAGGGGCCCTCAGGAATTGATAAAAAGTTACTGTAACAGTTTTT | D39 | *52 bp upstream* + *mreC* |
| TT830 | CAGTAGTCACCTTATCTCCCGCACTAATATCGC | | |
| **For construction of IU10220 and IU10222 (Δ*bgaA*::*tet*-$P_{Zn}$-$RBS^{ftsA}$-*mreC*$^{+}$) = $P_{Zn}$-*mreC*$^{+}$** | | | |
| TT657 | CGCCCCAAGTTCATCACCAATGACATCAAC | IU9613 | *bgaA'* *tet*-$P_{Zn}$-$RBS^{ftsA}$ |
| TT865 | GACATATTTTGATTTTTTAAAACGGTTCATTACATCGCTTCCTCTCTATCTTCCTTGTTA | | |
| TT866 | ACAAGGAAGATAGAGAGGAAGCGATGTAATGAACCGTTTTAAAAAATCAAAATATGTCAT | D39 | *mreC* |
| TT867 | AACTGGTTTATGAGAAAGTAAGTTCTTTTATGAATTCCCCACTAATTCTATCACATCTAC | | |
| TT868 | ATGTGATAGAATTAGTGGGGAATTCATAAAAGAACTTACTTTCTCATAAACCAGTTGCTG | | *bgaA'* to downstream |
| CS121 | GCTTTCTTGAGGCAATTCACTTGGTGC | | |
| **For construction of IU10224 *(*Δ*bgaA*::*tet*-$P_{Zn}$-$RBS^{ftsA}$-*rodZ*-FLAG*)*** | | | |
| TT657 | CGCCCCAAGTTCATCACCAATGACATCAAC | IU9613 | *bgaA'-tet-*$P_{Zn}$-*$RBS^{ftsA}$-rodZ-F* |
| TT863 | TATTTATCATCATCATCTTTATAATCATTTTTAGTAAAGGTTACAGTGATTTGTCCAGTC | | |
| TT864 | AATGATTATAAAGATGATGATGATAAATAAAAGAACTTACTTTCTCATAAACCAGTTGCT | | rodZ-F-downstream |
| CS121 | GCTTTCTTGAGGCAATTCACTTGGTGC | | |
| **For construction of IU11828 (*rodZ*-$HA^{3}$-$P_c$-*kan*)** | | | |
| P1384 | GAGGTAAGCGAGAAGTTTCTGAAGCGGATTGC | D39 | *rodZ* |
| TT928 | AAGCATAATCTGGAACATCATATGGATAATTTTTAGTAAAGGTTACAGTGATTTGTCCAG | | |

| | | | |
|---|---|---|---|
| TT929 | GACAAATCACTGTAACCTTTACTAAAAATTATCCATAT GATGTTCCAGATTATGCTTATC | IU7426 | HA$^{3}$-P$_{c}$-*kan* |
| TT404 | GTTCTTTTTTCATTCGTTTTTCCCTAAAACAATTCATCC AGTAAAATATAATATTTTATT | | |
| TT405 | AATATTATATTTTACTGGATGAATTGTTTTAGGGAAAAA CGAATGAAAAAAGAACAAATT | D39 | Downstrea m of *rodZ* |
| P1385 | ACAACACCTGCAATGGCCACACGTTGCTTT | | |
| **For construction of IU12345 (*ΔmreC* markerless)** | | | |
| P104 | AATGAGACGTGTTGCCATTGCAGG | D39 | upstream to 69bp *mreC* 5’ |
| TT983 | CATCTACATTATGAGTATCTGCACTCAAGAGAGCTGA CACAAGCAGAACAGTGA | | |
| TT984 | TGTTCTGCTTGTGTCAGCTCTCTTGAGTGCAGATACTC ATAATGTAGATGTGATAG | | 51bp *mreC* 3’ downstream |
| P107 | TGTCGCTTTCTCAGCAGCAAGACT | | |
| **For construction of IU12678 or IU12681 (Δ*bgaA*::*tet*-P$_{Zn}$-RBS$^{ftsA}$-*cozE*$^{+}$*)* = P$_{Zn}$-*cozE*$^{+}$** | | | |
| TT657 | CGCCCCAAGTTCATCACCAATGACATCAAC | IU8122 | *bgaA*’-*tet*-P$_{Zn}$-*RBS*$^{ftsA}$- |
| TT968 | CAAAAAAATAATTTATTTCTACGAAACATTACATCGCTT CCTCTCTATCTTCCTTGTTAT | | |
| TT969 | AAGGAAGATAGAGAGGAAGCGATGTAATGTTTCGTAG AAATAAATTATTTTTTTGGACCA | D39 | *cozE* |
| TT970 | CTGGTTTATGAGAAAGTAAGTTCTTTTACTTAGCTAAT TCTCTTTCTCGTTCTTTCATTA | | |
| TT971 | AAGAACGAGAAAGAGAATTAGCTAAGTAAAAGAACTT ACTTTCTCATAAACCAGTTGCTG | D39 | *bgaA”* to downstream |
| C121 | GCTTTCTTGAGGCAATTCACTTGGTGC | | |
| **For construction of IU12696 (*rodZ*Δ(4-68)aa markerless = (ΔHTH)** | | | |
| TT329 | CAACTGATATAGTTGGAAGTGAGGAGTCCATTTCCC | D39 | upstream to *rodZ* Δ(4-68)aa |
| TT999 | CAGAATCATAAGCATCCAAAACAATTTTTCTCATACTT GTCATCCCTTCTTTCTAG | | |
| TT1000 | AGAAGGGATGACAAGTATGAGAAAAATTGTTTTGGAT GCTTATGATTCTGGG | | 3’ *rodZ* to downstream |
| TT977 | CCATACCGATTTGACGACGTATATTCCAAACA | | |
| **For construction of IU12699 (*rodZ*Δ(196-261)aa markerless = (ΔDUF)** | | | |
| TT329 | CAACTGATATAGTTGGAAGTGAGGAGTCCATTTCCC | D39 | upstream to *rodZ*Δ *(*196-261)aa |
| ML1 | AAAGGTTACAGTGATTTGTCCAGTCTGTTGCAATTTAA CTGTTTCCTTACTTGTCTTATA | | |
| ML2 | GACAAGTAAGGAAACAGTTAAATTGCAACAGACTGGA CAAATCACTGTAACCTTTACTAA | | 3’ *rodZ* to downstream |
| P1385 | ACAACACCTGCAATGGCCACACGTTGCTTT | | |
| **For construction of IU12712 and IU12719 (Δ*bgaA*::*kan*-t1t2-P$_{ftsA}$-RBS$^{ftsA}$-*ftsA*)** | | | |
| P146 | TGGCCATTCATCGCTGGTCGTGCTGAAAT | IU9621 | 5’ *bgaA*’-Kan-T1T2 |
| SC484 | GAGCAAAAAAGAAAGCTCTGTGGTAGAAAC GCAAAAAGGCCATCCGTCAGG | | |
| SC483 | GACGGATGGCCTTTTTGCGTTTCTACCACA GAGCTTTCTTTTTTGCTCTTAGAGAG | D39 | P$_{ftsA}$-*ftsA*$^{+}$ |
| AJP49 | CAACTGGTTTATGAGAAAGTAAGTTCTTTTA TTCGTCAAACATGCTTCCGATC | | |

| | | | |
|---|---|---|---|
| AJP50 | CGGAAGCATGTTTGACGAATAAAAGAACTT ACTTTCTCATAAACCAGTTGC | D39 | 3' flanking fragment |
| CS121 | GCTTTCTTGAGGCAATTCACTTGGTGC | | |
| **For construction of IU12738 (*rodZ*(Δ21-257)aa markerless = Δ*rodZ*)** | | | |
| TT329 | CAACTGATATAGTTGGAAGTGAGGAGTCCATTTCCC | D39 | upstream to 60bp *rodZ* 5' |
| TT992 | TGAGCTGTTAATTTCGATAAATCAACACTCAATCCCTG ATTGATTCTAGCTAATCG | | |
| TT993 | GCTAGAATCAATCAGGGATTGAGTGTTGATTTATCGAA ATTAACAGCTCAGACTG | | 60bp *rodZ* 3' to downstream |
| P1385 | ACAACACCTGCAATGGCCACACGTTGCTTT | | |
| **For construction of IU12792 (*rodZ*(1-72)aa markerless)** | | | |
| TT329 | CAACTGATATAGTTGGAAGTGAGGAGTCCATTTCCC | D39 | upstream to *rodZ*Δ (73-273)aa |
| ML3 | ATTTGTTCTTTTTTCATTCGTTTTTCCTTAATCCAAAAC AATTTGGTCATCTAACTCAAC | | |
| ML4 | GTTGAGTTAGATGACCAAATTGTTTTGGATTAAGGAAA AACGAATGAAAAAAGAACAAAT | | 3' *rodZ* to downstream |
| P1385 | ACAACACCTGCAATGGCCACACGTTGCTTT | | |
| **For construction of IU12794 (*rodZ*(1-262)aa markerless)** | | | |
| TT329 | CAACTGATATAGTTGGAAGTGAGGAGTCCATTTCCC | D39 | upstream to *rodZ*Δ (262-273)aa |
| ML5 | TTTGTTCTTTTTTCATTCGTTTTTCCTTAAGCTGTTAAT TTCGATAAATCAACAGTCTGA | | |
| ML6 | CAGACTGTTGATTTATCGAAATTAACAGCTTAAGGAAA AACGAATGAAAAAAGAACAAAT | | 3' *rodZ* to downstream |
| P1385 | ACAACACCTGCAATGGCCACACGTTGCTTT | | |
| **For construction of IU12797 (*rodZ*(1-195)aa markerless)** | | | |
| TT329 | CAACTGATATAGTTGGAAGTGAGGAGTCCATTTCCC | D39 | upstream to *rodZ* Δ (196-273)aa |
| ML7 | TGTTCTTTTTTCATTCGTTTTTCCTTATTGCAATTTAAC TGTTTCCTTACTTGTCTTATA | | |
| ML8 | AGACAAGTAAGGAAACAGTTAAATTGCAATAAGGAAA AACGAATGAAAAAAGAACAAATT | | 3' *rodZ* to downstream |
| P1385 | ACAACACCTGCAATGGCCACACGTTGCTTT | | |
| **For construction of IU12799 (*rodZ*(1-135)aa::TAA-TAG-TGA markerless)** | | | |
| TT329 | CAACTGATATAGTTGGAAGTGAGGAGTCCATTTCCC | D39 | upstream- 5' *rodZ*-135aa |
| ML9 | AGGCTCCTCTGGTTGTCACTATTAAGTTTGAATATAGT TCCAAACATAATAAGTCACAAA | | |
| ML10 | GTTTGGAACTATATTCAAACTTAATAGTGACAACCAGA GGAGCCTTCTCTTTCTAATTAC | | TAA-TAG-TGA-3'rodZ downstream |
| P1385 | ACAACACCTGCAATGGCCACACGTTGCTTT | | |
| **For construction of IU12800 (rodZ(1-103)aa markerless)** | | | |
| TT329 | CAACTGATATAGTTGGAAGTGAGGAGTCCATTTCCC | D39 | upstream to *rodZ*- 103 aa |
| ML11 | TTCTTTTTTCATTCGTTTTTCCTTACTTCTTTTTCTTACT TGAACGTCTACGACCTGTCA | | |
| ML12 | GTCGTAGACGTTCAAGTAAGAAAAAGAAGTAAGGAAA AACGAATGAAAAAAGAACAAATT | | 3' *rodZ* to downstream |
| P1385 | ACAACACCTGCAATGGCCACACGTTGCTTT | | |
| **For construction of IU12803 (*rodZ*(1-134)aa markerless)** | | | |
| TT329 | CAACTGATATAGTTGGAAGTGAGGAGTCCATTTCCC | D39 | |

| | | | |
|---|---|---|---|
| ML13 | TTCTTTTTTCATTCGTTTTTCCTTATTGAATATAGTTCC AAACATAATAAGTCACAAAAA | | upstream to *rodZ-134aa* |
| ML14 | TGTGACTTATTATGTTTGGAACTATATTCAATAAGGAA AAACGAATGAAAAAAGAACAAA | | 3’ *rodZ* to downstream |
| P1385 | ACAACACCTGCAATGGCCACACGTTGCTTT | | |
| **For construction of IU12971 (∆*cozE*::$P_c$-*cat*)** | | | |
| TT962 | CCACCACGGTAAGCAGGCATACCTTCTAAC | D39 | Upstream + 5’ 90 bp *cozE* |
| TT974 | ACATTATCCATTAAAAATCAAACGGATCCTA CAAAGATCCCATCTGTCTCCATAGGTAAA | | |
| Kan rpsL forward | TAGGATCCGTTTGATTTTTAATGGATAATG | IU11119 | $P_c$-*cat* |
| Kan rpsL reverse | GGGCCCCTTTCCTTATGCTTTTG | | |
| TT975 | GTCCAAAAGCATAAGGAAAGGGGCCCTCCC GTTTGTATGAAAATCATAAAATAATGAAAG | D39 | 3’ 60 bp *cozE* + downstream |
| TT963 | GCCGCTAGACAAGGCTTAATCGTATCTCGC | | |
| **For construction of IU13454 *rodZ*(ΔHTH)-FLAG-$P_c$-*erm*** | | | |
| TT329 | CAACTGATATAGTTGGAAGTGAGGAGTCCATTTCCC | IU12696 | upstream to ΔHTH-FLAG |
| ML17 | GTTATTTATCATCATCATCTTTATAATCATTTTTAGTAA AGGTTACAGTGATTTGTCCAG | | |
| ML18 | ACAAATCACTGTAACCTTTACTAAAAATGATTATAAAG ATGATGATGATAAATAACCGGG | IU6293 | $P_c$-*erm* to downstream |
| P1385 | ACAACACCTGCAATGGCCACACGTTGCTTT | | |
| **For construction of IU13456 (*rodZ*(ΔDUF)-FLAG-$P_c$-*erm*)** | | | |
| TT329 | CAACTGATATAGTTGGAAGTGAGGAGTCCATTTCCC | IU12699 | upstream to ΔDUF-FLAG |
| ML17 | GTTATTTATCATCATCATCTTTATAATCATTTTTAGTAA AGGTTACAGTGATTTGTCCAG | | |
| ML18 | ACAAATCACTGTAACCTTTACTAAAAATGATTATAAAG ATGATGATGATAAATAACCGGG | IU6293 | $P_c$-*erm* to downstream |
| P1385 | ACAACACCTGCAATGGCCACACGTTGCTTT | | |
| **For construction of IU13457 (*rodZ*-FLAG markerless)** | | | |
| TT329 | CAACTGATATAGTTGGAAGTGAGGAGTCCATTTCCC | IU6293 | upstream to *rodZ*-FLAG |
| ML15 | CTTTTTTCATTCGTTTTTCCTTATTTATCATCATCATCTT TATAATCATTTTTAGTAAAG | | |
| ML16 | AAATGATTATAAAGATGATGATGATAAATAAGGAAAAA CGAATGAAAAAAGAACAAATTC | D39 | downstream |
| P1385 | ACAACACCTGCAATGGCCACACGTTGCTTT | | |
| **For construction of IU13555 (*rodZ*(1-72aa)-FLAG-$P_c$-*erm*)** | | | |
| TT329 | CAACTGATATAGTTGGAAGTGAGGAGTCCATTTCCC | D39 | upstream *rodZ*Δ (73-273)aa-FLAG |
| ML22 | CGGTTATTTATCATCATCATCTTTATAATCATCCAAAAC AATTTGGTCATCTAACTCAAC | | |
| ML23 | TGAGTTAGATGACCAAATTGTTTTGGATGATTATAAAG ATGATGATGATAAATAACCGGG | IU6293 | $P_c$-*erm* downstream |
| P1385 | ACAACACCTGCAATGGCCACACGTTGCTTT | | |

| | | | |
|---|---|---|---|
| **For construction of IU13556 (*rodZ*(1-134)aa-FLAG-P$_c$*-erm)*** | | | |
| TT329 | CAACTGATATAGTTGGAAGTGAGGAGTCCATTTCCC | D39 | upstream *rodZ* (1-134aa)-FLAG |
| ML20 | ATTTATCATCATCATCTTTATAATCTTGAATATAGTTCC AAACATAATAAGTCACAAAAA | | |
| ML21 | GACTTATTATGTTTGGAACTATATTCAAGATTATAAAGA TGATGATGATAAATAACCGGG | IU6293 | P$_c$-*erm* downstream |
| P1385 | ACAACACCTGCAATGGCCACACGTTGCTTT | | |
| **For construction of IU13577 (Δ*rodZ*-FLAG-P$_c$*-erm)*** | | | |
| TT329 | CAACTGATATAGTTGGAAGTGAGGAGTCCATTTCCC | IU12738 | upstream to Δ*rodZ*-FLAG |
| ML17 | GTTATTTATCATCATCATCTTTATAATCATTTTTAGTAA AGGTTACAGTGATTTGTCCAG | | |
| ML18 | ACAAATCACTGTAACCTTTACTAAAAATGATTATAAAG ATGATGATGATAAATAACCGGG | IU6293 | P$_c$-*erm* downstream |
| P1385 | ACAACACCTGCAATGGCCACACGTTGCTTT | | |
| **For construction of IU13655 (*rodZ*(ΔDUF)-FLAG markerless)** | | | |
| TT329 | CAACTGATATAGTTGGAAGTGAGGAGTCCATTTCCC | IU13456 | upstream to ΔDUF-FLAG |
| ML15 | CTTTTTTCATTCGTTTTTCCTTATTTATCATCATCATCTT TATAATCATTTTTAGTAAAG | | |
| ML16 | AAATGATTATAAAGATGATGATGATAAATAAGGAAAAA CGAATGAAAAAAGAACAAATTC | IUI3457 | downstream |
| P1385 | ACAACACCTGCAATGGCCACACGTTGCTTT | | |
| **For construction of IU13656 (*rodZ(*Δ21-257)-FLAG-markerless)** | | | |
| TT329 | CAACTGATATAGTTGGAAGTGAGGAGTCCATTTCCC | IU13577 | upstream to Δ*rodZ*-FLAG |
| ML15 | CTTTTTTCATTCGTTTTTCCTTATTTATCATCATCATCTT TATAATCATTTTTAGTAAAG | | |
| ML16 | AAATGATTATAAAGATGATGATGATAAATAAGGAAAAA CGAATGAAAAAAGAACAAATTC | IUI3457 | downstream |
| P1385 | ACAACACCTGCAATGGCCACACGTTGCTTT | | |
| **For construction of IU13658 (*rodZ*(1-72)aa-FLAG markerless)** | | | |
| TT329 | CAACTGATATAGTTGGAAGTGAGGAGTCCATTTCCC | IU13555 | upstream to *rodZ* (1-72aa)-FLAG |
| ML26 | TTCCTTATTTATCATCATCATCTTTATAATCATCCAAAA CAATTTGGTCATCTAACTCAA | | |
| ML27 | TTAGATGACCAAATTGTTTTGGATGATTATAAAGATGA TGATGATAAATAAGGAAAAACG | IUI3457 | downstream |
| P1385 | ACAACACCTGCAATGGCCACACGTTGCTTT | | |
| **For construction of IU13660 (*rodZ*(1-134)-FLAG markerless)** | | | |
| TT329 | CAACTGATATAGTTGGAAGTGAGGAGTCCATTTCCC | IU13556 | upstream to rodZ-135aa-FLAG |
| ML24 | TATTTATCATCATCATCTTTATAATCTTGAATATAGTTC CAAACATAATAAGTCACAAAA | | |
| ML25 | TATTATGTTTGGAACTATATTCAAGATTATAAAGATGAT GATGATAAATAAGGAAAAACG | IUI3457 | downstream |
| P1385 | ACAACACCTGCAATGGCCACACGTTGCTTT | | |
| **For construction of IU13680 (Δ*pbp1b*::P$_c$*-aad9*)** | | | |

| | | | |
|---|---|---|---|
| P222 | CGTTCGTGTGGCGCTGCTTCAAATTGTT | D39 | upstream to + 100 bp of *pbp1b* |
| P456 | CATTATCCATTAAAAATCAAACGGATCCTATTGAACCT TTCTTGCCAGGTCTAGCTGATT | | |
| kan rpsL forward | TAGGATCCGTTTGATTTTTAATGGATAATG | IU6987 | $P_c$-*aad9* |
| kan rpsL reverse | GGGCCCCTTTCCTTATGCTTTTG | | |
| P225 | CAAAAGCATAAGGAAAGGGGCCCTCTAGCGATAGCA GTAACTCAAGTACTACACGACCTT | D39 | 3' 57 bp *pbp1b* + downstream |
| P522 | AACGGCAACCACCAAAGGAGAAACCAAGGA | | |
| **For construction of IU13705 (*rodZ* (ΔHTH)-FLAG markerless)** | | | |
| TT329 | CAACTGATATAGTTGGAAGTGAGGAGTCCATTTCCC | IU13454 | upstream to ΔHTH-FLAG |
| ML15 | CTTTTTTCATTCGTTTTTCCTTATTTATCATCATCATCTT TATAATCATTTTTAGTAAAG | | |
| ML16 | AAATGATTATAAAGATGATGATGATAAATAAGGAAAAA CGAATGAAAAAAGAACAAATTC | IUI3457 | downstream |
| P1385 | ACAACACCTGCAATGGCCACACGTTGCTTT | | |
| **For construction of IU13837 (ΔbgaA::kan-T1T2-$P_{Zn}$-$RBS_{ftsA}$-*pgsA*$^{+}$) = $P_{Zn}$-*pgsA*** | | | |
| P146 | TGGCCATTCATCGCTGGTCGTGCTGAAAT | IU12788 | Δ*bgaA*-Pc-*kan*-t1t2-$RBS^{ftsA}$-$P_{Zn}$ |
| ML39 | AGATTGGGAATTTGTTCTTTTTTCATTACATCGCTTCCT CTCTATCTTCCTTGTTATAAT | | |
| ML38 | ATAACAAGGAAGATAGAGAGGAAGCGATGTAATGAAA AAAGAACAAATTCCCAATCTCTT | D39 | *pgsA* |
| ML41 | AGCAACTGGTTTATGAGAAAGTAAGTTCTTTCATTTCG AACCAAATGTCCCTTTAAATAC | | |
| ML40 | TTTAAAGGGACATTTGGTTCGAAATGAAAGAACTTACT TTCTCATAAACCAGTTGCTGCG | | *bgaA"* to downstream |
| CS121 | GCTTTCTTGAGGCAATTCACTTGGTGC | | |
| **For construction of IU13960 (Δ*pgsA*::$P_c$-*erm*)** | | | |
| P347 | GCAGACGATTTCGATCAACTTCCAAGTCC | D39 | upstream to 5' 60bp *pgsA'* |
| P349 | CATTATCCATTAAAAATCAAACGGATCCTAAATAGGTA TAAAGAGAATTCGACCTATTGT | | |
| kan rpsL forward | TAGGATCCGTTTGATTTTTAATGGATAATG | $P_c$-*erm* cassette | $P_c$-*erm* |
| kan rpsL reverse | GGGCCCCTTTCCTTATGCTTTTG | | |
| P350 | CAAAAGCATAAGGAAAGGGGCCCGGCTATGACTATTT CAAGGGTAGTGCC | D39 | 60bp 3' *pgsA*' downstream |
| P351 | TCACATTTTCTAGAGCAATTCCCATAGCTTATCC | | |
| **For construction of IU14522 and IU14524 (*rodZ*$^{+}$-$P_c$-[*kan-rpsL*$^{+}$]-60bp 3'-*rodZ*$^{+}$)** | | | |
| TT997 | TTACAGGAAATTACTTTAGAGGATGTCCTTGATGCTGG | D39 | upstream plus *rodZ* |
| ML47 | TTATCCATTAAAAATCAAACGGATCCTATTAATTTTTAG TAAAGGTTACAGTGATTTGTC | | |

| | | | |
|---|---|---|---|
| kan rpsL forward | TAGGATCCGTTTGATTTTTAATGGATAATG | $P_c$-[*kan-rpsL*$^+$] cassette | $P_c$-[*kan-rpsL*$^+$] |
| kan rpsL reverse | GGGCCCCTTTCCTTATGCTTTTG | | |
| ML48 | TAAACGTCCAAAAGCATAAGGAAAGGGGCCCGATTTATCGAAATTAACAGCTCAGACTGG | D39 | 60bp 3'-*rodZ* (repeat) downstream |
| P1385 | ACAACACCTGCAATGGCCACACGTTGCTTT | | |
| **For construction of IU14528 (∆*cozE*::$P_c$-[*kan-rpsL*$^+$])** | | | |
| TT962 | CCACCACGGTAAGCAGGCATACCTTCTAAC | D39 | Upstream + 90 bp of 5' *cozE* |
| TT974 | ACATTATCCATTAAAAATCAAACGGATCCTACAAAGATCCCATCTGTCTCCATAGGTAAA | | |
| kan rpsL reverse | TAGGATCCGTTTGATTTTTAATGGATAATG | $P_c$-[*kan-rpsL*$^+$] cassette | $P_c$-[*kan-rpsL*$^+$] |
| kan rpsL forward | GGGCCCCTTTCCTTATGCTTTTG | | |
| TT975 | GTCCAAAAGCATAAGGAAAGGGGCCCTCCCGTTTGTATGAAAATCATAAAATAATGAAAG | D39 | 3' 60 bp *cozE* + downstream |
| TT963 | GCCGCTAGACAAGGCTTAATCGTATCTCGC | | |
| **For construction of IU14697 (∆*pbp1b*)** | | | |
| P222 | CGTTCGTGTGGCGCTGCTTCAAATTGTT | D39 | Upstream + 5' 99 bp of *pbp1b* |
| TT1115 | TAGTACTTGAGTTACTGCTATCGCTAGATGAACCTTTCTTGCCAGGTCTAGC | | |
| TT1116 | AGACCTGGCAAGAAAGGTTCATCTAGCGATAGCAGTAACTCAAGTACTACACG | D39 | 3' 60 bp of *pbp1b* + downstream |
| P522 | AACGGCAACCACCAAAGGAGAAACCAAGGA | | |
| **For construction of IU15337 (*pbp2b*(Q56L)-HA-$P_c$-*kan*)** | | | |
| TT452 | GGAGGGTTGGCTGTGGGTGGCTACAAGAAC | D39 | 5' of *pbp2b* (Q56L) |
| TT1167 | TGAACTGCTTGTAATCTTGGTCAGACTAGCTGAGGCTAG | | |
| TT1168 | CTAGCCTCAGCTAGTCTGACCAAGATTACAAGCAGTTCA | IU6933 | 3' *pbp2b*-HA-$P_c$-*kan* |
| TT352 | TGAAGGACTGGAAAGACCACTGCACCTTCT | | |
| **For construction of IU15340 (*pbp2b*(T57A)-HA-$P_c$-*kan*** | | | |
| TT452 | GGAGGGTTGGCTGTGGGTGGCTACAAGAAC | D39 | 5' of *pbp2b* (T57A) |
| TT1169 | GAACTGCTTGTAATCTTGGCCTGACTAGCTGAGGCTAG | | |
| TT1170 | CTAGCCTCAGCTAGTCAGGCCAAGATTACAAGCAGTTC | IU6933 | 3' *pbp2b*-HA-$P_c$-*kan* |
| TT352 | TGAAGGACTGGAAAGACCACTGCACCTTCT | | |
| **For construction of IU15341 (*pbp2b*(T57N)-HA-$P_c$-*kan*** | | | |
| TT452 | GGAGGGTTGGCTGTGGGTGGCTACAAGAAC | D39 | 5' of *pbp2b* |

| | | | |
|---|---|---|---|
| TT1171 | CTGAACTGCTTGTAATCTTGTTCTGACTAGCTGAGGCTAG | | (T57N) |
| TT1172 | CTAGCCTCAGCTAGTCAGAACAAGATTACAAGCAGTTCAG | IU6933 | 3' *pbp2b*-HA-$P_c$-*kan* |
| TT352 | TGAAGGACTGGAAAGACCACTGCACCTTCT | | |
| **For construction of IU15343 (*pbp2b*(T57R)-HA-$P_c$-*kan*** | | | |
| TT452 | GGAGGGTTGGCTGTGGGTGGCTACAAGAAC | D39 | 5' of *pbp2b* (T57R) |
| TT1175 | ACTGCTTGTAATCTTGCGCTGACTAGCTGAGGCTAG | | |
| TT1176 | CTAGCCTCAGCTAGTCAGCGCAAGATTACAAGCAGT | IU6933 | 3' *pbp2b*-HA-$P_c$-*kan* |
| TT352 | TGAAGGACTGGAAAGACCACTGCACCTTCT | | |
| **For construction of IU15347 (*pbp2b*(I290A)-HA-$P_c$-*kan*** | | | |
| TT452 | GGAGGGTTGGCTGTGGGTGGCTACAAGAAC | D39 | 5' of *pbp2b* (I290A) |
| TT1177 | CATATTTATCCAGATGGGCTTCTTTTACCGAGCGTTT | | |
| TT1178 | AAACGCTCGGTAAAAGAAGCCCATCTGGATAAATATG | IU6933 | 3' *pbp2b*-HA-$P_c$-*kan* |
| TT352 | TGAAGGACTGGAAAGACCACTGCACCTTCT | | |
| **For construction of IU15628 *rodZ*(Y51A F55A Y59A)-Flag-markerless** | | | |
| TT329 | CAACTGATATAGTTGGAAGTGAGGAGTCCATTTCCC | D39 | upstream to *rodZ* (Y51A F55A Y59A) |
| ML56 | ATGCAGCTTTTTTCAAAGCAGAACGCGTAGCAAAAGGACTTGGAAGTTGATCGAAATCGT | | |
| ML57 | TGCTACGCGTTCTGCTTTGAAAAAAGCTGCATGGGCTGTTGAGTTAGATGACCAAATTGT | IU14594 | 3' *rodZ*-Flag to downstream |
| P1385 | ACAACACCTGCAATGGCCACACGTTGCTTT | | |
| **For construction of IU15907 ($P_c$-[*kan-rpsL*$^+$]-*rodA*$^+$)** | | | |
| P1543 | CAGGCCGTACTCTTCTGTCCTCTTTACTTCC | D39 | upstream *rodA* |
| ML84 | CATTATCCATTAAAAATCAAACGGATCCTATATTTATCAAAGTTCATTAAAAAATCTATC | | |
| Kan rpsL forward | TAGGATCCGTTTGATTTTTAATGGATAATG | $P_c$-[*kan-rpsL*$^+$] cassette | $P_c$-[*kan-rpsL*$^+$] |
| Kan rpsL reverse | GGGCCCCTTTCCTTATGCTTTTG | | |
| ML85 | AAACGTCCAAAAGCATAAGGAAAGGGGCCCGTATTGTATGAAAGTATAAGGTTAGTACAT | D39 | 5' of *rodA* |
| ML86 | AATAACCAGGACAGAGCCAATAAAGCC | | |
| **For construction of IU15928 (*iht*-L*6*-*pbp2b* markerless)** | | | |
| TT452 | GGAGGGTTGGCTGTGGGTGGCTACAAGAAC | D39 | upstream of *pbp2b* |
| ML82 | AGCCAAAAAAATTTCCAAACCTTTTTTATCCATTTCTAACTTAAAATCTTACTCTTAATT | | |
| AJP405 | GATAAAAAAGGTTTGGAAATTTTTTTGGCTTCTGCTGAAATTGGTACTGGTTTTCCATTT | IU14738 | *iht*-$L_6$ |
| YT104 | ACCAGAACCTTGACCAGATCCTGGTCCTTG | | |
| ML83 | CAAGGACCAGGATCTGGTCAAGGTTCTGGTAGACTGATTTGTATGAGAAAATTTAACAGC | D39 | 5' of *pbp2b* |

| | | | |
|---|---|---|---|
| TT352 | TGAAGGACTGGAAAGACCACTGCACCTTCT | | |
| **For construction of IU15970 (*iht*-L*6-rodA* markerless)** | | | |
| P1543 | CAGGCCGTACTCTTCTGTCCTCTTTACTTCC | D39 | upstream of *rodA* |
| ML87 | CAAAAAAATTTCCAAACCTTTTTATCCATATGTACTAACCTTATACTTTCATACAATAC | | |
| AJP405 | GATAAAAAAGGTTTGGAAATTTTTTTGGCTTCTGCTGAAATTGGTACTGGTTTTCCATTT | IU14738 | *iht*-$L_6$ |
| YT104 | ACCAGAACCTTGACCAGATCCTGGTCCTTG | | |
| ML88 | CAAGGACCAGGATCTGGTCAAGGTTCTGGTAAACGTTCTCTCGACTCTAGAGTCGATTAT | D39 | 5' of *rodA* |
| ML86 | AATAACCAGGACAGAGCCAATAAAGCC | | |
| **For construction of IU16344 (*iht*-$L_6$-*mreC* markerless)** | | | |
| P104 | AATGAGACGTGTTGCCATTGCAGG | D39 | Upstream of *mreC* |
| TT1232 | AAAAATTTCCAAACCTTTTTATCCATATCCCTACCTTTATATCAAAAACTGTTACAGTA | | |
| TT1233 | TGTAACAGTTTTTGATATAAAGGTAGGGATATGGATAAAAAAGGTTTGGAAATTTTTTTG | IU14738 | *iht*-$L_6$ |
| TT1234 | GACATATTTTGATTTTTTAAAACGGTTACCAGAACCTTGACCAGATCCTGGTCCTTGTCC | | |
| TT1235 | ACCAGGATCTGGTCAAGGTTCTGGTAACCGTTTTAAAAAATCAAAATATGTCATTATTGT | D39 | *mreC* to downstream |
| TT1236 | CCAAGCCTATAACAAAACAATAGACTAGGTAGAGATACTCTG | | |
| **For transformation assays** | | | |
| P222 | CGTTCGTGTGGCGCTGCTTCAAATTGTT | E193 | Δ*pbp1b* ::$P_c$-*erm* |
| P522 | AACGGCAACCACCAAAGGAGAAACCAAGGA | | |
| TT329 | CAACTGATATAGTTGGAAGTGAGGAGTCCATTTCCC | E655 | Δ*rodZ* ::$P_c$-*erm* |
| P1385 | ACAACACCTGCAATGGCCACACGTTGCTTT | | |
| P222 | CGTTCGTGTGGCGCTGCTTCAAATTGTT | K180 | Δ*pbp1b*::$P_c$-[*kan-rpsL*$^+$] |
| P522 | AACGGCAACCACCAAAGGAGAAACCAAGGA | | |
| P104 | AATGAGACGTGTTGCCATTGCAGG | IU1751 | Δ*mreCD* <>*aad9* |
| P107 | TGTCGCTTTCTCAGCAGCAAGACT | | |
| TT329 | CAACTGATATAGTTGGAAGTGAGGAGTCCATTTCCC | IU6987 | Δ*rodZ* ::$P_c$-*aad9* |
| P1385 | ACAACACCTGCAATGGCCACACGTTGCTTT | | |
| TT452 | GGAGGGTTGGCTGTGGGTGGCTACAAGAAC | IU7397 | Δ*pbp2b* <>aad9 |
| TT352 | TGAAGGACTGGAAAGACCACTGCACCTTCT | | |
| TT457 | ATTGTGGATGGTTTCCAAGGGATTCGTG AC | IU7814 | Δ*ftsZ*::aad9 |
| TT166 | TCATTGGGAGAGCCGGTTCCTGTGAAGAAT | | |
| P1348 | TCTTCTTGCAGCCTTGAAAGAGGTGGCAGT | IU9102 | Δ*mpgA* ::$P_c$-*aad9* |
| P1349 | AGAGCAAACTAGGAAACTAGCCGCAGGTTG | | |
| TT329 | CAACTGATATAGTTGGAAGTGAGGAGTCCATTTCCC | IU9931 | Δ*rodZ* |

| | | | |
|---|---|---|---|
| P1385 | ACAACACCTGCAATGGCCACACGTTGCTTT | | <>*aad9* |
| P174 | ATGTGGTGTATCCGCATTGGGACAGGAT | IU10294 | Δ*spd_1874* ::$P_c$-*cat* |
| P175 | AGCCGTAAGTCGCAGCACCAATCACAAA | | |
| P1543 | CAGGCCGTACTCTTCTGTCCTCTTTACTTCC | IU10943 | Δ*rodA* ::$P_c$-*erm* |
| P1544 | CGGGTGTTCAAGCTCTCTGGCTTCATTTTC | | |
| P104 | AATGAGACGTGTTGCCATTGCAGG | IU12268 | Δ*mreC* ::$P_c$-*erm* |
| P107 | TGTCGCTTTCTCAGCAGCAAGACT | | |
| TT962 | CCACCACGGTAAGCAGGCATACCTTCTAAC | IU12332 | Δ*cozE* ::$P_c$-*erm* |
| TT963 | GCCGCTAGACAAGGCTTAATCGTATCTCGC | | |
| TT329 | CAACTGATATAGTTGGAAGTGAGGAGTCCATTTCCC | IU12515 | Δ*rodZ*::$P_c$-[*kan-rpsL*$^+$] |
| P1385 | ACAACACCTGCAATGGCCACACGTTGCTTT | | |
| TT329 | CAACTGATATAGTTGGAAGTGAGGAGTCCATTTCCC | IU12738 | *rodZ* Δ(21-257)aa markerless |
| P1385 | ACAACACCTGCAATGGCCACACGTTGCTTT | | |
| TT962 | CCACCACGGTAAGCAGGCATACCTTCTAAC | IU12971 | Δ*cozE* ::$P_c$-*cat* |
| TT963 | GCCGCTAGACAAGGCTTAATCGTATCTCGC | | |
| P222 | CGTTCGTGTGGCGCTGCTTCAAATTGTT | IU13680 | Δ*pbp1b* ::$P_c$-*aad9* |
| P522 | AACGGCAACCACCAAAGGAGAAACCAAGGA | | |
| P347 | GCAGACGATTTCGATCAACTTCCAAGTCC | IU13960 | Δ*pgsA* ::$P_c$-*erm* |
| P351 | TCACATTTTCTAGAGCAATTCCCATAGCTTATCC | | |
| P146 | TGGCCATTCATCGCTGGTCGTGCTGAAAT | E46 | Δ*bgaA* ::$P_c$-*erm* |
| P147 | TACGCCTTCTATCATGCCTTTGATCGCCCGT | | |

| **For construction of *S. pneumoniae* B2H plasmids** | | |
|---|---|---|
| Primer | Sequence (5’-3’) | Template |
| **Construction of T25/T18-fusions to *S. pneumoniae rodZ* ΔHTH** | | |
| pKT25/pUT18C_*rodZ* ΔHTH_BF | CGGGATCCCATGAGAAAAATTGTTTTGGATGCTTA | IU12696 |
| pKT25/pUT18C_*rodZ*_ER | CGGAATTCTTAATTTTTAGTAAAGGTTACAGTGA | |
| **Construction of T25/T18-fusions to *S. pneumoniae rodZ* ΔDUF** | | |
| pKT25/pUT18C_*rodZ*_BF | CGGGATCCTATGAGAAAAAAAACAATTGGAGAGG | IU12699 |
| pKT25/pUT18C_*rodZ*_ER | CGGAATTCTTAATTTTTAGTAAAGGTTACAGTGA | |
| **Construction of T25/T18-fusions to *S. pneumoniae pbp1b*** | | |
| pKT25/pUT18C_*pbp1b*_XF | GCTCTAGAGATGCAAAATCAATTAAATGAATTAAA ACGAAAAATGCT | D39 |
| pKT25/pUT18C_*pbp1b*_BR | CGGGATCCTTATCGTCTCGCCCTTGAAGAAGAAG GTCGT | |
| **For verification and sequencing of *S. pneumoniae* B2H fusions** | | |
| pKT25_579F | GTTCGCCATTATGCCGCATC | |
| pKT25_802R | GGATGTGCTGCAAGGCGATT | |
| pUT18C_484F | GATGTACTGGAAACGGTGC | |
| pUT18C_660R | CTTAACTATGCGGCATCAGAGC | |
| pKNT25/pUT18_49F | CGCAATTAATGTGAGTTAGC | |
| pKNT25_328R | TTGATGCCATCGAGTACG | |
| pUT18_304R | CGAGCGATTTTCCACAACAA | |

| | |
|---|---|
| *pbp1b_656F1* | TAACGACCTATCTCAATGTG |
| *pbp1b_1245F2* | TGGAACAGGTCGTGTAGAAG |
| *pbp1b_1264R1* | CTTCTACACGACCTGTTCCA |

[a]Genomic DNA of D39 was used as templates for PCR reactions, except for $P_c$-[*kan*-*rpsL*$^+$] and $P_c$-*erm* cassettes (Tsui *et al.*, 2011).

**Table S5**. Overexpression of PgsA does not alleviate *ΔrodZ* lethality

| **Amplicon** | Number of colonies 20-24 h after transformation[a] | |
|---|---|---|
| | **- Zn** | **+Zn** |
| Recipient strain: IU1824 WT | | |
| 1. No DNA (- control ) | 0 | ND (not done) |
| 2. Δ*pbp1b*::$P_c$-*aad9* (+ control ) | 100-200 | ND |
| 3. Δ*rodZ<>aad9* | 0 | ND |
| Recipient strain: IU9613 *rodZ*$^+$// $P_{Zn}$-*rodZ*$^+$ | | |
| 4. No DNA (- control ) | 0 | 0 |
| 5. Δ*pbp1b*::$P_c$-*aad9* (+ control ) | 200-300 | 200-300 |
| 6. Δ*rodZ<>aad9* | 0 | 200-300 |
| Recipient strain: IU13837 *pgsA*$^+$//$P_{Zn}$-*pgsA*$^+$ | | |
| 7. No DNA (- control ) | 0 | 0 |
| 8. Δ*pbp1b*::$P_c$-*aad9* (+ control ) | 200-300 | 200-300 |
| 9. Δ*rodZ<>aad9* | 0 | 0 |
| Recipient strain: IU13960 Δ*pgsA*::$P_c$-*erm* //$P_{Zn}$-*pgsA*$^+$ | | |
| 10. No DNA (- control ) | 0 | 0 |
| 11. *Δpbp1b*::$P_c$-*aad9* (+ control ) | 0 | 200-300 |
| 12. Δ*rodZ<>aad9* | 0 | 0 |

[a]Recipient strains were constructed as described in Table S1. Transformations with 30 ng of the indicated amplicons were performed as described in *Experimental Procedures.* Zn inducer (0.4 mM $ZnCl_2$ + 0.04 mM $MnSO_4$) was added to media as indicated to increase expression of RodZ or PgsA in merodiploid strains. IU1824 and IU13837 or IU 9613 and IU13960 were initially grown in BHI lacking or containing Zn inducer, respectively. IU9613, IU13837, and IU13960 cells were collected by centrifugation and resuspended in transformation mix lacking or containing Zn inducer, which were subsequently plated in soft agar on TSAII-BA plates lacking or containing Zn inducer. The number of colonies obtained for 300 μL of transformation mix are shown. Similar results were obtained from three independent experiments.

**Table S6**. Mutations in the membrane proximal region of *S. pneumoniae* bPBP2b do not suppress Δ*rodZ* lethality

| | | | # of colonies 20 h after transformation of amplicons[a] | |
|---|---|---|---|---|
| **Recipient strain[b]** | **Zn (mM)** | **genotype[c]** | **Δ*pbp1b*::$P_c$-*aad9*** | **Δ*rodZ*::$P_c$-*aad9*** |
| IU9765 | 0 | Δ*bgaA*::*tet*-$P_{Zn}$-*rodZ*$^+$ | >500 | <5 faint |
| | 0.4 | | >500 | >500 |
| IU13440 | 0 | *pbp2b*(WT)-HA-$P_c$*kan*//$P_{Zn}$-*pbp2b* | >500 | <5 faint |
| IU15337 | 0 | *pbp2b*(Q56L)-HA-$P_c$*kan*//$P_{Zn}$-*pbp2b* | >500 | <5 faint |
| IU15340 | 0 | *pbp2b*(T57A)-HA-$P_c$*kan*//$P_{Zn}$-*pbp2b* | >500 | <5 faint |
| IU15341 | 0 | *pbp2b*(T57N)-HA-$P_c$*kan*//$P_{Zn}$-*pbp2b* | >500 | <5 faint |
| IU15343 | 0 | *pbp2b*(T57R)-HA-$P_c$*kan*//$P_{Zn}$-*pbp2b* | >500 | <5 faint |
| IU15347 | 0 | *pbp2b*(I290A)-HA-$P_c$*kan*//$P_{Zn}$-*pbp2b* | >500 | <5 faint |

[a]Transformations were performed as described in *Experimental Procedures.* For IU9765, Zn inducer (0.4 mM $ZnCl_2$ + 0.04 mM $MnSO_4$) was added to the transformation mix, which was then divided into plating soft agar and TSAII-BA plates containing or lacking Zn inducer. The other strains were transformed with no Zn addition. The number of colonies is normalized to 1mL of transformation mixture.

[b]All recipient strains are in the D39W Δ*cps* background, and all are Zn-independent for growth.

[c]Amino acids Q56 and T57 are in a similar position in a 3D model of *Spn* bPBP2b as the activating amino acid change in L61R in *Eco* bPBP2 (Rohs *et al.*, 2018). I290A is predicted to form a salt bridge with T57 in the 3D model of *Spn* bPBP2b.

**Table S7**. Qualitative scoring of RodZ WT, ΔHTH, and ΔDUF interactions by B2H assays

| T18[b] | T25-RodZ[a] | | | | T25[b] | T18-RodZ[a] | | |
|---|---|---|---|---|---|---|---|---|
| | WT | ΔHTH | ΔDUF | | | WT | ΔHTH | ΔDUF |
| RodZ WT[c] | ++++ | +++ | ++++ | | RodZ WT[c] | ++++ | ++ | ++++ |
| GpsB | +++ | + | +++ | | GpsB | +++ | + | ++ |
| MreC | +++ | ++ | +++ | | MreC | ++++ | +++ | +++ |
| MreD | +++ | +++ | +++ | | MreD | +/- | +/- | - |
| MpgA | +++ | +++ | +++ | | MpgA | ++ | ++ | ++ |
| bPBP2b | ++ | + | + | | bPBP2b | ++ | + | + |
| RodA | +++ | ++ | +++ | | RodA | +++ | + | +++ |
| aPBP1b | ++ | + | ++ | | aPBP1b | ++ | +/- | + |
| aPBP1a | ++++ | ++++ | ++++ | | aPBP1a | ++++ | ++ | ++ |
| aPBP2a | +++ | +++ | +++ | | aPBP2a | ++++ | ++ | ++ |
| bPBP2x | ++ | + | ++ | | bPBP2x | +++ | + | ++ |
| FtsW | ++ | + | - | | FtsW | ++ | + | - |
| EzrA | +++ | +++ | +++ | | EzrA | + | + | + |
| DivIVA | ++ | + | ++ | | DivIVA | +/- | +/- | +/- |
| StkP | + | +/- | + | | StkP | - | - | - |
| FtsA | + | + | + | | FtsA | - | - | - |
| FtsZ | - | - | - | | FtsZ | - | - | - |
| self | ++++ | ++ | ++++ | | self | ++++ | ++ | ++++ |

[a]T25-CyaA or T18-CyaA domain fused to the N-terminus of full-length RodZ$^{1-273}$ (WT), RodZ$^{\Delta 4-68}$(ΔHTH), or RodZ$^{\Delta 196-261}$(ΔDUF).

[b]T18-CyaA or T25-CyaA domain fused to N or C terminus of full-length selected proteins;

[c]Qualitative measure of β-galactosidase production. Co-transformations of strain BTH101 [*cya*-99] carrying appropriate plasmid pairs were spotted directly on LBKA+X-gal indicator plates, inspected for color development after 24, 30, and 36 h and scored similarly as reported in (Bendezu *et al.*, 2009): (-), white at 36 h; (+/-), white at 24 h, but light color afterwards; (+), white at 24 h, but medium color afterwards; (++), light color at 24 h and

medium/dark blue afterwards; (+++), medium blue at 24 h and dark blue afterwards; (++++), dark blue at 24 h and afterwards. No interactions were detected between the T18-CyaA or the T25-CyaA domains alone and the respective RodZ, RodZ$^{\Delta 4\text{-}68}$ (ΔHTH), and RodZ$^{\Delta 196\text{-}261}$ (ΔDUF) fusions (see Fig. S15A); In all B2H assays, T18-CyaA or T25-CyaA domain alone (T18 and T25), and T18-CyaA or T25-CyaA fused to the N terminus of the leucine zipper protein Zip (T18-Zip and T25-Zip) were used as negative (-) and positive (+) controls, respectively.

A

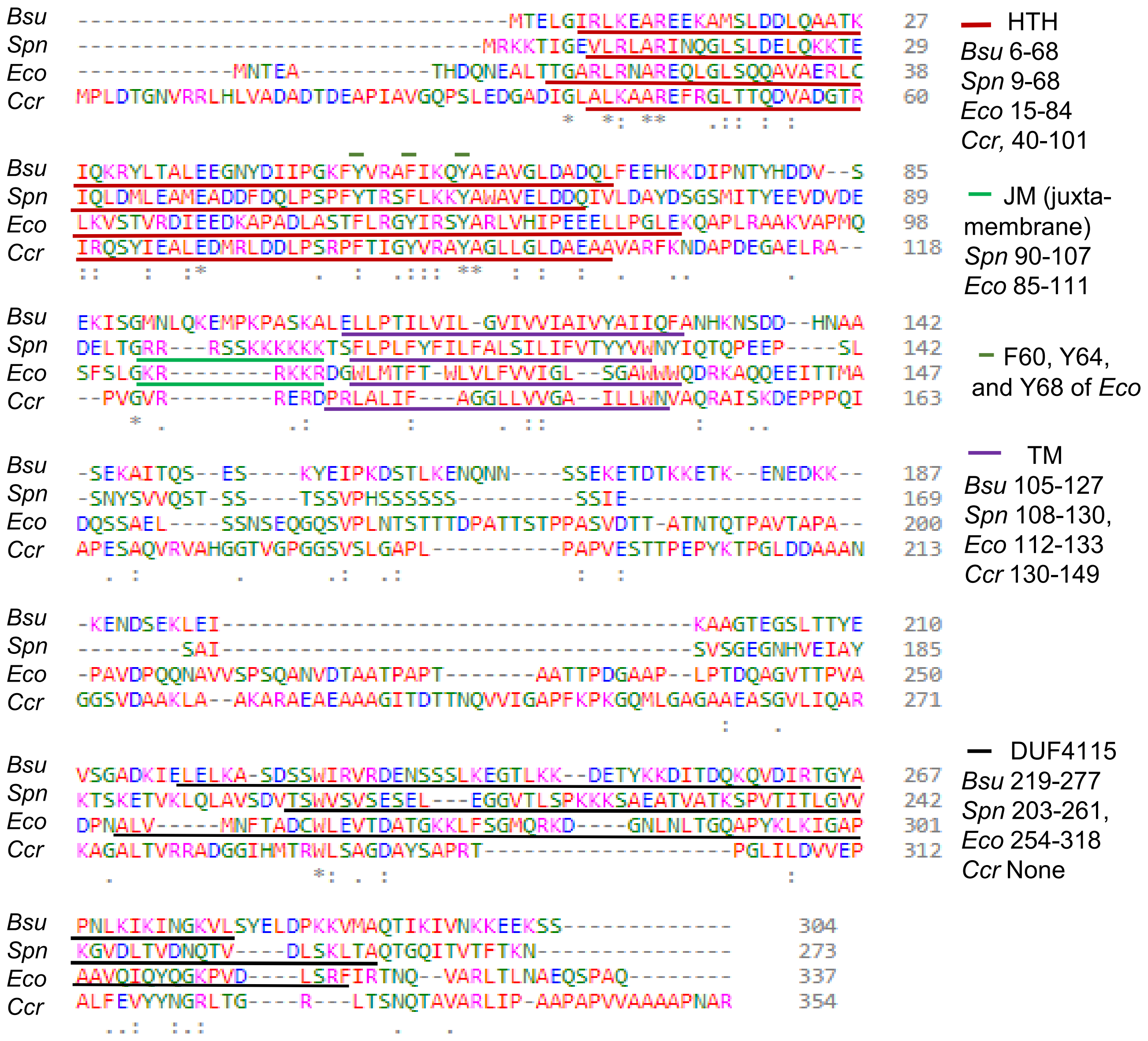


**Fig. S1. Amino acid (aa) alignments and AlphaFold2 structural predictions of RodZ from different bacteria. (A)** Clustal Omega amino acid alignment of RodZ of four bacteria. *B. subtilis* (str.168, QJR46138.1), *S. pneumoniae* (D39 SPD_2050, ABJ54044.1, *E. coli* (K-12, NP-417011.1), and *C. crescentus* (*Caulobacter vibrioides* CB15, ADW96154.1). HTH (helix-turn-helix) domains were identified as HTH_25 by NCBI conserved domain search. JM (juxta-membrane) domain of *E. coli* is described in (Bendezu *et al.*, 2009). F60 and Y64 of *E. coli* (green bars) interact with MreB (van den Ent *et al.*, 2010). TM (transmembrane) domains are determined with TMHMM server. DUFs (domains of unknown function) were identified as DUF4115 by NCBI conserved domain search. (Continued on next page)

B

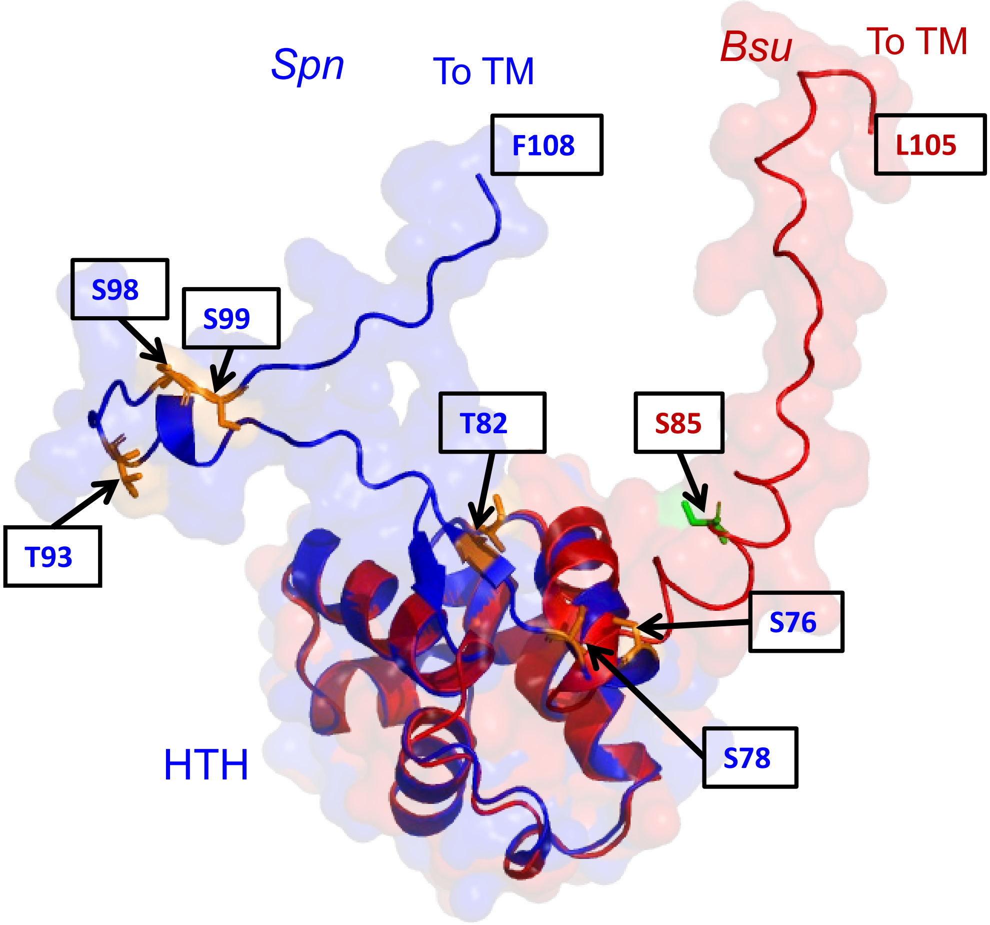


**Fig. S1. (B)** *In silico* structures predicted by AlphaFold2 of the N- termini of RodZ *(Spn)* (1-108 aa, blue) and RodZ (*Bsu*) (1-105 aa, red). The HTH domains are residues 5-76 and 6-69 residues of RodZ(*Spn)* and RodZ(*Bsu*), respectively. The regions between the HTH and the TM (77-107 of RodZ(*Spn*) and 70-104 of RodZ(*Bsu*) are not conserved (see Fig. S1A). S85 of RodZ(*Bsu*) is located in the region between the HTH and the TM domain and is reported to be phosphorylated (Sun and Garner, 2020). S76, S78, T82, T93, T98 and T99 are serine and threonine residues present in the region between HTH and the TM domain of RodZ(*Spn*).

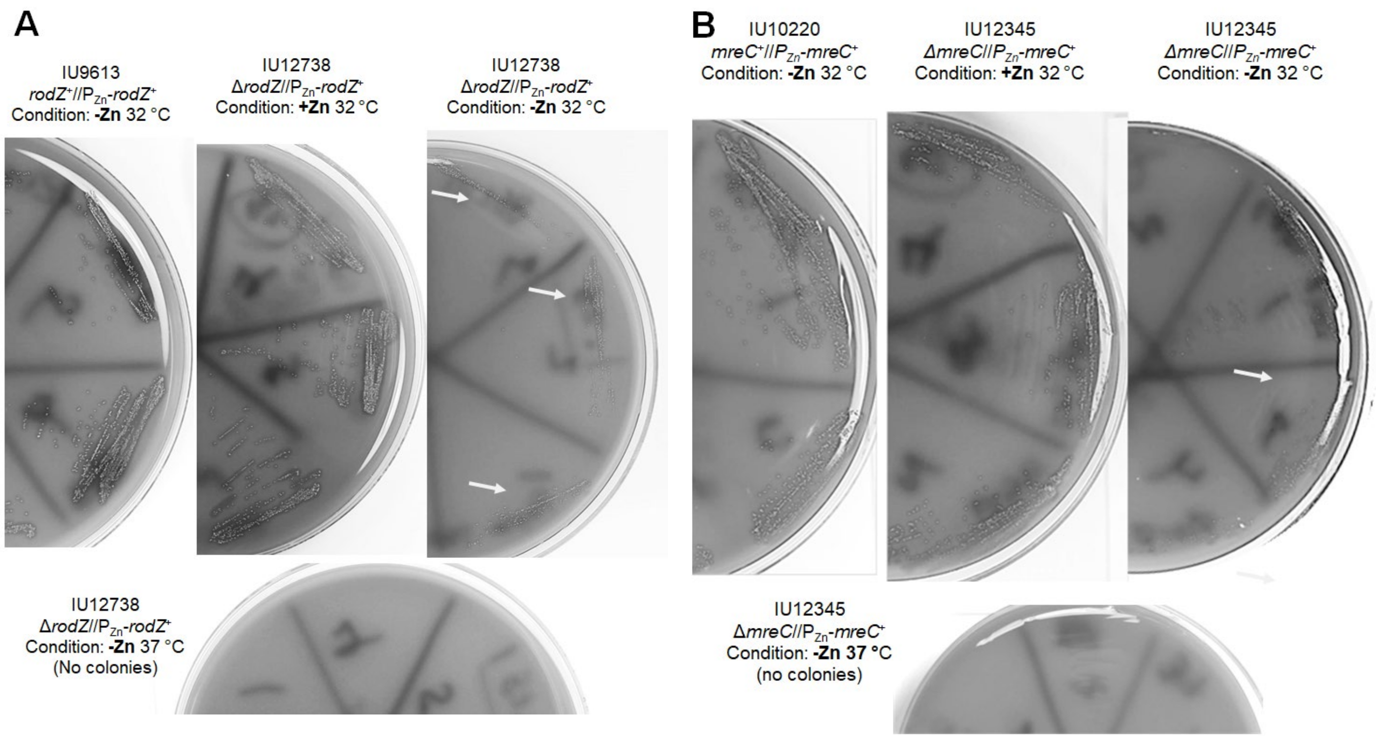


**C**

| Appearance of transformant colonies after 48 h incubation at 32°C or 37°C | | |
|---|---|---|
| Recipient strain IU1945 (WT) | | |
| | 32°C | 37°C |
| Amplicon:<br>1. No DNA (control) | 0 | 0 |
| 2. Δ*ftsZ*::*aad9* (septal) | 0 | 0 |
| 3. Δ*rodZ*<>*aad9* | + (small) | 0 |
| 4. Δ*mreCD*<>*aad9* | + (small) | 0 |
| 5. Δ*pbp2b*<>*aad9* | + (small) | 0 |
| 6. Δ*rodA*::$P_c$-*erm* | + (small) | 0 |

**Fig. S2. Colonies of *rodZ* and *mreC* depletion and mutant strains are detected at 32°C, but not at 37°C.** Merodiploid strains depleted of **(A)** RodZ or **(B)** MreC form tiny colonies at 32°C, but not at 37°C. Strains were streaked from frozen glycerol stocks onto TSAII-BA plates containing or lacking Zn inducer (0.4 mM $ZnCl_2$ + 0.04 mM $MnSO_4$) for 24 h, after which single colonies were re-steaked onto fresh plates (shown above) for 24 h. Tiny colonies of strains depleted for RodZ or MreC are indicated by arrows. **(C)** Transformation of deletion amplicons into WT strain IU1945 was performed at the temperatures indicated as described in *Experimental procedures*. Experiments were repeated several times with similar results.

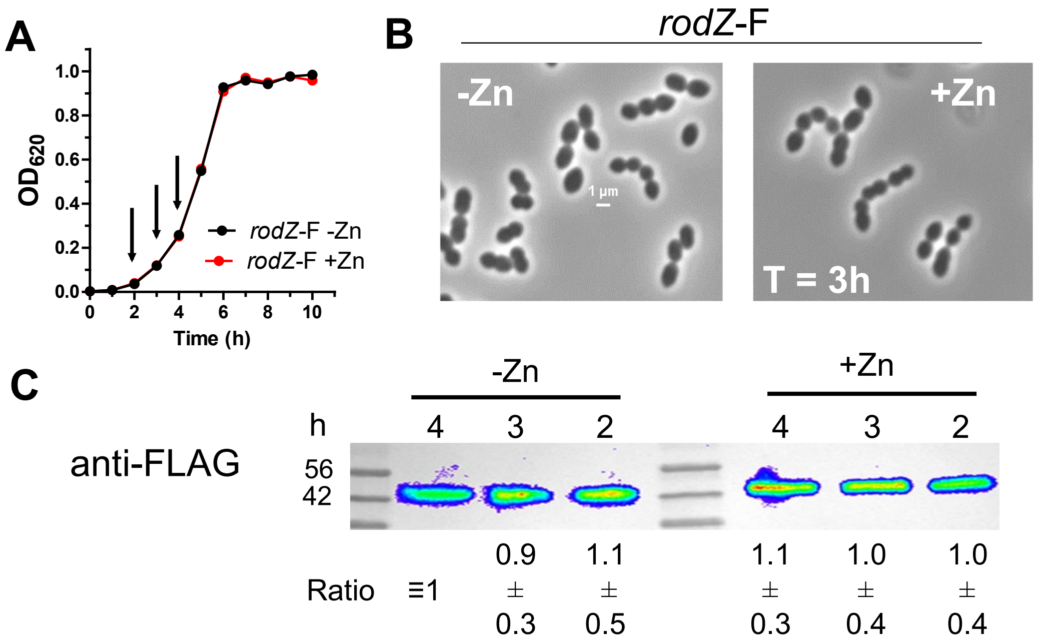


**Fig. S3. Zinc does not affect growth, cell morphology, or RodZ-F levels in the WT background. (A)** Representative growth curves of IU14594 (*rodZ*-F) ± Zn inducer (0.4 mM Zn +0.04 mM Mn) conditions. IU14594 was grown overnight in BHI at 37°C in the presence of 5% $CO_2$ without inducer. For day growth, samples were re-suspended in fresh BHI ± Zn to an $OD_{620}$ of ≈0.003. Arrows indicate time at which samples were harvested for western blot analysis. **(B)** Representative micrographs displaying IU14594 ± Zn at 3 h of growth. **(C)** Western blot showing relative RodZ-Flag amounts in IU14594 in the +Zn or -Zn conditions at 2h, 3h or 4h of growth. Western blotting was carried out with primary anti-Flag antibody and secondary HRP antibody labeling, and visualization with IVIS Living Image system. 3 µg of crude lysate was loaded in each lane. Quantitation of RodZ-F (average ± SEM) was obtained from two independent biological replicates.

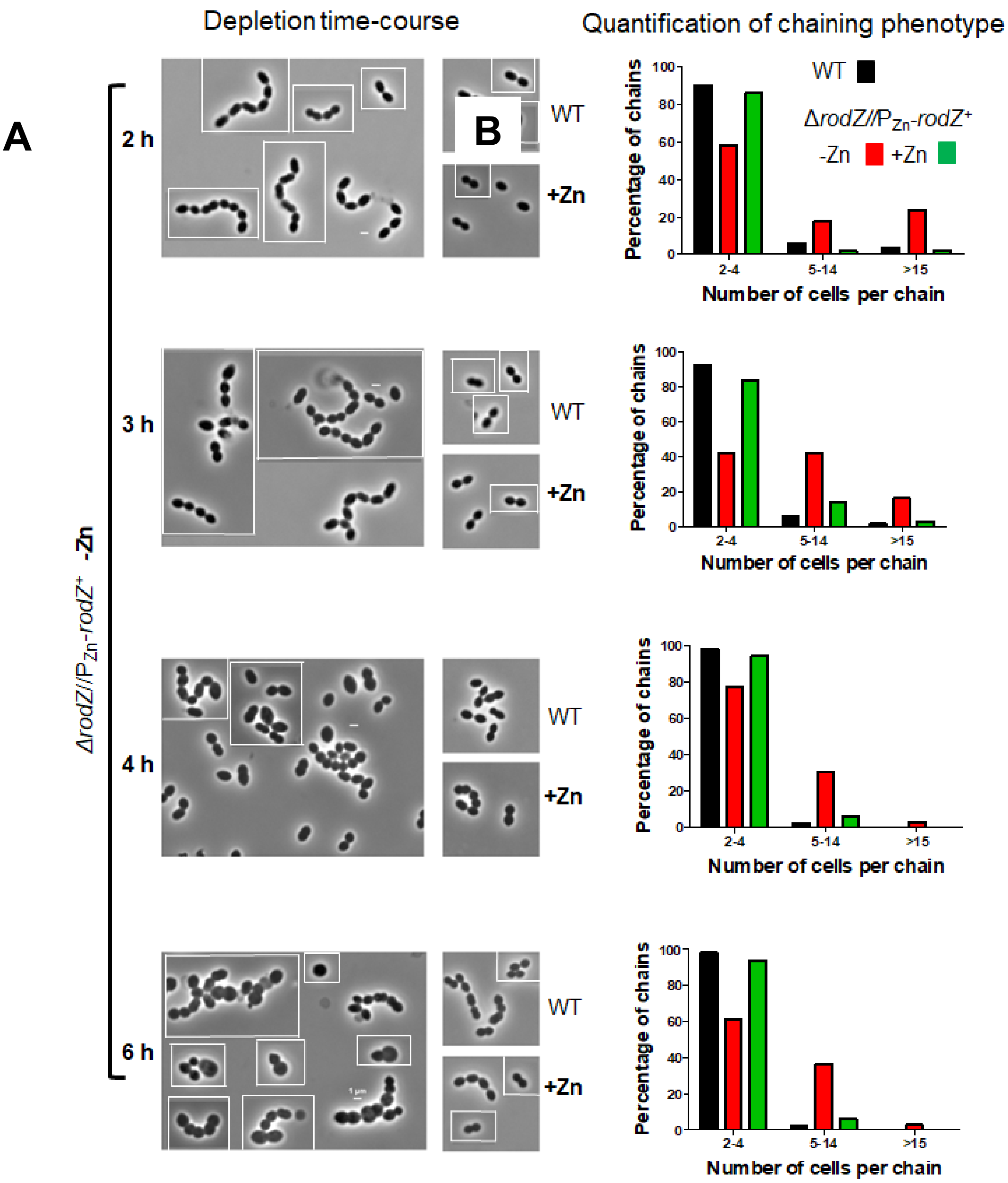


**Fig. S4. Cell morphology at various times after RodZ depletion. (A)** Representative images showing the loss of cell shape maintenance during RodZ depletion. Images were taken at 2, 3, 4 or 6 h after resuspension of IU1824 (WT) or IU12738 (Δ*rodZ*//P$_{Zn}$-*rodZ*$^{+}$) in BHI with or without Zn inducer (0.4 mM $ZnCl_2$ + 0.04 mM $MnSO_4$). Micrographs are mosaic and representative cells are shown. All scale bars represent 1 µm. Micrographs were composed using Illustrator and all images are to scale. Experiments were repeated 3-5 times with similar results. **(B)** Quantification of the chaining phenotype during depletion of RodZ. The number of cells in each chain were counted and categorized at various time-points. 100 chains were considered per sample for each time point. Data presented in the graphs are obtained from two independent experiments.

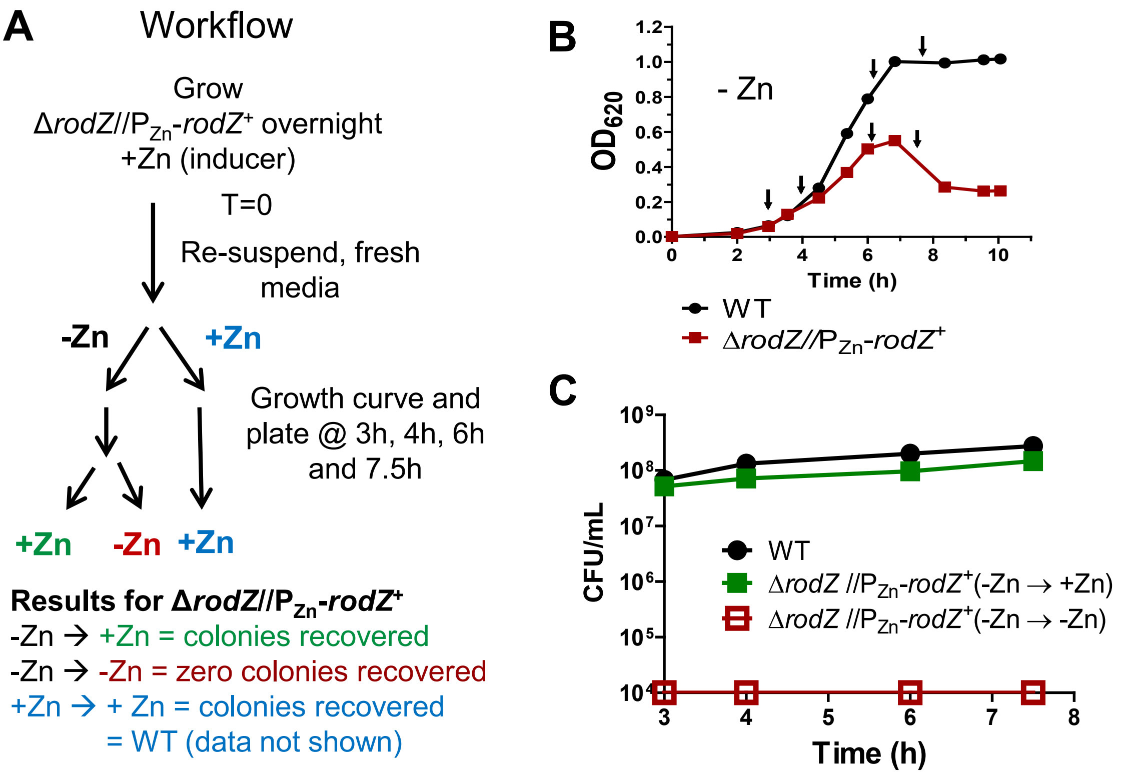


**Fig.S5. Cells depleted of RodZ for 7.5 h remain viable when plated onto TSAII-BA plates + Zn inducer. (A)** Schematic of the workflow of the OD/CFU experiment in which growth curves and CFU assays were performed as described in *Experimental procedures*. **(B)** and **(C)**, Representative growth curves and CFU determination results of WT (IU1824) and Δ*rodZ//*$P_{zn}$-*rodZ*⁺ (IU12738) strains. Strains were grown overnight in the presence of the inducer then re-suspended into fresh media with or without (+/-) Zn inducer (0.4 mM $ZnCl_2$ + 0.04 mM $MnSO_4$). During growth, samples were harvested and CFU assays were conducted for the +/- inducer conditions at 3, 4, 6 and 7.5 h. Plates were incubated at 37°C and counted for CFUs at 20-24 h. Note data points plotted for the -Zn → -Zn condition were illustrated as the level of detection (red symbol and line in (C) as CFUs were unrecoverable for the Δ*rodZ* strain in the -Zn → -Zn condition at a dilution of $10^{-4}$. For simplicity, the +Zn → +Zn conditions for the WT and Δ*rodZ* variant are not graphed. These data represent similar results obtained from 4 independent experiments.

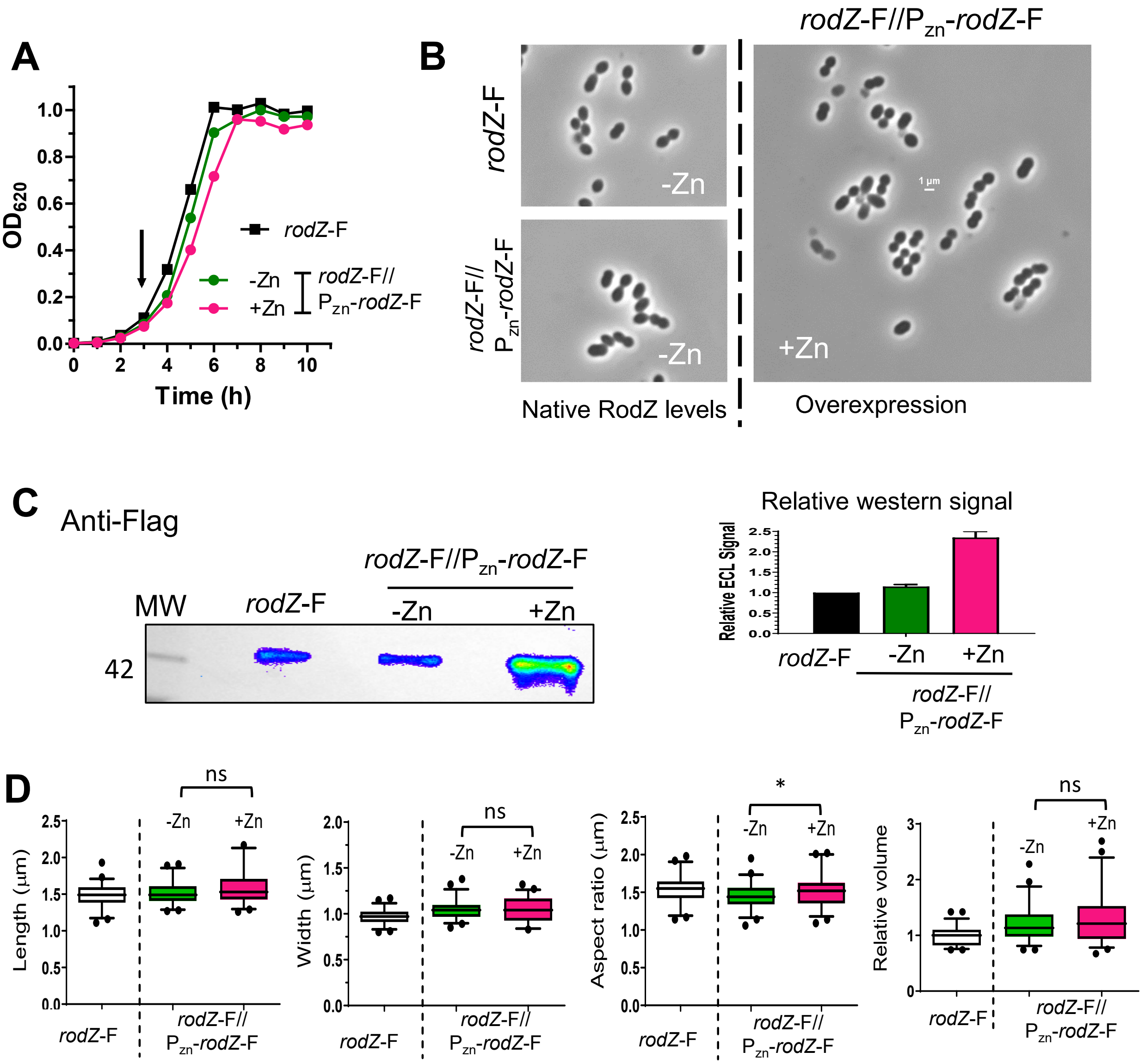


**Figure S6. Overexpression of RodZ does not alter growth or cellular morphology. (A)** and **(B)** Representative growth curves and microscopic images of IU14594 (*rodZ*-F) and IU16338 (*rodZ*-F//$P_{Zn}$-*rodZ*-F) with/without Zn inducer (0.4 mM $ZnCl_2$ + 0.04 mM $MnSO_4$). Arrow indicates time at which samples were harvested for microscopy and western blot analysis. Samples for microscopy and western analyses were taken at an $OD_{620}$ ≈0.15 – 0.2 at 3 h. Scale bar = 1 µm. **(C)** Quantitative western blot probed with anti-Flag as described in *Experimental procedures*. 3 µg of crude cell lysate was loaded on each lane. Right, graph displaying relative western signals. A to C are representative results from one of at least 3 independent biological replicates. **(D)** Box and whiskers plot (5-95 percentile) of cell length, width, aspect ratio, and relative volume measured for IU14594 and IU16338. ≈50 cells per sample were measured. Statistical analysis was conducted using two-tailed t-test between IU16338 under + and – Zn conditions. * $p < 0.05$; ns, non-significant.

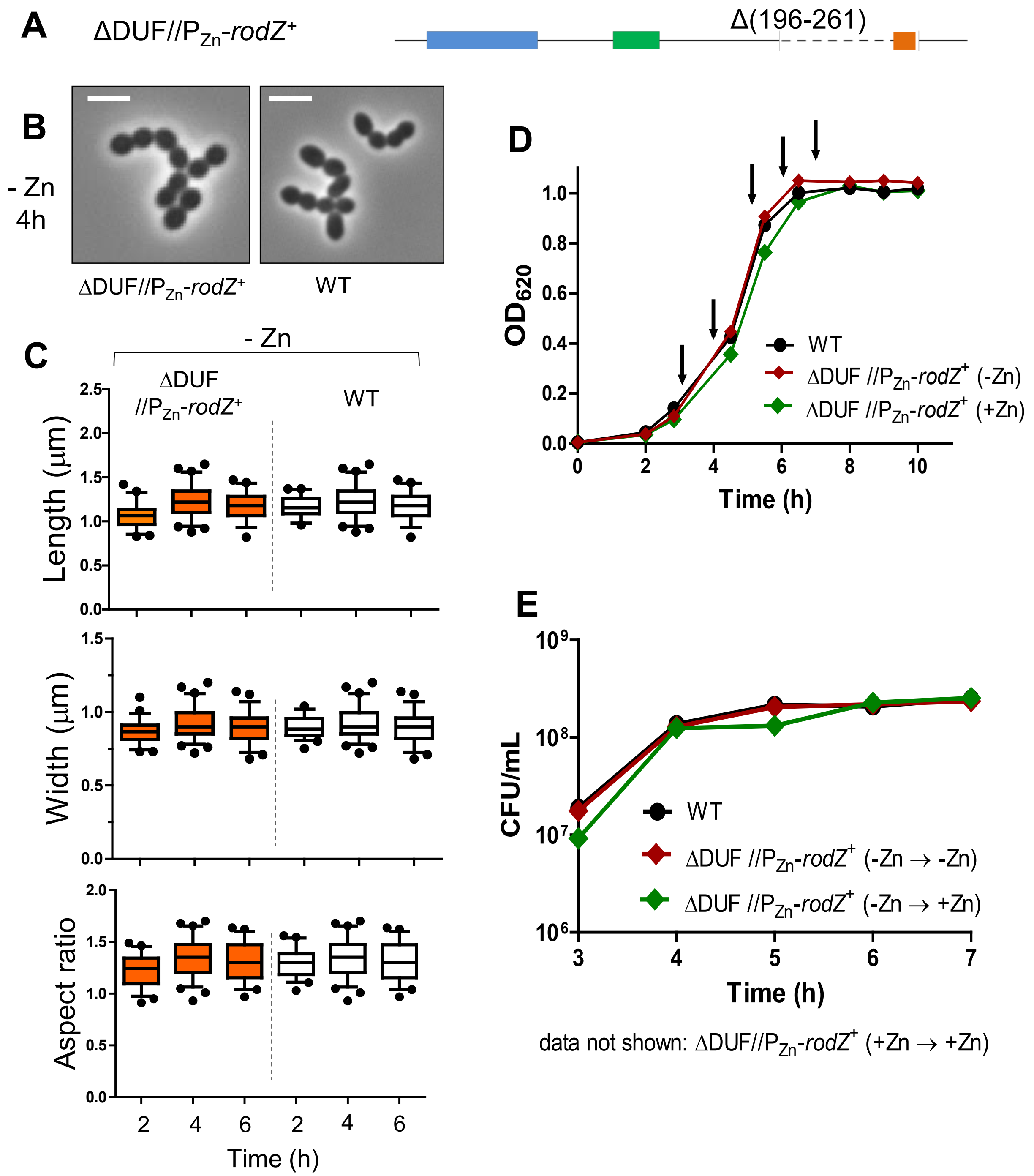


**Fig. S7. Deletion of the DUF domain does not alter cell shape, growth, or viability.** Analysis of the ΔDUF//P$_{Zn}$-*rodZ*$^+$ strain (IU12699) in comparison to WT (IU1824). **(A)** Representation of the DUF deletion at the native locus in the chromosome. **(B)** Representative cells of IU1824 and IU12699 imaged at 4 h. **(C)** Box and whiskers plot (5 to 95 percentile) of cell length, width, and aspect ratio of IU12699 in comparison to WT, both in the -Zn condition at 2, 4 or 6h of growth. For each sample and time point, 50-80 cells were measured. Statistical analysis using one-way ANOVA analysis showed no statistical significance between IU12699 and IU1824 in length, width and aspect ratio when samples from the same time points were compared. **(D)** Representative growth curves and **(E)** viability assay performed with strains IU1824 and IU12699. Arrows indicate time in which samples were harvested and plated to determine colony forming units. All experiments were performed with two or more biological replicates from which similar results were obtained.

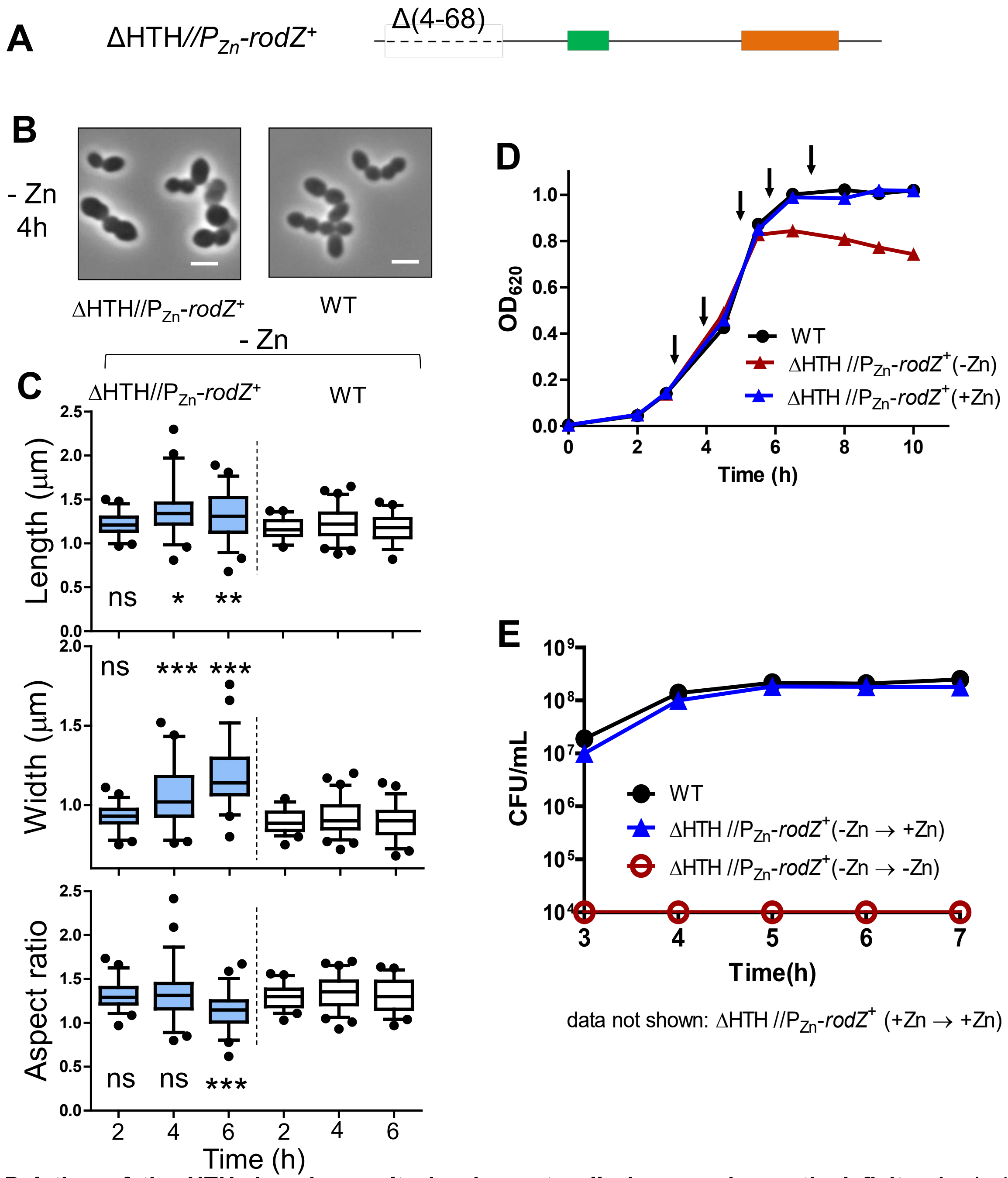


**Fig. S8. Deletion of the HTH domain results in aberrant cell shape and growth deficits**. Analysis of the ΔHTH//P$_{Zn}$-*rodZ*$^{+}$ strain (IU12696) in comparison to WT (IU1824). **(A)** Representation of the deletion at the native locus in the chromosome. **(B)** Representative cells of IU1824 and IU12696 imaged at 4 h. **(C)** Box and whiskers plot (5 to 95 percentile) of cell length, width, and aspect ratio of IU12696 in comparison to WT, both in the -Zn condition at 2, 4 or 6h of growth. For each sample and time point, 50-80 cells were measured. Statistical analysis was conducted using one-way ANOVA analysis by comparing IU12696 and IU1824 from the same time points. * $p < 0.05$; ** $p < 0.01$ *** $p < 0.001$, ns, non-significant. **(D)** Representative growth curves and **(E)** viability assay performed with strains IU1824 and IU12696. Arrows indicate time in which samples were harvested and plated to determine colony forming units. Note data points plotted for IU12696 -Zn condition were illustrated as the level of detection (red symbol and line in E as CFUs were unrecoverable for IU12696 under the -Zn condition at a dilution of $10^{-4}$. All experiments were performed with two or more biological replicates from which similar results were obtained.

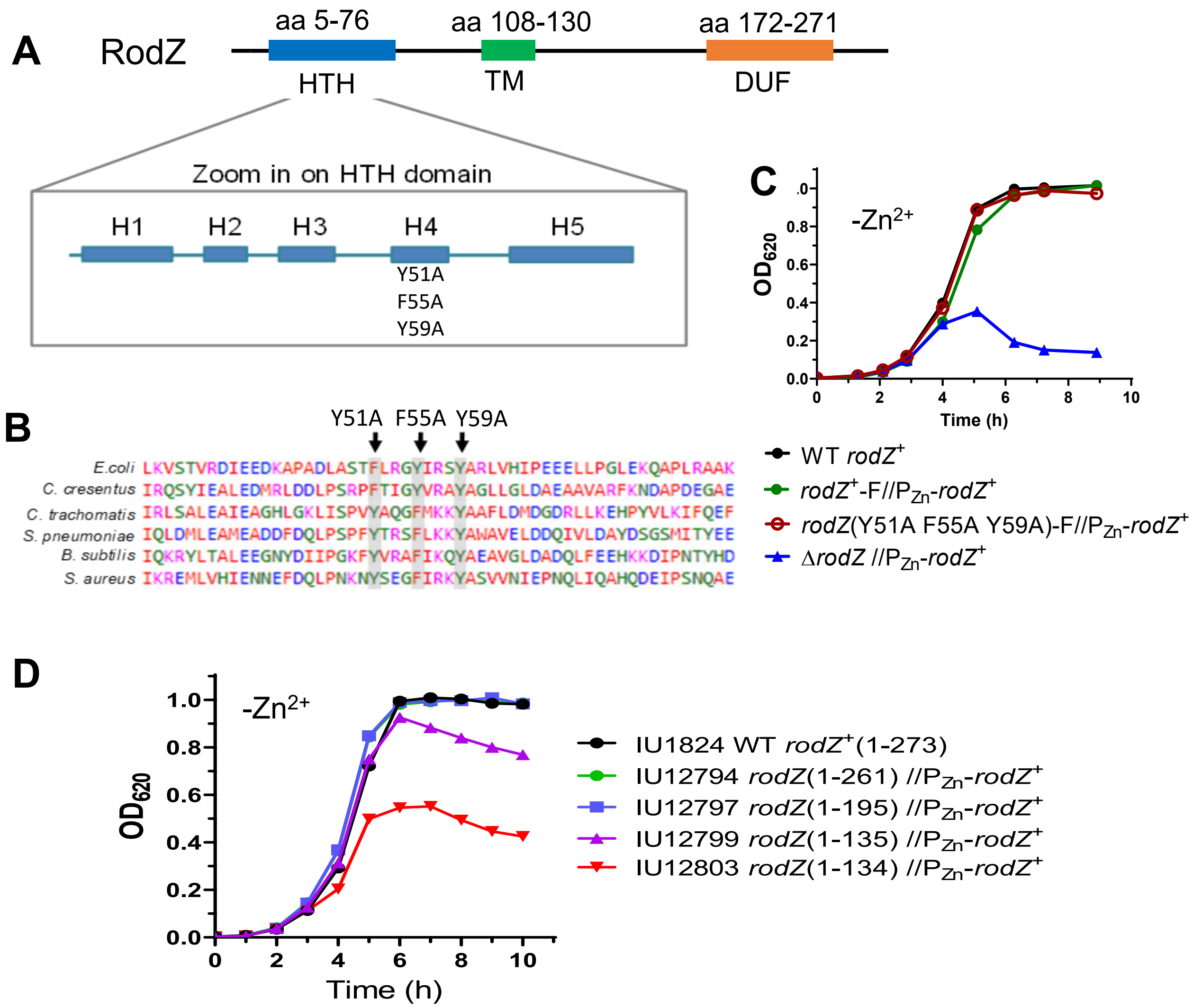


**Fig. S9. Conserved aromatic amino acids in RodZ required to bind MreB in *E.coli* are not essential in *S. pneumoniae*. (A)** Schematic of RodZ of *S. pneumoniae* zooming in on the HTH domain. Constructed point mutations of RodZ in *S. pneumoniae* are indicated. **(B)** Gray boxes and arrows highlight the conserved aromatic residues present in RodZ of different bacteria. Point mutations of the RodZ triple mutant constructed in *S. pneumoniae* are shown. **(C)** Representative growth curve of IU1824 (WT), IU13457 (*rodZ*-F//$P_{zn}$-*rodZ*$^+$), IU15628 (*rodZ*(Y51A F55A Y59A)-F//$P_{zn}$-*rodZ*$^+$) and IU12738 (Δ*rodZ*-F//$P_{zn}$-*rodZ*$^+$). **(D)** Growth curves of RodZ truncation variants capable of growing in the absence of the Zn inducer (0.04 mM $Zn^{2+}$/0.04 mM $Mn^{2+}$). For C and D, strains were grown overnight in BHI broth +Zn inducer, and resuspended in BHI broth –Zn inducer. Similar growth curves were obtained for each strain in two or more independent biological replicates.

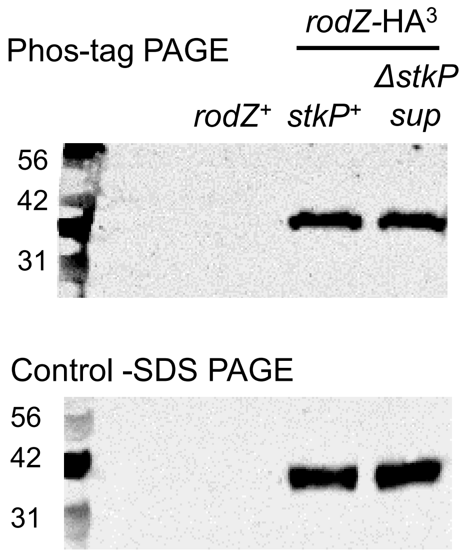


**Fig. S10. Phosphorylation of RodZ by the StkP protein kinase is not detected in *S. pneumoniae*.** Top panel, phos-tag PAGE western blot of a non-HA-tagged control strain IU15987 (*rodZ*$^{+}$ Δ*stkP sup* (with suppressor mutation), IU11828 (*stkP*$^{+}$ *rodZ*-HA$^{3}$) and IU17883 (Δ*stkP sup rodZ*-HA$^{3}$). RodZ-HA$^{3}$ migrates at the same location in the *stkP*$^{+}$ and Δ*stkP* mutant samples, indicating lack of StkP-dependent phosphorylation of RodZ. Bottom panel, control western of SDS PAGE showing identical RodZ-HA$^{3}$ amounts in the *stkP*$^{+}$ and Δ*stkP* strains. Phos-tag PAGE and western blotting procedures using anti-HA as the primary antibody are described in *Experimental procedures*. The experiment was performed twice independently.

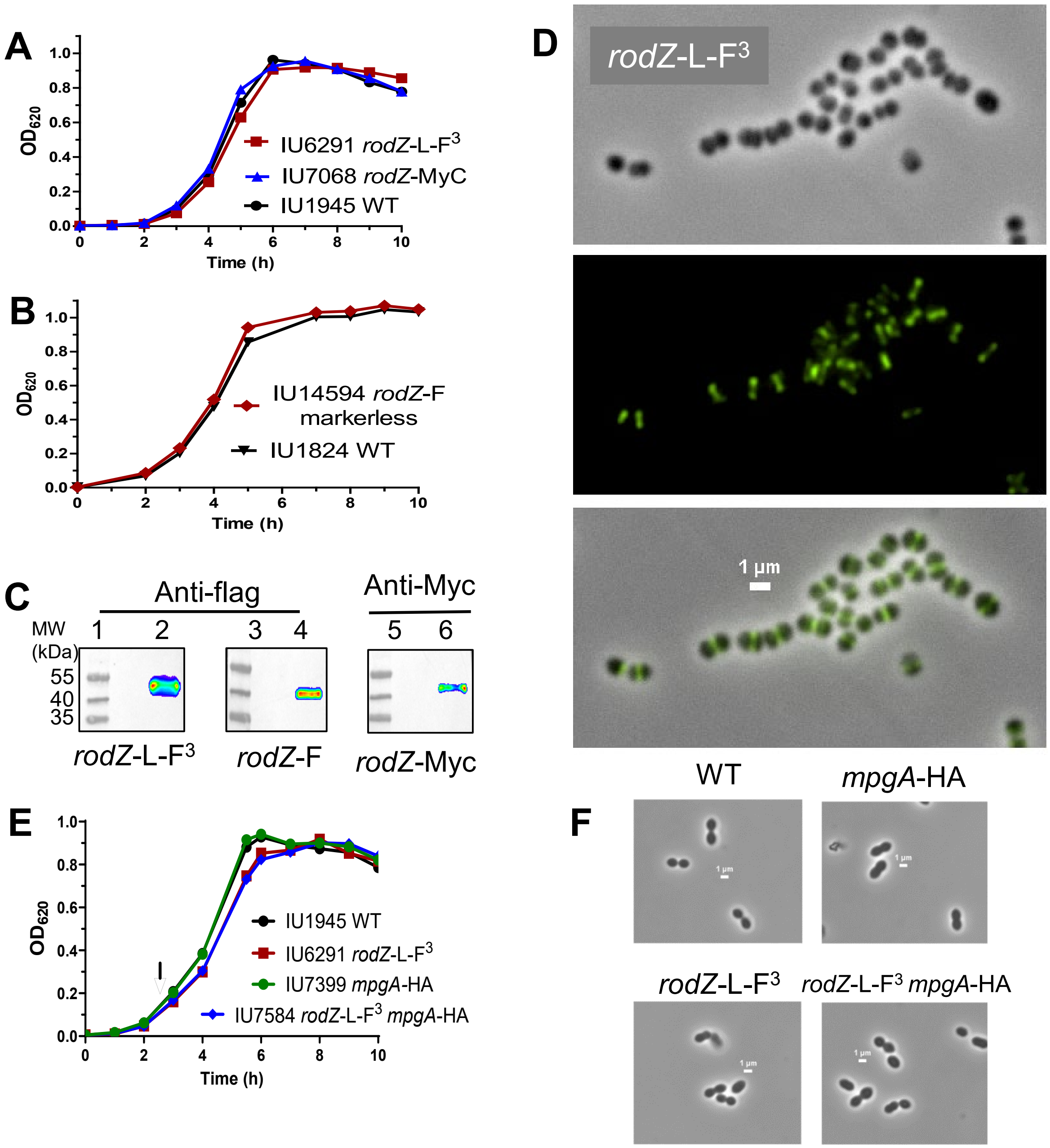


**Fig. S11. Epitope-tagged RodZ variants used for IFM and co-IP experiments are functional. (A)** and **(B)** Representative growth curves of strains IU6291 (*rodZ*-L-F$^3$), IU7068 (*rodZ*-MyC), and IU14594 (*rodZ*-F markerless) in comparison to the respective WT control strain, IU1824 or IU1945. **(C)** Visualization of epitope-tagged RodZ via western blot**.** Samples from lanes 1 to 6 are IU1945, IU6291, IU1824, IU14594, IU1945 and IU7068. **(D)** Representative immunofluorescence microscopy (IFM) images of IU6291 (*rodZ*-L-F$^3$). Panels shown from top to bottom are: phase contrast microscopy, fluorescence microscopy and overlay of phase/FITC. **(E)** Representative growth curves and (F) microscopic images of IU7584 (*rodZ*-L-F3 *mpgA*-HA) used for co-IP experiments. Cells were imaged at $OD_{620}$ of ≈0.1-0.2. Arrow indicates time of harvest. All growth and microscopy experiments were performed with three independent replicates with similar results.

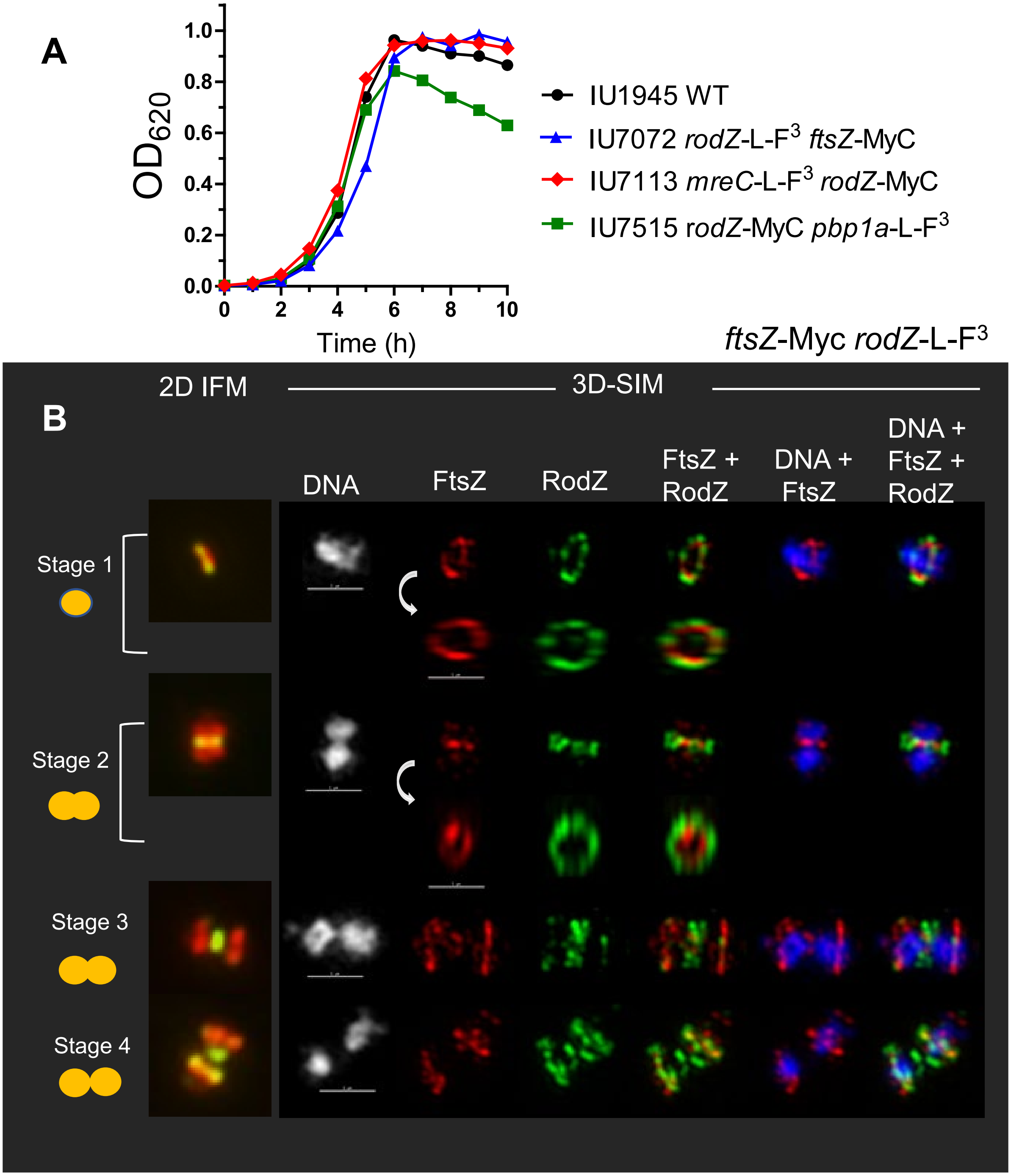


**Fig. S12. RodZ and FtsZ have overlapping, but different localization patterns in *Spn*. (A)** Representative growth curves of doubly epitope-tagged strains used in immunofluorescence microscopy (IFM) experiments and 3D-SIM. **(B)** Different localization patterns of RodZ and FtsZ during later stages of pneumococcal cell division. Representative 2D-IFM (left column) and 3D-SIM IFM and DAPI images of strain IU7072 (*ftsZ*-Myc *rodZ*-L-F3) at different division stages. DNA (DAPI stained image) is false-colored white or blue in columns 1 or 5, respectively. FtsZ and RodZ are pseudo-colored as red and green respectively, and overlapping signal is colored yellow. The first row of each panel represents images captured in the XY plane, while second row images were obtained by rotating a section of the mid-cell region around the X or Y axis. In stage 3 cells, FtsZ has begun to re-locate to equators, while RodZ remains largely at the septum (arrows). Images are representative of >20 examined cells in different division stages from two experiment. Scale bar = 1 µm.

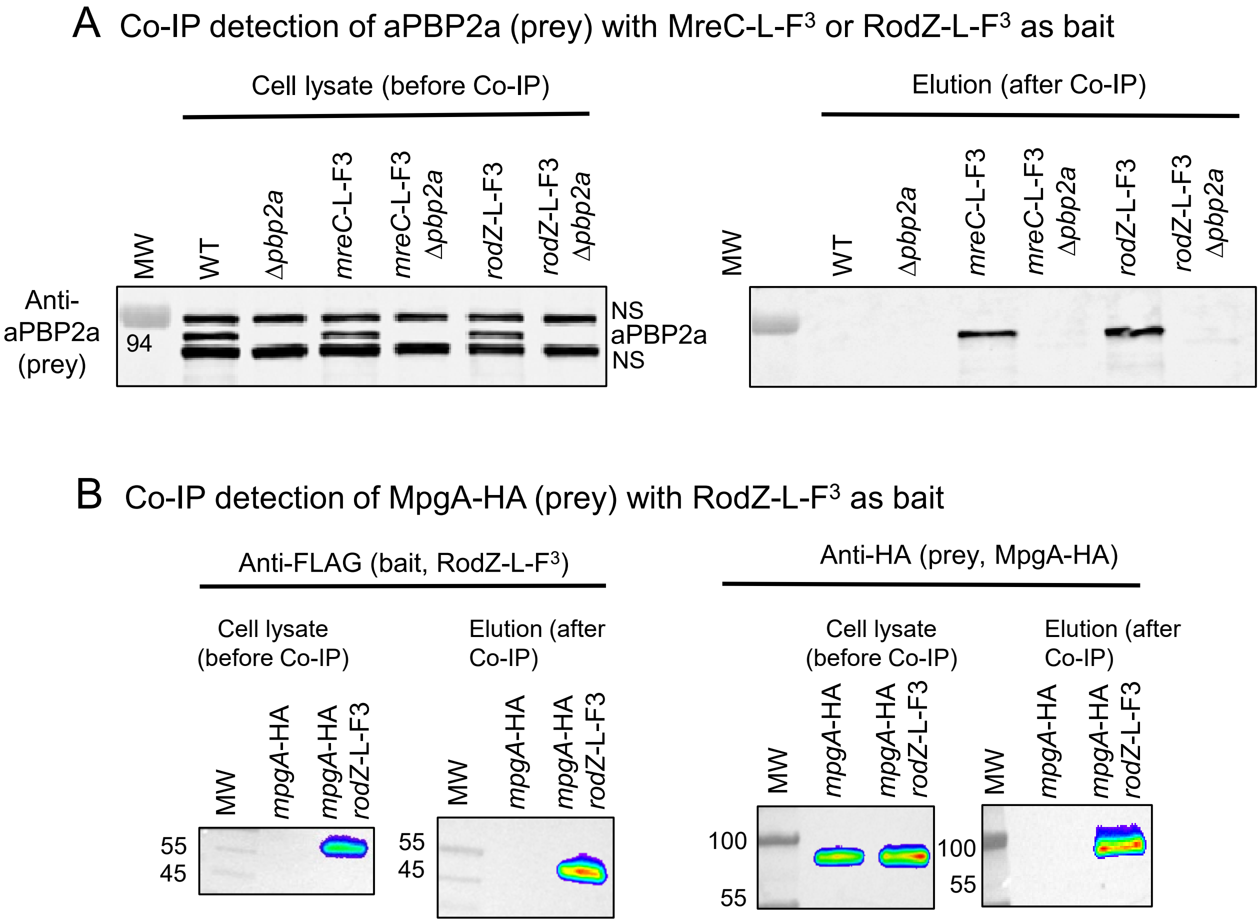


**Fig. S13. Complexes of RodZ with other proteins detected by co-IP. (A)** Co-IP detection of aPBP2a (prey) with MreC-L-$F^3$ or RodZ-L-$F^3$ as bait. Strains used were IU1945 (WT), K166 (Δ*pbp2a*), IU4970 (*mreC*-L-$F^3$), IU17817 (*mreC*-L-$F^3$ Δ*pbp2a*), IU6291 (*rodZ*-L-$F^3$), and IU17821 (*rodZ*-L-$F^3$ Δ*pbp2a*). Since anti-aPBP2a cross-reacts with two other proteins (NS) in addition to aPBP2a, *pbp2a* deletion strains were used in these studies to identify the aPBP2a-specific band. 6 µg (4 µl) of each lysate sample (input) were loaded in the left lanes, while 15 µL of each elution output sample were loaded in to right lanes. Expected molecular weight of aPBP2a is 81 kDa. Note that non-specific bands are not present in the output samples. **(B)** Co-IP detection of MpgA-HA (prey) with RodZ-L-$F^3$ as bait. Strains used were IU7399 (*mpgA*-HA), and IU7484 (*mpgA*-HA *rodZ*-L-$F^3$). ≈55 µg of lysate sample (input), or 15 µL of elution output sample were loaded on to each lane. Expected molecular weight of RodZ-L-$F^3$ and MpgA-$HA^3$ are 34, and 62 kDa, respectively. Blot images in A and B are representative of 4 and 3 independent experiments, respectively (continued on next page).

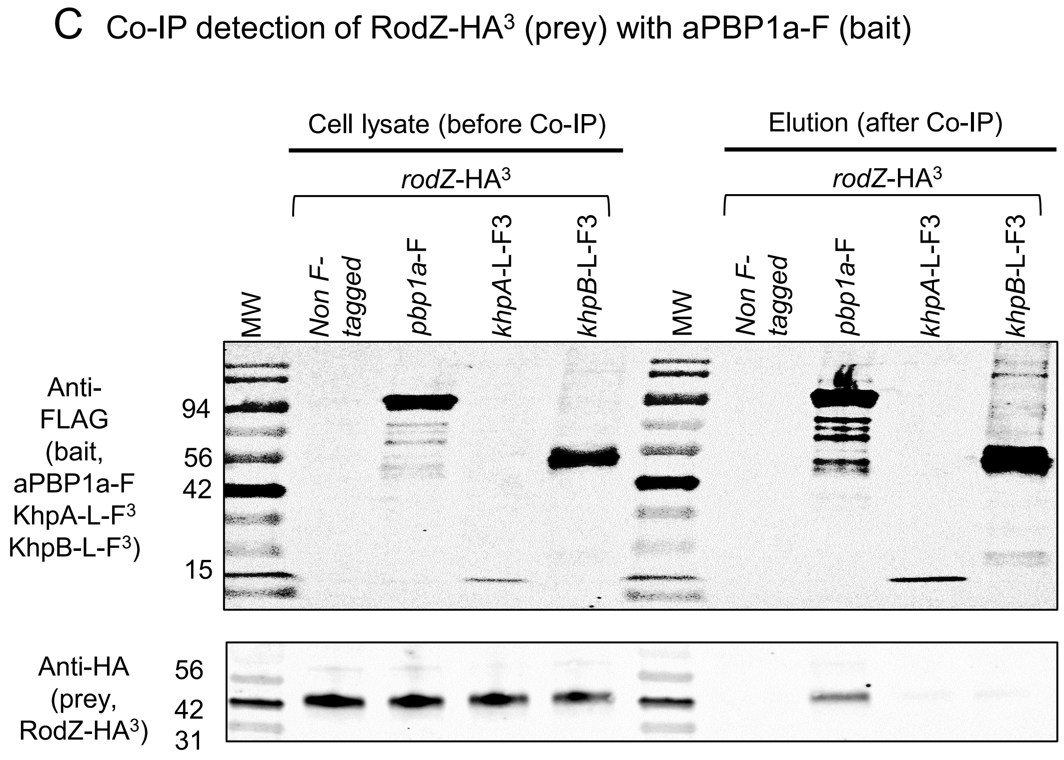


**Fig. S13C. (C)** Co-IP detection of RodZ-HA$^3$ (prey) with aPBP1a-F as bait, but not with KhpA-L-F$^3$ or KhpB-L-F$^3$ as bait. Strains used were IU11828 (*rodZ*-HA$^3$), IU11925 (*rodZ*-HA$^3$ *pbp1a*-F), IU17873 (*rodZ*-HA$^3$ *khpA*-L-F$^3$) and IU17877 (*rodZ*-HA$^3$ *khpB*-L-F$^3$). 9 µg (6 ul) of each lysate sample (input) was loaded in the left lanes, while 25 µL of elution each output sample was loaded in the right lanes. Expected molecular weight of aPBP1a-F, KhpA-L-F$^3$, KhpB-L-F$^3$, and RodZ-HA$^3$ are 81, 13, 35, and 34 kDa, respectively. Blot images in (C) and (B) are representative of 3 independent experiments. For A to C, average values of prey protein ratios are shown in Table 3.

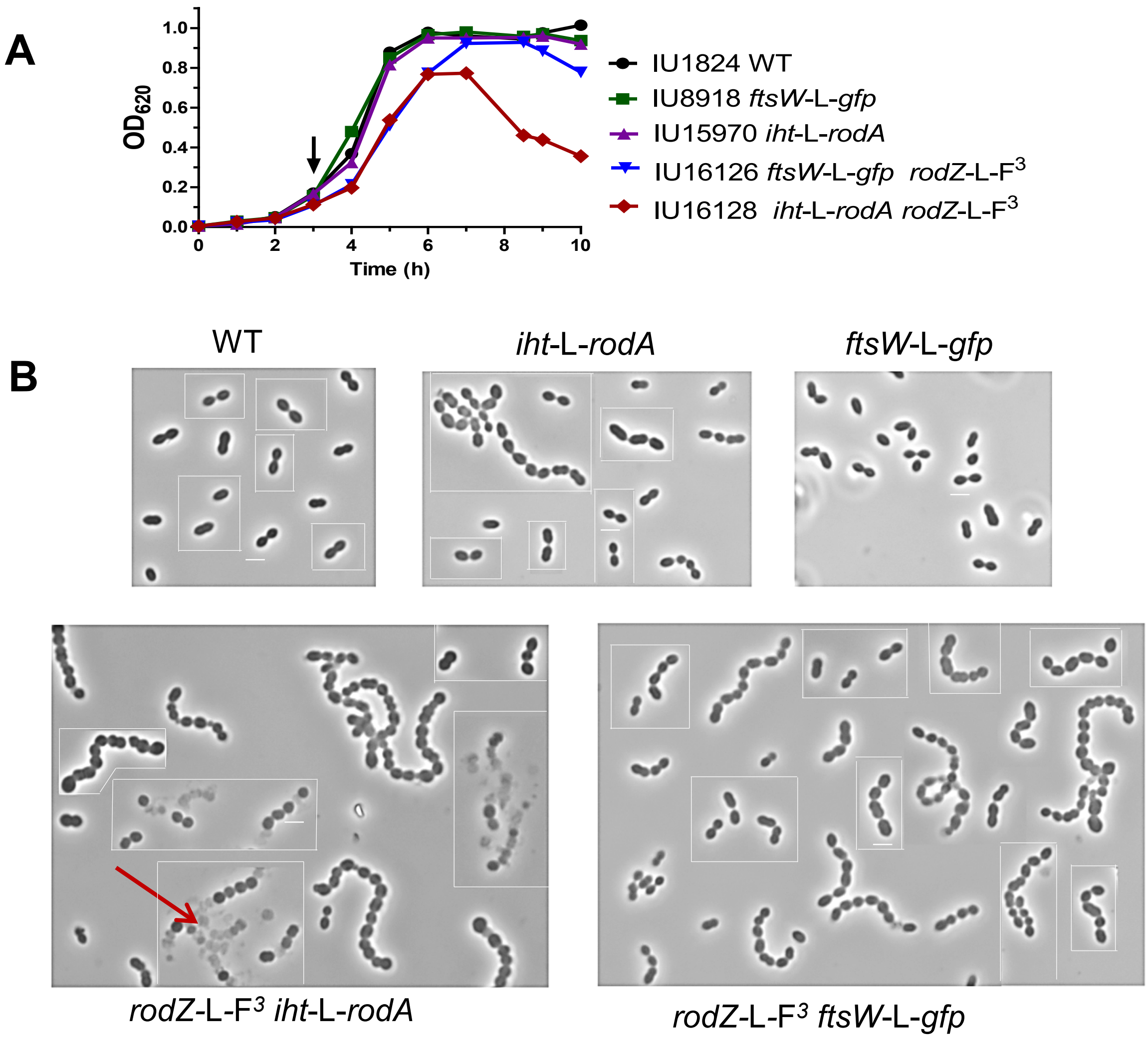


**Fig. S14. Co-IP detection of FtsW-GFP (prey) with RodZ-L-F$^3$ (bait). (A)** Representative growth curves and **(B)** microscopic images of tagged *rodA* or *ftsW* strains for co-IP experiments. Single tagged *ftsW*-L-*gfp* and *iht*-L-*rodA* strains grew similarly the to WT strain, while double-tagged *rodZ*-L-F$^3$ *iht*-L-*rodA* showed lysis in ≈50% of the cells (red arrow). As a result, *rodZ*-L-F$^3$ *iht*-L-*rodA* strain was not used for further experiments. Growth curves and microscopy experiments were performed twice with similar results. (Continued on next page)

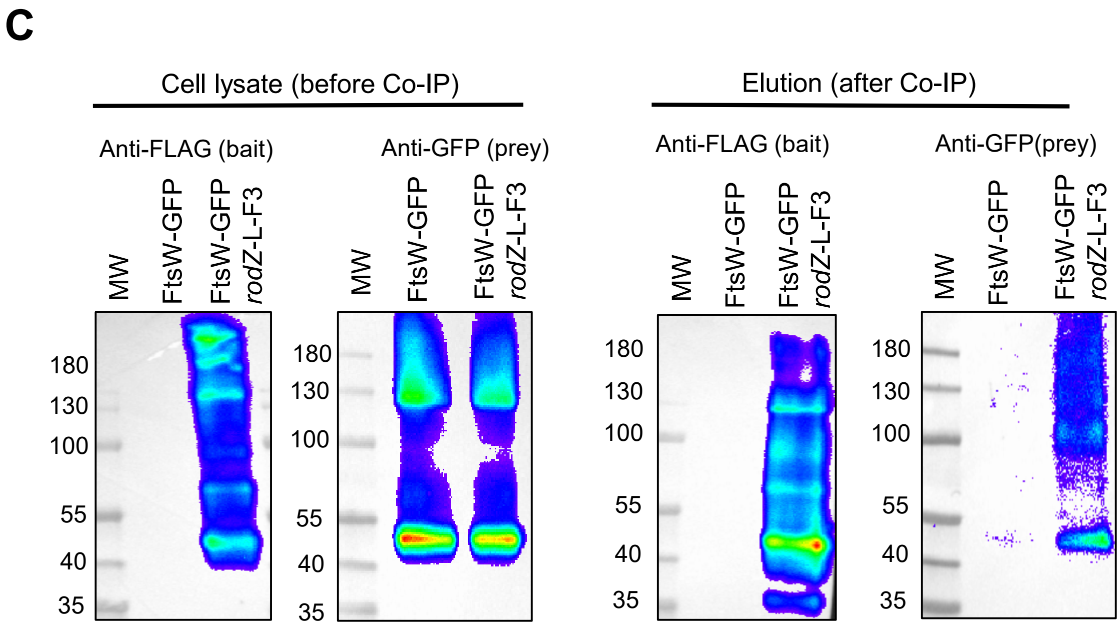


| Molecular Mass Reference | MW (KDa) |
|---|---|
| RodZ-L-F3 | 34 |
| FtsW | 45 |
| GFP | 27 |
| FtsW-GFP | 73 |
| 1:1 of RodZ-L-F3 and FtsW-GFP | 107 |

**Fig. S14. (C)** FtsW-GFP (prey) is eluted with RodZ-L-F$^3$ (bait) in co-IP from cross-linked *S. pneumoniae* cells of strain IU16126 (right lanes) compared with untagged control strain (IU8918; left lanes). Concentrated intact cells were cross-linked with 0.1% (wt/vol) paraformaldehyde, and the cross-linkages were not reversed by heating before loading onto the gels. FtsW-GFP complex with RodZ-L-F$^3$ was detected using anti-GFP in the output sample of IU16126, but not in IU8918. ≈50-60 µg of crude cell lysate and 40 µl of elution samples (output) were loaded in each lane. Growth curves, microscopy and co-IP experiments were performed twice with similar results.

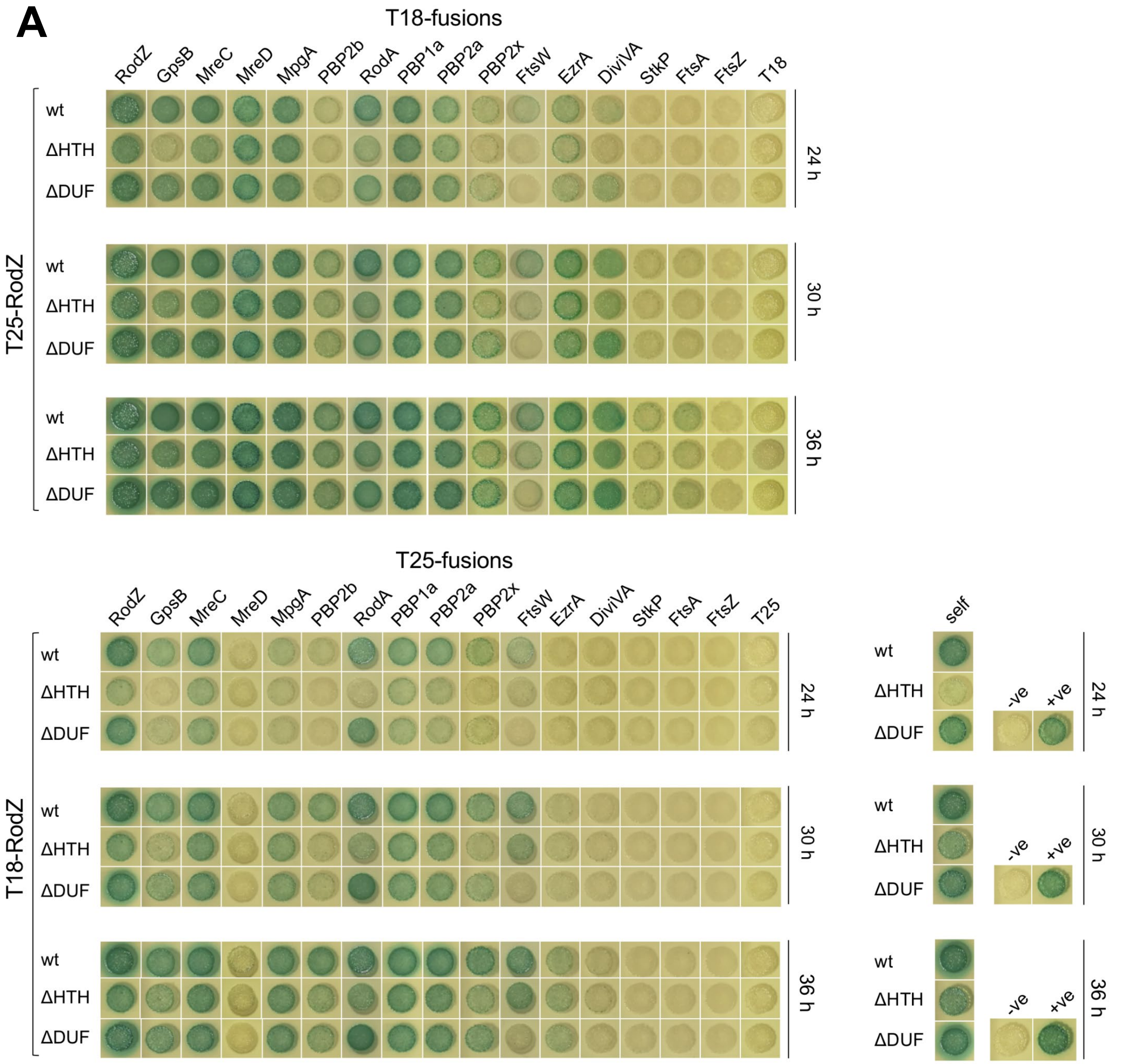


**Fig. S15A. Decreased interactions of RodZ (ΔHTH) and RodZ (ΔDUF) compared to RodZ WT with certain cell elongation and division proteins by B2H**. **(A)** B2H assay showing the interactions of RodZ(ΔHTH) and RodZ(ΔDUF) with selected cell elongation and division proteins and themselves in comparison with those of RodZ WT. The agar plates were photographed after 24, 30 and 36 h at 30°C in a time course experiment. No complete loss of interactions with any of the protein partners tested was observed for either RodZ ΔHTH and RodZ ΔDUF, although decreased interactions with some of the tested proteins were clear after 24 and 30 h for both truncated variants. Notably, RodZ ΔHTH shows a significant decrease in self-interactions with respect to RodZ WT and RodZ ΔDUF. The results of these experiments are summarized in Figure 9C.

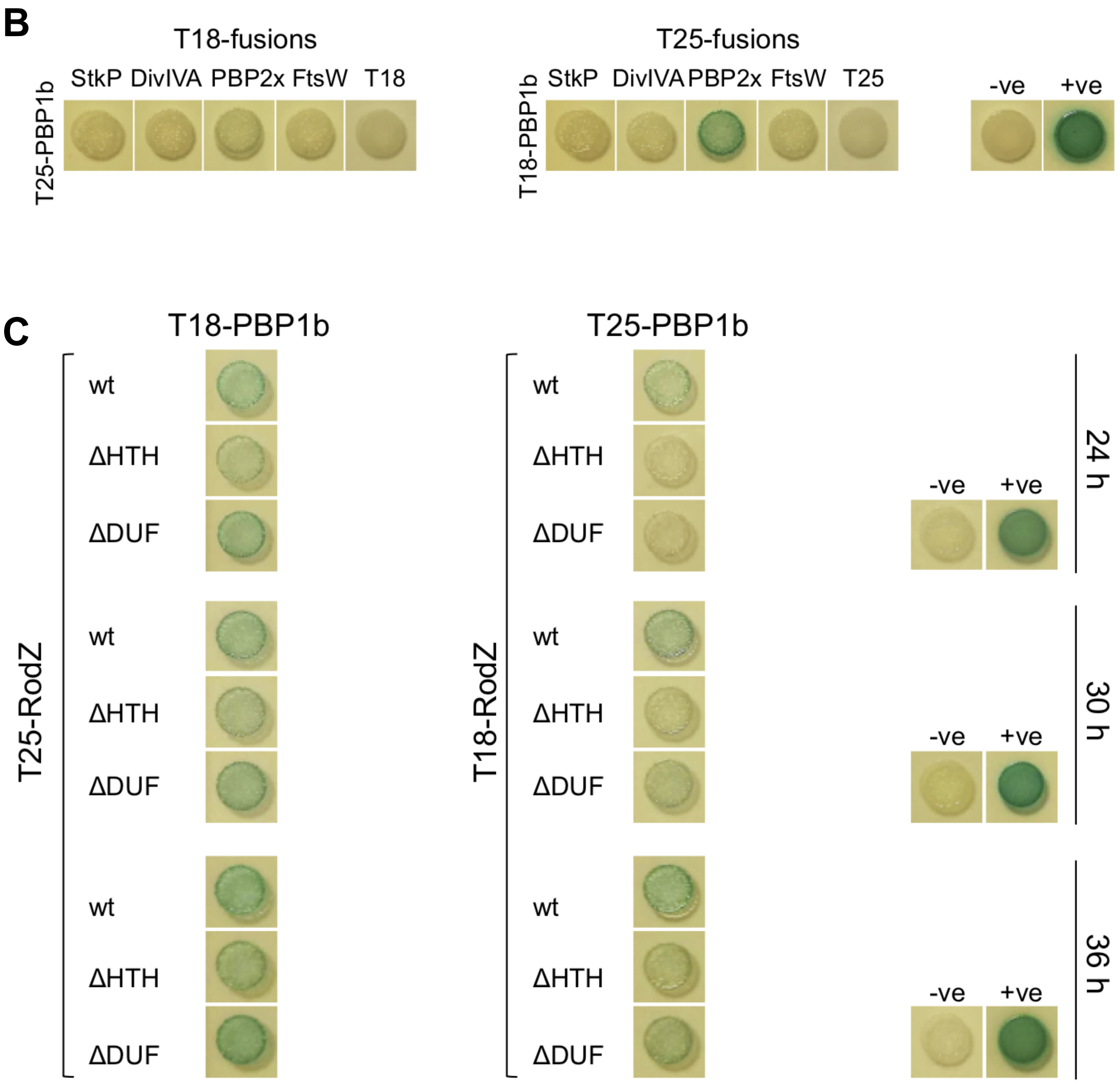


**Fig. S15BC. RodZ ΔHTH shows decreased interactions with aPBP1b with respect to RodZ WT.** B2H assays showing: **(B)** remaining interactions of aPBP1b with selected cell division proteins after 40 h at 30°C; and **(C)** interactions of aPBP1b with RodZ(ΔHTH) and RodZ(ΔDUF) in comparison with RodZ WT. The agar plates were photographed after 24, 30, and 36 h at 30°C in a time course experiment. No complete loss of interactions with aPBP1b was observed for either RodZ(ΔHTH) and RodZ(ΔDUF), although decreased interactions between RodZ(ΔHTH) and aPBP1b are clear at all time points.

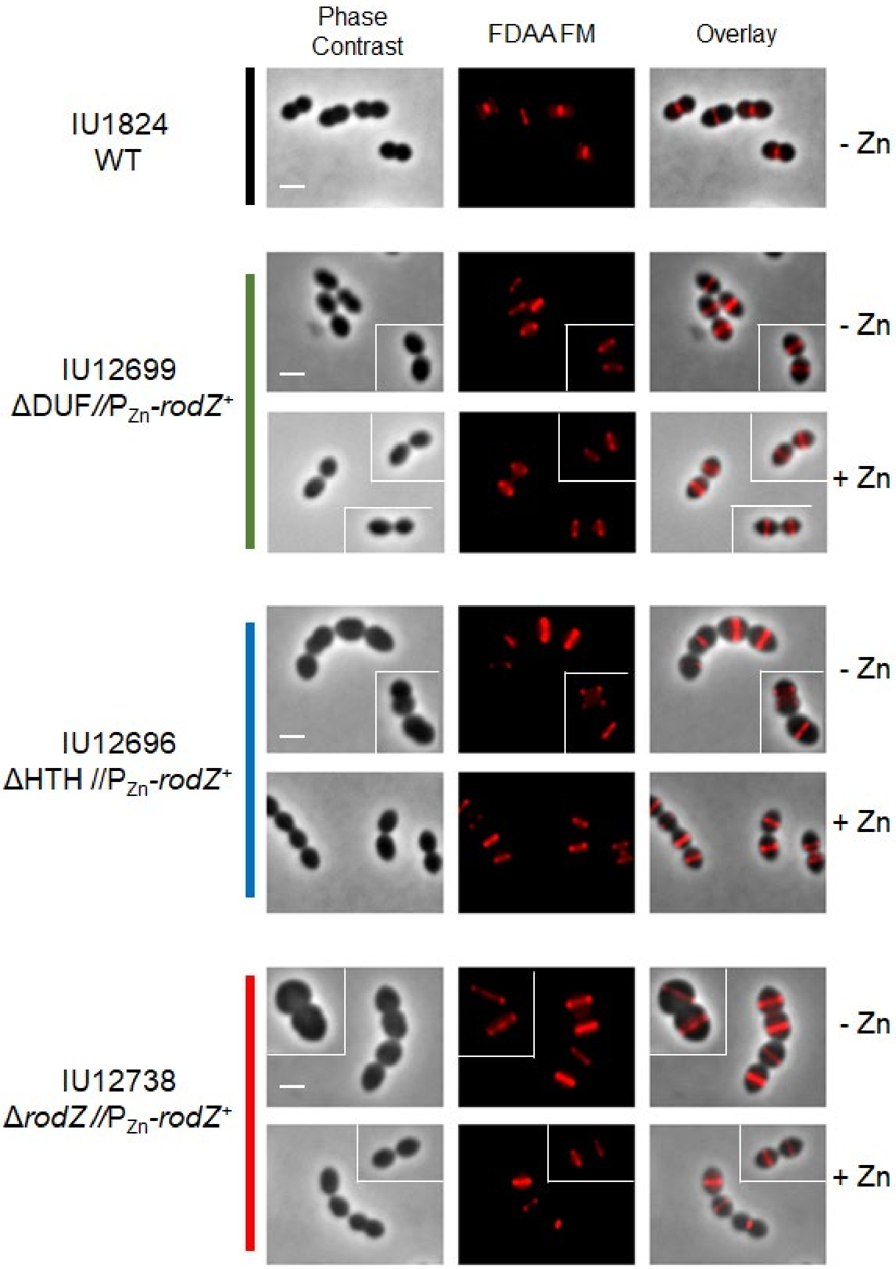


**Fig. S16. Fluorescent D-amino acids (FDAA) incorporation occurs at midcell upon RodZ WT depletion in cells expressing RodZ(ΔDUF) or RodZ(ΔHTH) or deleted for *rodZ*.** FDAA labels areas of new transpeptidase (TP) activity catalyzed by penicillin-binding proteins (PBPs). Overnight cultures of IU1824, IU12696, IU12699, and IU12738 were grown in BHI and resuspended to an $OD_{620}$ ≈0.003 in fresh BHI with or without Zn inducer (0.4 mM $ZnCl_2$ + 0.04 mM $MnSO_4$). At 4 h, samples were labeled with 250 µM TADA ( pseudo-colored red) for 2.5 min, then washed and visualized using fluorescence microscopy. Images are mosaic and representative of cells observed within a single field. Similar results are obtained from 3 independent biological experiments.

#### *mreC*-L-F$^3$ and *pbp2b*-HA growth curves

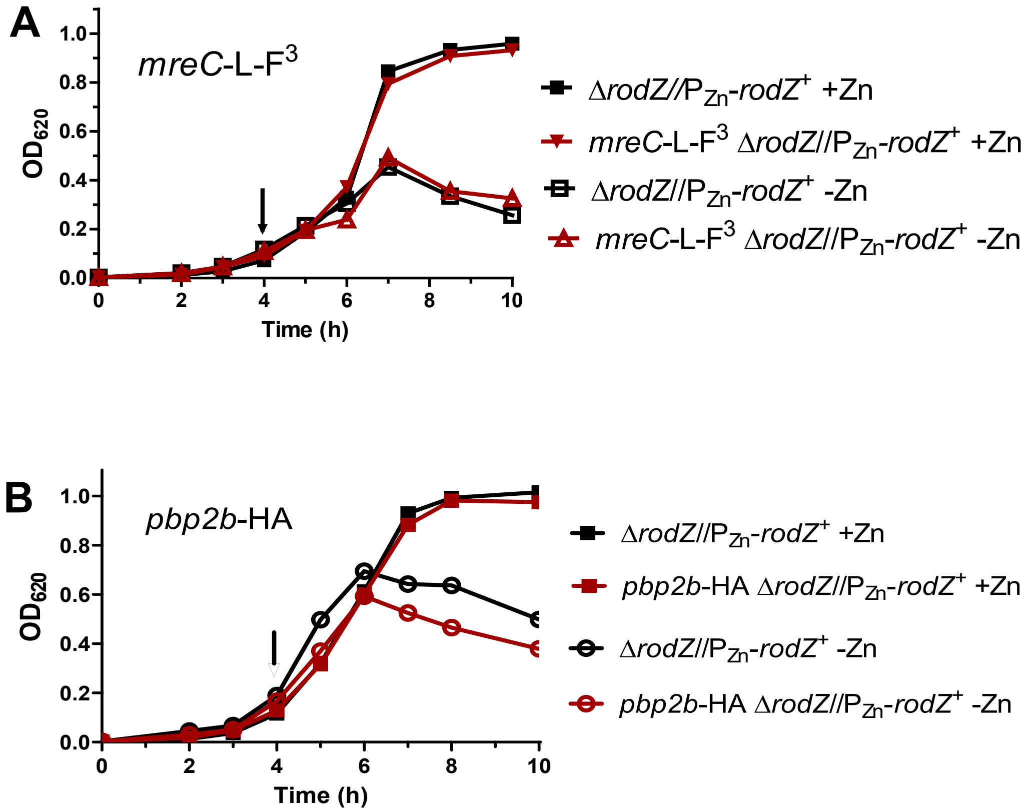


**Fig. S17. Representative growth curves of strains expressing MreC-L-F$^3$ (A) or bPBP2b-HA (B) depleted for RodZ.** Strains used: IU14158 (*mreC*-L-F$^3$ Δ*rodZ*//P$_{Zn}$-*rodZ*$^+$), IU14431 (*pbp2b*-HA Δ*rodZ*//P$_{Zn}$-*rodZ*$^+$), and untagged parent strain IU12738 (Δ*rodZ*//P$_{Zn}$-*rodZ*$^+$). Overnight cultures were grown in BHI broth with Zn inducer (0.4 mM ZnCl + 0.04 mM $MnSO_4$), and resuspended in fresh BHI broth with or without inducer. Arrows indicate the time at which samples were harvested for imaging. Growth curves were performed three times with similar results.

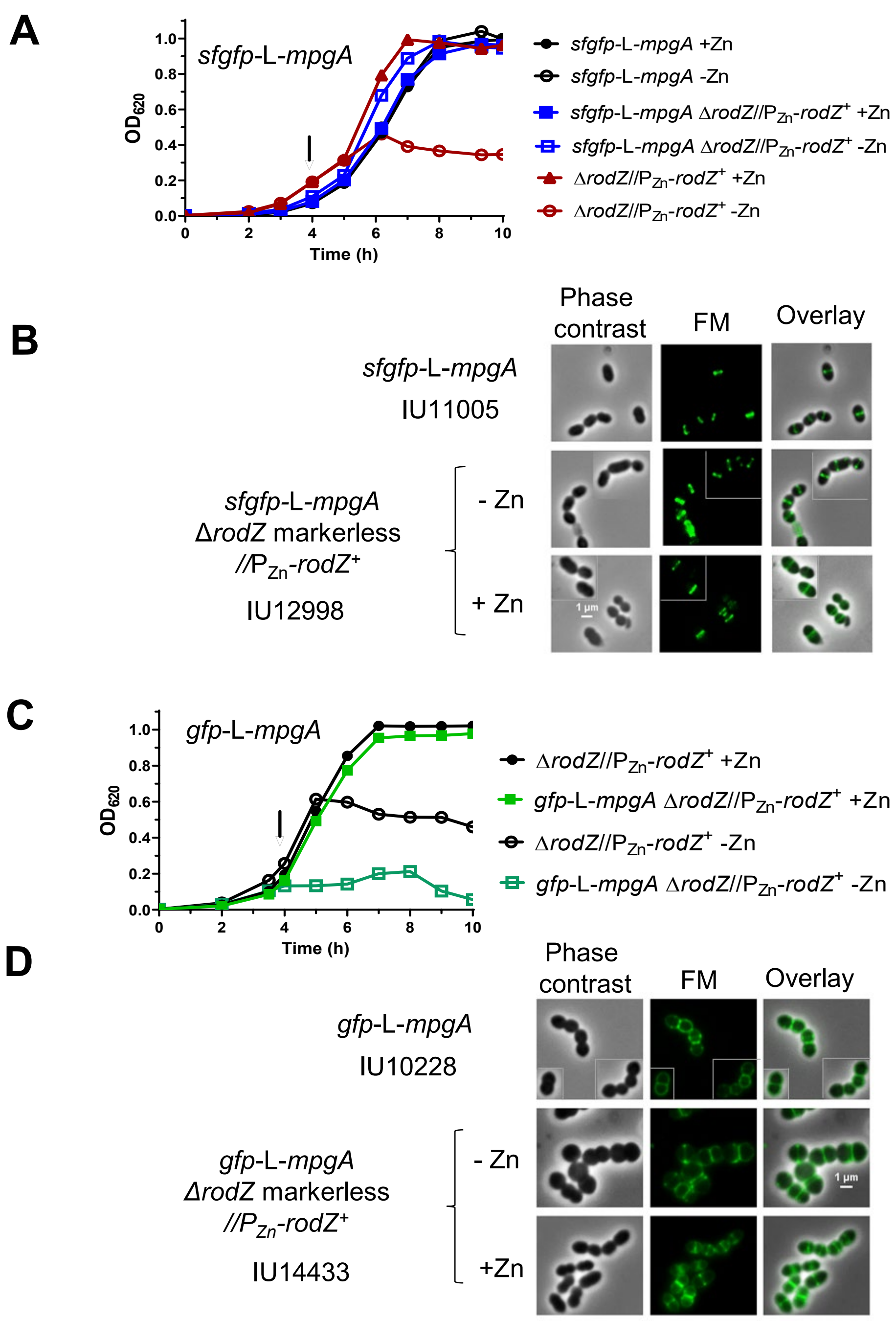


**Fig S18. *sfgfp-mpgA* (formerly *mpgA*(*Spn*)) suppresses Δ*rodZ* lethality, while *gfp-mpgA* exacerbates Δ*rodZ* lethality.** **(A)** Representative growth curves of strains IU11005 (*sfgfp*-L-*mpgA*), IU12738 (Δ*rodZ*//P$_{zn}$-*rodZ*$^{+}$), and IU12998 (*sfgfp*-L-*mpgA* Δ*rodZ*//P$_{zn}$-*rodZ*$^{+}$) grown in the presence or absence of Zn inducer (0.4 mM ZnCl + 0.04 mM $MnSO_4$) as described for Fig. S17. **(B)** Representative images of *sfgfp*-L-*mpgA* strains at 4h of growth (arrow in growth curves). **(C)** Representative growth curves of strains IU12738 (Δ*rodZ*//P$_{zn}$-*rodZ*$^{+}$) and IU14433 (*gfp*-L-*mpgA* Δ*rodZ*//P$_{zn}$-*rodZ*$^{+}$) grown in the presence or absence of Zn inducer. (D) Mosaic of representative images at 4 h of growth (arrow on growth curve). Growth curves and microscopy experiments were performed three times with similar results.

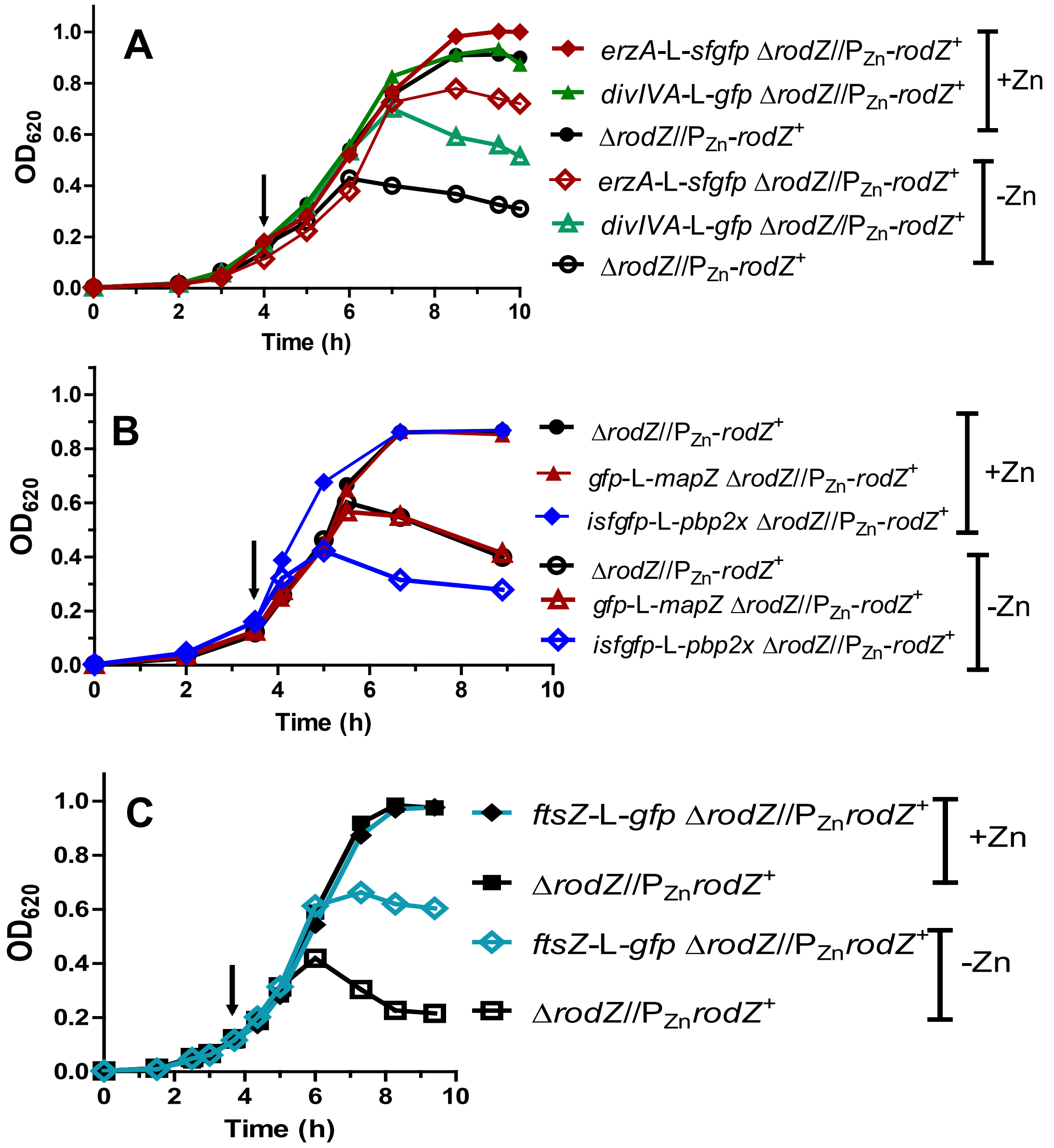


**Fig S19. FtsZ, EzrA, bPBP2x, DivIVA, and MapZ maintain midcell localization upon RodZ depletion.** Representative growth curves of strains grown with or without Zn inducer. **(A)** IU12738 (Δ*rodZ*//$P_{zn}$-*rodZ*$^+$), IU13058 (*ezrA*-L-*sfgfp* Δ*rodZ*//$P_{zn}$-*rodZ*$^+$), and IU13061 (*divIVA*-L-*gfp* Δ*rodZ*//$P_{zn}$-rod*Z*$^+$). **(B)** IU12738, IU13062 (*gfp*-L-*mapZ* Δ*rodZ*//$P_{zn}$-*rodZ*$^+$ ), and IU13000 (*isfgfp*-L-*pbp2x* Δ*rodZ*//$P_{zn}$-*rodZ*$^+$). **(C)** IU12738 and IU12993 (*ftsZ*-L-*sfgfp* Δ*rodZ*//$P_{zn}$-*rodZ*$^+$). Arrows indicate time points (4 h) of sampling microscopy. **(D)** and **(E)** (continued on next pages) Mosaics of representative micrographic images at 4 h of growth. Fluorescence microscopy was done as described in *Experimental procedures*. Similar results were obtained in three independent biological replicates. (Continued on next pages)

#### (D) FtsZ, EzrA and PBP2x localization during RodZ depletion

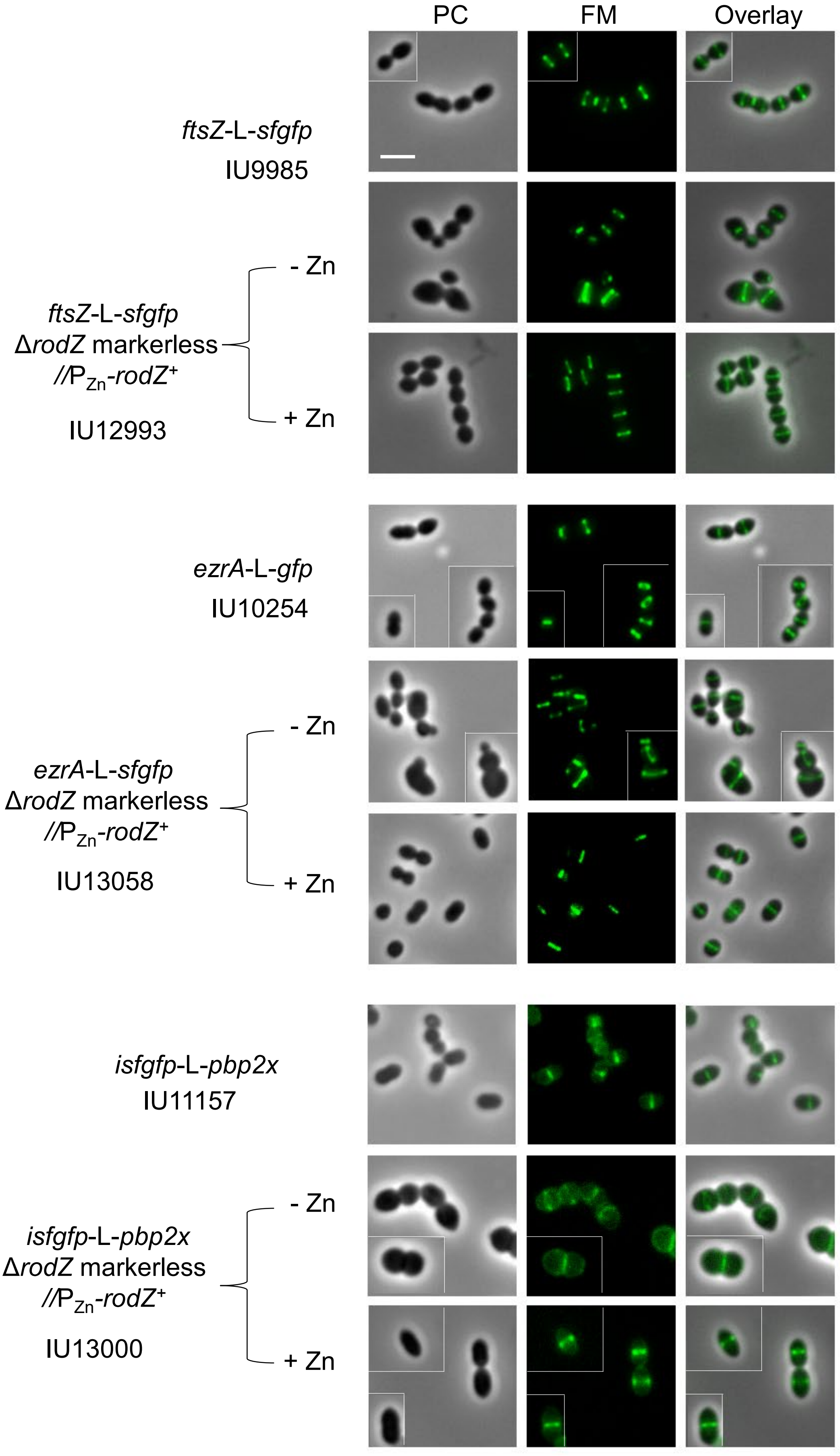

### (E) DivIVA and MapZ localization during RodZ depletion

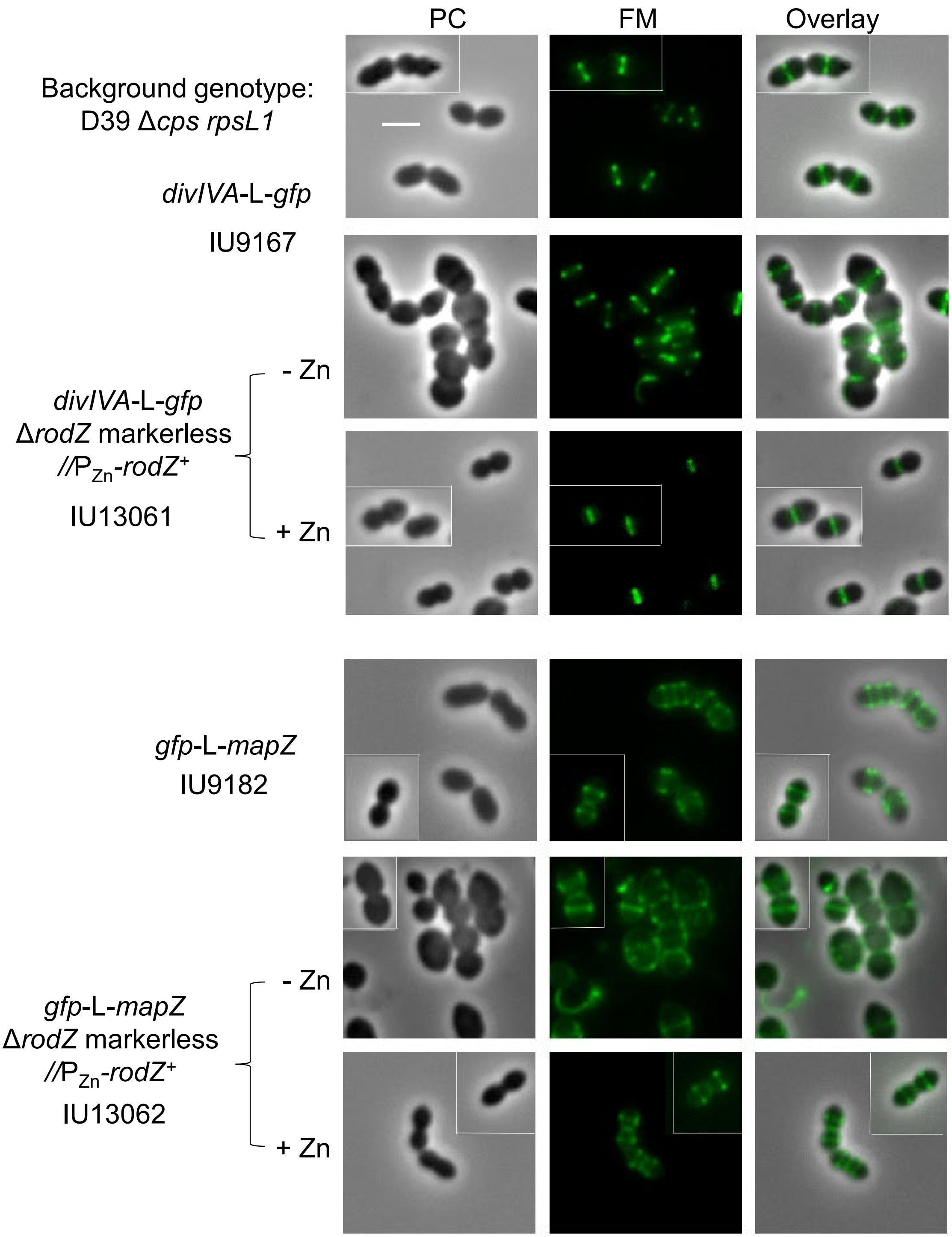

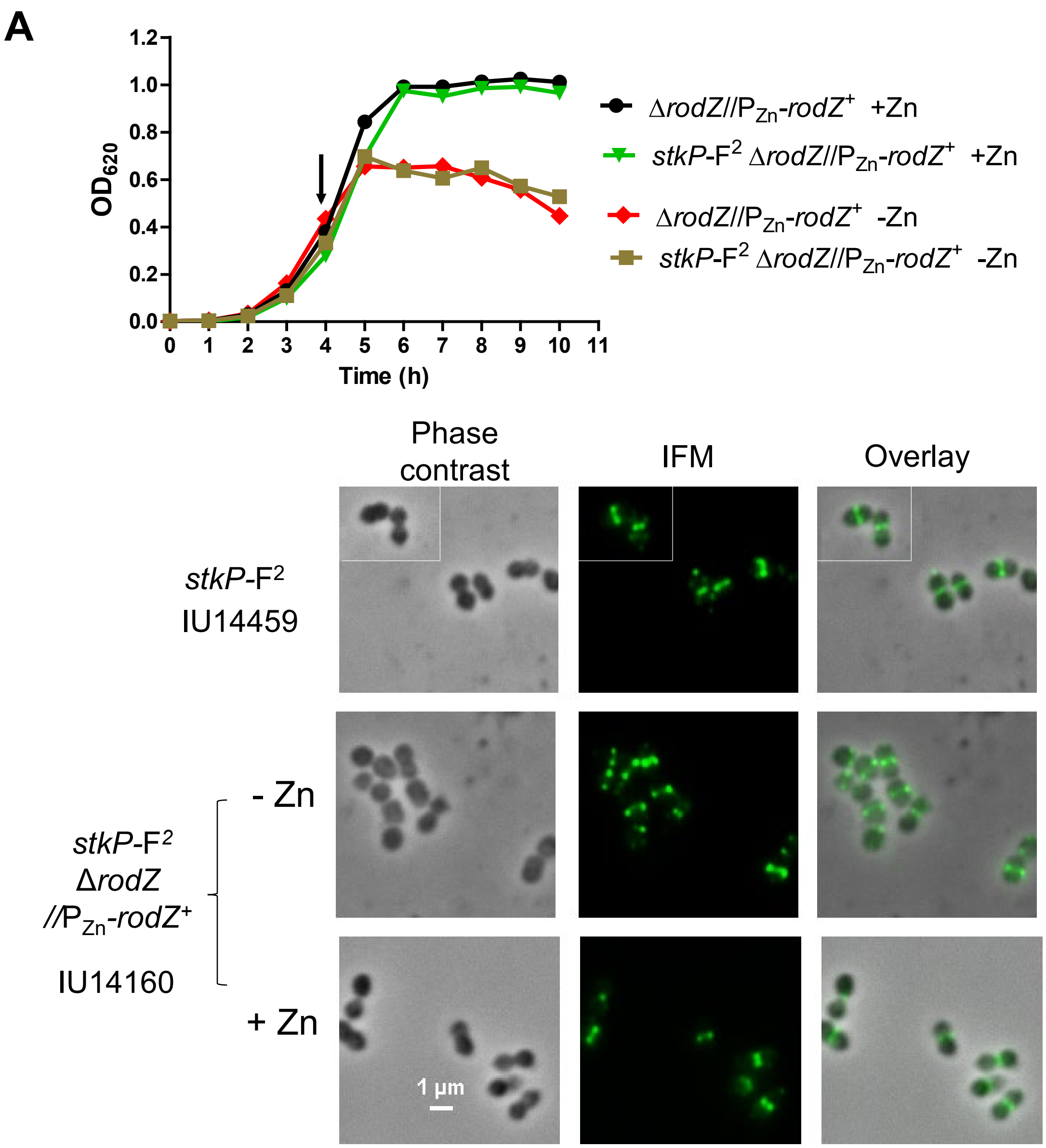


**Fig. S20. StkP, aPBP1a, and FtsA maintain midcell localization upon RodZ depletion. (A)** StkP. Representative growth curves of IU14160 (*stkP*-F$^2$ Δ*rodZ*//P$_{Zn}$-*rodZ*$^+$) and IU12738 (Δ*rodZ*//P$_{Zn}$-*rodZ*$^+$) with or without Zn inducer (0.4 mM $ZnCl_2$ + 0.04 mM $MnSO_4$). Arrow indicates the time at which samples were taken for imaging. Representative images are shown of IU14459 (*stkP*-F$^2$) and IU14160 (*stkP*-F$^2$ Δ*rodZ*//P$_{Zn}$-*rodZ*$^+$) cells harvested at 4 h and processed for IFM as described in *Experimental procedures*. Growth curves and IFM experiments were performed three times independently with similar results. (Continued on next page).

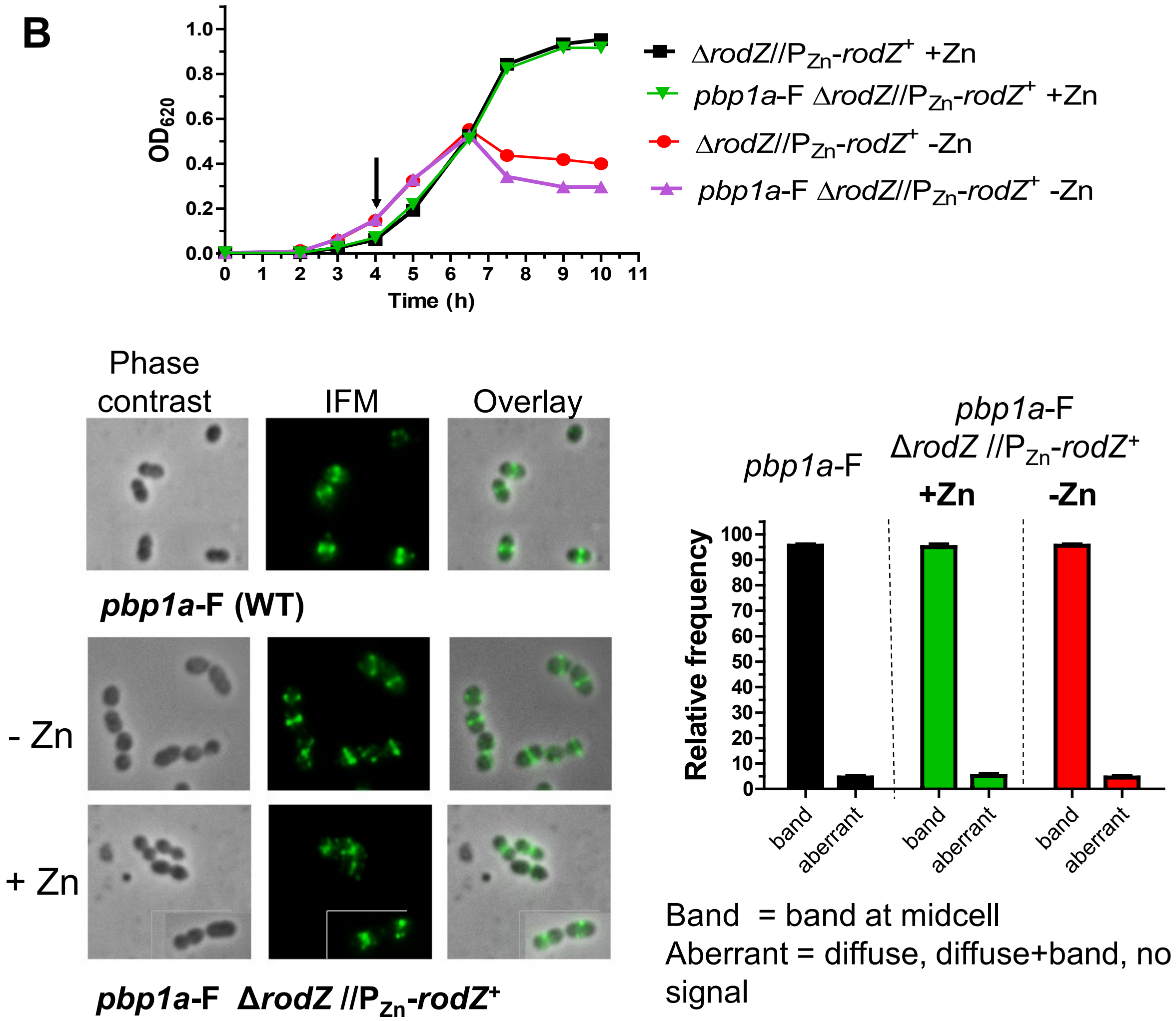


**Fig. S20. (B)** aPBP1a. Representative growth curves of IU14496 (*pbp1a*-F Δ*rodZ*//$P_{Zn}$-*rodZ*$^{+}$) and IU12738 (Δ*rodZ*//$P_{Zn}$-*rodZ*$^{+}$) with or without Zn inducer (0.4 mM $ZnCl_2$ + 0.04 mM $MnSO_4$). Arrow indicates the time at which samples were harvested for imaging. Representative images are shown of IU14494 (*pbp1a*-F) and IU14496 (*pbp1a*-F Δ*rodZ*//$P_{Zn}$-*rodZ*$^{+}$) cells harvested at 4 h and processed for IFM as described in *Experimental procedures*. Quantitation of localization pattern of aPBP1a based on IFM images is graphed for IU14494, and IU14496 grown in the presence or absence of the Zn inducer. 100 cells from two independent experiments were scored for each strain and condition. (Continued on next page)

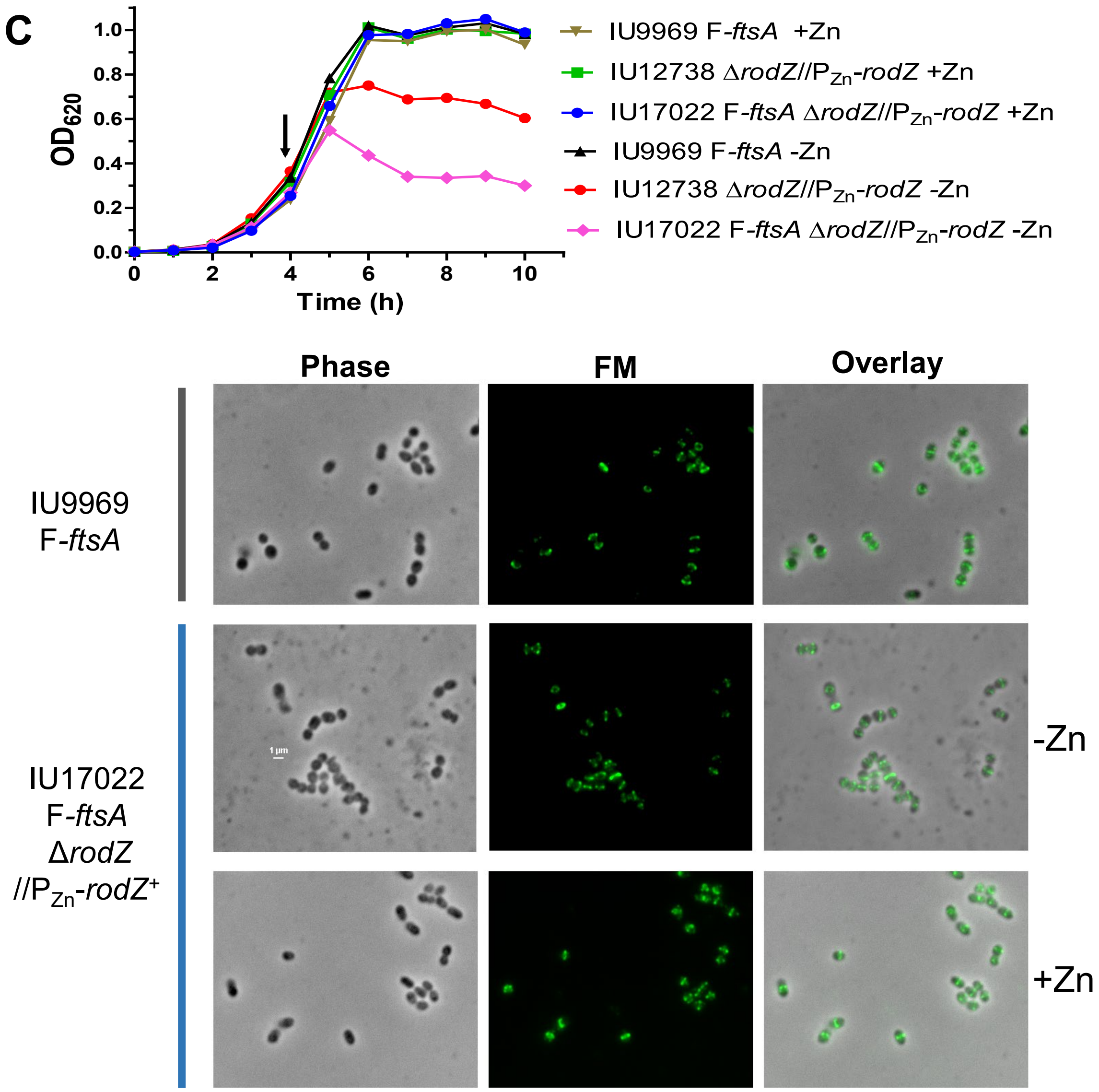


**Fig. S20. (C)** FtsA. Representative growth curves and IFM images of IU9969 (F-*ftsA*), IU17022 (F-*ftsA* Δ*rodZ*//$P_{Zn}$-*rodZ*$^+$) and IU12738 (Δ*rodZ*//$P_{Zn}$-*rodZ*$^+$) at 4h of growth. Growth curves and IFM images are representative of 2 independent experiments. Two other strains with genotypes of *ftsA'-sfgfp-ftsA'* Δ*rodZ*//$P_{Zn}$-*rodZ*$^+$ (IU14199) and *gfp-ftsA* Δ*rodZ*//$P_{Zn}$-*rodZ*$^+$ (IU17024) constructed to study the localization of FtsA during RodZ depletion were not used because these strains showed aberrant morphologies in the presence of the inducer and highly defective growth in the absence of the inducer.

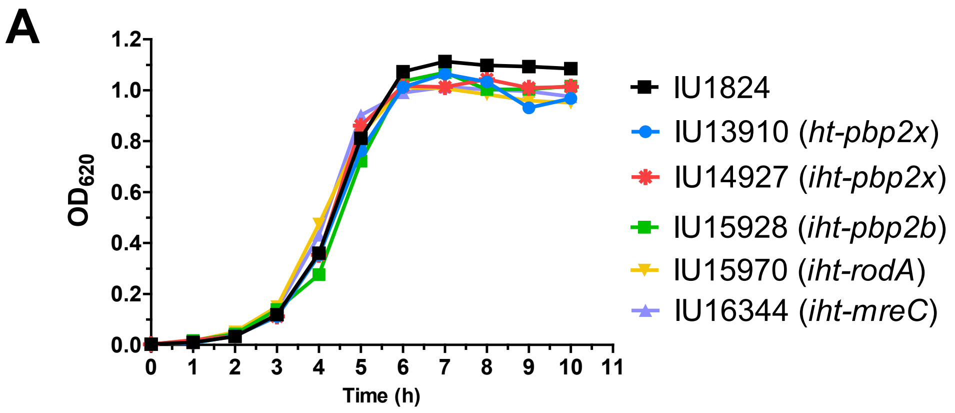


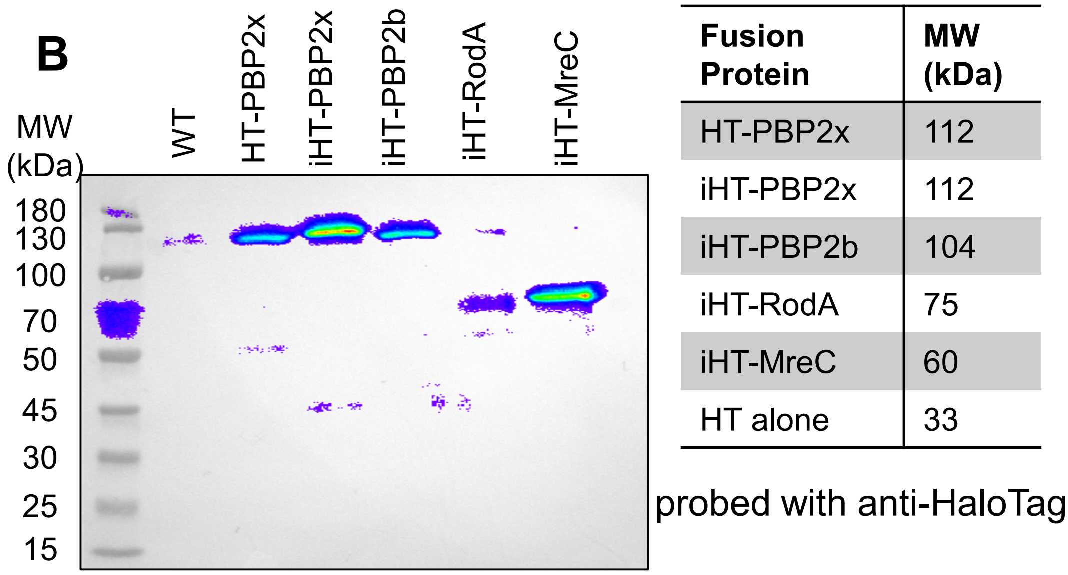


| Fusion Protein | MW (kDa) |
|---|---|
| HT-PBP2x | 112 |
| iHT-PBP2x | 112 |
| iHT-PBP2b | 104 |
| iHT-RodA | 75 |
| iHT-MreC | 60 |
| HT alone | 33 |

**Fig. S21. HT-fusion proteins are functional in strains grown in BHI at 37°C. (A)** Growth curves of HT-fusion strains in BHI medium. Strains used are IU1824 (WT), IU13910 (*ht-pbp2x*), IU14927 (*iht-pbp2x*), IU15928 (*iht-pbp2b*), IU15970 (*iht-rodA*), and IU16344 (*iht-mreC*). **(B)** Western blot of fusion strains. 4 µg of lysates obtained from cells grown to $OD_{620}$ ≈0.15 to 0.2 were loaded in each lane. Monoclonal anti-HT mouse antibody and secondary anti-mouse antibody conjugated to HRP were used at 1:1000, and 1:3300 dilutions, respectively. Growth curves and western blot results are representative of 2 and 3 experiments, respectively. (Continued on next page)

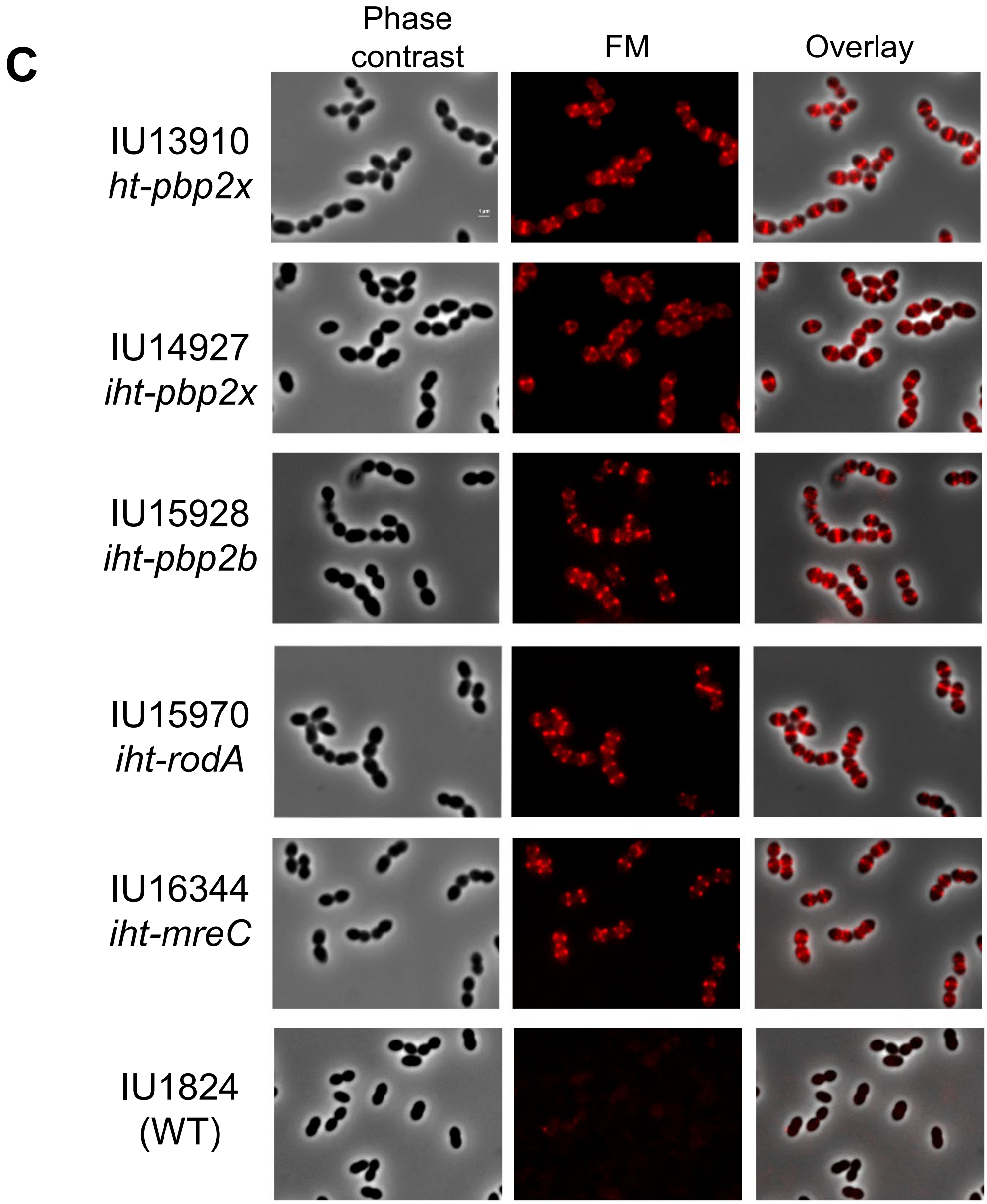


**Fig. S21. (C)** Localization patterns of HaloTag fusions to bPBP2x, bPBP2b, RodA, and MreC. Strains used are IU1824 (WT), IU13910 (*ht-pbp2x*), IU14927 (*iht-pbp2x*), IU15928 (*iht-pbp2b*), IU15970 (*iht-rodA*), and IU16344 (*iht-mreC*). Strains were harvested at mid-log phase (4h of growth), labeled with saturating concentration of HT-TMR ligand (0.83 μM) and visualized with phase contrast microscopy and fluorescence microscopy as described in *Experimental procedures*. Figure panels from left to right: phase contrast microscopy (cell bodies black), fluorescence microscopy (labeled fusion proteins red, Texas Red filter) and overlay. All images are to scale with scale bar representing 1 μm. Micrographs are representative of three biological replicates. (Continued on next page)

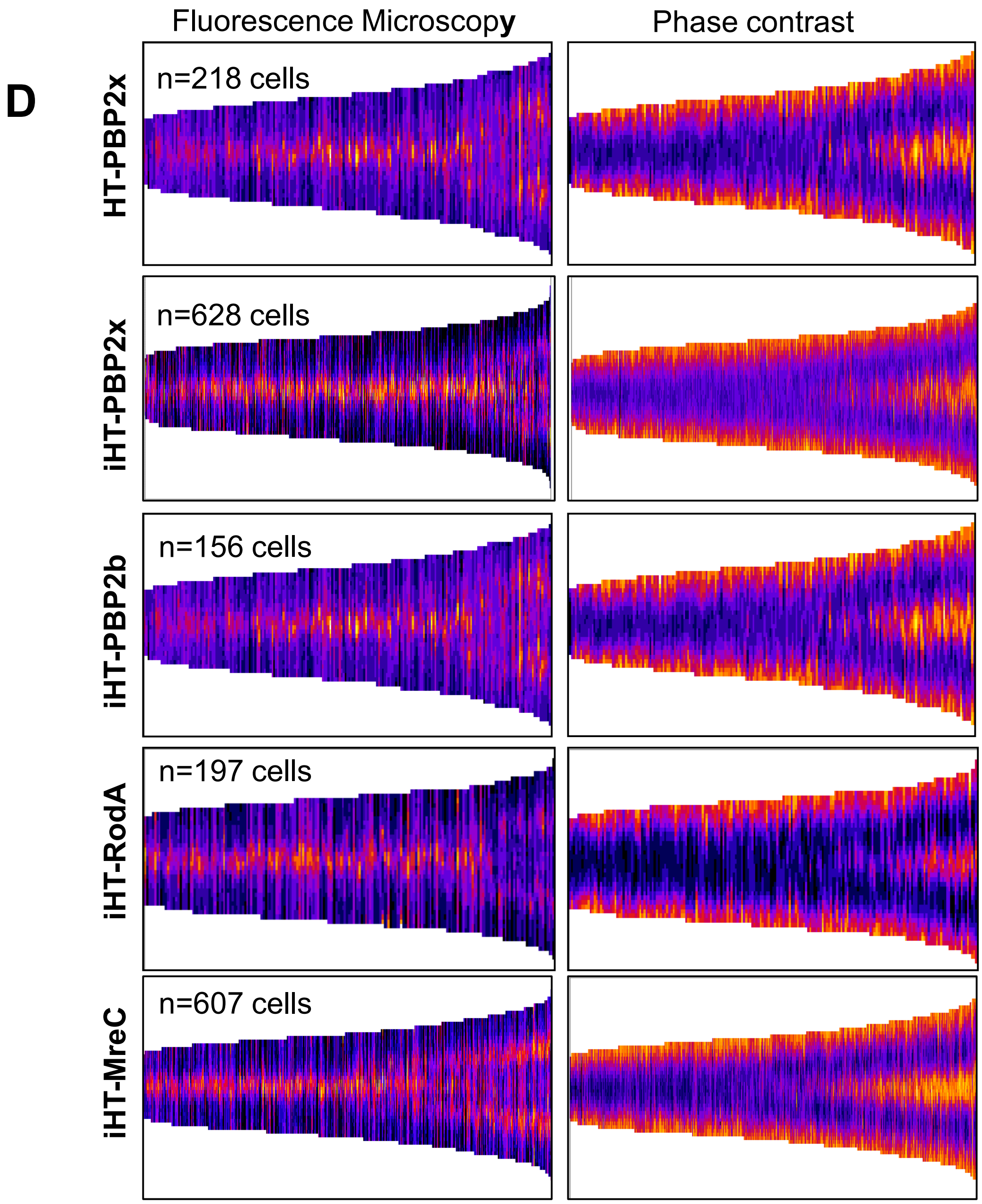


**Fig. S21. (D)** Demographs based on fluorescence intensity (HT-fusions) and light scattering (cell body) generated from strains expressing HT-fusion proteins as listed in legend to Fig. S21C. Cell images were processed by using MicrobeJ to generate demographs as described in (Perez et al., 2019) and *Experimental procedures.* Cells are sorted by length from shorter (left) to longer (right), corresponding to pre-divisional single cells to late-divisional daughter cells about to separate, respectively. n indicates the number of cells aligned within a given demograph. Each demograph is representative of one of 3 independent biological replicates, which provide similar results.

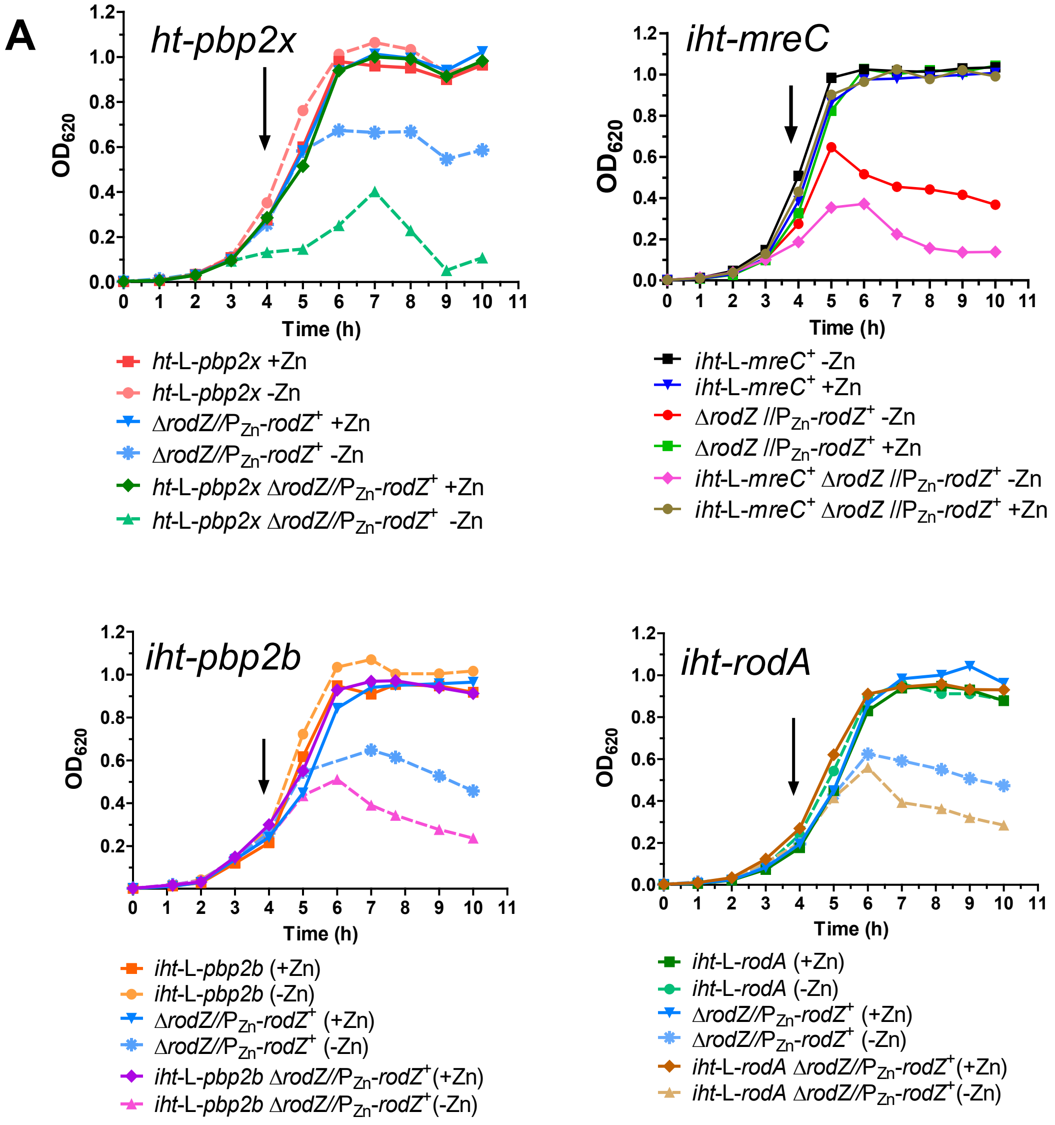


**Fig S22. bPBP2b and RodA, but not bPBP2x, mislocalize upon RodZ depletion.** **(A)** Growth curves of HT-fusion strains. Strains used are IU12738 (Δ*rodZ*//$P_{Zn}$-*rodZ*$^+$), IU13910 (*ht-pbp2x*), IU16062 (*ht-pbp2x* Δ*rodZ*//$P_{Zn}$-*rodZ*$^+$), IU16344 (*iht-mreC*), IU16920 (*iht-mreC* Δ*rodZ*//$P_{Zn}$-*rodZ*$^+$), IU15928 (*iht-pbp2b*), IU16058 (*iht-pbp2b* Δ*rodZ*//$P_{Zn}$-*rodZ*$^+$), IU15970 (*iht-rodA*), and IU16060 (*iht-rodA* Δ*rodZ*//$P_{Zn}$-*rodZ*$^+$). Growth curves were performed 3 times with similar results. (Continued on next page)

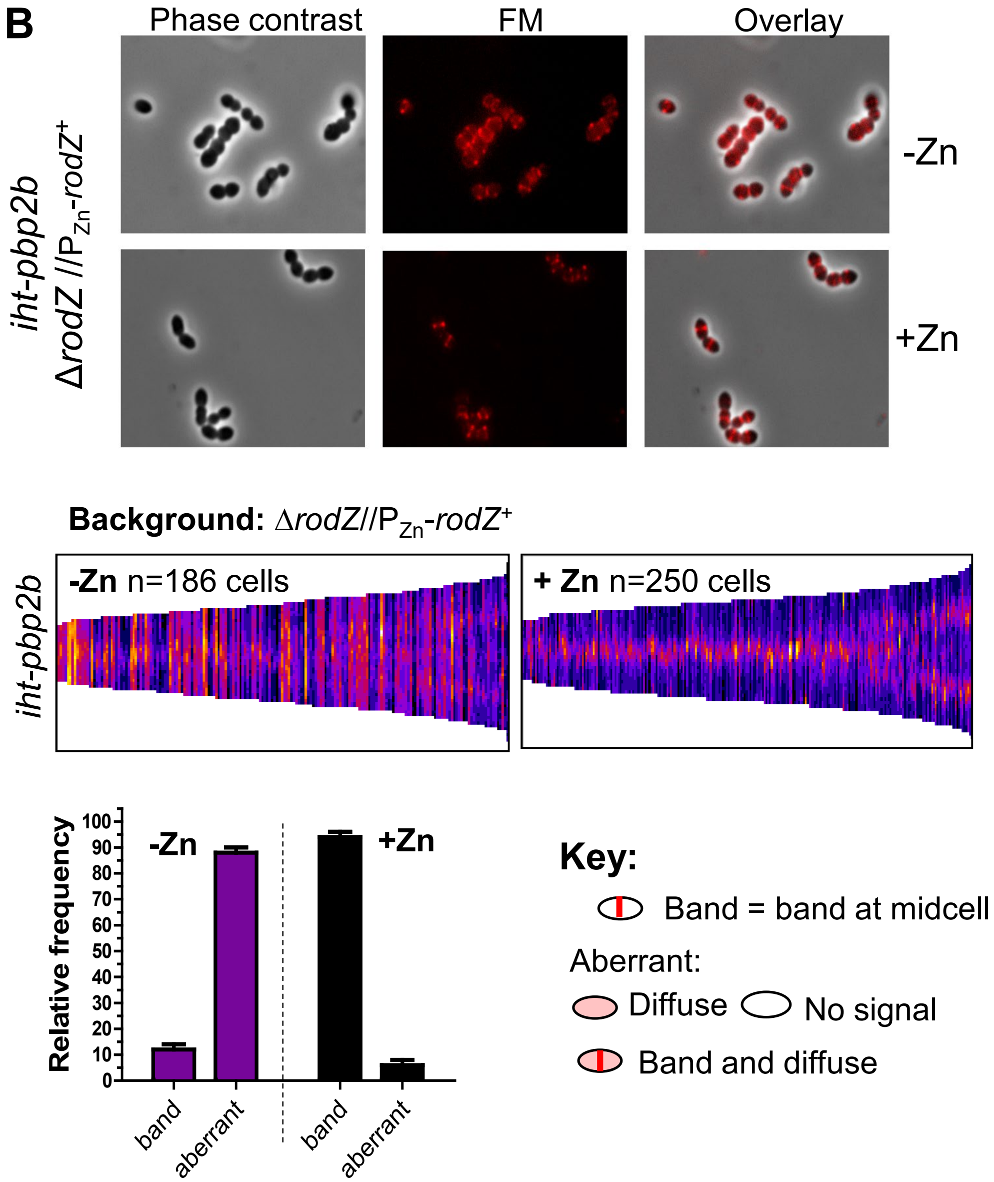


**Fig S22. (B)** bPBP2b mislocalizes upon RodZ depletion**.** IU16058 (*iht-pbp2b ΔrodZ*//P$_{Zn}$-*rodZ*$^{+}$) was grown with (complementation) or without inducer (depletion) Zn inducer (0.4 mM $ZnCl_2$ + 0.04 mM $MnSO_4$). Cells were harvested, and labeled with HT-TMR ligand at 4 h of growth. Top panels: representative micrographs showing phase contrast, FM and overlay images of iHT-PBP2b localization. Middle panels: demographs showing fluorescence intensity of iHT-PBP2b localization in the absence and presence of RodZ. n indicates the number of cells aligned in a given demograph. Bottom panels: bar graph displaying iHT-PBP2b localization patterns characterized from micrograph images. 100 cells from two independent experiments were considered for each strain and condition. Key illustrates criteria used to classify observed localization patterns as aberrant or WT status. This figure represents 3 independent biological replicates from which similar results were obtained. (Continued on next page)

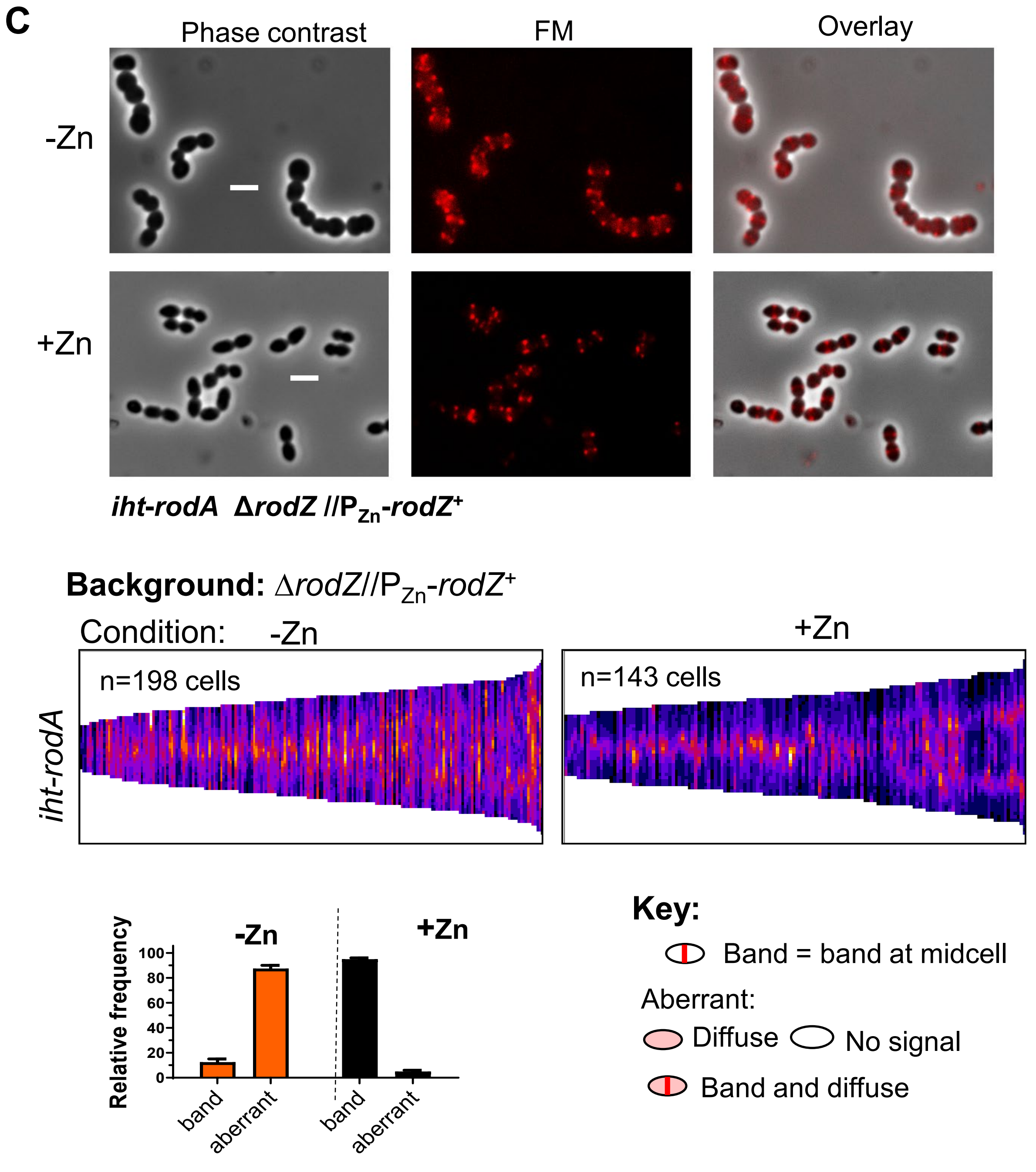


**Fig S22. (C)** RodA mislocalizes upon RodZ depletion. IU16060 (*iht-rodA ΔrodZ*//P$_{Zn}$-*rodZ*$^{+}$) was grown with or without inducer, labeled with TMR for 2D-fluorscence microscopy. Panel arrangements are the same as Fig. S21B. Scale bar = 2 µm. Representative figure from 3 independent biological replicates which gave similar results.

(Continued on next page)

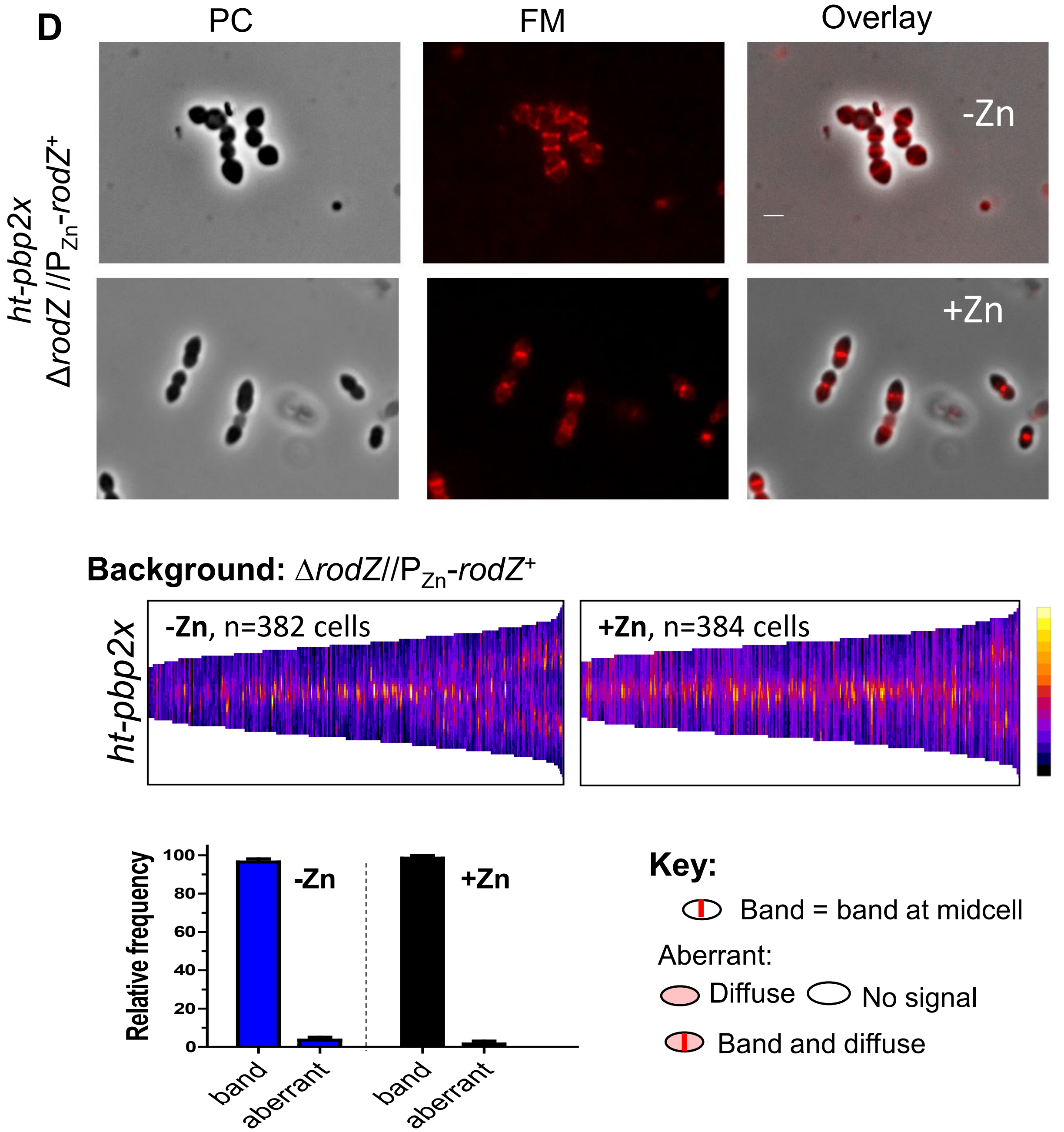


**Fig S22. (D)** bPBP2x maintains midcell localization upon RodZ depletion. IU16062 (*ht-pbp2x* Δ*rodZ*//P$_{Zn}$-*rodZ*$^+$) was grown with or without inducer and labeled with HT-TMR ligand for 2D-fluorescence microscopy. Scale bar on micrograph = 1 µm. Panel arrangements are the same as Fig. S21B. This figure represents 3 independent biological replicates from which similar results were obtained.

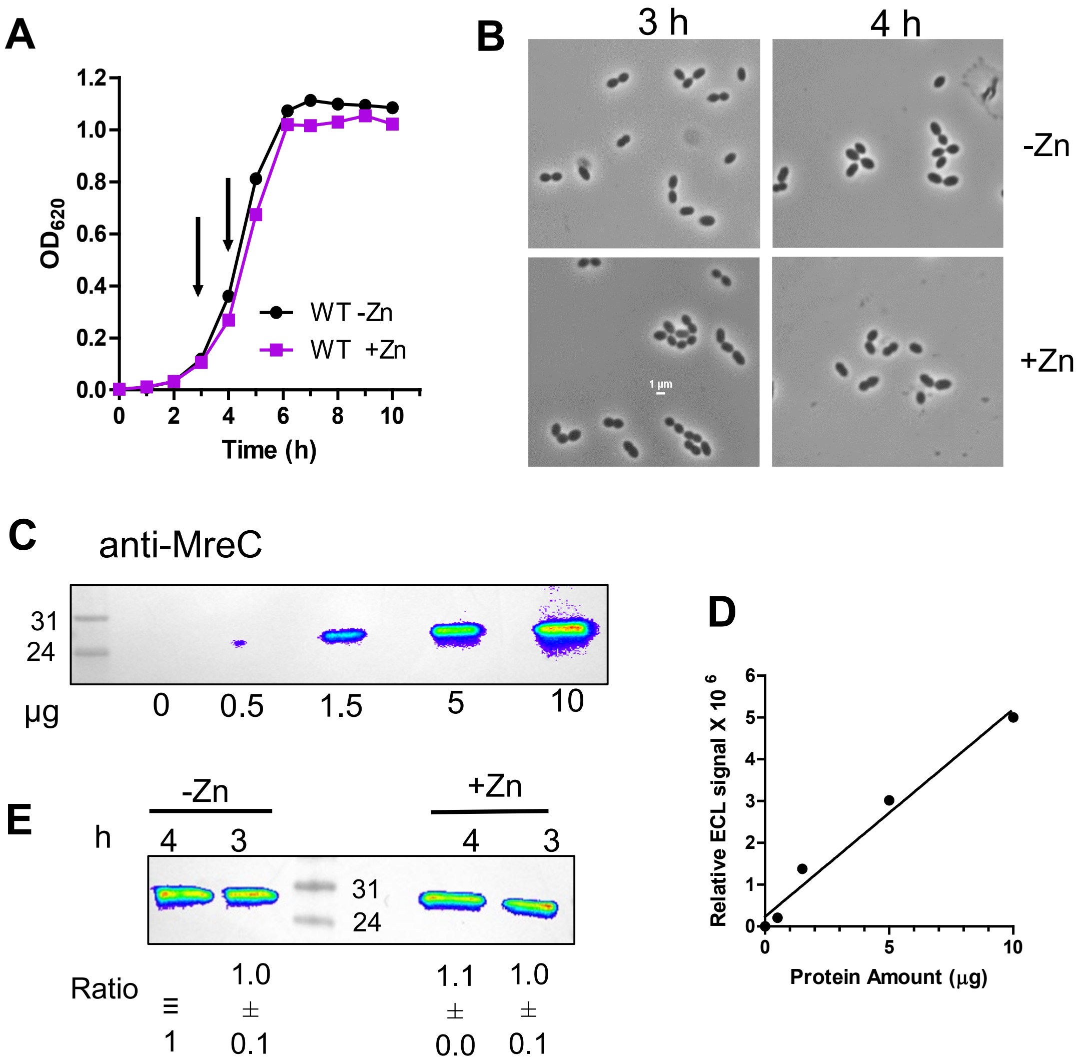


**Fig. S23. Zinc does not affect growth, cell morphology or MreC amount in WT cells. (A)** Representative growth curves of WT (IU1824) ± Zn inducer (0.4 mM $ZnCl_2$ + 0.04 mM $MnSO_4$) conditions. Arrows indicate times at which samples were harvested for western blot analysis. **(B)** Representative micrographs displaying IU1824 ± Zn at 3 and 4 h of growth. **(C)** and **(D)** Western blot showing direct relationship between µg protein loaded per lane (0, 0.5, 1.5, 5 or 10 µg) and signal intensities obtained with primary anti-MreC antibody and secondary HRP antibody labeling, and visualization with IVIS Living Image system. **(E)** Relative MreC amounts in IU1824 in the +Zn or -Zn conditions at 3 h or 4 h of growth. 6 µg of crude lysate was loaded in each lane. Relative quantitation of MreC (average ± SEM) was obtained from two independent biological replicates.

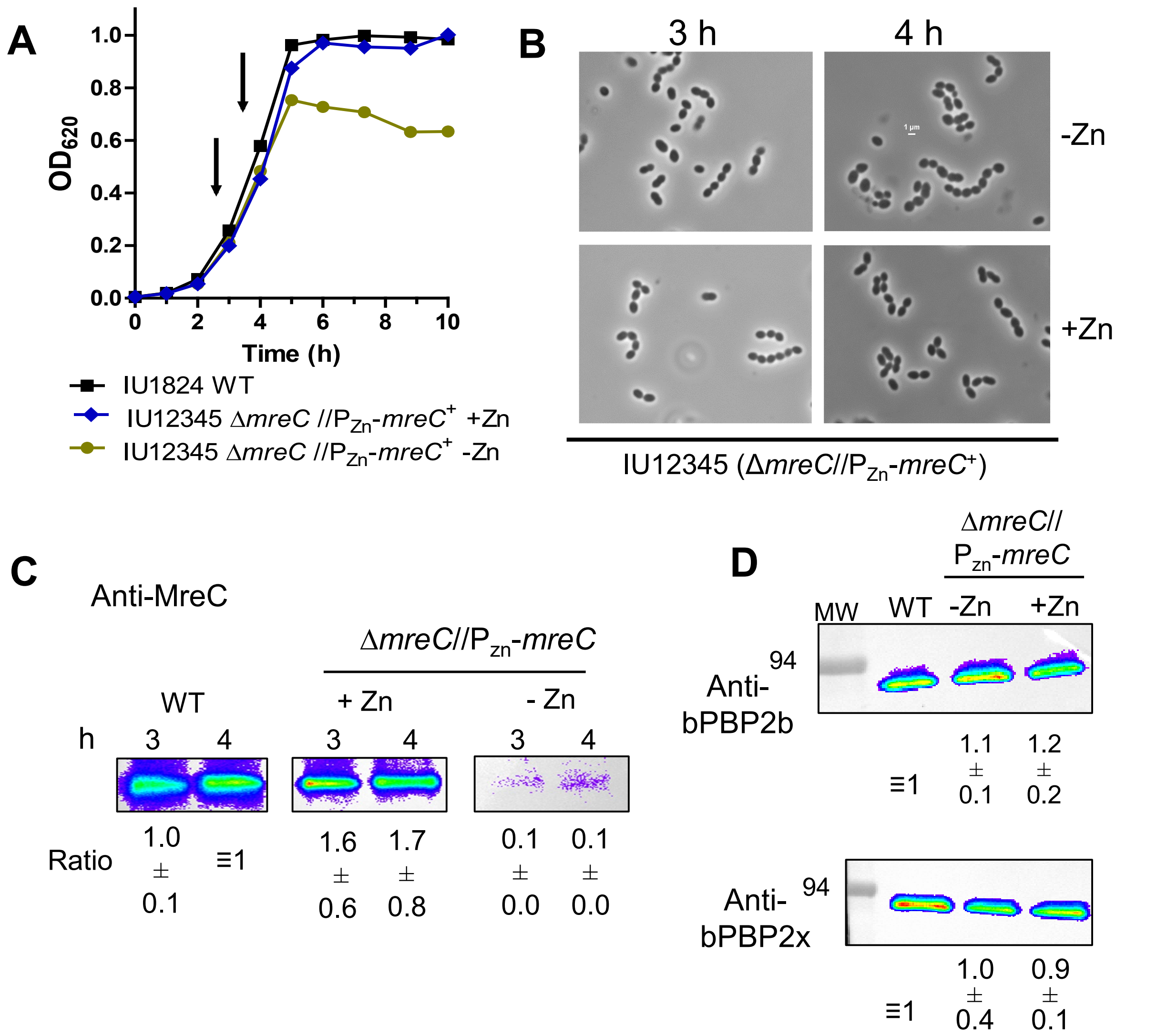


**Fig. S24. MreC protein amounts decrease to a nearly undetectable level, whereas bPBP2b and bPBP2x remain unchanged upon MreC depletion for 3 h. (A)** Representative growth curves of WT (IU1824) and depletion strain Δ*mreC*//P$_{Zn-}$*mreC*$^+$ (IU12345). IU1824 and IU12345 were grown overnight in BHI with or without Zn inducer (0.4 mM ZnCl + 0.04mM $MnSO_4$), respectively, and diluted into BHI with no Zn for IU1824, and into BHI with or without Zn for IU12345. Cultures were harvested at 3 or 4 h for western analysis (arrows). **(B)** Representative micrographs of IU1824 and IU12345 at 3h and 4h. Scale bar = 1 µm. **(C)** Western blot showing MreC expression from native chromosomal site in IU1824, or from the ectopic site in the presence or absence of inducer in IU12345 at 3h and 4h of growth. 6 µg of crude cell lysates were loaded in each lane. **(D)** bPBP2b and bPBP2x protein levels are not altered under MreC depletion condition. Protein samples were obtained from IU1824 (WT), or IU12345 grown in the presence or absence of inducer (+Zn or –Zn) for 4h. 3 µg of crude cell lysates were loaded in each lane. For C and D western blotting was carried out with primary antibodies to MreC, bPBP2b, or bPBP2x, secondary HRP antibody, and visualized with IVIS Living Image system. Ratios indicate protein amounts (average ± SEM) in IU12345 relative to WT from 2 independent biological replicates.

A

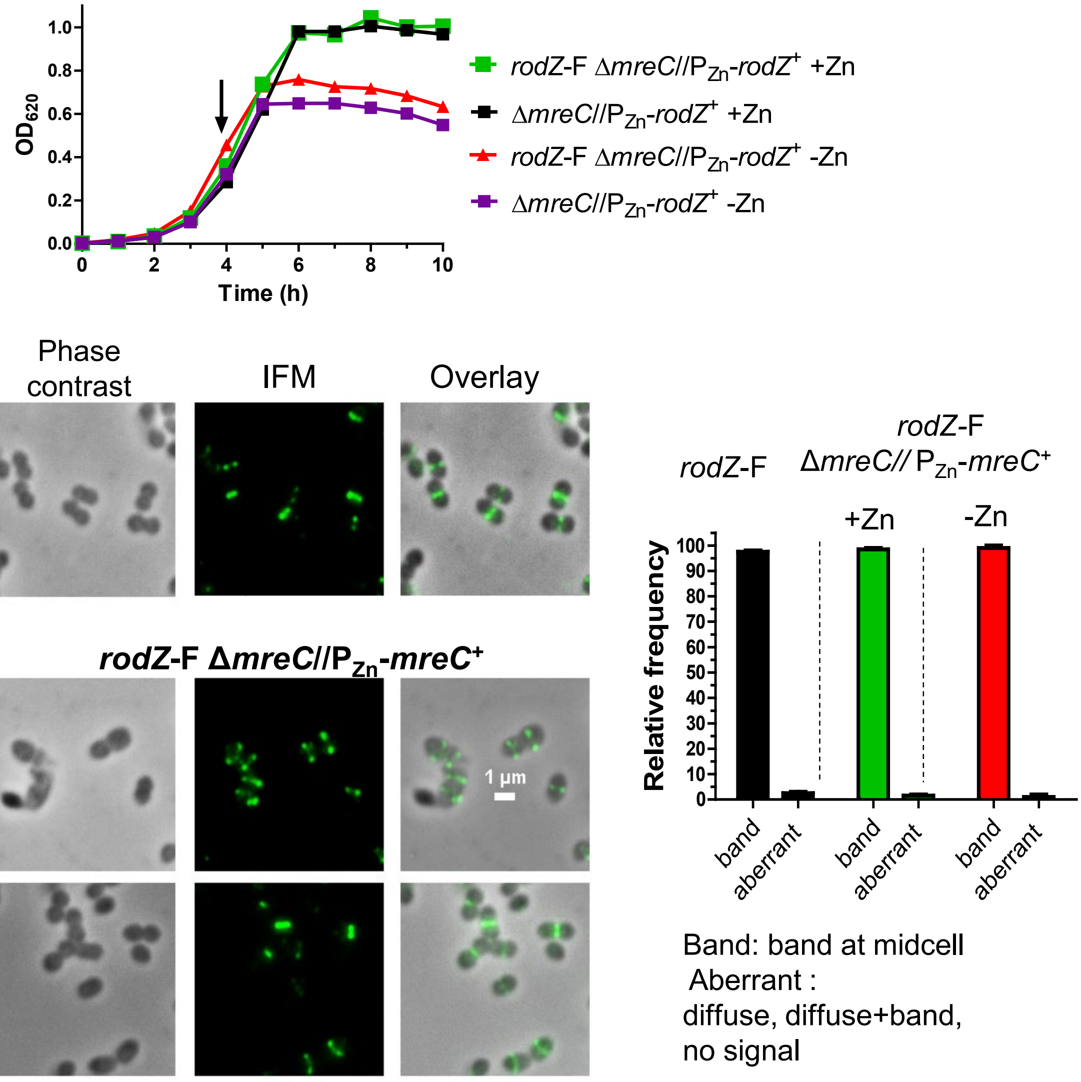


**Fig. S25. RodZ and aPBP1a maintain midcell localization, while bPBP2b mislocalizes upon MreC depletion. (A)** RodZ remains at midcell during MreC depletion. Representative growth curves of IU12345 (Δ*mreC*//P$_{Zn}$-*mreC*$^+$) and IU14598 (*rodZ*-F Δ*mreC*//P$_{Zn}$-*mreC*$^+$) grown in the presence or absence of Zn inducer (0.4 mM $ZnCl_2$ + 0.04 mM $MnSO_4$). Localization patterns are shown of RodZ during depletion of MreC for 4 h. 2D IFM of IU14594 (*rodZ*-F) and IU14598 were performed as outlined in *Experimental procedures.* Panels shown from left to right are: phase, FITC (antibody labeled FLAG-tagged RodZ), and overlay (phase + FITC). Quantification of the observed RodZ localization pattern is graphed for WT and Δ*mreC*//P$_{Zn}$-*mreC*$^+$ at 4h. For each sample and condition, 100 cells from two experiments were manually examined and classified. Growth curves and IFM images are representative of 3 independent biological experiments.(Continued on next page)

**B**

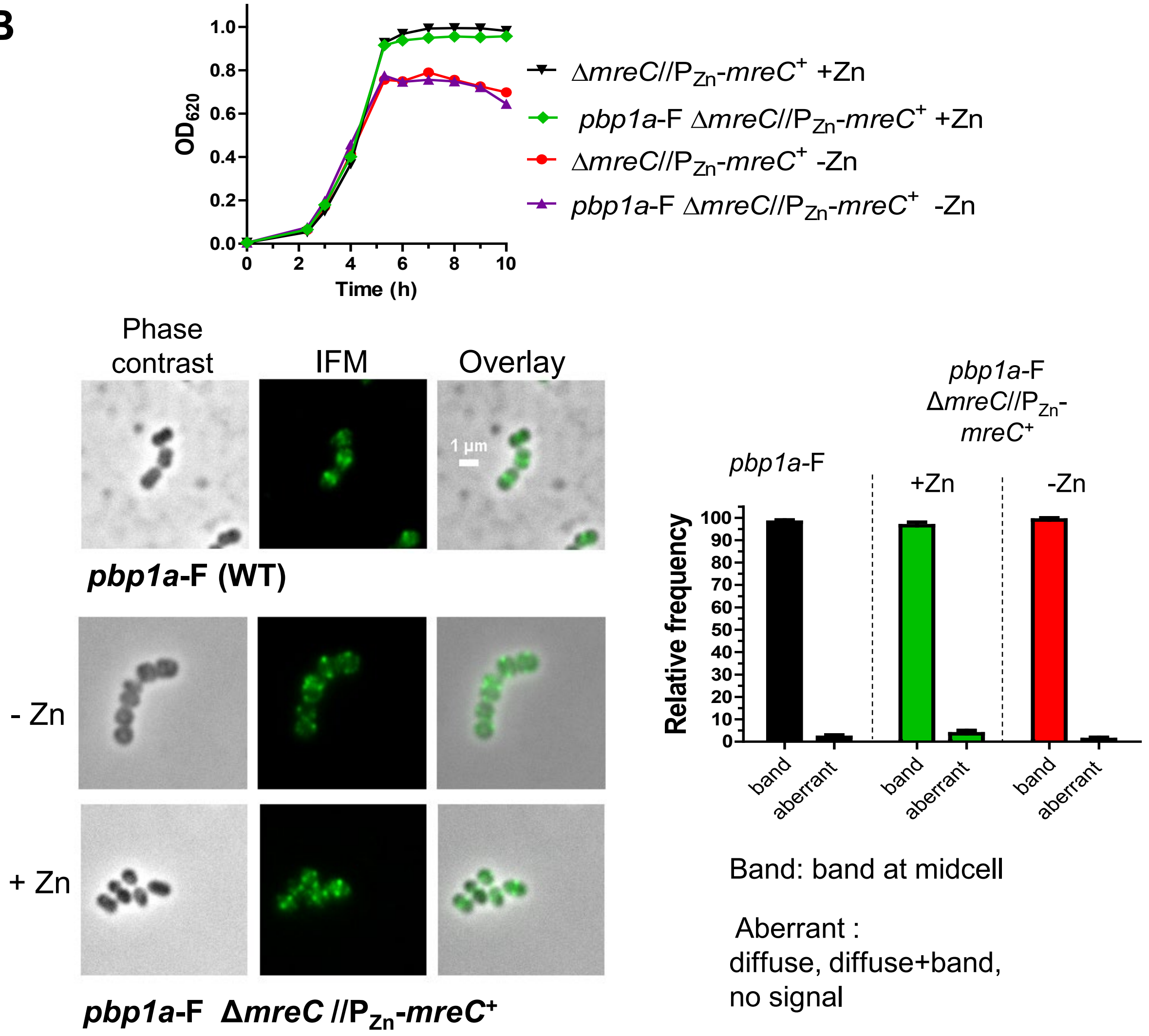


**Fig. S25**. **(B)** aPBP1a remains at midcell upon MreC depletion. (C) Representative growth curves of IU12345 (Δ*mreC*//$P_{Zn}$-*mreC*⁺) and IU15901 (*pbp1a*-F Δ*mreC*//$P_{Zn}$-*mreC*⁺) grown in the presence or absence of Zn inducer (0.4 mM $ZnCl_2$ + 0.04 mM $MnSO_4$). Arrow indicates the time at which samples were harvested and processed for IFM imaging. Localization patterns are shown of PBP1a-F during depletion of MreC at 4 h. 2D IFM of IU14494 (*pbp1a*-F) and IU15901 were performed as outlined in *Experimental procedures.* Panels shown from left to right are: phase, FITC (antibody labeled FLAG-tagged aPBP1a), and overlay (phase/FITC). Quantification of the observed PBP1a-F localization pattern is graphed for WT and Δ*mreC*//$P_{Zn}$-*mreC*⁺ at 4 h. For each sample and condition, 100 cells from two experiments were manually examined and classified. Growth curves and IFM images are representative of 2 independent biological experiments. (Continued on next page)

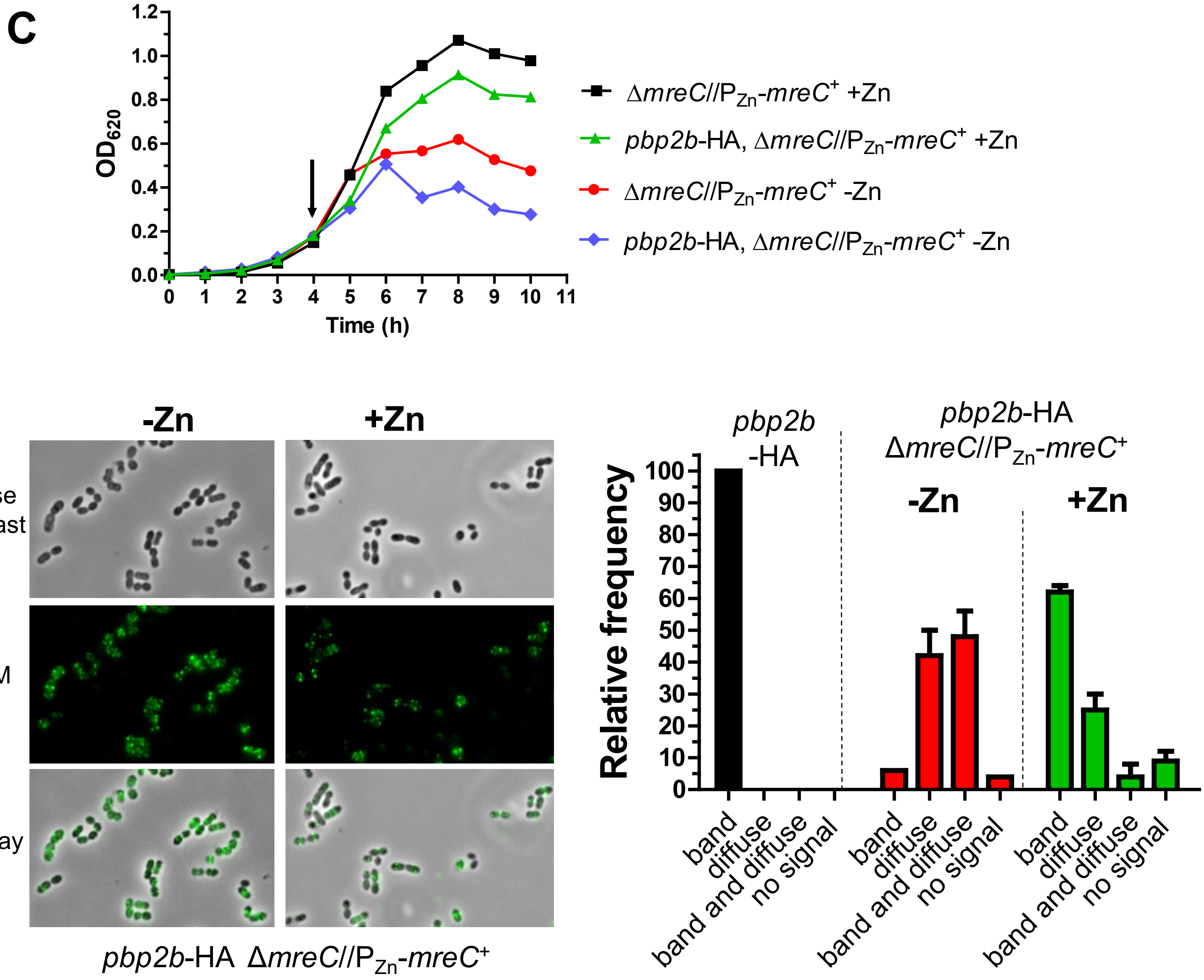


**Fig. S25. (C)** bPBP2b mislocalizes upon MreC depletion. Representative growth curves of IU12345 (Δ*mreC*//P$_{Zn}$-*mreC*$^{+}$) and IU14773 (*pbp2b*-HA Δ*mreC*//P$_{Zn}$-*mreC*$^{+}$). Arrow indicates the time at which samples were harvested and processed for IFM imaging. Representative IFM images are shown of IU14773 grown in the presence or absence of Zn inducer (0.4 mM $ZnCl_2$ + 0.04 mM $MnSO_4$). Quantification is shown for bPBP2b-HA localization in IU14455 (*pbp2b*-HA) and IU14773. 100 cells were categorized from two separate experiments as described in *Experimental procedures.*

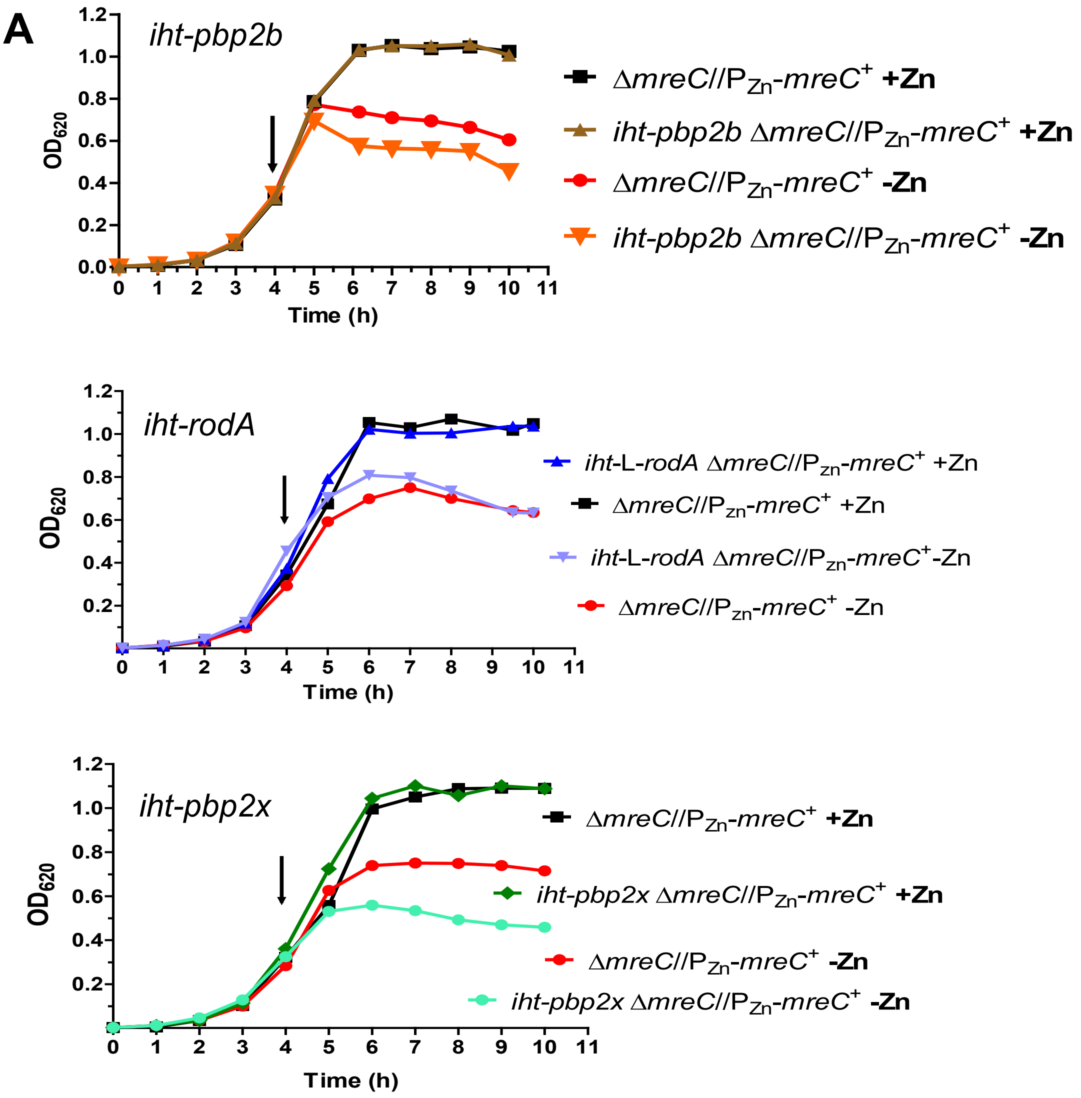


**Fig. S26**. **RodA and bPBP2b mislocalize upon MreC depletion, while bPBP2x localizes to midcell. (A)** Representative growth curves of strains expressing HT-fusion constructs in *mreC*-depletion background. Cells were harvested at 4 h of growth for labeling with HT-TMR. Strains used are IU12345 (Δ*mreC*//$P_{Zn}$-*mreC*$^+$), IU16281 (*iht-pbp2b* Δ*mreC*//$P_{Zn}$-*mreC*$^+$), IU16283 (*iht-rodA* Δ*mreC*//$P_{Zn}$-*mreC*$^+$), and IU16326 (*iht-pbp2x* Δ*mreC*//$P_{Zn}$-*mreC*$^+$). Growth curves were performed at least 2 times with similar results. (Continued on next page)

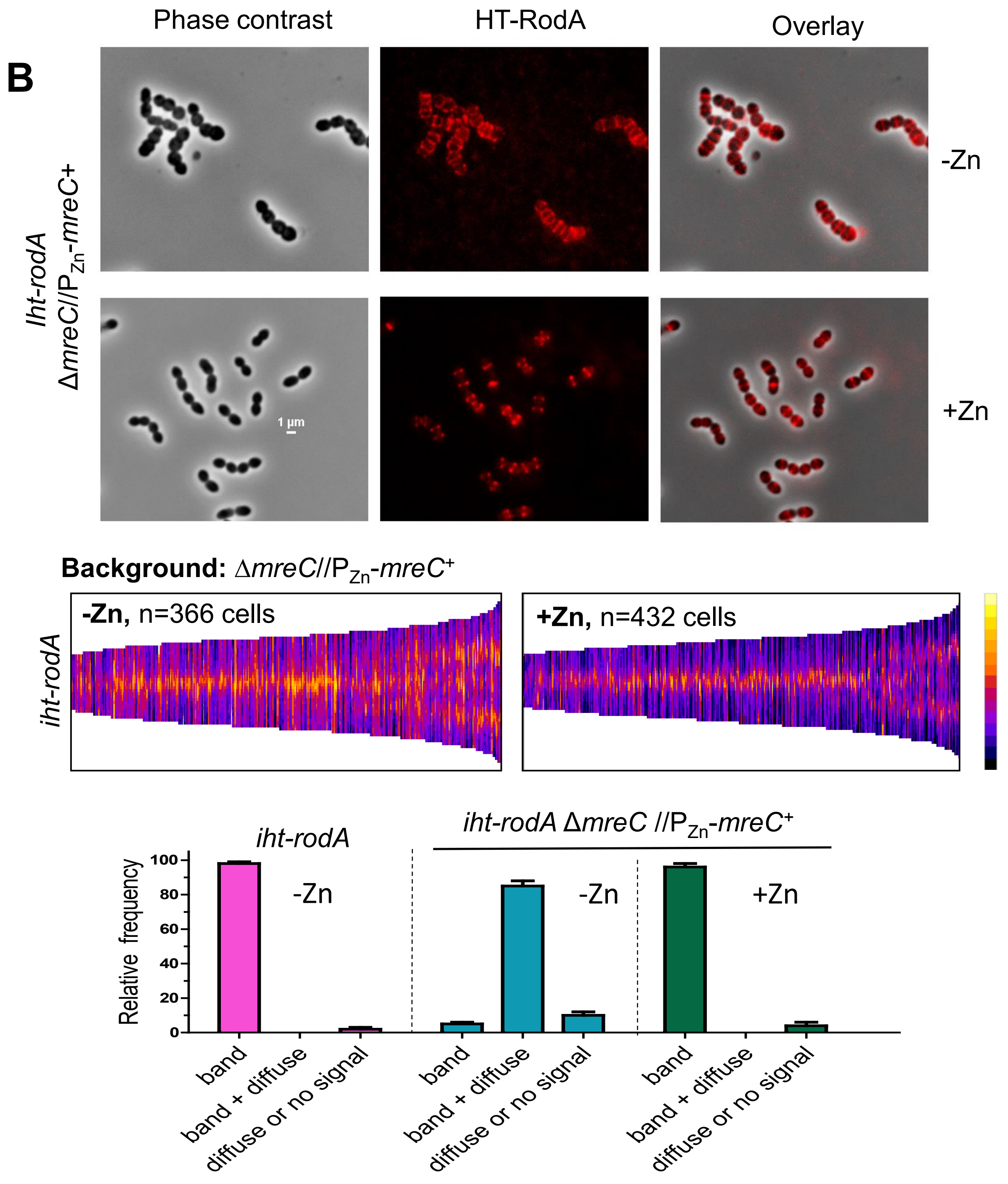


**Fig S26. (B)** RodA mislocalizes from midcell upon MreC depletion. IU16283 (*iht-rodA* Δ*mreC*//$P_{Zn}$-*mreC*$^+$) was grown with or without Zn inducer, and labeled with TMR for 2D-FM microscopy. Panels are arranged similarly to Figure 13 that shows bPBP2b mislocalization upon MreC depletion. Scale bar = 1 µm. This experiment was performed twice independently with similar results.(Continued on next page)

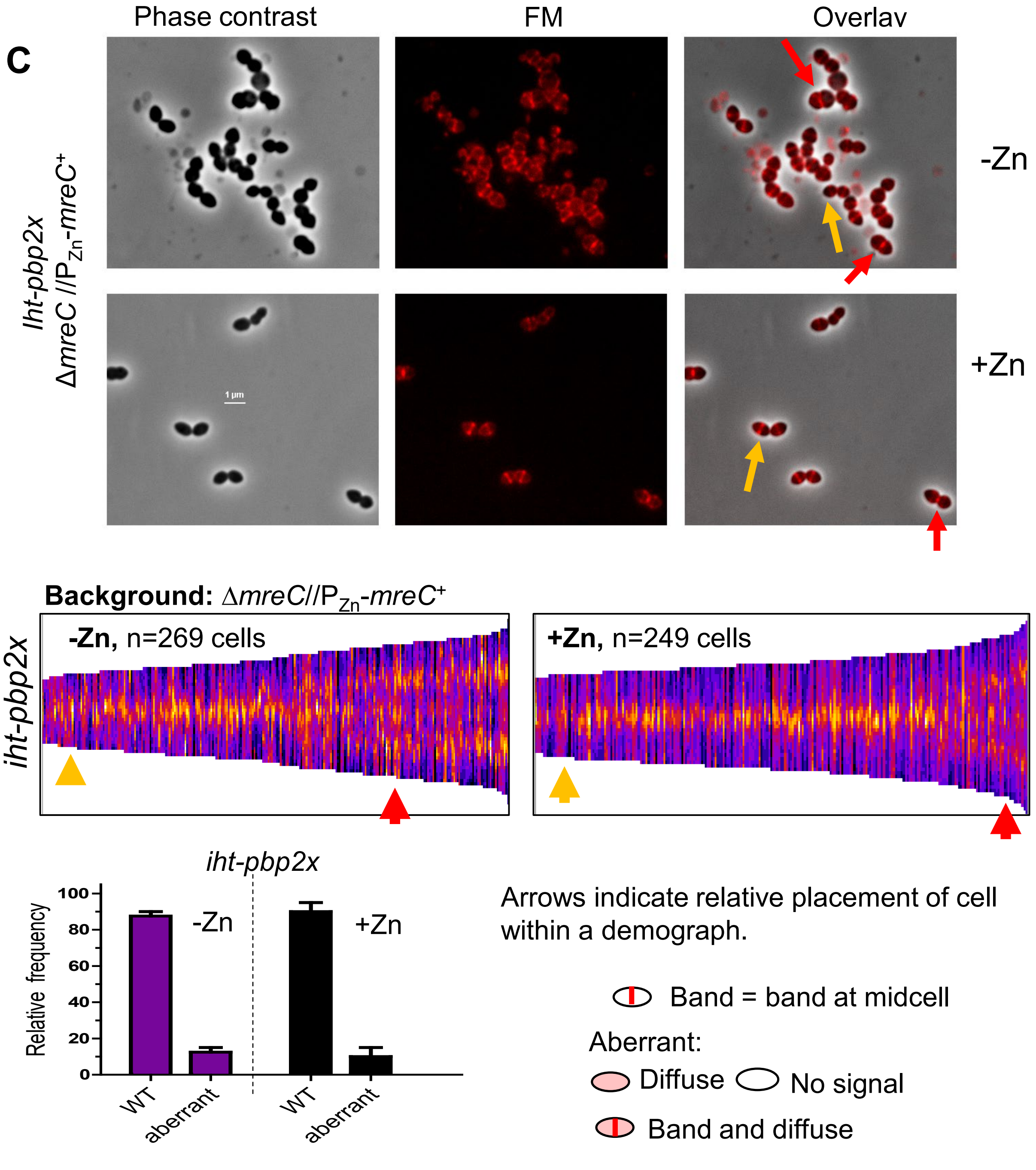


**Fig S26. (C)** bPBP2x localizes normally at midcell upon MreC depletion. IU16326 (*ht-pbp2x* Δ*mreC*//P$_{Zn}$-*mreC*$^+$) was grown with or without inducer and labeled with TMR for 2D-FM. Scale bar = 1 µm. Yellow and red arrows point to representative cells and indicate placement in the resulting demographs. This experiment was performed twice independently with similar results.

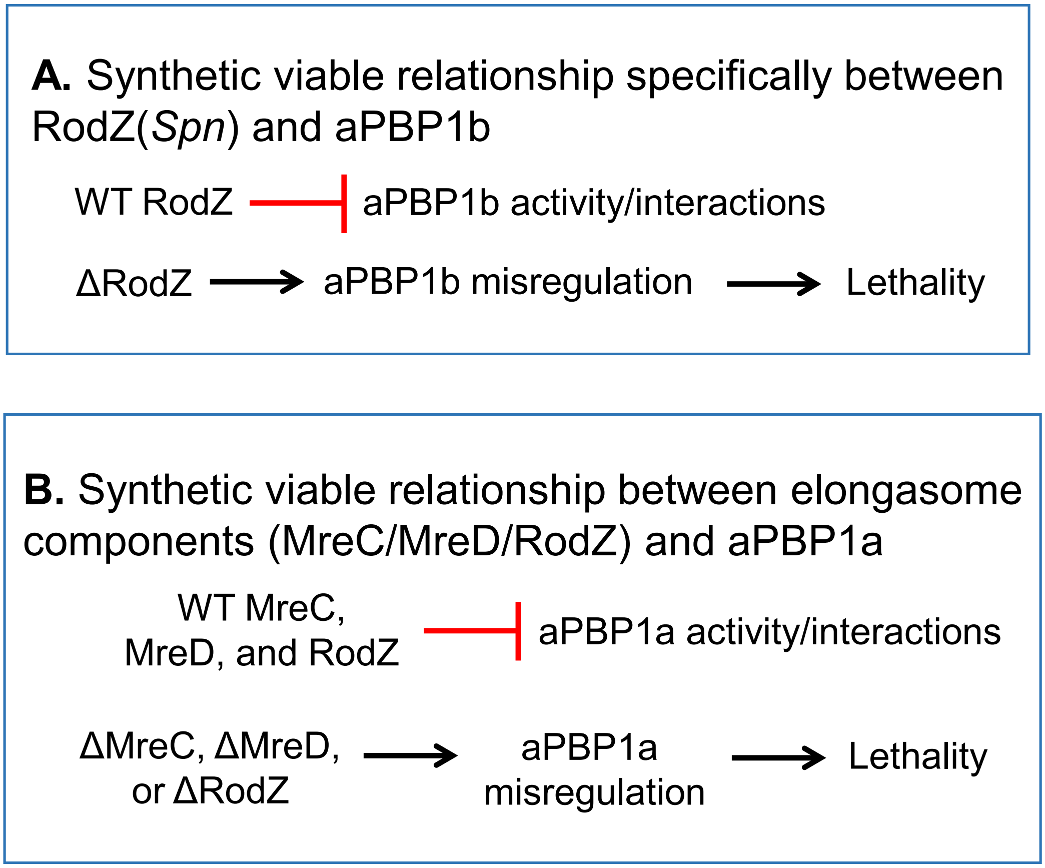


**Fig. S27.** Alternative model for synthetic-viable genetic relationships of aPBP1b and aPBP1a with members of the pPG elongasome of *S. pneumoniae*. **(A)** The absence of aPBP1b suppresses the essentiality of RodZ, but not that of MreC/D, whereas **(B)** the absence of aPBP1a suppresses the requirement for RodZ, MreC, or MreD. In this model, the synthetic-viable relationship results from negative regulation of aPBP1b or aPBP1a activity and/or interactions by the indicated members of the pneumococcal pPG elongasome in WT cells. In the absence of RodZ, aPBP1b and aPBP1a misregulation occurs and contributes to cell lethality. In the absence of MreC or MreD, aPBP1a, but not aPBP1b, misregulation occurs, contributing to cell lethality. The misregulation is alleviated by the absence of the aPBPs. See text for additional details and other models.
